# The *Glossina* Genome Cluster: Comparative Genomic Analysis of the Vectors of African Trypanosomes

**DOI:** 10.1101/531749

**Authors:** Geoffrey M Attardo, Adly M.M Abd-Alla, Alvaro Acosta-Serrano, James E Allen, Rosemary Bateta, Joshua B Benoit, Kostas Bourtzis, Jelle Caers, Guy Caljon, Mikkel B Christensen, David W Farrow, Markus Friedrich, Aurélie Hua-Van, Emily C Jennings, Denis M Larkin, Daniel Lawson, Michael J Lehane, Vasileios P Lenis, Ernesto Lowy-Gallego, Rosaline W. Macharia, Anna R Malacrida, Heather G Marco, Daniel Masiga, Gareth L Maslen, Irina Matetovici, Richard P Meisel, Irene Meki, Veronika Michalkova, Wolfgang J Miller, Patrick Minx, Paul O Mireji, Lino Ometto, Andrew G Parker, Rita Rio, Clair Rose, Andrew J Rosendale, Omar Rota-Stabelli, Grazia Savini, Liliane Schoofs, Francesca Scolari, Martin T Swain, Peter Takáč, Chad Tomlinson, George Tsiamis, Jan Van Den Abbeele, Aurelien Vigneron, Jingwen Wang, Wesley C Warren, Robert M Waterhouse, Matthew T Weirauch, Brian L Weiss, Richard K Wilson, Xin Zhao, Serap Aksoy

**Affiliations:** Institute of Biological, Environmental and Rural Sciences, Aberystwyth University, Aberystwyth, Ceredigion, United Kingdom; Department of Biochemistry, Biotechnology Research Institute - Kenya Agricultural and Livestock Research Organization, Kikuyu, Kenya; CAS Center for Influenza Research and Early-warning (CASCIRE), Chinese Academy of Sciences, Beijing, China; Center for Autoimmune Genomics and Etiology and Divisions of Biomedical Informatics and Developmental Biology, Cincinnati Children’s Hospital Medical Center, Cincinnati, Ohio, United States; Laboratoire Evolution, Genomes, Comportement, Ecologie, CNRS, IRD, Univ. Paris-Sud, Université Paris-Saclay, Gif-sur-Yvette, France; VectorBase, European Molecular Biology Laboratory, European Bioinformatics Institute (EMBL-EBI), Cambridge, Cambridgeshire, United Kingdom; Department of Biological Sciences, Florida International University, Miami, Florida, United States; Department of Sustainable Ecosystems and Bioresources, Research and Innovation Centre, Fondazione Edmund Mach, San Michele all'Adige (TN), Italy; School of Life Sciences, Fudan University, Shanghai, China; Department of Life Sciences, Imperial College London, London, United Kingdom; Biomedical Sciences, Institute of Tropical Medicine, Antwerp, Belgium; Molecular Biology and Bioinformatics Unit, International Center for Insect Physiology and Ecology, Nairobi, Kenya; Insect Pest Control Laboratory, Joint FAO/IAEA Division of Nuclear Techniques in Food & Agriculture, Vienna, Vienna, Austria; Centre for Geographic Medicine Research, Coast, Kenya Medical Research Institute, Kilifi, Kenya; Department of Biology - Functional Genomics and Proteomics Group, KU Leuven, Leuven, Belgium; Department of Vector Biology, Liverpool School of Tropical Medicine, Liverpool, Merseyside, United Kingdom; Department of Cell and Developmental Biology, Medical University of Vienna, Vienna, Austria; Department of Biology, Mount St. Joseph University, Cincinnati, Ohio, United States; Department of Comparative Biomedical Sciences, Royal Veterinary College, London, United Kingdom; Institute of Zoology, Slovak Academy of Sciences, Bratislava, Slovakia; Laboratory of Microbiology, Parasitology and Hygiene, University of Antwerp, Antwerp, Belgium; Department of Entomology and Nematology, University of California, Davis, Davis, California, United States; Department of Biological Sciences, University of Cape Town, Rondebosch, South Africa; Department of Biological Sciences, University of Cincinnati, Cincinnati, Ohio, United States; Department of Biology and Biochemistry, University of Houston, Houston, Texas, United States; Department of Ecology & Evolution, and Swiss Institute of Bioinformatics, University of Lausanne, Lausanne, Switzerland; Centre for Biotechnology and Bioinformatics, University of Nairobi, Nairobi, Kenya; Department of Environmental and Natural Resources Management, University of Patras, Agrinio, Etoloakarnania, Greece; Department of Biology and Biotechnology, University of Pavia, Pavia, Italy; Schools of Medicine and Dentistry, University of Plymouth, Plymouth, United Kingdom; Department of Animal Systematics, Ústav zoológie SAV; Scientica, Ltd., Bratislava, Slovakia; McDonnell Genome Institute, Washington University School of Medicine, St. Louis, Missouri, United States; Department of Biological Sciences, Wayne State University, Detroit, Michigan, United States; Department of Biology, West Virginia University, Morgantown, West Virginia, United States; Department of Epidemiology of Microbial Diseases, Yale School of Public Health, New Haven, Connecticut, United States; Bond Life Sciences Center, University of Missouri, Columbia, Missouri, United States

**Keywords:** Tsetse, trypanosomiasis, hematophagy, lactation, disease, neglected, symbiosis

## Abstract

**Background:** Tsetse flies (*Glossina sp.*) are the sole vectors of human and animal trypanosomiasis throughout sub-Saharan Africa. Tsetse are distinguished from other Diptera by unique adaptations, including lactation and the birthing of live young (obligate viviparity), a vertebrate blood specific diet by both sexes and obligate bacterial symbiosis. This work describes comparative analysis of six *Glossina* genomes representing three sub-genera: *Morsitans* (*G. morsitans morsitans (G.m. morsitans)*, *G. pallidipes*, *G. austeni*), *Palpalis* (*G. palpalis*, *G. fuscipes*) and *Fusca* (G. *brevipalpis*) which represent different habitats, host preferences and vectorial capacity.

**Results:** Genomic analyses validate established evolutionary relationships and sub-genera. Syntenic analysis of *Glossina* relative to *Drosophila melanogaster* shows reduced structural conservation across the sex-linked X chromosome. Sex linked scaffolds show increased rates of female specific gene expression and lower evolutionary rates relative to autosome associated genes. Tsetse specific genes are enriched in protease, odorant binding and helicase activities.

Lactation associated genes are conserved across all *Glossina* species while male seminal proteins are rapidly evolving. Olfactory and gustatory genes are reduced across the genus relative to other characterized insects. Vision associated Rhodopsin genes show conservation of motion detection/tracking functions and significant variance in the Rhodopsin detecting colors in the blue wavelength ranges.

**Conclusions:** Expanded genomic discoveries reveal the genetics underlying *Glossina* biology and provide a rich body of knowledge for basic science and disease control. They also provide insight into the evolutionary biology underlying novel adaptations and are relevant to applied aspects of vector control such as trap design and discovery of novel pest and disease control strategies.

## Background

Flies in the genus *Glossina* (tsetse flies) are vectors of African trypanosomes, which are of great medical and economic importance in Africa. Sleeping sickness (Human African Trypanosomiasis or HAT) is caused by two distinct subspecies of the African trypanosomes transmitted by tsetse. In East and Southern Africa, *Trypanosoma brucei rhodesiense* causes the acute *Rhodesiense* form of the disease, while in Central and West Africa *T*. *b*. *gambiense* causes the chronic *Gambiense* form of the disease, which comprises about 95% of all reported HAT cases. Devastating epidemics in the 20^th^ century resulted in hundreds of thousands of deaths in sub-Saharan Africa [1], but more effective diagnostics now indicate that data concerning sleeping sickness deaths are subject to gross errors due to under-reporting [2]. With hindsight, it is thus reasonable to infer that millions died from sleeping sickness during the colonial period. Loss of interest and funding for control programs within the endemic countries resulted in a steep rise in incidence after the post-independence period of the 1960s. In an ambitious campaign to control the transmission of Trypanosomiasis in Africa, multiple groups came together in a public/private partnership. These include the WHO, multiple non-governmental organizations, Sanofi Aventis and Bayer. The public sector groups developed and implemented multi-country control strategies and the companies donated the drugs required for treatment of the disease. The campaign reduced the global incidence of *Gambiense* HAT to <3,000 cases in 2015 [3]. Based on the success of the control campaign there are now plans to eliminate *Gambiense* HAT as a public health problem by 2030 [4]. In contrast, control of *Rhodesiense* HAT has been more complex as disease transmission involves domestic animals, which serve as reservoirs for the parasite. Hence elimination of the *Rhodesiense* disease will require treatment or elimination of domestic reservoirs, and/or reduction of tsetse vector populations. These strategies play a key part while medical interventions are used largely for humanitarian purposes. In addition to the public health impact of HAT, Animal African Trypanosomiasis (AAT or Nagana) limits the availability of meat and milk products in large regions of Africa. It also excludes effective cattle rearing from ten million square kilometers of Africa [5] with wide implications for land use, i.e. constraints on mixed agriculture and lack of animal labor for ploughing [6]. Economic losses in cattle production are estimated at 1-1.2 billion dollars US and total agricultural losses caused by AAT are estimated at 4.75 billion dollars US per year [7, 8].

Achieving disease control in the mammalian host has been difficult given the lack of vaccines. This is due to the process of antigenic variation the parasite displays in its host. Hence, accurate diagnosis of the parasite and staging of the disease are important. This is of particular importance due to the high toxicity of current drugs available for treatment of late-stage disease although introduction of a simpler and shorter nifurtimox and eflornithine combination therapy (NECT)[9] and discovery of new oral drugs, such as fexinidazole [10] and acoziborole, are exciting developments. Although powerful molecular diagnostics have been developed in research settings, few have yet to reach the patients or national control programs [11]. Further complicating control efforts, trypanosomes are showing resistance to available drugs for treatment [12, 13]. While vector control is essential for zoonotic *Rhodesiense* HAT, it has not played a major role in *Gambiense* HAT as it was considered too expensive and difficult to deploy in the resource poor settings of HAT foci. However, modelling, historical investigations and practical interventions demonstrate the significant role that vector control can play in the control of *Gambiense* HAT [14–16], especially given the possibility of long-term carriage of trypanosomes in both human and animal reservoirs [17, 18]. The African Union has made removal of trypanosomiasis via tsetse fly control a key priority for the continent [19]. Within the *Glossinidae*, 33 extant taxa are described from 22 species in 4 subgenera. The first three sub-genera *Austenina Townsend*, *Nemorhina Robineau-Desvoidy* and *Glossina Wiedemann* correspond to the *Fusca*, *Palpalis*, and *Morsitans* species groups, respectively [20].

The fourth subgenus *Machadomia* was established in 1987 to incorporate *G. austeni*. The relationship of *G. austeni* Newstead with respect to the *Palpalis* and *Morsitans* complex flies remains controversial [21]. While molecular taxonomy shows that *Palpalis*-and *Morsitans*-species groups are monophyletic, the *Fusca* species group emerges as sister group to all remaining Glossinidae. *Morsitans* group taxa are adapted to drier habitats relative to the other two subgenera [22]. *Palpalis* group flies tend to occur in riverine and lacustrine habitats. *Fusca* group flies largely inhabit moist forests of West Africa. The host-specificity of the different species groups vary, with the *Palpalis* group flies displaying strong anthrophilicity while the others are more zoophilic in preference. The principal vectors of HAT include *G. palpalis* s.l., *G. fuscipes* and *G. m. morsitans* s.l. The riverine habitats of *Palpalis* group flies and their adaptability to peridomestic environments along with human blood meal preferences make them excellent vectors for HAT. Other species belonging to the *Morsitans* group (such as *G. pallidipes*) can also transmit human disease, but principally play an important role in AAT transmission. In particular, *G. pallidipes* has a wide distribution and a devastating effect in East Africa. Also, of interest is *G. brevipalpis*, an ancestral tsetse species within the Fusca species complex. This species exhibits poor vectorial capacity with *T. brucei* relative to *G. m. morsitans* in laboratory infection expriments using colonized fly lines [23]. Comparison of the susceptibility of *G. brevipalpis* to *Trypanosoma congolense* (a species that acts as a major causative agent of AAT) also showed it has a much lower rate of infection relative to *Glossina austeni* [24].

To expand the genetic/genomic knowledge and develop new and/or improved vector control tools a consortium, the International Glossina Genome Initative (IGGI), was established in early 2004 to sequence the *G. m. morsitans* genome [25]. In 2014 the first tsetse fly genome from the *Glossina m. morsitans* species was produced [26]. This project facilitated the use of modern techniques such as transcriptomics and enabled functional investigations at the genomic level into tsetse’s viviparous reproductive physiology, obligate symbiosis, trypanosome transmission biology, olfactory physiology and the role of saliva in parasite transmission. The species *Glossina m. morsitans* was chosen for the first genome discovery effort as it is relatively easy to maintain under laboratory conditions and many physiological studies had been based on this species. To study the genetics underlying tsetse species-specific traits, such as host preference and vector competence, we have now assembled five additional representative genomes from the species complexes of *Glossina*: Morsitans (*G. m. morsitans*, G. *pallidipes*,), Morsitans/Machadomia (*G. austeni*), Palpalis *(G. palpalis, G. fuscipes)* and Fusca (*G. brevipalpis*). These species represent flies with differences in geographical localization, ecological preferences, host specificity and vectorial capacity (Summarized in Figure 1). Here we report on the evolution and genetics underlying this genus by comparison of their genomic architecture and predicted protein-coding sequences and highlight some of the genetic differences that hold clues to the differing biology between these species.

**Figure 1:**
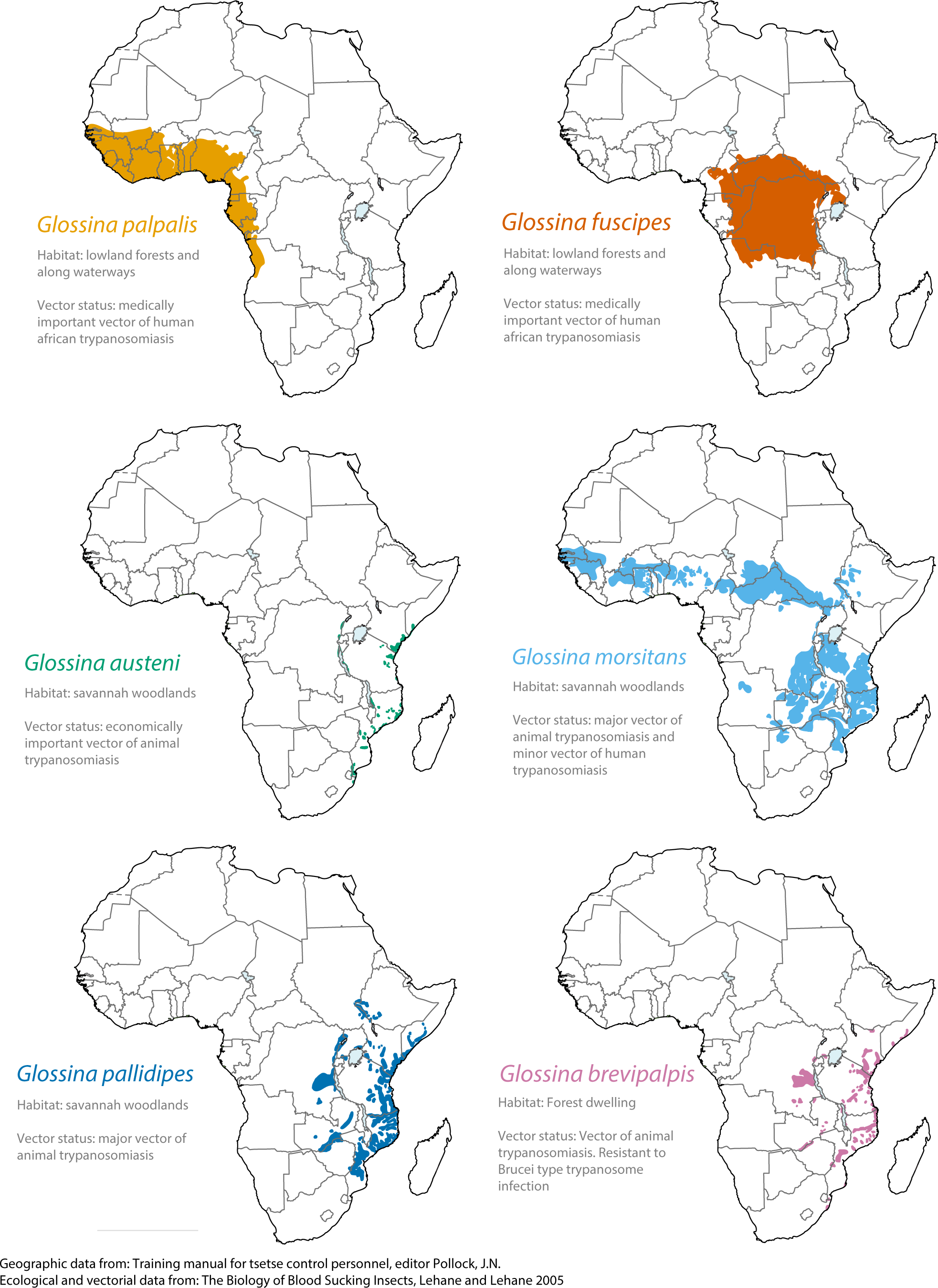
Geographic distribution, ecology and vectorial capacity of sequenced *Glossina* species. Visual representation of the geographic distribution of the sequenced *Glossina* species across the African continent. Ecological preferences and vectorial capacities are described for each associated group.

## Results and Discussion

### Multiple genetic comparisons confirm *Glossina* phylogenetic relationships and the inclusion of *G. austeni* as a member of the *Morsitans* sub-Genus

Sequence similarity between the genomes was analyzed using whole genome nucleotide alignments of supercontigs and predicted coding sequences from the five new *Glossina* genomes as well as those from the *Musca domestica* genome using *G. m. morsitans* as a reference (Figure 2A). The results indicate that *G. pallidipes* and *G. austeni* are most similar at the sequence level to *G. m. morsitans*. This is followed by the species in the *Palpalis* sub-genus (*G. fuscipes* and *G. palpalis*). The remaining species (*G. brevipalpis*) shows the least sequence conservation relative to *G. m. morsitans* followed by the outgroup species *M. domestica*. The lower sequence similarity between *G. brevipalpis* and the other tsetse species reinforces its status as a distant relative to the *Morsitans* and *Palpalis* sub-genera.

**Figure 2:**
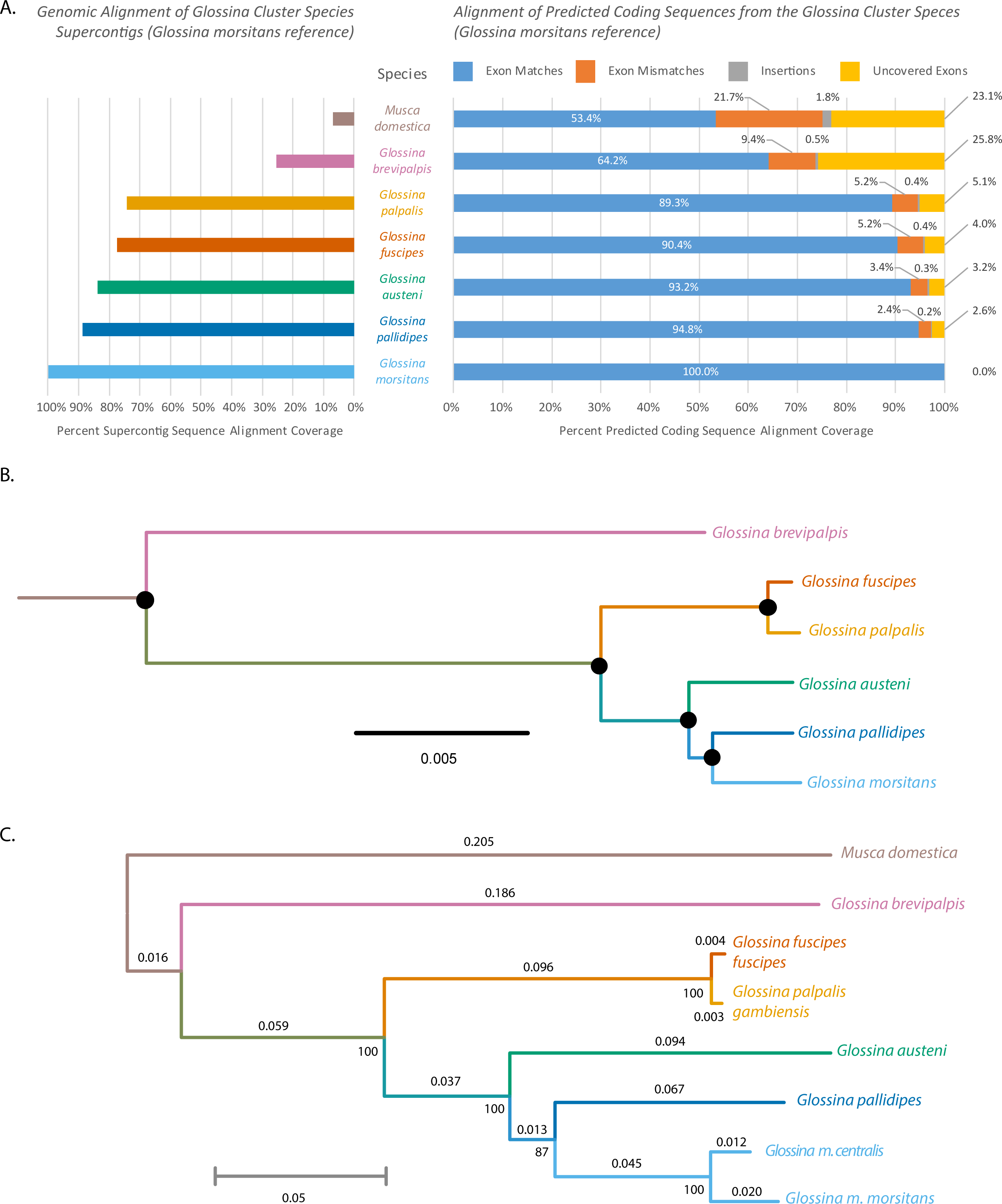
*Glossina* whole genome alignment, phylogenetic analysis of orthologous protein-coding nuclear genes and phylogenetic analysis of mitochondrial sequences. A. Analysis of whole genome and protein-coding sequence alignment. The left graph reflects the percentage of total genomic sequence aligning to the *G. m. morsitans* reference. The right side of the graph represents alignment of all predicted coding sequences from the genomes with coloration representing matches, mismatches, insertions and uncovered exons. B. Phylogenic tree from conserved protein-coding sequences. Black dots at nodes indicate full support from Maximum likelihood (Raxml), Bayesian (Phylobayes), and coalescent-aware (Astral) analyses. Raxml and Phylobayes analyses are based on an amino acid dataset of 117,782 positions from 286 genes from 12 species. Astral analyses is based on a 1125 nucleotide dataset of 478,617 positions from the 6 *Glossina* (full trees are in Supplemental Figure 1A-C). C. Molecular phylogeny derived from whole mitochondrial genome sequences. The analysis was performed using the maximum likelihood method with MEGA 6.0.

Alignment of the predicted coding sequences produced a similar result to that observed in the whole genome alignment in terms of similarity to *G. m. morsitans* (Figure 2A). Of interest is that more than 25% of the *G. m. morsitans* exon sequences were not align-able with *G. brevipalpis*, indicating that they were either lost, have diverged beyond alignability or were in an unsequenced region in *G. brevipalpis*. In addition, *G. brevipalpis* has on average ~5000 fewer predicted protein-coding genes than the other species. Given the low GC content of the *G. brevipalpis* sequenced genome it is possible that some of the regions containing these sequences lie within heterochromatin. The difficulties associated with sequencing heterochromatic regions may have excluded these regions from our analysis; however, it also implies that if these protein coding genes are indeed present they are located in a region of the genome with low transcriptional activity.

We inferred the phylogeny and divergence times of *Glossina* using a concatenated alignment of 286 single-copy gene orthologs (478,000 nucleotide positions) universal to *Glossina* (Figure 2B). The tree recovered from this analysis has support from both Maximum Likelihood and Bayesian analyses, using respectively homogeneous and heterogeneous models of replacement. A coalescent-aware analysis further returned full support, indicating a speciation process characterized by clear lineage sorting (Supplemental Figure 1). These results suggest an allopatric speciation process characterized by a small founder population size followed by little to no introgression among newly formed species.

Furthermore, we assembled complete mitochondrial (mtDNA) genome sequences for each species as well as *Glossina morsitans centralis* as references for use in distinguishing samples at the species, sub-species or haplotype levels. All the mtDNA genomes encode large (16S rRNA) and small (12S rRNA) rRNAs, 22 tRNAs and 13 protein-coding genes. Phylogenetic analysis of the resulting sequences using the Maximum Likelihood method resulted in a tree with congruent topology to that produced by analysis of the concatenated nuclear gene alignment (Figure 2C). A comparative analysis of the mtDNA sequences identified variable marker regions with which to identify different tsetse species via traditional sequencing and/or high-resolution melt analysis (HRM) (Supplemental Figure 2). Analysis of the amplicons from this region using HRM facilitated the discrimination of these products based on their composition, length and GC content. Use of HRM on these variable regions successfully resolved differences between test samples consisting of different tsetse species as well as individuals with different haplotypes or from different populations (Supplemental Figure 3). This method provides a rapid, cost effective and relatively low-tech way of identifying differences in field caught tsetse for the purposes of population genetics and measurement of population diversity.

**Figure 3:**
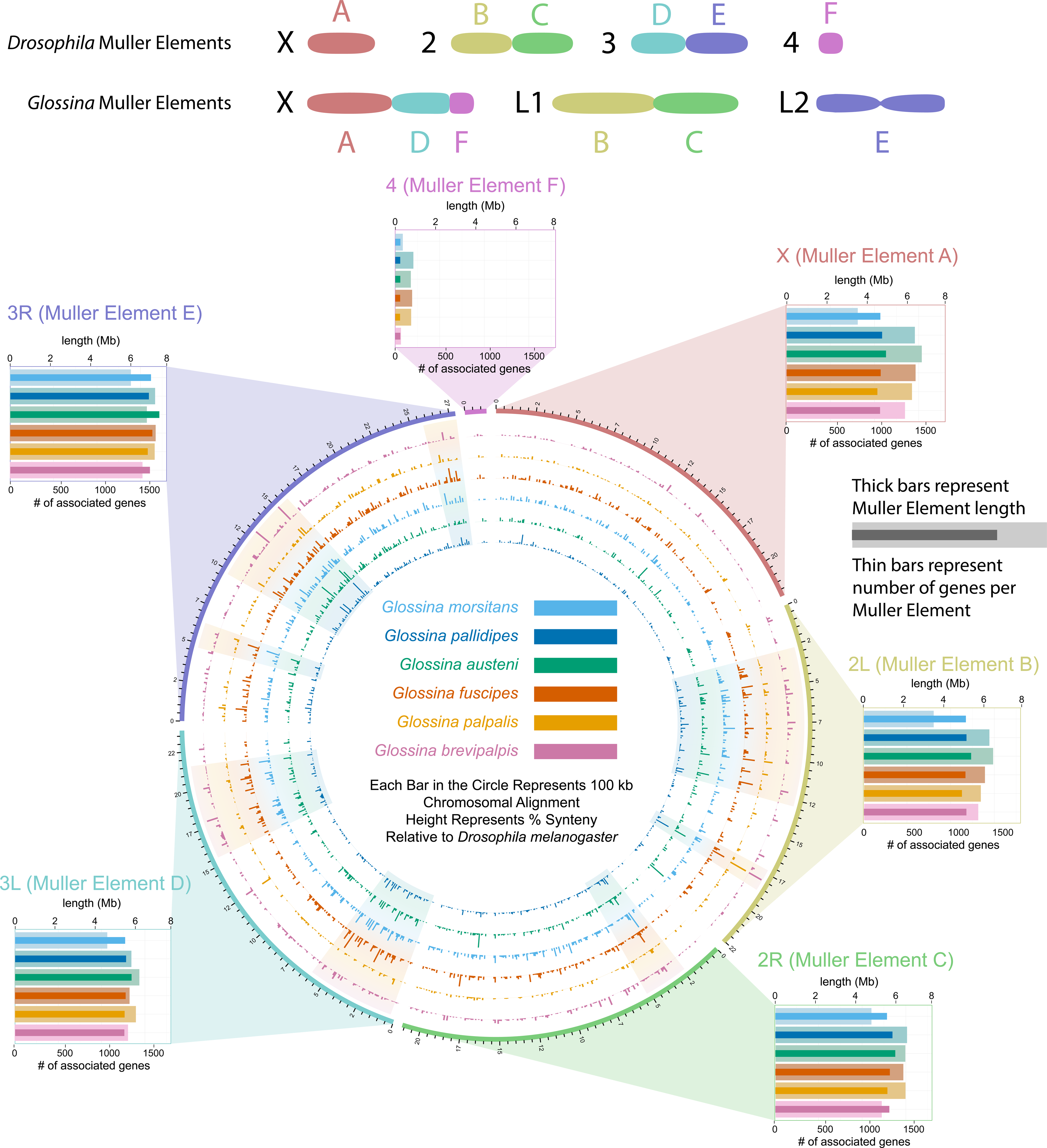
Visualization of syntenic block analysis data and predicted Muller Element sizes. Level of syntenic conservation between tsetse scaffolds and *Drosophila* chromosomal structures (Muller Elements). The color-coded concentric circles consisting of bars represent the percent of syntenic conservation of orthologous protein-coding gene sequences between the *Glossina* genomic scaffolds and *Drosophila* Muller elements. Each bar represents 100 kb of aligned sequence and bar heights represent the percent of syntenic conservation. The graphs on the periphery of the circle illustrate the combined predicted length and number of genes associated with the Muller elements for each tsetse species. The thin darkly colored bars represent the number of 1:1 orthologs between each *Glossina* species and *D. melanogaster*. The thicker lightly colored bands represent the predicted length of each Muller element for each species. This was calculated as the sum of the lengths of all scaffolds mapped to those Muller elements.

The trees derived from the nuclear and mitochondrial phylogenetic analyses agree with previously published phylogenies for tsetse [27–29] and the species delineate into groups representing the defined *Fusca*, *Palpalis* and *Morsitans* sub-genera.

A contentious issue within the taxonomy of *Glossina* is the placement of *G. austeni* within the *Machadomia* sub-Genus. Comparative anatomical analysis of male genitalia places *G. austeni* within the *morsitans* sub-Genus. However, female *G. austeni* genitalia bear anatomical similarities to members of the *Fusca* sub-genus. In addition, *G. austeni’s* habitat preferences and some external morphology resemble those of the *palpalis* sub-Genus [28]. Recent molecular evidence suggests that *G. austeni* are closer to the *morsitans* sub-genus [27, 29]. The data generated via the three discrete analyses described above all support the hypothesis that *G. austeni* is a member of the *Morsitans* sub-genus rather than the *Palpalis* sub-genus and belongs as a member of the *Morsitans* group rather than its own discrete sub-Genus.

### Comparative analysis of *Glossina* with *Drosophila* reveals reduced synteny and female specific gene expression on X-linked scaffolds

The scaffolds in each *Glossina spp.* genome assembly were assigned to chromosomal arms based on orthology and relative position to protein-coding sequences in the *D. melanogaster* genome (*Drosophila*) [30]. The *Glossina* and *Drosophila* genomes contain six chromosome arms (Muller elements A-F) [31–33]. We assigned between 31-52% of annotated genes in each species to a Muller element, which we used to assign >96% of scaffolds to Muller elements in each species (Figure 3 and Supplemental table 1). From these results, we inferred the relative size of each Muller element in each species by counting the number of annotated genes assigned to each element and calculating the cumulative length of all assembled scaffolds assigned to each element. Using either measure, we find that element E is the largest and element F is the shortest in all species, consistent with observations in *Drosophila* [34].

Mapping of the *Glossina* scaffolds to the *Drosophila* Muller elements reveals differing levels of conservation of synteny (homologous genomic regions with maintained orders and orientations) across these six-species relative to *Drosophila*. In *G. m. morsitans*, the X chromosome is composed of Muller elements A, D, and F as opposed to the *Drosophila* X which only contains A and sometimes D [33], and all other *Glossina* species besides *G. brevipalpis* have the same karyotype [35]. We therefore assume that the same elements are X-linked in the other *Glossina* species (apart from *G. brevipalpis*). This analysis reveals that scaffolds mapping to *Drosophila* Muller element A show a reduced overall level of syntenic conservation relative to the other Muller elements while the scaffolds mapping to *Drosophila* Muller element D (part of the *Glossina* X chromosome, but not the *D. melanogaster* X) retain more regions of synteny conservation. We hypothesize that the lower syntenic conservation on element A reflects a higher rate of rearrangement because it has been X-linked for more time (both in the *Drosophila* and *Glossina* lineages) than element D (only in *Glossina*) and rearrangement rates are higher on the X chromosome (element A) in *Drosophila* [34]

To examine the relationship between gene expression and DNA sequence evolution, we compared gene expression levels between the X chromosome and autosomes using sex specific RNA-seq libraries derived from whole males, whole non-lactating females and whole lactating females for all the *Glossina* species apart from *G. pallidipes*. Consistent with previous results from *G. m. morsitans* [33], the ratio of female:male expression is greater on the X chromosome than autosomes across species (Supplemental Figure 4). In addition, there is a deficiency of genes with male-biased expression (up-regulated in males relative to females) on the X-linked elements in all species (Supplemental Figure 5). Reduced levels of male-biased gene expression have also been observed in mosquitoes and is a conserved feature of the *Anopheles* genus [36]. The X chromosome is hemizygous in males, which exposes recessive mutations to natural selection and can accelerate the rate of adaptive substitutions and facilitate the purging of deleterious mutations on the X chromosome [37, 38]. Using dN/dS values for annotated genes, we fail to find any evidence for this faster-X effect across the entire phylogeny or along any individual lineages (Supplemental Figure 6). The faster-X effect is expected to be greatest for genes with male-biased expression because they are under selection in males [37], but we find no evidence for faster-X evolution of male-biased genes in any of the *Glossina* species. In contrast, there is some evidence for “slower-X” evolution amongst female-biased genes (Supplemental Figure 7), suggesting that purifying selection is more effective at purging deleterious mutations on the X chromosome [39]. Genes with female-biased expression tend to be broadly expressed [40], suggesting that pleiotropic constraints on female-biased genes increase the magnitude of purifying selection and produce the observed slower-X effect [41].

**Figure 4:**
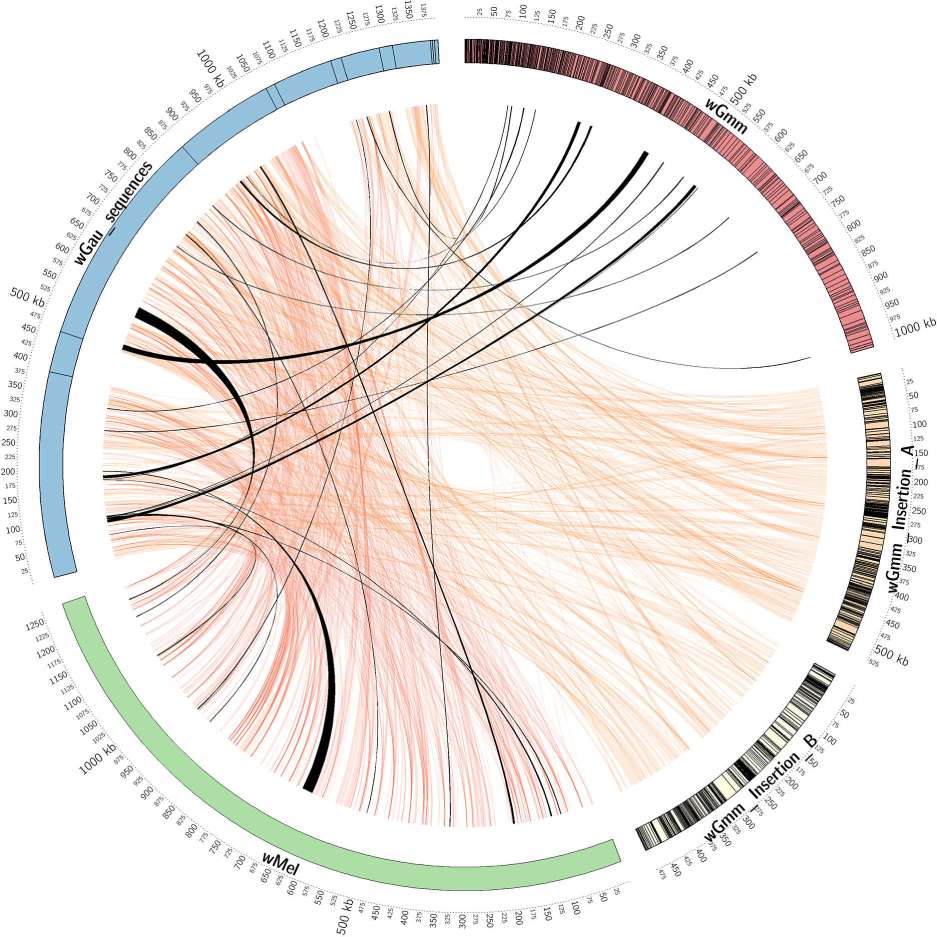
Homology map of the *Wolbachia* derived cytoplasmic and horizontal transfer derived nuclear sequences. Circular map of the *G. austeni Wolbachia* horizontal transfer derived genomic sequences (*wGau* - blue), the *D. melanogaster Wolbachia* cytoplasmic genome sequence (*wMel* - green), the *G. m. morsitans Wolbachia* cytoplasmic genome sequence (*wGmm* - red), and the *Wolbachia* derived chromosomal insertions A & B from *G. m. morsitans* (*wGmm* Insertion A and Insertion B yellow and light yellow respectively). The outermost circle represents the scale in kbp. Contigs for the *wGau* sequences, *wGmm* and the chromosomal insertions A & B in *G. m. morsitans* are represented as boxes. Regions of homology between the *G. austeni* insertions and the other sequences are represented by orange ribbons. Black ribbons represent syntenic regions between the *wGau* insertions and the cytoplasmic genomes of *wGmm* and *wMel*.

**Figure 5:**
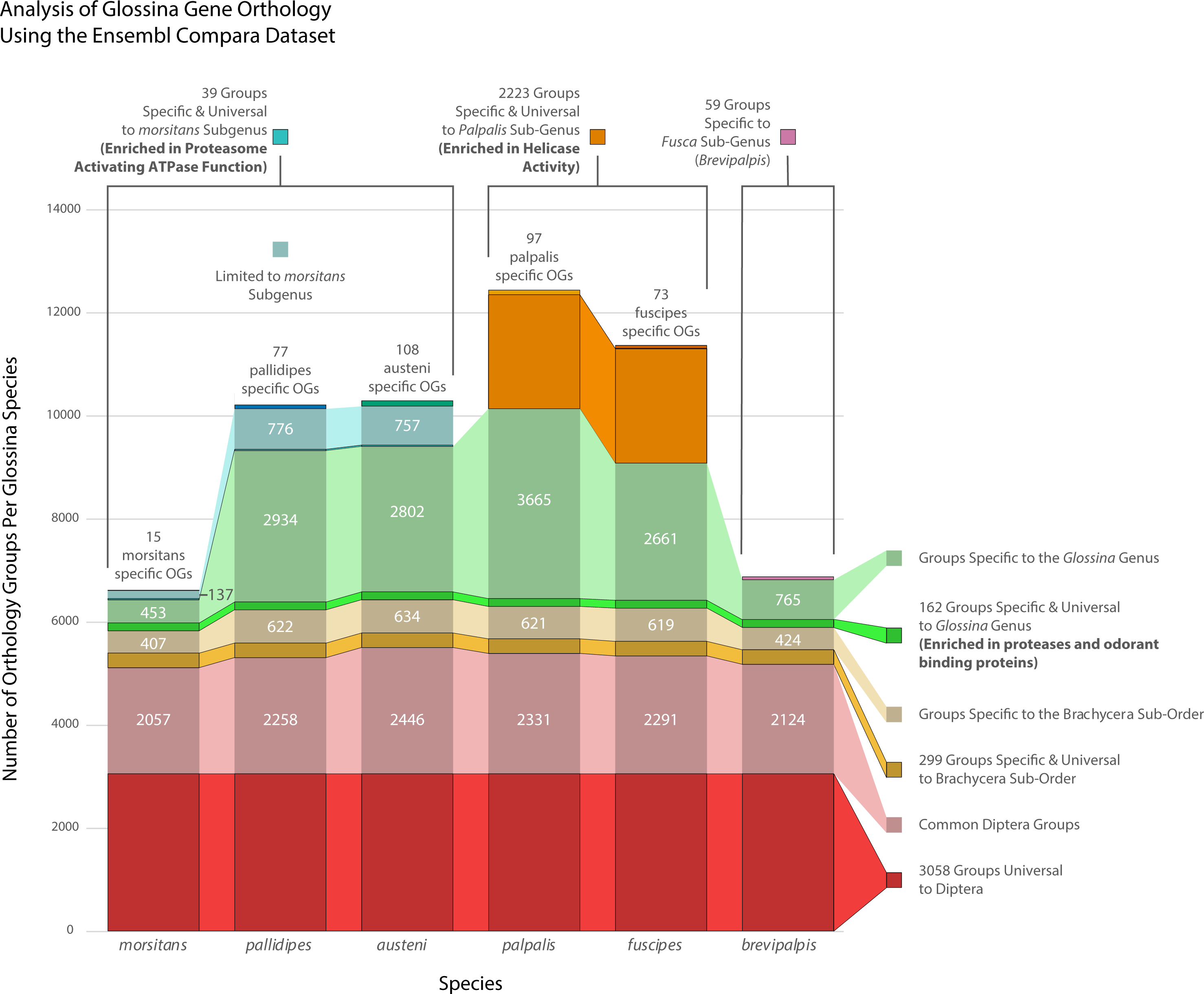
Constituent analysis of *Glossina* associated gene orthology groups. Visualization of relative constitution of orthology groups containing *Glossina* gene sequences. Combined bar heights represent the combined orthogroups associated with each *Glossina* species. The bars are color-coded to reflect the level of phylogenetic representation of clusters of orthogroups at the order, sub-order, genus, sub-genus and species. Saturated bars represent orthology groups specific and universal to a phylogenetic level. Desaturated bars represent orthogroups specific to a phylogenetic level but lack universal representation across all included species. Gene ontology analysis of specific and universal groups can be found in Table 3.

**Figure 6:**
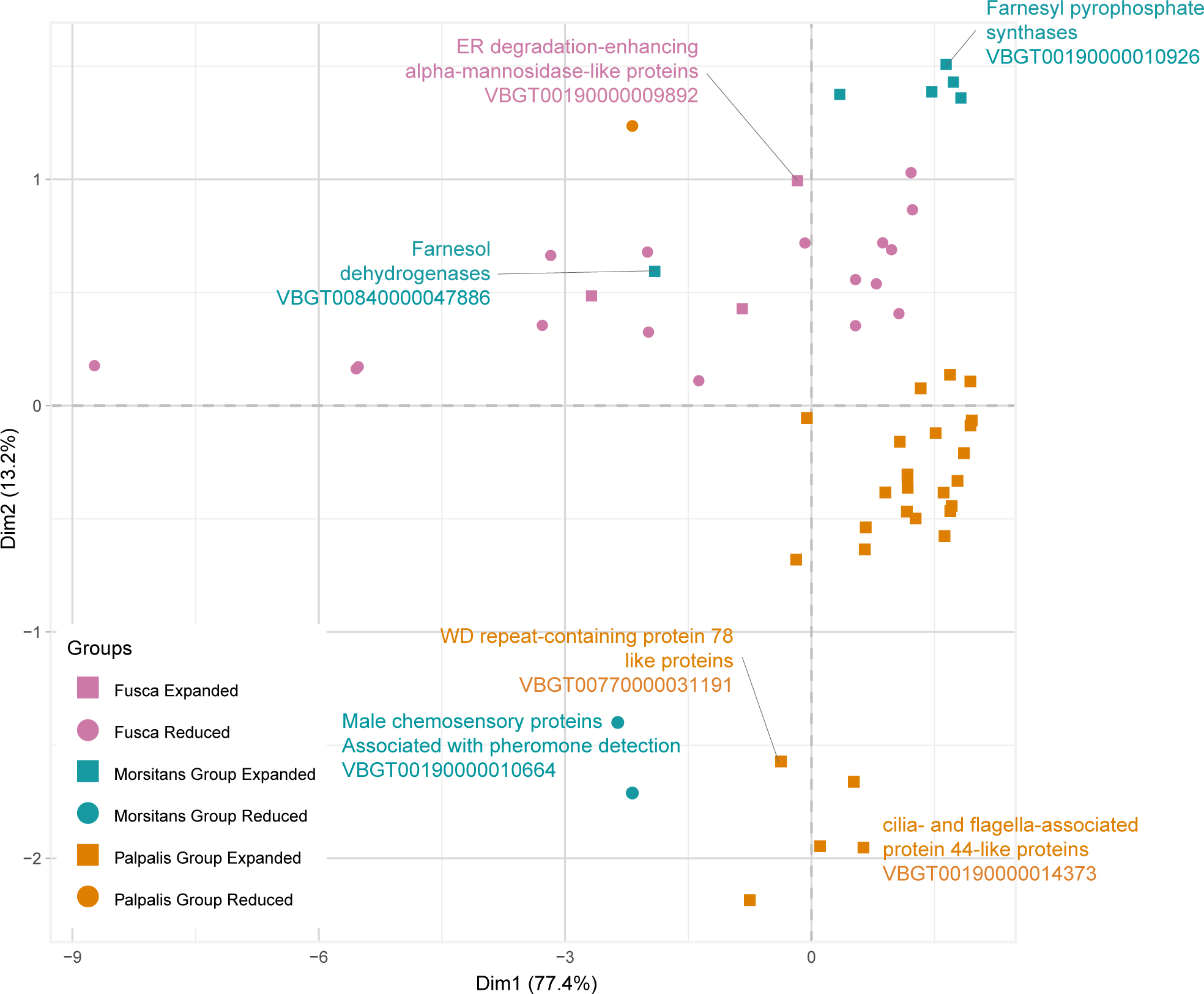
Sub-genus specific gene family expansions/retractions. Principal component analysis-based clustering of gene orthology groups showing significant differences in the number of representative sequences between the six *Glossina* species. Orthology groups included have sub-genus specific expansions/contractions as determined by CAFE test (P-value < 0.05). Detailed information regarding the functional associations of the unlabeled groups is provided in Supplemental Figures 10+11 and in Supplemental Table 4.

**Figure 7:**
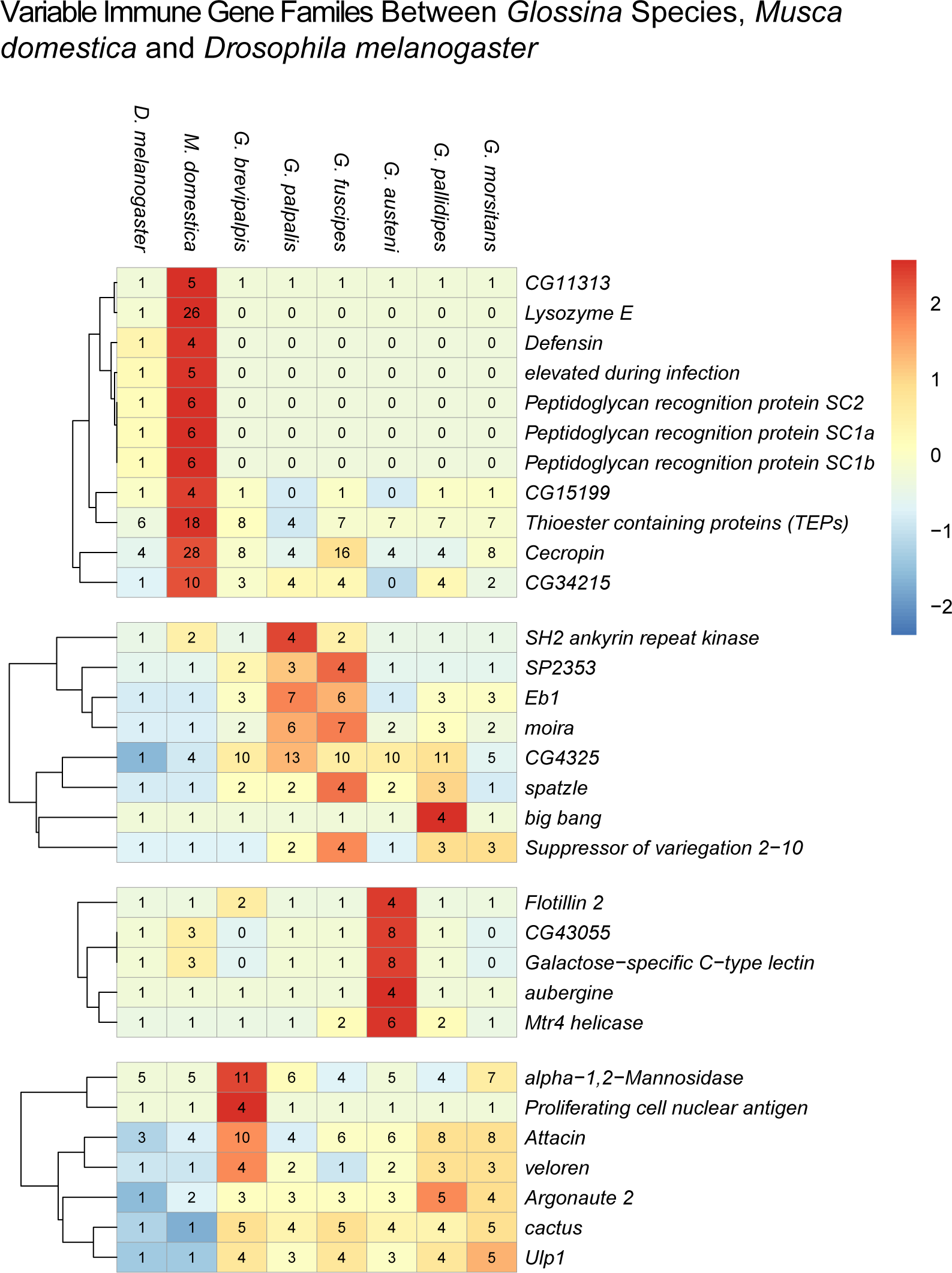
Heat map of counts of *Glossina* homologs to *Drosophila* immune genes. Counts of *Glossina* sequences within ortholog groups containing *Drosophila* genes annotated with the “Immune System Process” GO tag (GO:0002376).

The exception to these observations is element F. Element F, the smallest X-linked element, has low female expression and an excess of genes with male-biased expression (Supplemental Figure 8). In contrast with the other X-linked Muller elements in *Glossina*, the dN/dS ratios of all Element F associated genes (male biased and unbiased) suggest that they are evolving faster than the rest of the genome across all tsetse lineages (Supplemental Figure 9). The F elements in *Drosophila* species, while not being X-linked, show similar properties in that they have lower levels of synteny, increased rates of inversion, and higher rates of protein coding sequence evolution, suggesting that the F element is rapidly evolving in flies within Schizophora [42].

**Figure 8:**
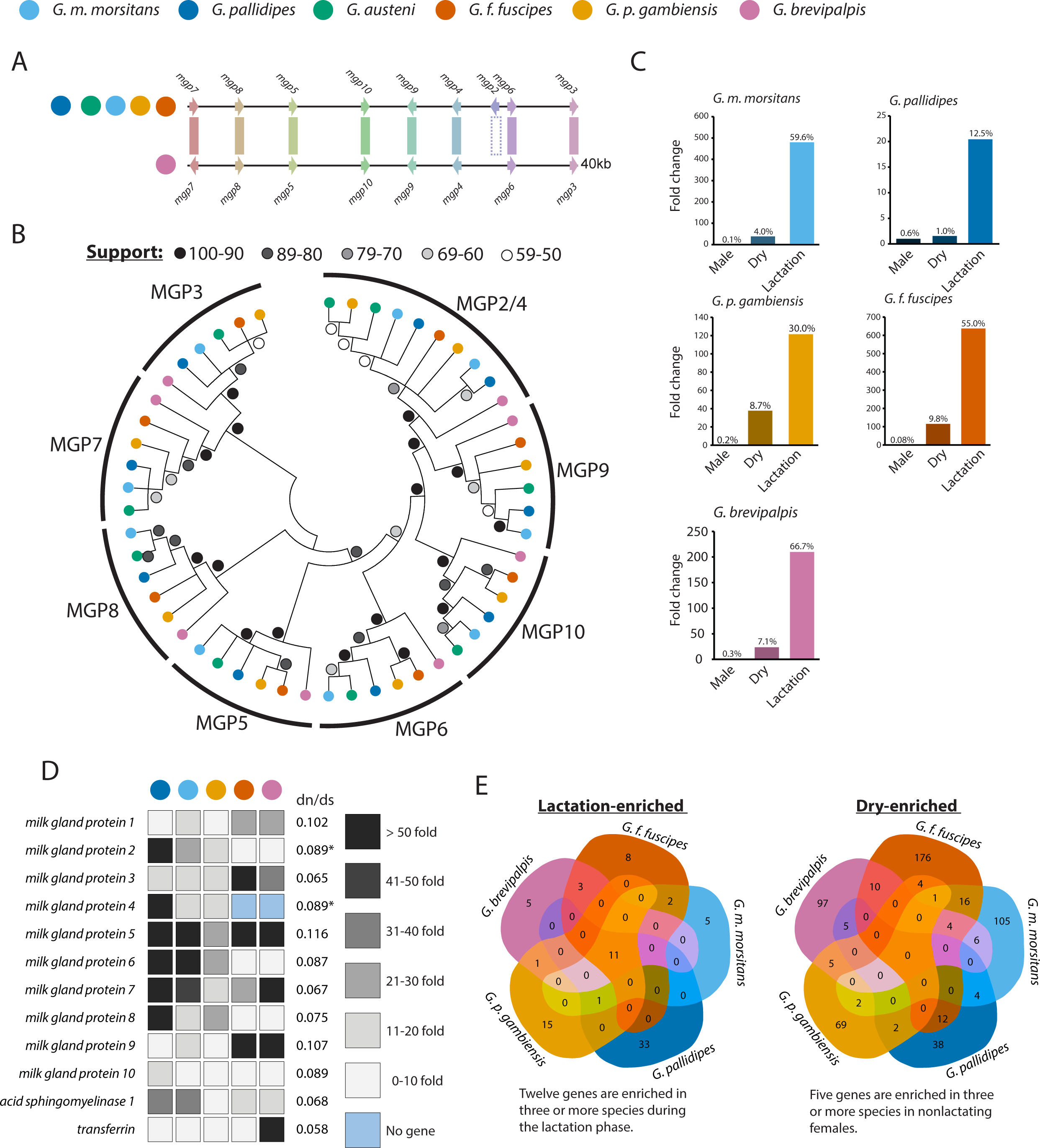
Conservation of synteny, sequence homology and stage/sex specific expression of tsetse milk proteins between species. Overview of the conservation of tsetse milk protein genes and their expression patterns in males, non-lactating and lactating females. A.) Syntenic analysis of gene structure/conservation in the *mgp2-10* genetic locus across *Glossina* species. B.) Phylogenetic analysis of orthologs from the *mgp2-10* gene family. C.) Combined sex and stage specific RNA-seq analysis of relative gene expression of the 12 milk protein gene orthologs in males, non-lactating and lactating females of 5 *Glossina* species. D.) Visualization of fold change in individual milk protein gene orthologs across 5 species between lactating and non-lactating female flies. Gene sequence substitution rates are listed for each set of orthologous sequences. E.) Comparative enrichment analysis of differentially expressed genes between non-lactating and lactating female flies.

**Figure 9:**
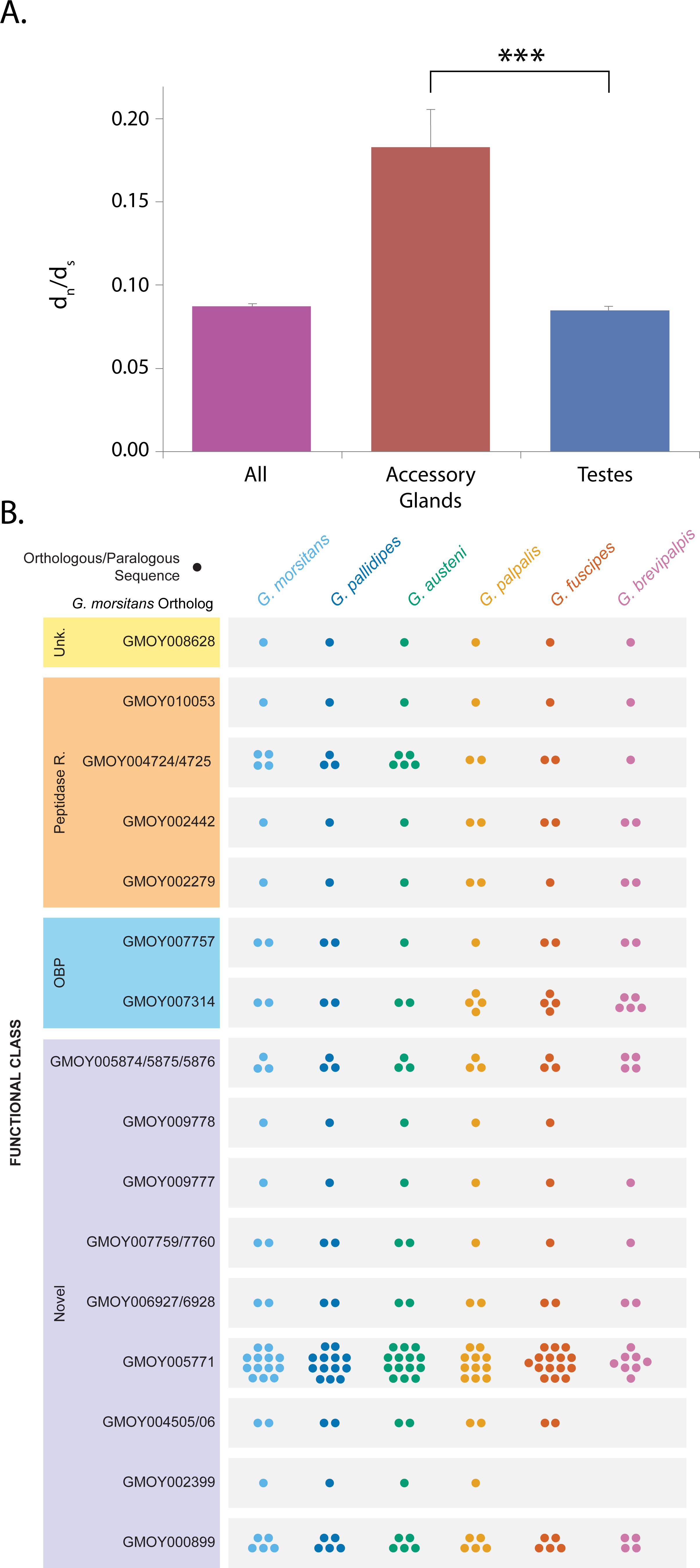
Comparative analysis of *Glossina* male accessory gland (MAG) protein family memberships. Graphical representation of the putative *Glossina* orthologs and paralogs to characterized MAG genes from *G. m. morsitans*. The genes are categorized by their functional classes as derived by orthology to characterized proteins from *Drosophila* and other insects.

### The *G. austeni* genome contains *Wolbachia* derived chromosomal insertions (Figure 4)

A notable feature of the *G. m. morsitans* genome was the integration of large segments of the *Wolbachia* symbiont genome via horizontal gene transfer (HGT). Characterization of the *G. m. morsitans* HGT events revealed that the chromosomal sequences with transferred material contained a high degree of nucleotide polymorphisms, coupled with insertions, and deletions [43]. These observations were used in this analysis to distinguish cytoplasmic from chromosomal *Wolbachia* sequences during the *in-silico* characterization of the tsetse genomes. Analysis of the six assemblies revealed that all contain *Wolbachia* sequences although *G. pallidipes, G. fuscipes*, *G. palpalis* and *G. brevipalpis* had very limited DNA sequence that displayed homology with *Wolbachia*. Furthermore, analysis of fly lines from which the sequenced DNA was obtained with *Wolbachia* specific primers PCR-amplification was negative. This is in line with PCR-based screening of *Wolbachia* infections in natural populations, further indicating that these short segments could be artifacts or contaminants [44]. However, *G. austeni* contains more extensive chromosomal integrations of *Wolbachia* DNA (Supplemental Table 2).

All *Wolbachia* sequences, chromosomal and cytoplasmic, identified in *G. austeni* were mapped against the reference genomes of *Wolbachia* strains *wMel*, *wGmm*, and the chromosomal insertions A & B in *G. m. morsitans* (Figure 4). The *G. austeni* chromosomal insertions, range in size from 500 - 95,673 bps with at least 812 DNA fragments identified *in silico*. Sequence homology between *w*Mel, *w*Gmm, and the chromosomal insertions A and B in *G. m. morsitans morsitans* varied between 98 – 63%, with the highest sequence homologies observed with chromosomal insertions A and B from *G. m. morsitans*. The similarity between the genomic insertions in *G. m. morsitans* and *G. austeni* relative to cytoplasmic *Wolbachia* sequences suggests they could be derived from an event in a common ancestor. The absence of comparable insertions in *G. pallidipes* (a closer relative to *G. m. morsitans*) indicate that either these insertions occurred independently or that the region containing the insertions was not assembled in *G. pallidipes*. Additional data from field based *Glossina* species/sub-species is required to determine the true origin of these events.

### Analysis of *Glossina* genus and sub-genus specific gene families reveals functional enrichments

All annotated *Glossina* genes were assigned to groups (orthology groups - OGs) containing predicted orthologs from other insect and arthropod species represented within Vectorbase. A global analysis of all the groups containing *Glossina* genes was utilized to determine the gene composition of these flies relative to their Dipteran relatives and between the Glossina sub-genera. An array of twelve Diptera are represented within this analysis including *Anopheles gambiae* (Nematocera), *Aedes aegypti* (Nematocera), *Lutzomyia longipalpis* (Nematocera), *Drosophila melanogaster* (Brachycera), *Stomoxys calcitrans* (Brachycera) and *Musca domestica* (Brachycera).

The tsetse associated OGs are represented by groups ranging from universal to the Diptera included in the analysis to species specific to the individual tsetse species. The composition of these OGs breaks down to a core of 3,058 OGs with constituents universal to Diptera (93430 genes), 299 OGs specific and universal to Brachyceran flies (4975 genes) and 162 OGs specific and universal to *Glossina* (1548 genes). A dramatic feature identified by this analysis is the presence of 2,223 OGs specific and universal to the *Palpalis* sub-genus (*G. fuscipes* and *G. palpalis* 4948 genes). This contrasts with the members of the *Morsitans* sub-genus (*G. m. morsitans*, *G. pallidipes* and *G. austeni*) in which there are 137 specific and universal OGs (153 genes) (Figure 5, Supplemental table 3, Supplemental data 2+3).

To understand the functional significance of the *Glossina* specific OGs, we performed an analysis of functional enrichment of gene ontology (GO) terms within these groups. Many of the *Glossina* specific genes are not currently associated with GO annotations as they lack characterized homologs in other species. As such these sequences were not included in this analysis. However, ~60% of the genes within the combined *Glossina* gene repertoire are associated with GO annotations, which allowed for analysis of a sizable proportion of the dataset.

### *Glossina* genus universal and specific genes are enriched in genes coding for proteases and odorant binding proteins

The orthology groups containing genes specific and universal to the *Glossina* genus are enriched in odorant binding and serine-type endopeptidase activities. The universality of these genes within *Glossina* and their absence from the other surveyed Dipteran species suggests they are currently associated with tsetse specific adaptations.

The ontology category with the lowest p-value represents proteolysis associated genes. This category encompasses 92 *Glossina* specific proteases with predicted serine-type endopeptidase activity. The abundance of this category may be an adaptation to the protein-rich blood specific diet of both male and female flies. A similar expansion of serine proteases is associated with blood feeding in mosquitoes and the presence of an equivalent expansion in tsetse may represent an example of convergent evolution [45]. This class of peptidases is also associated with critical functions in immunity, development and reproduction in Diptera [46–49].

The other enriched GO term common to all *Glossina* is for genes encoding odorant binding proteins (OBPs). Of the 370 OBPs annotated within *Glossina*, 55 lack orthologs in species outside of *Glossina*. The primary function of OBPs is to bind small hydrophobic molecules to assist in their mobilization in an aqueous environment. These proteins are primarily associated with olfaction functions as many are specifically expressed in chemosensory associated tissues/organs where they bind small hydrophobic molecules and transport them to odorant receptors [50, 51]. However, functional analyses in *G. m. morsitans* have associated an OBP (OBP6) with developmental activation of hematopoiesis during larvigenesis in response to the mutualistic *Wigglesworthia* symbiont [52]. In addition, many of the OBPs identified in this analysis are characterized as *Glossina* specific seminal proteins with male accessory gland specific expression patterns. They are primary constituents of the spermatophore structure produced by the male tsetse during mating [53]. The genus specific nature of these OBPs suggests that they are key components of reproductive adaptations of male tsetse.

### The *Palpalis* sub-genus contains a large group of sub-genus specific genes

A large group of genes specific and universal to members of the Palpalis sub-genus (*G. palpalis* and *G. fuscipes*) was a defining feature of the orthology analysis. The expansion includes 2223 OGs and encompasses 4948 genes between *G. palpalis* and *G. fuscipes*. Homology based analysis of these genes by comparison against the NCBI NR database revealed significant (e-value < 1×10^−10^) results for 603 of the genes. Within this subset of genes, ~ 5% represent bacterial contamination from tsetse’s obligate endosymbiont *Wigglesworthia*. Sequences homologous to another well-known bacterial symbiont *Spiroplasma* were found exclusively in *G. fuscipes*. This agrees with previous observations of *Spiroplasma* infection of colonized and field collected *G. fuscipes* flies [54].

Four genes bear homology to viral sequences (*GPPI051037/GFUI045295* and *GPPI016422/GFUI028200*). These sequences are homologus to genes from Ichnoviruses. These symbiotic viruses are transmitted by parasitic Ichneumonid wasps with their eggs to suppress the immune system of host insects [55]. These genes may have originated from a horizontal transfer event during an attempted parasitization.

Another feature of note is the abundance of putative proteins with predicted helicase activity. Of the 603 genes with significant hits, 64 (10.5%) are homologous to characterized helicases. Functional enrichment analysis confirms the enrichment of helicase activity in this gene set. These proteins are associated with the production of small RNA’s (miRNAs, siRNAs and piRNAs) which mediate posttranscriptional gene expression and the defensive response against viruses and transposable elements. Of the 64 genes, 41 were homologous to the armitage (*armi*) helicase. Recent work in *Drosophila* shows that *armi* is a reproductive tissue specific protein and is responsible for binding and targeting mRNAs for processing into piRNAs by the PIWI complex [56]. The reason for the accumulation of this class of genes within the *Palpalis* sub-genus is unknown. However, given the association of these proteins with small RNA production they could be associated with a defensive response against viral challenges or overactive transposable elements. A similar phenomena is seen in *Aedes aegypti* where components of the PIWI pathway have been amplified and function outside of the reproductive tissues to generate piRNAs against viral genes [57].

### Analysis of gene family variations reveals sub-genus specific expansions and contractions of genes involved in sperm production and chemosensation

In addition to unique gene families, we identified orthology groups showing significant variation in gene numbers between *Glossina* species. Of interest are groups showing significant sub-genus specific expansions or contractions, which may represent lineage specific adaptations. General trends that we observed in these groups show the largest number of gene family expansions within the *Palpalis* sub-genus and the largest number of gene family contractions within *G. brevipalpis* (a member of the *Fusca* sub-genus) (Figure 6 - For completely annotated figures with descriptions and group IDs see Supplemental Figures 11 and 12, respectively).

#### *Palpalis sub-genus specific expansion of sperm associated genes* (Supplemental data 4)

Members of the *Palpalis* sub-genus had a total of 29 gene family expansions and 1 contraction relative to the other 4 tsetse species. Of the three sub-Genera, this represents the largest number of expansions and parallels with the large number of *Palpalis* specific orthology groups. Two gene families expanded within the *Palpalis* group (VBGT00770000031191 and VBGT00190000014373) encode WD repeat containing proteins. The *Drosophila* orthologs contained within these families (*cg13930*, *dic61B*, *cg9313*, *cg34124*) are testes specific and associated with cilia/flagellar biosynthesis and sperm production [58]. Alteration/diversification of sperm associated proteins could explain the split of the *Palpalis* sub-genus from the other *Glossina* and the potential incipient speciation documented between *G. palpalis* and *G. fuscipes* [59].

#### The Morsitans sub-genera shows reductions in chemosensory protein genes

Within the *Morsitans* sub-genus six gene families are expanded and two are contracted relative to the other tsetse species. Of interest, one of the contracted gene families encodes chemosensory proteins (VBGT00190000010664) orthologous to the CheB and CheA series of proteins in *D. melanogaster*. The genes encoding these proteins are expressed exclusively in the gustatory sensilla of the forelegs of male flies and are associated with the detection of low volatility pheromones secreted by the female in higher flies [60]. Of interest is that the number of genes in *G. palpalis* (14), *G. fuscipes* (15) and G. *brevipalpis* (14) are expanded within this family relative to *D. melanogaster* (12), *M. domestica* (10) and *S. calcitrans* (4). However, the *Morsitans* group flies *G. m. morsitans* (7), *G. pallidipes* (7) and *G. austeni* (5) all appear to have lost some members of this family. The functional significance of these changes is unknown. However, it could represent an optimization of the male chemosensory repertoire within the *Morsitans* sub-genus.

In terms of expanded gene families in *Morsitans*, we find two encoding enzymes associated with the terpenoid backbone biosynthesis pathway (VBGT00190000010926 -farnesyl pyrophosphate synthase and VBGT00840000047886 – farnesol dehydrogenase). This pathway is essential for the generation of precursors required for the synthesis of the insect hormone Juvenile Hormone (JH). In adult *G. m. morsitans*, JH levels play an important role in regulating nutrient balance before and during pregnancy. High JH titers activate lipid biosynthesis and accumulation in the fat body prior to lactation. During lactation, JH titers fall, resulting in the catabolism and mobilization of stored lipids for use in milk production [61].

### Comparative analysis of the immune associated genes in *Glossina* species reveals specific expansions, contractions and losses relative to *Musca domestica* and *Drosophila melanogaster*

Tsetse flies are exposed to bacterial, viral, protozoan and fungal microorganisms exhibiting a broad spectrum of beneficial, commensal, parasitic and pathogenic phenotypes within their host. Yet, the diversity and intensity of the microbial challenge facing tsetse flies is limited relative to that of related Brachyceran flies such as *D. melanogaster* and *M. domestica* in terms of level of exposure, microbial diversity and host microbe relationships. While tsetse larvae live in a protected environment (maternal uterus) feeding on maternally produced lactation secretions, larval *D. melanogaster* and *M. domestica* spend their entire immature development in rotting organic materials surrounded by and feeding on a diverse array of microbes. The adult stages also differ in that tsetse feed exclusively on blood which exposes them to a distinct yet limited array of microbial fauna. The immune function and genetic complement of *D. melanogaster* is well characterized and provides the opportunity to compare the constitution of orthologous immune gene sequences between *M. domestica* and the *Glossina* species [62]. Orthology groups containing *Drosophila* genes associated with the ‘Immune System Process’ GO tag (GO:0002376) were selected and analyzed to measure the presence/absence or variance in number of orthologous sequences in *Glossina* (Figure 7 + Supplemental Table 5).

Several orthologs within this ontology group are highly conserved across all species and are confirmed participants with the fly’s antimicrobial immune response. These genes include the peptidoglycan recognition proteins (PGRPs) (with the exception of the PGRP SC1+2 genes) [63], prophenoloxidase 1, 2 and 3 [52], the reactive oxygen intermediates *dual oxidase* and *peroxiredoxin 5* [64, 65], and antiviral (RNAi pathway associated) *dicer 2* and *argonaute 2*. The antimicrobial peptide encoding genes *attacin* (variants A and B) and *cecropin* (variants A1, A2, B and C) are found within *Glossina*, but have diverged significantly (the highest % identity based on blastx comparison = 84%) from closely related fly taxa [66–68].

#### Glossina species are missing immune gene families present in D. melanogaster and M. domestica

Several gene families are missing within the *Glossina* species although expanded within *M. domestica* (Figure 7). These include *lysozyme E*, *defensin*, *elevated during infection*, and the *PGRP-SC1+2* gene families. These may be adaptations to the microbe rich diet and environment in which *M. domestica* larvae and adults exist. The expansion of immune gene families in *M. domestica* relative to *D. melanogaster* was previously documented in the publication of the *M. domestica* genome [69]. However, the added context of the *Glossina* immune gene complement highlights the significance of the expansion of these families relative to their loss in all *Glossina* species. The loss of these families may represent the reduced dietary and environmental exposure to microbial challenge associated with the dramatic differences in life history between these flies.

#### Glossina species show immune gene family expansions associated with the Toll and IMD pathways

In contrast, we observed several *Glossina* immune related gene families which are expanded relative to orthologous families in *Drosophila* and *M. domestica* (Figure 7). Duplications of this nature often reflect evolutionarily important aspects of an organism’s biology, and in the case of tsetse, may have resulted from the fly’s unique association with parasitic African trypanosomes. Prominent among the expanded immune related *Glossina* genes are those that encode *Attacin A* and *Attacin B*, which are IMD pathway produced effector antimicrobial molecules, and *Cactus*, a component of the Toll signaling pathway. Similarly, the most highly expanded immune related gene across *Glossina* species are the orthologs of *Drosophila* CG4325. RNAi-based studies in *Drosophila* indicate that CG4325 is a regulator of both the Toll and IMD signaling pathways [70]. Significant expansion of this gene family in *Glossina* substantiates previously acquired data that demonstrated the functional importance of the Toll and IMD pathways in tsetse’s response to trypanosome challenge [71, 72]. Finally, all six *Glossina* genomes encode multiple copies of *moira*. This gene, which is involved in cell proliferation processes [73], is differentially expressed upon trypanosome infection when compared to uninfected *G. m. morsitans* [74]. In an effort to eliminate parasite infections, tsetse flies produce reactive oxygen intermediates that cause collateral cytotoxic damage [64]. Additionally, trypanosome infection of tsetse’s salivary glands induces expression of fly genes that encode proteins associated with stress and cell division processes, further indicating that parasite infection results in extensive damage to host cells. Expansion of *moira* gene copy number in *Glossina’s* genome may reflect the fly’s need to maintain epithelial homeostasis in the face of damage caused by trypanosome infections.

#### G. brevipalpis has a species-specific expansion of immune associated proteins

An interesting highlight from this analysis is the identification of a gene expansion associated with alpha-mannosidase activity (VBGT00190000009892). An orthologous *Drosophila* gene (α*-Man-Ia*) is an essential component in the encapsulation response by hemocytes to attack by parasitoid wasps. This enzyme modifies lamellocyte surface glycoproteins to facilitate the recognition and encapsulation of foreign bodies. As described in the *G. m. morsitans* genome paper and here, there is evidence of parasitization by parasitoid wasps in the genomes of these flies in the form of integrated gene sequences homologous to polynavirus genes [26]. The expansion of these proteins could be an evolutionary response to pressure induced by parasitization although the current status of tsetse associated parasitoids is unknown.

### Tsetse reproductive genetics

#### Milk protein genes are universal and tightly conserved in Glossina (Figure 8 + Supplemental table 6)

The intrauterine development and nourishment of individual larval offspring is a defining characteristic of the *Hippoboscoidea* superfamily, which includes the *Glossinidae* (Tsetse flies), *Hippoboscidae* (Ked flies), *Nycteribiidae* (Bat flies), and *Streblidae* (Bat flies) families [75]. Nutrient provisioning is accomplished by the secretion of a milk-like substance from specialized glands into the uterus where the larval flies consume the milk. Dry weight of tsetse milk is roughly 50% protein and 50% lipids [76]. A compiled list of the milk protein orthologs from the six species of tsetse have been assembled (Supplemental Table 6).

Milk protein genes 2-10 (*mgp2-10*) in *G. m. morsitans* are the largest milk protein gene family. These genes are tsetse specific, lack conserved functional protein domains and their origin is currently unknown. However, experimental evidence suggests they act as lipid emulsification agents and possible phosphate carrier molecules in the milk [77]. Search for orthologous sequences to these genes revealed 1:1 orthologs to each of the 9 genes in the 5 new *Glossina* species except for *G. brevipalpis* which lacks an orthologous sequence for the *mgp2* gene. These genes are conserved at the levels of both synteny and sequence (Figure 8A+B). Comparative expression analysis of these genes (and the other characterized milk protein orthologs: *milk gland protein 1*, *acid sphingomyelinase* and *transferrin* [78, 79]) in male, non-lactating and lactating females shows sex and lactation specific expression profiles across the five species for which sex-specific RNA-seq data was available (Figure 8C+D). Comparison of sequence variation across species for these genes by dN/dS analysis indicates that they are under heavy negative selective pressure (Figure 8D). Enrichment analysis based on comparison of lactation-based RNA-seq data confirms that these 12 orthologous sequences are enriched in lactating flies across all *Glossina* (Figure 8E). The *mgp2-10* gene family is a unique and conserved adaptation that appears essential to the evolution of lactation in the *Glossina* genus. Determination of the origins of this protein family requires genomic analyses of other members of the Hippoboscoidea superfamily that exhibit viviparity along with other species closely related to this group.

### Tsetse seminal protein genes are rapidly evolving and vary in number and sequence conservation between species (Figure 9)

Recent proteomic analysis of male seminal proteins in *G. m. morsitans* revealed an array of proteins transferred from the male to the female as components of the spermatophore [80]. Cross referencing of the proteomic data with tissue specific transcriptomic analyses of the testes and male accessory glands (MAGs) allowed us to identify the tissues from which these proteins are derived. Many of the MAG associated proteins are *Glossina*-specific and are derived from gene families with multiple paralogs. These sequences were used to identify and annotate orthologous sequences in the other five *Glossina* species. In contrast to the milk proteins, sequence variance and differences in paralog numbers varies in male reproductive genes between the six *Glossina* species.

This is particularly evident in the genes with MAG biased/specific expression. MAG biased/specific genes are represented by 22 highly expressed gene families encoding characterized seminal fluid proteins (SFP). We investigated the evolutionary rate of reproductive genes over-expressed in the MAGs and testes, relative to a set of 5,513 *G. m. morsitans* genes, orthologous between the six species (Figure 9A). The average dN/dS ratio is higher in MAG biased genes than in testes biased genes or the entire *Glossina* ortholog gene set suggesting that the MAG genes are under relaxed selective constraints. In addition, we found high heterogeneity in the selective pressure across MAG genes. This is specifically evident in the tsetse specific genes *GMOY002399*, *GMOY007759*, *GMOY004505* and *GMOY005874* (a protein with OBP like conserved cysteine residues) as well as the OBP ortholog *GMOY007314*. All five genes encode seminal fluid proteins as confirmed by the proteomic analysis of the spermatophore [80].

In addition to sequence variability the number of paralogs per species differs as well (Figure 9B). This is similar to comparative analysis observations in *Anopheles* and *Drosophila* species [81, 82]. This variance is especially evident in *Glossina* specific protein families (i.e. *GMOY002399*, *GMOY004505/4506*, *GMOY005771*). In particular, there are a large number of gene orthologs/paralogs to the *GMOY005771* gene across all *Glossina* species revealing a large family of MAG genes of unknown function. The number of orthologs/paralogs differs significantly between *Glossina* species. In addition, the two *G. m. morsitans* paralogs *GMOY004724* and *GMOY004725* (predicted peptidase regulators), appear to display a higher number of putative gene duplications in the *Morsitans* sub-genus relative to the *Palpalis* and *Fusca* sub-Genera. Conservation appears instead to be more evident across testes genes that code for proteins associated with conserved structural and functional components of sperm.

Overall, comparison of the MAG biased genes across *Glossina* reveals that this group shows substantial variability in terms of genomic composition and rate of evolution. This is in agreement with other studies indicating that male accessory proteins evolve at high rates due to intraspecific competition between males or sexually antagonistic coevolution between males and females [83].

### Olfactory associated protein-coding genes are conserved and reduced in number relative to other Diptera

Comparative analyses of genes responsible for perireceptor olfaction activities revealed high conservation of the repertoire among the six species. The genes appear to scatter across their respective genomes with only a few duplicates occurring in clusters [84]. *Glossina* species expanded loci that include Gr21a (responsible for CO_2_ detection) [85], Or67d (mediates *cis*-vaccenyl acetate reception) and Obp83a, (thought to be olfactory specific) [86]. The expanded loci suggest involvement of gene duplication and/or transposition in their emergence [84]. All six species lack sugar receptors likely as a result of tsetse’s streamlined blood-feeding behavior.

Although our analysis did not reveal major discrepancies among the species, *G. brevipalpis* has lost three key gustatory receptors (Gr58c, Gr66a and Gr32a) compared to other species. In addition, *G. brevipalpis* showed higher structural gene rearrangements that could be attributed to its evolutionary distance relative to the other tsetse species [87].

### A salivary protein gene shows sub-genus specific repeat motifs (Figure 10)

Efficient acquisition of a blood meal by tsetse relies on a broad repertoire of physiologically active saliva components inoculated at the bite site. These proteins modulate early host responses, which, in addition to facilitating blood feeding can also influence the efficacy of parasite transmission [88, 89]. The differences in the competence of different tsetse fly species to develop mature *T. brucei* salivary gland infections may also be correlated with species-specific variations in saliva proteins. Tsetse saliva raises a species-specific IgG response in their mammalian hosts [90]. This response could potentially function as a biomarker to monitor exposure of host populations to tsetse flies [91].

**Figure 10:**
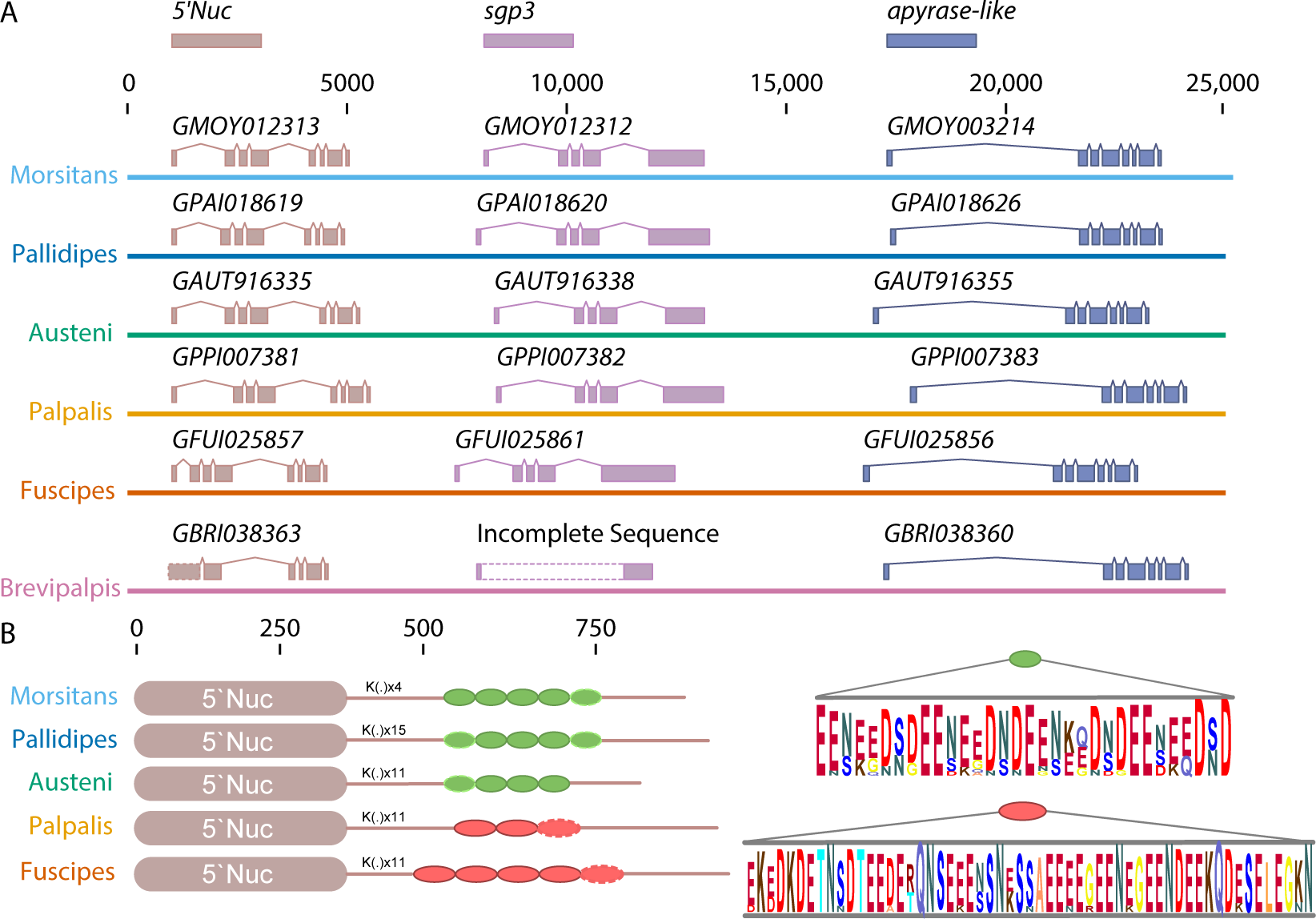
5’Nuc/apyrase salivary gene family organization and sequence features across Glossina species. A.) Chromosomal organization of the 5’Nuc/apyrase family orthologs on genome scaffolds from the six *Glossina* species. The brown gene annotations represent 5’Nuc gene orthologs; purple gene annotations represent *sgp3* gene orthologs and the blue gene annotations an apyrase-like encoding gene. The broken rectangular bars on the *G. brevipalpis* scaffold indicate that the sequence could not be determined due to poor sequence/assembly quality. B.) Schematic representation of sgp3 gene structure in tsetse species. The K(.) denotes a repetition of a Lysine (K) and another amino acid (Glutamic acid, Glycine, Alanine, Serine, Asparagine or Arginine). The green oval represents a repetitive motif found in *Morsitans* sub-genus; the red oval represents a repetitive motif found in *Palpalis* group. The dashed line indicates a partial motif present. For each of the two motifs the consensus sequence is shown in the right by a Logo sequence. The poor sequence/assembly quality of the *G. brevipalpis* scaffold prevented inclusion of this orthology in the analysis.

The *sgp3* gene [92] is characterized in all the tsetse species by two regions: a metallophosphoesterase/5`nucleotidase and a repetitive glutamate/aspartate/asparagine-rich region (Figure 10A). The complete sequence for this gene from *G. brevipalpis* could not be obtained due to a gap in the sequence. The metallophosphoesterase/5`nucleotidase region is highly conserved between all tsetse species. However, the sequences contain sub-genus specific (*Morsitans* and *Palpalis*) repeat motifs within the glutamate/aspartate/asparagine region. The motifs differ in size (32 amino acids in the *Morsitans* group and 57 amino acids in the *Palpalis* group) and amino acid composition (Figure 10B). Moreover, within each sub-genus, there are differences in the number of repetitive motifs. Within the *Morsitans* group, *G. m. morsitans* and *G. pallidipes* have five motifs while *G. austeni* has only four. In the *Palpalis* group, *G. palpalis* has three repetitive motifs and *G. fuscipes* five. Between the metallophosphoesterase/5`nucleotidase and the glutamate/aspartate/asparagine-rich regions there are a series of amino acids doublets comprising a lysine at the first position followed on the second position by another amino acid (glutamic acid, glycine, alanine, serine, asparagine or arginine). These differences may account for the differential immunogenic ‘sub-Genus-specific’ antibody response caused by Sgp3 in *Morsitans* and *Palpalis* group flies [90].

### Comparison of vision associated Rhodopsin genes reveals conservation of motion tracking receptors and variation in receptors sensitive to blue wavelengths (Figure 11)

**Figure 11:**
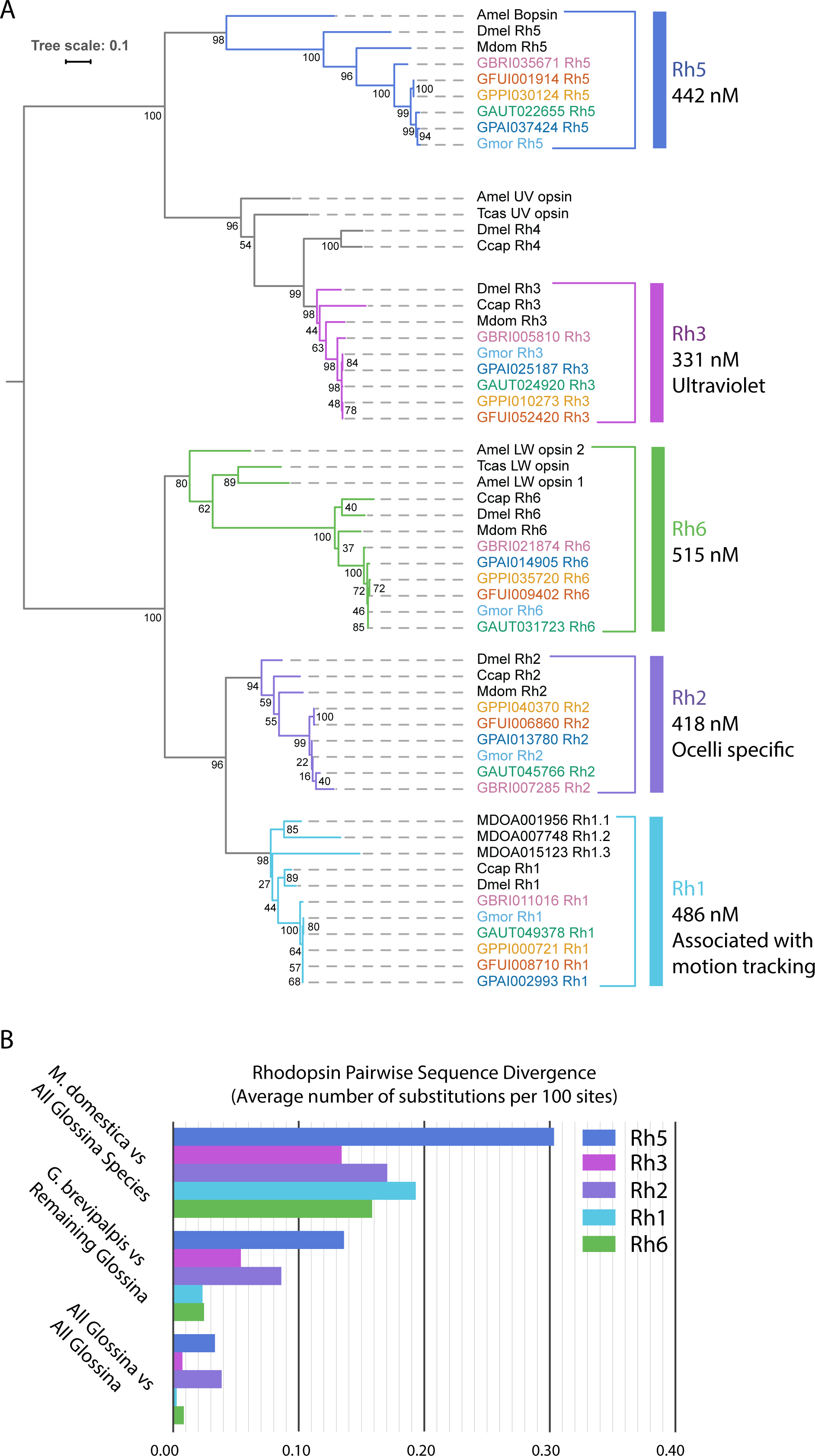
Phylogenetic and sequence divergence analysis of Glossina vision associated proteins. Phylogenetic and sequence conservation analysis of the vision associated *Rhodopsin* G-protein coupled receptor genes in *Glossina* and orthologous sequences in other insects. A.) Phylogenetic analysis of Rhodopsin protein sequences. B.) Pairwise analysis of sequence divergence between *M. domestica* and *Glossina* species and within the *Glossina* genus.

Vision plays an important role in host and mate seeking by flies within the *Glossina* genus. This aspect of their biology is a critical factor in the optimization and development of trap/target technologies [93, 94]. Analysis of the light sensitive Rhodopsin proteins across the *Glossina* species reveals orthologs to those described in the *G. m. morsitans* genome (Figure 11A). The expanded analysis provided by these additional genomes corroborates observations made for the original *G. m. morsitans* genome, including the conservation of the blue sensitive *Rh5* rhodopsin and the loss of one of the two dipteran UV-sensitive Rhodopsins: *Rh4* [26]. The availability of the new genomes provides complete sequences for an additional long wavelength sensitive Rhodopsin gene, *Rh2*. Prior to this analysis the recovery of a complete sequence from *G. m. morsitans* was not possible due to poor sequence quality at its locus.

Rhodopsin protein sequence divergence among the six *Glossina* species and *M. domestica* (as an outgroup) was investigated by calculating pairwise sequence divergence. As expected, the average pairwise sequence divergence between *M. domestica* and any *Glossina* species is higher than maximum sequence divergence among *Glossina* species for any of the five investigated Rhodopsin subfamilies, ranging between 0.13 to 0.3 substitutions per 100 sites. Average sequence divergence of *G. brevipalpis* to other *Glossina* is consistently lower than *Musca* vs *Glossina* but also higher than the average pairwise distances between all other *Glossina*, suggesting the older evolutionary lineage of *G. brevipalpis* (Figure 11B).

Three interesting aspects emerge in the comparison between subfamilies at the level of sequence divergence between *Glossina* species. The *Rh1* subfamily, which is deployed in motion vision, has the lowest average sequence divergence suggesting the strongest level of purifying selection. *Rh2*, which is expressed in the ocelli, and *Rh5*, which is expressed in color-discriminating inner photoreceptors, are characterized by conspicuously higher than average sequence divergence among *Glossina* species. This observation could account for the varying attractivity of trap and targets to different tsetse species.

## Conclusions

The comparative genomic analysis of these six *Glossina* species highlights important aspects of *Glossina* evolution and provides further insights into their unique biology. Additional comparative analyses of the genome assemblies, repetitive element composition, genes coding for neuropeptides and their receptors, cuticular protein genes, transcription factor genes and peritrophic matrix protein genes are available in the associated supplemental materials text, figures and data files (Supplemental Material; Supplemental Figures 10 and 13, Supplemental Tables 15, 16, 17, 18, 19; Supplemental Data 6 and 7). The results derived from the analysis of these genomes are applicable to many aspects of tsetse biology including host seeking, digestion, immunity, metabolism, endocrine function, reproduction and evolution. This expanded knowledge has important practical relevance. Indeed, tsetse control strategies utilize trapping as a key aspect of population management. These traps use both olfactory and visual stimuli to attract tsetse. The findings of a reduced contingent of olfactory associated genes and the variability of color sensing Rhodopsin genes provide research avenues into improvements of trap efficacy. Deeper understanding of the important chemosensory and visual stimuli associated with the different species could facilitate the refinement of trap designs for specific species. The findings associated with *Glossina* digestive biology, including the enrichment of proteolysis-associated genes and identification of *Glossina* specific expansions of immune associated proteins provide new insights and avenues of investigation into vector competence and vector/parasite relationships. Analysis of the female and male reproduction associated genes reveals the differential evolutionary pressures on females and males. The conservation of female milk proteins across species highlights the fact that this unique biology is optimized and under strong negative evolutionary pressure. In counterpoint, male accessory gland derived seminal proteins appear to have evolved rapidly between *Glossina* species and with little conservation relative to other Diptera in gene orthology and functional conservation. Tsetse reproduction is slow due to their unique viviparous adaptations, making these adaptations a potential target for the development of new control measures. The knowledge derived from these comparisons provide context and new targets for functional analysis of the genetics and molecular biology of tsetse reproduction. In addition to the practical aspects of the knowledge derived from these analyses, they also provide a look at the genetics underlying the evolution of unique adaptive traits and the resources to develop deeper understanding of these processes.

## Materials and Methods

### Aim

The aim of these studies was to generate and mine the genomic sequences of six species of tsetse flies with different ecological niches, host preferences and vectorial capacities. The goals of the analyses performed here are to identify novel genetic features specific to tsetse flies and to characterize differences between the *Glossina* species to correlate genetic changes with phenotypic differences in these divergent species. This was accomplished by the analyses described below.

### *Glossina* strains

All genomes were sequenced from DNA obtained from 2-4 lines of flies originating from individual pregnant females and their female offspring. Species collections were derived from laboratory strains with varied histories (See Supplemental Table 8). The *G. pallidipes, G. palpalis* and *G. fuscipes* flies were maintained in the laboratory at the Slovak Academy of Sciences in Bratislava, Slovakia. The *G. brevipalpis* strain were maintained in the Insect Pest Control Laboratory of the Joint FAO/IAEA Division of Nuclear Techniques in Food and Agriculture, Seibersdorf, Austria. Finally, *G. austeni* were obtained from the Tsetse Trypanosomiasis Research Institute in Tanga, Tanzania. Females were given two blood meals supplemented with 20 mg/ml tetracycline to cure them of symbionts to eliminate non-tsetse derived DNA.

### Genomic sequencing and assembly

Total genomic DNA was isolated from female pools for each species. High quality/ high molecular weight DNA was isolated from individual flies using Genomic-tip purification columns (QIAGEN) and the associated buffer kit. Samples were treated according to the protocol for tissue-based DNA extraction. The pooled individual DNA isolates were utilized for sequencing on Illumina HiSeq2000 instruments. The sequencing plan followed the recommendations provided in the ALLPATHS-LG assembler [95]. Using this model, we targeted 45x sequence coverage each of fragments (overlapping paired reads ~180bp length) and 3kb paired end (PE) sequences as well as 5x coverage of 8kb PE sequences. The first draft assembly scaffold gaps of each species were closed where possible with mapping of the same species assembly input sequences (overlapping paired reads ~180bp length) and local gap assembly [96]. Contaminating sequences and contigs 200bp or less were removed (Supplemental Table 9).

### Scaffold mapping to Muller Elements and Sex Specific Muller Element Expression Biases

We mapped scaffolds in each *Glossina spp.* genome assembly to chromosomes using homology relationships with *D. melanogaster* (Supplemental Table 1). This method exploits the remarkable conservation of chromosome arm (Muller element) gene content across flies [33, 97, 98]. We used the 1:1 orthologs between each *Glossina* species and *D. melanogaster* from OrthoDB [99] to assign scaffolds from each species to Muller elements, applying an approach previously developed for house fly [30]. For each species, a gene was assigned to a Muller element if it was a 1:1 ortholog with a *D. melanogaster* gene. Then, each scaffold was assigned to a Muller element if the majority (>50%) of genes with 1:1 orthologs on that scaffold were assigned to a single Muller element.

We used the RNA-seq data (described below) to compare gene expression in males and females. Expression comparisons were between male flies and either lactating (L) or non-lactating (NL) females.

### Repeat feature annotation

Repeat libraries for each species were generated using RepeatModeler [100]. The resultant libraries were used to annotate the genome with RepeatMasker [101], alongside tandem and low complexity repeats identified with TRF [102] and DUST [103]. The proportion of the genome covered by repeats is shown in (Supplemental Table 10), with the figures for *G. m. morsitans* provided for comparison.

### Automated gene annotation

Gene annotation was performed with MAKER [104], using the first 2 rounds to iteratively improve the training of the *ab initio* gene predictions derived from the combined Benchmarking Universal Single-Copy Orthologs (BUSCO) [105] and Core Eukaryotic Genes Mapping Approach (CEGMA) [106] HMMs, which were aligned to the genome assemblies using GeneWise [107]. RNA-seq data for each species (described below) were used to build a reference-guided transcriptome assembly with Tophat [108] and Cufflinks [109]. The initial MAKER analysis produced unrealistically high numbers of gene models, so InterProScan [110] and OrthoMCL [111] were used to identify gene predictions which lacked strong evidence. Only gene models that met one or more of the following criteria were retained: (a) an annotation edit distance < 1 [112]; (b) at least one InterPro domain (other than simple coils or signal peptides); (c) an ortholog in the *Glossina* species complex. This process resulted in a reduction of 12-25% in the number of gene models for each species (Supplemental Table 11). Genes from all six species were assigned to 15,038 orthology groups via the Ensembl Compara ‘GeneTrees’ pipeline [113].

For all types of ncRNA except tRNA and rRNA genes, we predicted RNA gene models by aligning sequences from Rfam [114] against the genome using BLASTN [115]. The BLAST results were then used to seed Infernal [116] searches of the aligned regions with the corresponding Rfam covariance models. rRNA genes were predicted with RNAmmer [117], and tRNA genes with tRNAScan-SE [118].

### Manual gene annotation

*Glossina* sequence data and annotation data were loaded into the Apollo [119] community annotation instances in VectorBase [120]. Manual annotations, primarily from a workshop held in Kenya in 2015, underwent both manual and automated quality control to remove incomplete and invalid modifications, and then merged with the automated gene set. Gene set versions are maintained at (www.vectorbase.org) for each organism. All highlighted cells relate to the current gene set version indicated in the table. Statistics for older gene set versions are provided along with the relevant version number.

### Genome completeness analysis (BUSCO and CEGMA Analysis)

Quality of the genome assembly and training of the *ab initio* predictors used in the gene prediction pipeline was determined using the diptera_odb9 database which represents 25 Dipteran species and contains a total of 2799 BUSCO (Benchmarking Universal Single-Copy Orthologs) genes derived from the OrthoDB v9 dataset [105] (Supplemental Table 12).

### Identification of Horizontal Gene Transfer Events

All genome sequence files for *G. pallidipes*, *G. palpalis*, *G. fuscipes*, *G. austeni*, and *G. brevipalpis* used for the whole genome assembly were also introduced into a custom pipeline for the identification of putative Horizontal Gene Transfer (HGT) events between *Wolbachia* and tsetse. *Wolbachia* sequences were filtered out from WGS reads using a combination of MIRA [121] and NextGenMap [122] mapping approaches. The reference sequences used were *w*Mel (AE017196), *w*Ri (CP001391), *w*Bm (AE017321), *w*Gmm (AWUH01000000), *w*Ha (NC_021089), *w*No (NC_021084), *w*Oo (NC_018267), *w*Pip (NC_010981), and the chromosomal insertions A and B in *G. morsitants morsitans.* All filtered putative *Wolbachia*-specific sequences were further examined using blast and custom-made databases.

To identify the chromosomal *Wolbachia* insertions, the following criteria were used: Sequences that (relative to the reference genomes): (a) exhibit high homology to insertion sequences A & B from *G. m. morsitans*, (b) exhibit a high degree of nucleotide polymorphisms (at least 10 polymorphisms/100bp) with the reference genomes, and (c) contain a high degree of polymorphism coupled with insertions and/or deletions. *Wolbachia* specific sequences for each *Glossina* species were assembled with MIRA using a *de novo* approach. For *G. pallidipes, G. palpalis, G. fuscipes*, and *G. brevipalpis* assembled sequences corresponding only to cytoplasmic *Wolbachia* were identified. Genomic insertions were only observed in assembled sequences from *G. austeni* (Supplemental Table 2). The statistics for the *G. austeni* assembled sequences are as follows: N50 4493, N90 1191, and mean contig length, 2778bps. During the process of identifying HGT events in *G. fuscipes*, we also recovered *Spiroplasma* sequences but none of the recovered sequences were chromosomal.

### Whole-genome pairwise alignment

We generated all possible pairwise alignments between the six *Glossina* species (including *G.m. morsitans*) and an outgroup, *M. domestica*, using the Ensembl Compara software pipeline [123]. LASTZ [124] was used to create pairwise alignments, which were then joined to create 'nets' representing the best alignment with respect to a reference genome [125]. *G. m. morsitans* was always used as the reference for any alignment of which it was a member, otherwise the reference genome was randomly assigned. Coverage statistics and configuration parameters for all alignments are available at https://www.vectorbase.org/compara_analyses.html.

### Glossina phylogeny prediction

We identified orthologous genes across the six *Glossina* species and six outgroups (*M. domestica*, *D. melanogaster*, *D. ananassae*, *D. grimshawi*, *L. longipalpis*, and *A. gambiae*) by employing a reciprocal-best-hit (RBH) approach in which *G. m. morsitans* was used as focal species. We identified 286 orthologs with a clear reciprocal relationship among the 12 species. All orthologs were aligned individually using MAFFT [126] and concatenated in a super-alignment of 478.617 nucleotide positions. The nucleotide alignment was translated in the corresponding amino acids and passed through Gblocks [127] (imposing “half allowed gap positions” and leaving remaining parameters at default) to obtain a dataset of 117.783 amino acid positions. This dataset was used for a Maximum Likelihood analysis in RAxML [128] employing the LG+G+F model of replacement, and for a Bayesian analysis using Phylobayes [129, 130] employing the heterogeneous CAT+G model of replacement. We further performed a coalescent-aware analysis using Astral [131] and the 286 single gene trees obtained using Raxml [128] and analyzing the alignments at the nucleotide level with the GTR+G model of replacement.

### Mitochondrial genome analysis and phylogeny

The mtDNA genomes of *G. m. centralis* and *G. brevipalpis* were sequenced using the Illumina HiSeq system and about 15 kb of mitochondrial sequence of each species was obtained. These sequences were used to identify the mtDNA sequences within the sequenced tsetse genomes (*G. pallidipes, G. m. morsitans, G. p. gambiensis, G. f. fuscipes* and *G. austeni*) from the available genomic data. Sanger sequencing confirmed the mtDNA genome sequence of each tsetse species. This involved PCR amplification of the whole mtDNA genome using fourteen pairs of degenerate primers designed to cover the whole mitochondrial genomes of the sequences species (Supplemental Table 13). The PCR products were sent for Sanger sequencing. The sequences obtained by Sanger and Illumina sequencing for each species were assembled using the SegMan program from the lasergene software package (DNAStar Inc., Madison, USA). The phylogenetic analysis based on these sequences was performed using maximum likelihood method with the MEGA 6.0 [132].

### Synteny analysis

The synteny analysis was derived from whole genome alignments performed as follows using tools from the UCSC Genome Browser [133]. The LASTZ software package (version 1.02.00) generated the initial pairwise sequence alignments with the parameters: *E=30, H=2000, K=3000, L=2200, O=400* and the default substitution matrix. From these alignments, Kent’s toolbox (version 349) [133] was used to generate chain and nets (higher-level abstractions of pairwise sequence alignments) with the parameters: *-verbose=0 -minScore=3000 and - linearGap=medium*. The multiple alignment format (MAF) files were built with MULTIZ for TBA package (version 01.21.09) [134], using the chains and nets, along with the phylogenetic relationships and distances between species. Using the MAF files, pairwise homologous synteny blocks (HSBs) were automatically defined using the SyntenyTracker software [135]. Briefly, the SyntenyTacker software defines an HSB as a set of two or more consecutive orthologous markers in homologous regions of the two genomes, such that no other defined HSB is within the region bordered by these markers. There are two exceptions to this rule: the first involves single orthologous markers not otherwise defined within HSBs; and the second involves two consecutive singleton markers separated by a distance less than the resolution threshold (10 kb for this analysis). As the 10 Kb blocks were too small for visualisation in Circos [136], they were aggregated into larger 100 Kb histogram blocks, where each 100 Kb Circos block shows the fraction of sequence identified as syntenic for a particular species when aligned to *D. melanogaster*. Synteny blocks are available for visualisation from the Evolution Highway comparative chromosome browser: http://eh-demo.ncsa.uiuc.edu/drosophila/.

### Orthology and paralogy inference and analysis

Phylogenetic trees were inferred with the Ensembl Compara 'GeneTrees' pipeline [123, 137] using all species from the VectorBase database of arthropod disease vectors [120]. The trees include 33 non-*Glossina* species, such as *D. melanogaster*, which act as outgroup comparators. All analyses are based on the VectorBase April 2016 version of the phylogenetic trees. Representative proteins from all genes were clustered and aligned, and trees showing orthologs and paralogs were inferred with respect to the NCBI taxonomy tree (http://www.ncbi.nlm.nih.gov/taxonomy).

The 15,038 predicted gene trees containing *Glossina* sequences were parsed to quantify the trees based on their constituent species. Raw tree files (Supplemental Data 1) were parsed using a custom PERL script (Supplemental File 1) to determine gene counts for representative Dipteran species for each gene tree. Count data were imported into Excel and filtered using pivot tables to categorize orthology groups based upon species constitution (Supplemental Data 2).

The orthology groups were broken into cohorts based on the phylogenetic composition of species within each group. The *Glossina* containing orthology groups were categorized as follows: common to Diptera (including the Nematocera sub-order), Brachycera sub-order specific, *Glossina* genus specific, *Glossina* sub-genus specific (*Morsitans* and *Palpalis*) or *Glossina* species specific. Each category is sub-divided into two groups, universal groups that contain representative sequences from all species within the phylogenetic category or partial orthology groups containing sequences from some but not all members of the phylogenetic category. *Glossina* gene IDs and associated FASTA sequences associated with groups of interest were extracted using a custom Perl script for gene ontology analysis (Supplemental File 2)

### Gene Ontology (GO) analysis

Gene associated GO terms were obtained from the VectorBase annotation database via the BioMart interface. Genes from *Glossina* genus, and sub-genus specific orthology groups were isolated and tested for enrichment of GO terms. Analysis for GO terms for enrichment was performed with the R package “topGO”. The enriched genes were separated into species specific lists compared against the entirety of predicted protein-coding genes from the respective species. Significance of enrichment was determined using Fisher’s Exact Test (Supplemental Table 3 + Supplemental Data 3).

### Identification and analysis of gene expansions/contractions

Gene trees containing orthologs/paralogs representing each of the six *Glossina* species were analyzed to identify sub-genus associated gene expansions/contractions. Gene trees were considered for analysis if the variance in the number of orthologs/paralogs between the 6 species was greater than 2. Variable gene trees were tested for phylogenetic significance relative to the predicted *Glossina* phylogeny using the CAFE software package [138] to reject potentially inaccurate variance predictions due to erroneous gene annotations. Gene trees with a CAFE score of <0.05 were considered significant (Supplemental table 4 + Supplemental data 4).

Sequences from gene trees satisfying the variance and CAFE thresholds were extracted with a custom PERL script (Supplemental File 2) and analyzed by BLASTP analysis against an insect specific subset of the NCBI NR database. Gene trees were annotated with the most common description associated with the top BLAST hits of its constituent sequences. Gene trees were subjected to PCA analysis in R using the FactoMineR and Factoextra packages using species specific gene counts as input data. The results were plotted and annotated with their associated BLAST derived descriptions.

### RNA-seq data

Total RNA was isolated for each of the six tsetse species from whole male and whole female (non-lactating and lactating) for RNA-seq library construction. Poly(A)+ RNA was isolated, then measured with an Agilent Bioanalyzer for quality. Samples were considered to be of high quality if they showed intact ribosomal RNA peaks and lacked signs of degradation. Samples passing quality control were used to generate non-normalized cDNA libraries with a modified version of the Nu-GEN Ovation^®^ RNASeq System V2 (http://www.nugeninc.com). We sequenced each cDNA library (0.125 lane) on an Illumina HiSeq 2000 instrument (~36 Gb per lane) at 100 bp in length.

RNA-seq analyses were conducted based on methods described in Benoit et al. [77], Rosendale et al.[139], and Scolari et al.[140] with slight modifications. RNA-seq datasets were acquired from whole males, whole dry females, and whole lactating females. The SRA numbers for each of the libraries are listed in (Supplemental Table 14).

RNA-seq datasets were quality controlled using the FastQC (Babraham Bioinformatics) software package. Each set was trimmed/cleaned with CLC Genomics (Qiagen) and quality was re-assessed with FastQC. Each dataset was mapped to the predicted genes from each *Glossina* genome with CLC Genomics. Each read required at least 95% similarity over 50% of length with three mismatches allowed. Transcripts per million (TPM) was used as a proxy for gene expression. Relative transcript abundance differences were determined as the TPM in one sample relative to the TPM of another dataset (*e.g.* male/lactating Female). A proportion based statistical analysis [141] followed by Bonferroni correction at 0.05 was used to identify genes with significant sex and stage specific transcript enrichment. This stringent statistical analysis was used as only one replicate was available for each treatment.

Enriched transcripts in lactating and dry transcriptomes from the species examined were compared to orthologous sequences in *G. m. morsitans* [142]. Overlap was determined by comparison of the enrichment status of orthologous sequences in the *Glossina* species tested. The results of this analysis are visualized in a Venn diagram (http://bioinformatics.psb.ugent.be/webtools/Venn/). Determination of dN/dS values and production of phylogenetic trees was conducted with the use of DataMonkey [143, 144] for dN/dS analyses and MEGA5 for alignment and tree construction [145].

### Cuticular Protein Analysis

The predicted peptide sequences from each species were analyzed by BLASTp analysis [115] against characteristic sequence motifs derived from several families of cuticle proteins [146]. Predicted cuticle proteins were further analyzed with CutProtFam-Pred, a cuticle protein prediction tool described in Ioannidou et al. [147], to assign genes to specific families of cuticle proteins. To find the closest putative homolog to cuticle protein genes in Glossina, genes were searched (BLASTp) against Refseq protein database from the National Center for Biotechnology Information (NCBI). The protein sequences with the lowest e-value were considered the closest putative homologs (Supplemental Data 5).

### Transcription factor identification and annotation

Likely transcription factors (TFs) were identified by scanning the amino acid sequences of predicted protein-coding genes for putative DNA binding domains (DBDs). When possible, we predicted the DNA binding specificity of each TF using the procedures described in Weirauch *et al.* [148]. Briefly, we scanned all protein sequences for putative DBDs using the 81 Pfam [149] models listed in Weirauch and Hughes [150] and the HMMER tool [151], with the recommended detection thresholds of Per-sequence Eval < 0.01 and Per-domain conditional Eval < 0.01. Each protein was classified into a family based on its DBDs and their order in the protein sequence (e.g., bZIPx1, AP2×2, Homeodomain+Pou). We then aligned the resulting DBD sequences within each family using clustalOmega [152], with default settings. For protein pairs with multiple DBDs, each DBD was aligned separately. From these alignments, we calculated the sequence identity of all DBD sequence pairs (*i.e.* the percent of AA residues that are identical across all positions in the alignment). Using previously established sequence identity thresholds for each family [148], we mapped the predicted DNA binding specificities by simple transfer. For example, the DBD of the *G. austeni* GAUT024062-PA protein is identical to the DBD of *D. melanogaster* mirr (FBgn0014343). Since the DNA binding specificity of mirr has already been experimentally determined, and the cutoff for Homeodomain family of TFs is 70%, we can infer that GAUT024062-PA will have the same binding specificity as mirr. All associated data can be found in (Supplemental Data 6)

## Supporting information

Text for Supplemental Analyses

Supplemental Table 1 Statistics on gene and scaffold mapping to Muller elements

Supplemental table 2 Wolbachia sequences

Supplemental Table 3 Gene Ontology (GO Molecular Function) Term Enrichment

Supplemental Table 4 sub Genus Specific Gene Expansions and Contractions

Supplemental Table 5 Count of Glossina orthologs to Immune Activity Associated Drosophila genes

Supplemental Table 6 Glossina Milk Protein Genes

Supplemental Table 7 Glossina Male Accessory Protein Genes

Supplemental Table 8 Glossina strains and colonies of origin

Supplemental Table 9 Glossina species contig and scaffold assembly statistics

Supplemental Table 10 Total measured repetitive elements among Glossina genomes

Supplemental Table 11 Predicted gene sets for new Glossina genomes

Supplemental Table 12 BUSCO Genomic Analysis Results

Supplemental Table 13 Primers used to amplify the mitochondrial genome

Supplemental Table 14 SRA Numbers

Supplemental Table 15 Repeat and TE Analysis

Supplemental Table 16 Neuropeptide genes

Supplemental Table 17 Protein receptor genes for bioactive neuropeptides

Supplemental Table 18 Family specific counts of putative cuticle protein genes

Supplemental Table 19 List of all peritrophin and peritrophin-like proteins

Supplemental Data 1 Raw_Ensemble_Dipteran_Orthology_Data

Supplemental Data 2 Orthology_Group_Analysis

Supplemental Data 3 topGO Results of Genus and sub-genus specific Gene IDs

Supplemental Data 4 Variable_Orthology_Group_Analysis

Supplemental Data 5 Immune Gene Analysis

Supplemental Data 6 Cuticle Protein Genes

Supplemental Data 7 Transcription Factor Data

Supplemental Figure 4 - X-Linkage of Male and Female Biased Genes

Supplemental Figure 5 - Autosome-X Expression Levels

Supplemental Figure 6 - dN-dS Autosome vs X

Supplemental Figure 7_dnds_sexbias_XA_scaff

Supplemental Figure 8 - Male Biased Expression per Muller Element

Supplemental Figure 9 - dN-dS per Muller Element

Supplemental Figure 10 - Genomic Composition of Glossina Transposable Elements

Supplemental Figure 11 - Sub-Genus specific Gene Families PCA Plot Descriptions

Supplemental Figure 12 - Sub-Genus specific Gene Families PCA Plot Indentifiers

Supplemental Figure 13 - Transcription Factor Constitution in Glossina

Supplemental File 1 Ensemble_Orthology_Data_Parser_PERL_Script

Supplemental File 2 Orthology_group_sequence_and_ID_extractor_PERL_Script

## Abbreviations

DBD: DNA binding domain
HAT: Human African Trypanosomiasis
AAT: Animal African Trypanosomiasis
mtDNA: mitochondrial DNA
rRNA: ribosomal RNA
tRNA: transfer RNA
HGT: horizontal gene transfer
OG: orthology group
GO: gene ontology
OBP: odorant binding protein
miRNA: micro RNA
siRNA: small interfering RNA
piRNA: piwi interacting RNA
MGP: milk gland protein
MAG: male accessory gland
SFP: seminal fluid protein

## Declarations

Ethics approval and consent to participate:

Not applicable

## Consent for publication

Not applicable

## Availability of data and material

The genomes, transcriptomes and predicted protein-coding sequences are available from VectorBase via the following link https://www.vectorbase.org/taxonomy/glossina. The raw RNA-seq datasets generated and/or analyzed during the current study are available from the NCBI SRA database repository, at the following link https://www.ncbi.nlm.nih.gov/sra/SRP158014. All data generated during the analyses of these dataset are included in this published article and its supplementary information files.

## Competing interests

### Funding

This work was supported by NIH Grants D43 TW007391, U01AI115648, R01AI051584, R03TW008413 and R03TW009444 to SA. Grant R21AI109263 to GA and SA from NIH-NIAID, Grant U54HG003079 from NIH-NHGRI to RKW and SA, McDonnell Genome Institute at Washington University School of Medicine. Partial funding from the National Research Foundation to HGM (Grant # 10924). Swiss National Science Foundation grant PP00P3_170664 to RMW. This research was partially supported by the Slovak Research and Development Agency under the contract No. APVV-15-0604 entitled “Reduction of fecundity and trypanosomiasis control of tsetse flies by the application of sterile insect techniques and molecular methods”.

### Contributions (^&^ = Annotation group leader, ^$^ = Project leader)

Analysis of whole genomic sequences and database management: D.L., E.L., G.L.M., M.B.; BUSCO and Female reproductive gene analysis: E.C.J., J.B.B.^&^, V.M.; Chemosensory gene analysis: D.M., P.O.M., R.W.M.; Cuticular protein gene analysis: A.J.R., D.W.F.; Gene orthology and expansion analyses: G.M.A.^$&^, J.E.A.; Genome sequencing, assembly and analysis: WCW^$^, C.T., P.M., R.K.W.; Horizontal gene transfer analysis: G.T., K.B.; Immune gene analysis: A.V., B.L.W., J.W., R.B.; Male reproductive gene analysis: A.R.M., F.S., G.S.; Mitochondrial DNA sequence analysis: A.M.M.A., I.M., A.G.P.; Molecular evolution and phylogenetic analyses: L.O., O R-S; Neuropeptide and G protein-coupled receptor analysis: H.G.M.^&^, J.C.^&^, L.S.; Rhodopsin gene analysis: M.F.; Orthology and comparative genomics analysis advice, manuscript editing: R.M.W.; Peritrophin gene analysis: A.A-S.^&^, C.R.; Project conception: S.A.^$^, M.J.L.^$^; Project funding: S.A. ^$^, Project management: S.A. ^$^, W.C.W., Provision of experimental material: P.T., S.A. ^$^, A.M.M.A.; Salivary protein gene analysis: J.V.D.A., I.M.^&^, G.C., X.Z.; Symbiont associated gene analysis: R.R.^&^; Syntenic analysis of genomes: M.T.S., D.M.L., V.P.E.L.; Transcription factor and DNA binding motif prediction: M.T.W.; Transposable element analysis: A.H.V., W.J.M.; X chromosome and sex linked expression analysis: R.P.M.

## Acknowledgements

We thank the production-sequencing group of McDonnell Genome Institute at Washington University for library construction, sequencing and data curation. Great thanks to the members of the Comparative Genomics workshop held at the Biotechnology Research Institute - Kenya Agricultural and Livestock Research Organization, Kikuyu, Kenya. Including: Muna Abry, Willis Adero, Erick Aroko, Joel Bargul, Tania Bishola, Lorna Jemosop Chebon, Appolinaire Djikeng, John Irungu, Everlyn Kamau, Christine Kamidi, Caleb Kibet, Esther Kimani, Kelvin Kimenyi, Mathuriin Koffi, Benard Kulohoma, Clarence Mangera, Abraham Mayoke, David Mburu, Grace Murilla, Mary Murithi, Ramadhan Mwakubambanya, Sarah Mwangi, Nelly Ndungu, Joyce Njuguna, Benson Nyambega, Faith Obange, Samuel Ochieng, Edwin Ogola, Owallah (Martin) Ogwang, Sylvance Okoth, Luicer Olubayo, Irene Onyango, Fred Osowo, David Price, Martin Rono, Sharon Towett, Kelvin Wachiuri, Kevin Wamae and Mark Wamalwa.

The workshop was sponsored by the D43 TW007391 award from the Fogarty International Center to S.A. and was facilitated by: Yale School of Public Health, Kenya Agricultural and Livestock Research Organization (KALRO), International Centre of Insect Physiology and Ecology (ICIPE), South African National Bioinformatics Institute (SANBI), International Livestock Research Institute (ILRI), Biosciences Eastern and Central Africa.

---

**Supplemental Figure 1:** Maximum Likelihood, Bayesian and Astral based phylogenetic analysis of a concatenated single gene ortholog alignment.

**Supplemental Figure 2:** Tsetse variable mitochondrial DNA sequence and species delineation by high resolution melt curve analysis

**Supplemental Figure 3:** Application of high-resolution melt curve analysis to distinguish Tsetse haplotypes/populations

**Supplemental Figure 4:** The percent of female-, male-, and un-biased genes that are on an X chromosome scaffold (Muller elements A, D, or F) is plotted for each species. Sex-biased expression was measured between males and either lactating (L) or non-lactating (NL) females. Asterisks indicate a significant difference between the percent of sex-biased genes that are X-linked when compared to unbiased genes (*P<0.05 in Fisher’s exact test).

**Supplemental Figure 5:** The distribution of the log2 (fold-change between females and males) is plotted for autosomal and X-linked genes in each species. Female-male gene expression comparisons are between males and either lactating or non-lactating females.

**Supplemental Figure 6:** Rates of non-synonymous to synonymous substitution (dN/dS) along different evolutionary lineages within the *Glossina* genus.

**Supplemental Figure 7:** Rates of non-synonymous to synonymous substitution (dN/dS) rates of female, male and non-sex biased genes across the X and autosomal muller elements.

**Supplemental Figure 8:** The distribution of male biased gene expression across the predicted *Glossina* Muller Elements. Bar heights represent the percentage of genes per element with male biased gene expression.

**Supplemental Figure 9:** Rates of non-synonymous to synonymous substitution (dN/dS) rates across the predicted muller elements.

**Supplemental Figure 10: Repetitive element constitution of Glossina genomes.** Analysis of repetitive element composition across the six *Glossina* species. A.) Graphical representation of the constitution and sequence coverage by the various classes of identified repetitive elements. B.) Relative constitution of DNA Terminal Inverted Repeat (TIR) families across the *Glossina* genomes. C.) Relative constitution of Long Interspersed Nuclear Elements (LINEs) across the *Glossina* genomes.

**Supplemental Figure 11: Sub-genus specific gene family expansions/retractions (with functional annotations).** Principal component analysis-based clustering of gene orthology groups showing significant differences in the number of representative sequences between the six *Glossina* species.

**Supplemental Figure 12: Sub-genus specific gene family expansions/retractions (with orthology group number annotations).** Principal component analysis-based clustering of gene orthology groups showing significant differences in the number of representative sequences between the six *Glossina* species.

**Supplemental Figure 13: Distribution of transcription factor families across insect genomes.** Heatmap depicting the abundance of transcription factor (TF) families across a collection of insect genomes. Each entry indicates the number of TF genes for the given family in the given genome, based on presence of DNA binding domains. Color key is depicted at the top (light blue means the TF family is completely absent) – note log (base 2) scale.

