## Supplementary material for "The *Glossina* Genome Cluster: Comparative Genomic Analysis of the Vectors of African Trypanosomes": Text for Supplemental Analyses

**Genome assemblies and global features of note**

The genomic sequences for the tsetse species described here originated from mother and daughter lines for each respective *Glossina* species. Sequencing and assembly of the resulting reads produced scaffolds of varied sizes, contiguity and coverage (Supplemental Table 9). The total assembled sequencing coverage varied between 40-58x for each species. The average assembled size was 359 Mb with the greatest contiguity measured for *G. pallidipes*, which comprised the fewest contigs (n=7,275) with an N50 contig length of 167 kb. On average, the new *Glossina* assemblies resulted in fewer contigs (17,604 vs 24,071) at a higher level of contiguity (72 vs 49kb) than the original *G.* *morsitans* assembly. This is likely due to the advancements in the sequencing technologies and software utilized to sequence and assemble these genomes relative to the original *G.* *morsitans* genome. The *G.* *morsitans* genome also has fewer predicted genes relative to the more recently produced genomes, suggesting that additional sequencing on this species would be informative.

The GC content of these genomes range from 27% (*G. brevipalpis*) to 35% (*G. pallidipes*) (Table 1). Genomic regions with low GC content are associated with heterochromatic DNA which is often transcriptionally inactive [85]. It is possible that the lower GC content in *G. brevipalpis* relative to other tsetse species could result in additional regions of lower transcriptional activity. In addition, *G. brevipalpis* has the smallest genome of the newly sequenced *Glossina* species yet 35% of it consists of repetitive sequences.

The completeness and accuracy of gene model predictions within the genomes was determined by Benchmarking Universal Single-Copy Orthologs (BUSCO) analysis (Supplemental Table 12). This analysis revealed high levels of representation of universal orthologs in all *Glossina* species. The scores for the genomes ranged from 92% representation (*G. morsitans*) to between 97-98% (the remaining five *Glossina* species). The lower level of representation within *G. morsitans* probably results from the fact that it was assembled from sequence data derived from multiple older technologies using the now unsupported Celera assembly software [85].

**Repeat analysis and transposable element composition (Supplemental Figure 10 + Supplemental table 14)**

A comparative analysis was performed on the quantities and types of repetitive elements contained within the six tsetse genomes. The analysis reveals a similar content across the six genomes in terms of the number of consensus sequences and subclass diversity. The total percentage of masked repeats ranging from 20.87% (*G. brevipalpis*) to 27.65% (*G. pallidipes*) (Supplemental table 13). On average about 25% of the tsetse genomes are comprised of repeated elements consisting of simple repeats, tandem and satellite, low complexity DNAs and dispersed transposable elements (TEs). These elements mainly belong to three main subclasses (Long Interspersed Nuclear Elements - LINEs, Terminal Inverted Repeats – TIRs and Helitrons) in addition to unknown repetitive elements (Supplemental Figure 10A).

Although the overall number of elements was lower for *G. brevipalpis* in comparison to other *Glossina* species, the relative proportion of each subclass was comparable, with a diverse array of element types found within DNA and LINE subclasses. The *G. brevipalpis* genome was also found to contain the lowest overall repeat content, particularly so for LINEs; however, the opposite trend was observed for low complexity repeats, which had a greater number of simple repeats, which are likely micro-satellites.

A large proportion of yet unclassified repeats was observed, especially for *G. morsitans*. For all tsetse genomes, 3 subclasses of TEs predominate - DNA transposons (Class II TIRs) (Supplemental Figure 10B), Helitrons (Class II Rolling Circle -RC), and LINEs (Class I non LTR) (Supplemental Figure 10C). Other Class I elements such as LTR or SINEs are very scarce. However, the majority of LTR elements present in *Glossina* genomes are Gypsy elements. For DNA transposons, more than half correspond to the mariner element from the large super family of Tc1-mariner-Pogo. This family was also the most diversified.

Helitron elements are the most abundant in coverage. Helitrons are associated with long-term activity and persistent vertical transmission. These elements are thought to contribute to speciation and genome evolution due to their tendency to capture gene sequences during their duplication via rolling circle replication. These events can result in the generation of protogenes that form the basis of new gene families [91]. In this context, there are several noted *Glossina* specific gene expansions. These include gene families such as the milk protein genes [70] and male accessory protein genes (see below in the section on reproductive biology).

The total assembled repeats did not correlate with assembly contiguity measures, = in other words, high repeat content did not equate to lower assembly contiguity. Nonetheless, given highly repeat-rich regions are largely inaccessible to short read length sequencing and assembly methods, our approximation of repeat element content in *Glossina* is likely an underestimation and more detailed distribution and measures of transposable elements will require further experimentation.

**Functional enrichment of *Glossina* specific genes (Figure 5 + Supplemental table 3)**

*Morsitans sub-Genus Specific and Universal Genes*

Analysis of genes specific and universal to the *Morsitans* sub-Genus (containing *G. morsitans*, *G. pallidipes* and *G. austeni*) revealed enrichment in proteasome activating ATPase function. In addition to the protease enrichment observed in the Genus at large, this may be an adaptation to aid in protein digestion. Proteasome function is also linked with the process of autophagy [104]. Autophagy plays an important role in the process of lactation in tsetse. The milk gland tissue of female flies undergoes autophagic renewal at the end of each gonotrophic cycle. This process breaks down and restores the secretory cells in the gland immediately after larval deposition to ensure optimal function during the next lactation cycle [105].

**Sub-genus specific gene family expansions and contractions (Figure 6, Supplemental Table 4 and Supplemental figures** **11+12)**

*Palpalis sub-genus specific gene family variations*

A gene family encoding phosphoribosylformylglycinamidine synthases (VBGT00190000012418) is also increased in the *Palpalis* group. This enzyme, also known as *ade2* in *Drosophila*, is part of the *de novo* biosynthetic pathway for purine nucleotides. In tsetse, the function of this pathway is negatively impacted by the loss of the obligate *Wigglesworthia* symbiont and its associated B-vitamin cofactors, upon which nucleotide biosynthesis pathways rely [110, 111]. Purines in addition to being components of DNA act as essential enzymatic co-factors (ATP/GTP, S-adenosylmethionine). Amplification of an enzyme associated with *de novo* biosynthesis of purines reinforces the significance of their role in tsetse biology and may be an optimization of the pathway within the context of the symbiotic relationship with *Wigglesworthia*.

A family of thiamine (vitamin B1) dependent fatty acid alpha oxidases is also expanded within the *Palpalis* group (VBGT00190000011125). These enzymes function to remove single carbons from fatty acid chains in a thiamine dependent manner to facilitate beta-oxidation. They are also capable of acting on methylated fatty acids, which prevent beta-oxidation. Interestingly, the *Drosophila* ortholog to this gene is most highly expressed in the digestive tract of developing larvae, adult females and embryos. These enzymes could be important for larval milk digestion and are likely impacted by the thiamine deficiencies noted in *Wigglesworthia* free flies.

**Tsetse saliva genes (Figure 10)**

The analysis performed here encompasses five gene families encoding the following saliva protein classes: i) 5’nucleotidase-related proteins, ii) tsetse thrombin inhibitors, iii) tsetse salivary gland proteins, iv) adenosine-deaminases and v) the tsetse antigen-5 proteins. Comparison of these classes across these six tsetse fly species indicates that orthologues are present in all the genomes and that (in most cases) these sequences show sub-genus specific i.e. *Morsitans*, *Palpalis* and *Fusca* features. This agrees with previous findings documenting the differential sialome protein profiles of the respective sub-genera [127]. Sequences from *G. brevipalpis* show the strongest sequence divergence. This correlates with the *Fusca* group’s early evolutionary divergence in tsetse fly evolution.

In the host hemostatic reactions, ATP- and ADP-mediated platelet aggregation is essential and blood sucking insects have ATP/ADP-hydrolysing [ATP(D)ase; apyrase] enzymes present in their salivary secretions [128]. Each of the *Glossina* species examined here have three genes coding for proteins with a 5’-nucleotidase signature. These genes are located in a conserved 26 kb locus and include the *5’Nuc*, *sgp3* and *apyrase1-like* genes (Figure 10A). A second *apyrase2-like* gene is located on a separate scaffold.

The 5’Nuc orthologues are highly conserved across the species with similarities ranging from 92%-94% within the *Morsitans* group, 97% within the Palpalis species. Similarity comparison of these sequences relative to *G. brevipalpis* revealed sequence similarity of ~70%. Analysis of the predicted translation products of the representative sequences from the *Morsitans* and *Palpalis* groups identified the first 25 amino acids of these proteins as a putative signal peptide corresponding to the secretory nature of saliva proteins [128]. However, no signal peptide was detected for the *5’Nuc* genomic sequence of the *Fusca* group flies. All the amino acids predicted in *G. morsitans* to be involved in catalytic activity, co-factor and substrate binding [128] are conserved amongst all tsetse species, with the exception of a substrate binding amino acid Arg^358^ that is replaced by phenylalanine in *G. austeni*. It is not clear whether the substitution will have an impact on its apyrase activity, immunogenic properties or vector competence.

**Neuropeptides and protein hormone receptors (Supplemental Tables 16+17)**

*Neuropeptides*

Neuropeptides are amino acid-based molecules that can enter into the circulatory system and act as signalling molecules as well as neuromodulators and neurotransmitters within the central nervous system. Neuropeptides regulate key biological processes including metabolism and homeostasis, reproduction, growth and development, behaviour and feeding amongst others. Previous work identified 39 neuropeptide genes in *G. morsitans* including the conserved core neuropeptide genes found in all insect genomes, as well as a variable cohort of neuropeptide genes only found in some insect genomes [136].

The current analysis expands the scope of the previous analysis of neuropeptides [137] and their receptors by including the new tsetse fly species and *M. domestica*. The housefly differs from tsetse across multiple aspects of their biology including reproduction, development and feeding biology. This provides the opportunity to determine if these differences are reflected in the neuropeptide/receptor complement.

The core set of neuropeptides are present in most of the species examined (Supplementary Table 16). *G. brevipalpis* is an obvious outsider in this analysis – consistently showing the lowest protein identity to *G. morsitans* and missing a total of 6 neuropeptide genes (Allatostatin B, CCHamide-1, DH (calcitonin), ETH, Leucokinin and PDF). These genes are present in the other *Glossina* species.

The lack of certain neuropeptides in *G. morsitans* is reinforced by the consistent absence in the other *Glossina* species. The missing peptide genes include adipokinetic hormone/corazonin-like neuropeptide (ACP - function unknown), allatotropin (stimulation of juvenile hormone synthesis and cardiac activity), inotocin (control of water balance) [138] and sulfakinin (control of feeding and of larval locomotion/odor preferences) [136]. Indeed, almost none of the above neuropeptides are associated with the genomes of the other *Glossina* species or in *M. domestica*.

This analysis also identified a number of neuropeptide genes not previously located or confirmed in the *G. morsitans* genome [23]. These include: Myoinhibiting peptide (allatostatin B) and CCH amide 1 (possible modulator of sensory perception and olfactory behavior in starved *D. melanogaster*) [139] which were located in the genome of *M. domestica* and all the *Glossina* species with the exception of *G. brevipalpis*. The orcokinin gene (regulation of circadian rhythms) is present in all the *Glossina* species but missing in *M. domestica*. In addition, some recently discovered neuropeptide families were included in the current gene searches, and so natalisin (regulation of reproductive behavior and fecundity) [140] and trissin (function unknown) are now documented in all six *Glossina* species.

*Neuropeptide Receptors*

Most neuropeptides carry out their function by activating specific G-protein coupled receptors (GPCRs), guanylate cyclase receptors and receptor tyrosine kinases [141]. We annotated over 40 GPCRs in the *Glossina* genome databases (Supplemental Table 17), including orphan receptors with as yet unknown ligand(s). We confirm here that the vasopressin/oxytocin related neuropeptide, inotocin, and its receptor were lost during dipteran evolution as this system is also absent in all other dipteran genomes investigated so far. The allatotropin, ACP neuropeptides and their receptors are also absent in the *Glossina* species, *M. domestica* and *Drosophila*, but occur in mosquito genomes, pointing to a more recent loss of these neuropeptidergic systems in the Schizophora clade. The sulfakinin receptor is missing in all the *Glossina* species suggesting evolutionary loss of this signaling system in tsetse flies possibly due to the evolution of tsetse’s blood feeding behaviour/biology.

**Cuticular Proteins (Supplemental Table 18)**

Query of the *Glossina* genomes with sequence motifs characteristic of cuticle protein families revealed 584 genes encoding putative cuticle proteins. Each species represented between 101-130 genes which were assigned to one of eight families (CPR, CPAP1, CPAP3, CPF, CPCFC, CPLCA, CPLCG, and TWDL; Supplemental data 6). The total number of cuticle protein genes identified in *Glossina* is within the range observed in other insects (Supplemental table 14), but less than that of other dipterans [79, 142]. As with other insects, the number of CPR genes (60-85) constituted the largest group of cuticle protein genes in the *Glossina* genomes; however, this number is two- to fourfold less than in other dipterans and partially accounted for the reduced number of total cuticle protein genes. In general, the number of genes in each cuticle protein family did not correlate with evolutionary relatedness in the dipterans. *Glossina* had a similar number of genes in the CPAP1 and TWDL families as the mosquito species (Culicidae), whereas the number of CPLCG genes was more similar to the other Brachycera flies (Supplemental table 14). This variability in gene numbers is evident in the TWDL (Tweedle) family, which shows an expansion in the dipterans [79]. The TWDL genes tended to group by taxa within Diptera and gene expansions are evident in *Musca domestica*, *Drosophila melanogaster*, *Ceratitis capitata*; however, *Glossina* lack these large expansions. Further study is needed to determine if this difference is due to multiple, independent events of gene duplication within brachyceran flies, or if *Glossina* lost genes that were present in a common ancestor. Cuticular protein gene expansions likely reflects adaptive evolution [143]. The lack of expansion in *Glossina* may reflect their unique strategy of intrauterine larval development. The relative lack of cuticle protein synthesis during the first two larval instars of *G. morsitans* [144] further suggests that the protection of the mother limits the need for extensive cuticle proteins. However, there is currently a lack in understanding of the precise role of these protein families, and the functional implications of their reduction requires further elucidation.

**Transcription factors**

To compare repertoires of predicted DNA binding transcriptional regulatory proteins we searched the proteins from the tsetse genomes for putative DNA binding domains. These proteins were then classified according to the class of binding domain to provide a picture of the relative abundance of these important regulatory proteins. The analysis identified a total of between 700 (*G. morsitans*) and 806 (*G. fuscipes*) putative TFs in the *Glossina* genomes, which is similar to other insect genomes (e.g., 701 for *D. melanogaster*). Likewise, for the most part, the number of members of each *Glossina* TF family is comparable to that of other insects (Supplemental Figure 13).

We next inferred DNA binding motifs for these TFs using a previously described procedure [80] (Supplemental data 7). For the following, we use *G. pallidipes*, which has the median number of TFs among the six species, as an example. Similar results were obtained for the other species. Of the 783 *G. pallidipes* TFs, we were able to infer motifs for 266 (34%) (Supplemental data 6), mostly based on DNA binding specificity data from *D. melanogaster* (215 TFs), but also from species as distant as mouse (3 TFs) and zebrafish (2 TFs). Many of the largest TF families have inferred motifs for a substantial proportion of their TFs, including Homeodomain (63 of 89, 71%), bHLH (47 of 52, 90%), and nuclear receptors (10 of 18, 56%). As expected, the largest gap is for C2H2 zinc fingers (only 32 of 270, ~12%), which evolve quickly by shuffling their many zinc finger arrays, resulting in largely dissimilar DBD sequences across organisms [145]. This catalog of TFs and their predicted DNA binding motifs offers an opportunity to begin examining and comparing gene regulatory networks in the *Glossina* genomes

**Peritrophic Matrix (Supplemental table 19)**

Peritrophins and peritrophin-like proteins (PLPs) [131, 132], typified by the presence of chitin-binding domains (CBDs), are important structural components of the peritrophic matrix (PM), a semi-permeable barrier separating the lumen from the midgut epithelial cells [133]. The insect PM is involved in regulating immunity genes and appear to act as a physical barrier to pathogens [134, 135].

Comparative analyses of peritrophins and PLPs across all six major *Glossina* species show a high degree of conservation with slight differences in the numbers and characteristics of these proteins. *Glossina* have less than half the number of peritrophins and PLPs relative to *Musca, Stomoxys* and *Drosophila*, with these latter Dipterans having similar numbers of genes to various mosquito species. The higher number of genes found in these species could reflect the major differences in the feeding habits between them and *Glossina*. The digestive tracts of *Glossina* are adapted to a restricted diet of lactation secretions and blood, while other Dipteran species have much more diverse diets and greater exposure to dietary pathogens and hazards. The streamlining of peritrophins in *Glossina* could potentially explain their susceptibility to infection by trypanosomes.
