## Supplemental Table 1 Statistics on gene and scaffold mapping to Muller elements for "The *Glossina* Genome Cluster: Comparative Genomic Analysis of the Vectors of African Trypanosomes"

| Species | genes | mapped genes | % genes mapped | scaffolds | mapped scaffolds | % scaffolds mapped |
| --- | --- | --- | --- | --- | --- | --- |
| *G. austeni* | 19,747 | 6,829 | 34.58% | 671 | 650 | 96.87% |
| *G. brevipalpis* | 14,641 | 6,450 | 44.05% | 448 | 435 | 97.10% |
| *G. fuscipes* | 20,138 | 6,487 | 32.21% | 793 | 764 | 96.34% |
| *G. morsitans* | 12,442 | 6,430 | 51.68% | 1,773 | 1,703 | 96.05% |
| *G. pallidipes* | 19,297 | 6,519 | 33.78% | 544 | 528 | 97.06% |
| *G. palpalis* | 20,160 | 6,322 | 31.36% | 918 | 884 | 96.30% |
