## Supplemental table 2 Wolbachia sequences for "The *Glossina* Genome Cluster: Comparative Genomic Analysis of the Vectors of African Trypanosomes"

**Supplemental table 2.** *Wolbachia* sequences identified in *G. morsitans*, *G. austeni*, *G. fuscipes*, *G. pallidipes*, *G. brevipalpis*, and *G. palpalis*

|  | Cytoplasmic | Chromosomal |
| --- | --- | --- |
| *G. morsitans* |  | 1,013,719 bp |
| *G. pallidipes* | 2,580 bp | No |
| *G. austeni* | 90,846 bp | 960,436 bp |
| *G. palpalis* | 1,050 bp | No |
| *G. fuscipes* | 14,228 bp | No |
| *G. brevipalpis* | 55,476 bp | No |
