## Supplemental Table 3 Gene Ontology (GO Molecular Function) Term Enrichment for "The *Glossina* Genome Cluster: Comparative Genomic Analysis of the Vectors of African Trypanosomes"

**Supplemental Table 3: Gene Ontology (GO Molecular Function) Term Enrichment in the *Glossina* Genus and associated sub-Genera**

| **GO.ID** | **Term** | **Annotated** | **Significant** | **Expected** | **Rank in classic Fisher** | **Classic Fisher** | **Elim Fisher** | **Topgo Fisher** | **Parentchild Fisher** |
| --- | --- | --- | --- | --- | --- | --- | --- | --- | --- |
| **Glossina Specific and Universal Enriched GO Terms (Molecular Function)** | | | | | | | | | |
| ***G. morsitans*** | | | | | | | | | |
| GO:0005549 | odorant binding | 78 | 8 | 0.49 | 1 | 2.10E-08 | 2.10E-08 | 2.10E-08 | 1.10E-08 |
| GO:0004252 | serine-type endopeptidase activity | 145 | 7 | 0.91 | 3 | 2.90E-05 | 2.90E-05 | 2.90E-05 | 0.1626 |
| GO:0004190 | aspartic-type endopeptidase activity | 16 | 2 | 0.1 | 8 | 0.0044 | 0.0044 | 0.0044 | 0.1108 |
| ***G. pallidipes*** | | | | | | | | | |
| GO:0004252 | serine-type endopeptidase activity | 180 | 16 | 1 | 1 | 3.10E-14 | 3.10E-14 | 3.10E-14 | 0.00062 |
| GO:0005549 | odorant binding | 69 | 7 | 0 | 7 | 4.00E-07 | 4.00E-07 | 4.00E-07 | 6.20E-08 |
| GO:0002162 | dystroglycan binding | 1 | 1 | 0 | 10 | 0.0069 | 0.0069 | 0.0069 | 0.00438 |
| ***G. austeni*** | | | | | | | | | |
| GO:0004252 | serine-type endopeptidase activity | 178 | 17 | 1.24 | 2 | 1.60E-15 | 1.60E-15 | 1.60E-15 | 0.0065 |
| GO:0005549 | odorant binding | 75 | 8 | 0.52 | 7 | 3.90E-08 | 3.90E-08 | 3.90E-08 | 6.90E-09 |
| GO:0004190 | aspartic-type endopeptidase activity | 34 | 4 | 0.24 | 10 | 8.40E-05 | 8.40E-05 | 8.40E-05 | 0.1745 |
| ***G. fuscipes*** | | | | | | | | | |
| GO:0004252 | serine-type endopeptidase activity | 179 | 19 | 1.09 | 1 | 1.60E-19 | 1.60E-19 | 1.60E-19 | 1.50E-05 |
| GO:0005549 | odorant binding | 79 | 11 | 0.48 | 5 | 8.60E-13 | 8.60E-13 | 8.60E-13 | 2.20E-14 |
| GO:0002162 | dystroglycan binding | 1 | 1 | 0.01 | 10 | 0.0061 | 0.0061 | 0.0061 | 0.0024 |
| ***G. palpalis*** | | | | | | | | | |
| GO:0004252 | serine-type endopeptidase activity | 177 | 20 | 1.18 | 1 | 2.90E-20 | 2.90E-20 | 2.90E-20 | 5.10E-05 |
| GO:0005549 | odorant binding | 77 | 9 | 0.51 | 7 | 1.40E-09 | 1.40E-09 | 1.40E-09 | 7.20E-11 |
| GO:0003678 | DNA helicase activity | 18 | 2 | 0.12 | 10 | 0.0062 | 0.0062 | 0.0062 | 0.028 |
| ***G. brevipalpis*** | | | | | | | | | |
| GO:0005549 | odorant binding | 81 | 12 | 0.54 | 1 | 8.50E-14 | 8.50E-14 | 8.50E-14 | 1.00E-14 |
| GO:0004252 | serine-type endopeptidase activity | 163 | 13 | 1.09 | 2 | 2.30E-11 | 2.30E-11 | 2.30E-11 | 0.003 |
| GO:0008036 | diuretic hormone receptor activity | 2 | 2 | 0.01 | 8 | 4.30E-05 | 4.30E-05 | 4.30E-05 | 4.00E-05 |
| **Morsitans sub-genus specific and universal enriched GO Terms (Molecular Function)** | | | | | | | | | |
| ***G. morsitans*** | | | | | | | | | |
| GO:0036402 | proteasome-activating ATPase activity | 10 | 2 | 0.01 | 1 | 6.90E-05 | 6.90E-05 | 6.90E-05 | 0.011 |
| ***G. pallidipes*** | | | | | | | | | |
| GO:0036402 | proteasome-activating ATPase activity | 9 | 2 | 0.01 | 1 | 2.70E-05 | 2.70E-05 | 2.70E-05 | 0.0074 |
| ***G. austeni*** | | | | | | | | | |
| GO:0036402 | proteasome-activating ATPase activity | 11 | 3 | 0.02 | 1 | 3.10E-07 | 3.10E-07 | 3.10E-07 | 0.00076 |
| **Palpalis sub-Genus Specific and Universal Enriched GO Terms (Molecular Function)** | | | | | | | | | |
| ***G. fuscipes*** | | | | | | | | | |
| GO:0003677 | DNA binding | 696 | 23 | 11.59 | 4 | 0.00107 | 1.07E-03 | 0.00015 | 0.0476 |
| GO:0004672 | protein kinase activity | 248 | 11 | 4.13 | 13 | 0.00278 | 0.00278 | 0.00048 | 0.0018 |
| GO:0008026 | ATP-dependent helicase activity | 14 | 3 | 0.23 | 7 | 0.00144 | 0.00144 | 0.0016 | 0.0018 |
| ***G. palpalis*** | | | | | | | | | |
| GO:0004672 | protein kinase activity | 265 | 14 | 4.68 | 9 | 0.00023 | 0.00023 | 5.50E-06 | 0.00011 |
| GO:0003678 | DNA helicase activity | 18 | 3 | 0.32 | 25 | 0.00362 | 0.00362 | 0.00362 | 0.06133 |
| GO:0008026 | ATP-dependent helicase activity | 12 | 2 | 0.21 | 35 | 0.0182 | 0.0182 | 0.0182 | 0.01478 |
