## Supplemental Table 4 sub Genus Specific Gene Expansions and Contractions for "The *Glossina* Genome Cluster: Comparative Genomic Analysis of the Vectors of African Trypanosomes"

**Supplemental Table 4:** Overview of sub-Genus Specific Gene Expansions and Contractions (Numbers represent the number of paralogous sequences per species)

| **VectorBase Gene Tree ID Number** | **Functional Definition** | **Sub-Genus** | **Expansion or Contraction** | ***Glossina brevipalpis*** | ***Glossina austeni*** | ***Glossina morsitans*** | ***Glossina pallidipes*** | ***Glossina palpalis*** | ***Glossina fuscipes*** | **CAFE Derived Family-wide P-value** |
| --- | --- | --- | --- | --- | --- | --- | --- | --- | --- | --- |
| VBGT00190000009793 | PREDICTED: 116 kDa U5 small nuclear ribonucleoprotein component [Bactrocera oleae] | Fusca | Contraction | 5 | 11 | 7 | 8 | 8 | 12 | 0.001 |
| VBGT00190000009794 | PREDICTED: laccase-5-like isoform X3 [Bombus impatiens] | Fusca | Contraction | 4 | 8 | 5 | 7 | 5 | 7 | 0.011 |
| VBGT00190000009808 | V-type proton ATPase catalytic subunit A isoform 2 [Lucilia cuprina] | Fusca | Contraction | 6 | 11 | 9 | 11 | 9 | 12 | 0.011 |
| VBGT00190000010180 | PREDICTED: zinc finger MIZ domain-containing protein 2 isoform X1 [Stomoxys calcitrans] | Fusca | Contraction | 2 | 3 | 5 | 6 | 4 | 6 | 0.004 |
| VBGT00190000011642 | PREDICTED: kinesin-like protein KIF20B isoform X1 [Bactrocera oleae] | Fusca | Contraction | 6 | 13 | 11 | 17 | 13 | 13 | 0.004 |
| VBGT00190000014894 | hypothetical protein FF38_05426 [Lucilia cuprina] | Fusca | Contraction | 1 | 6 | 4 | 4 | 5 | 4 | 0.031 |
| VBGT00190000015811 | hypothetical protein FF38_05834 [Lucilia cuprina] | Fusca | Contraction | 1 | 4 | 5 | 6 | 3 | 6 | 0.004 |
| VBGT00190000016440 | PREDICTED: uncharacterized protein LOC105227462 [Bactrocera dorsalis] | Fusca | Contraction | 2 | 6 | 4 | 6 | 5 | 6 | 0.049 |
| VBGT00190000016667 | hypothetical protein FF38_03619 [Lucilia cuprina] | Fusca | Contraction | 1 | 5 | 2 | 7 | 2 | 4 | 0 |
| VBGT00190000016741 | PREDICTED: C3 and PZP-like alpha-2-macroglobulin domain-containing protein 8 [Musca domestica] | Fusca | Contraction | 1 | 5 | 3 | 5 | 4 | 3 | 0.001 |
| VBGT00750000029331 | hypothetical protein FF38_06312 [Lucilia cuprina] | Fusca | Contraction | 1 | 4 | 3 | 5 | 5 | 5 | 0.028 |
| VBGT00770000031214 | PREDICTED: zinc finger protein 699-like [Stomoxys calcitrans] | Fusca | Contraction | 10 | 18 | 18 | 18 | 19 | 16 | 0.039 |
| VBGT00820000045940 | PREDICTED: transitional endoplasmic reticulum ATPase TER94 isoform X1 [Bactrocera oleae] | Fusca | Contraction | 20 | 25 | 25 | 21 | 24 | 21 | 0.045 |
| VBGT00820000045950 | PREDICTED: activating signal cointegrator 1 complex subunit 3 [Musca domestica] | Fusca | Contraction | 7 | 13 | 12 | 16 | 15 | 9 | 0 |
| VBGT00820000046004 | PREDICTED: dnaJ homolog subfamily A member 4 [Musca domestica] | Fusca | Contraction | 13 | 19 | 16 | 17 | 15 | 19 | 0.02 |
| VBGT00820000046040 | hypothetical protein FF38_07185 [Lucilia cuprina] | Fusca | Contraction | 2 | 4 | 5 | 6 | 5 | 7 | 0.013 |
| VBGT00840000047885 | Peroxisomal multifunctional enzyme type 2 [Lucilia cuprina] | Fusca | Contraction | 7 | 11 | 12 | 9 | 8 | 11 | 0.004 |
| VBGT00190000009711 | PREDICTED: aldose reductase-like [Musca domestica] | Fusca | Expansion | 16 | 10 | 8 | 12 | 9 | 11 | 0.008 |
| VBGT00190000009892 | PREDICTED: ER degradation-enhancing alpha-mannosidase-like protein 3 [Musca domestica] | Fusca | Expansion | 11 | 7 | 6 | 4 | 4 | 5 | 0.011 |
| VBGT00190000012881 | PREDICTED: GATA zinc finger domain-containing protein 11 isoform X2 [Bactrocera cucurbitae] | Fusca | Expansion | 13 | 7 | 4 | 7 | 7 | 7 | 0.013 |
| VBGT00190000010664 | trypsin-like serine protease precursor [Glossina morsitans morsitans] | Morsitans | Contraction | 15 | 5 | 7 | 7 | 14 | 15 | 0.001 |
| VBGT00820000045989 | putative cytochrome P450 28d1 [Lucilia cuprina] | Morsitans | Contraction | 11 | 7 | 5 | 7 | 14 | 17 | 0 |
| VBGT00190000009926 | PREDICTED: lysine-specific demethylase lid [Musca domestica] | Morsitans | Expansion | 4 | 7 | 7 | 6 | 4 | 3 | 0.036 |
| VBGT00190000010724 | putative cysteine desulfurase, mitochondrial [Lucilia cuprina] | Morsitans | Expansion | 1 | 4 | 4 | 4 | 1 | 1 | 0.041 |
| VBGT00190000010913 | PREDICTED: cytochrome c oxidase subunit 4 isoform 1, mitochondrial [Stomoxys calcitrans] | Morsitans | Expansion | 2 | 5 | 5 | 3 | 1 | 2 | 0.001 |
| VBGT00190000010926 | PREDICTED: farnesyl pyrophosphate synthase [Musca domestica] | Morsitans | Expansion | 1 | 5 | 5 | 3 | 1 | 1 | 0.004 |
| VBGT00190000011677 | PREDICTED: RUS1 family protein C16orf58 homolog [Musca domestica] | Morsitans | Expansion | 1 | 5 | 2 | 4 | 1 | 1 | 0.004 |
| VBGT00840000047886 | PREDICTED: dehydrogenase/reductase SDR family protein 7-like [Musca domestica] | Morsitans | Expansion | 7 | 10 | 11 | 10 | 8 | 11 | 0.028 |
| VBGT00190000009725 | Argonaute-1, isoform A [Drosophila melanogaster] | Palpalis | Contraction | 13 | 8 | 12 | 12 | 7 | 8 | 0.025 |
| VBGT00190000009849 | PREDICTED: valine--tRNA ligase isoform X2 [Ceratitis capitata] | Palpalis | Expansion | 4 | 7 | 5 | 4 | 7 | 9 | 0.01 |
| VBGT00190000009945 | PREDICTED: serine palmitoyltransferase 1 [Stomoxys calcitrans] | Palpalis | Expansion | 4 | 6 | 3 | 5 | 12 | 8 | 0 |
| VBGT00190000010218 | TM2 domain-containing protein almondex [Lucilia cuprina] | Palpalis | Expansion | 1 | 4 | 3 | 3 | 7 | 10 | 0 |
| VBGT00190000010265 | PREDICTED: probable phosphorylase b kinase regulatory subunit alpha isoform X7 [Stomoxys calcitrans] | Palpalis | Expansion | 2 | 2 | 3 | 2 | 6 | 7 | 0.017 |
| VBGT00190000010375 | PREDICTED: mismatch repair endonuclease PMS2 [Musca domestica] | Palpalis | Expansion | 3 | 2 | 2 | 2 | 6 | 7 | 0.043 |
| VBGT00190000010512 | Zinc transporter foi, partial [Lucilia cuprina] | Palpalis | Expansion | 1 | 2 | 2 | 3 | 5 | 9 | 0 |
| VBGT00190000010757 | PREDICTED: midasin [Musca domestica] | Palpalis | Expansion | 1 | 4 | 1 | 1 | 7 | 7 | 0 |
| VBGT00190000010980 | Transcription-associated protein 1 [Lucilia cuprina] | Palpalis | Expansion | 4 | 2 | 2 | 7 | 17 | 13 | 0 |
| VBGT00190000011125 | PREDICTED: 2-hydroxyacyl-CoA lyase 1 [Stomoxys calcitrans] | Palpalis | Expansion | 1 | 1 | 1 | 1 | 7 | 6 | 0.004 |
| VBGT00190000011176 | PREDICTED: pre-mRNA-splicing factor Slu7 [Stomoxys calcitrans] | Palpalis | Expansion | 1 | 2 | 2 | 2 | 11 | 12 | 0 |
| VBGT00190000011233 | PREDICTED: cytoplasmic FMR1-interacting protein [Musca domestica] | Palpalis | Expansion | 1 | 1 | 2 | 1 | 6 | 6 | 0.003 |
| VBGT00190000011761 | PREDICTED: replication protein A 70 kDa DNA-binding subunit [Stomoxys calcitrans] | Palpalis | Expansion | 1 | 3 | 2 | 1 | 5 | 6 | 0.001 |
| VBGT00190000011829 | PREDICTED: phosphopantothenoylcysteine decarboxylase [Musca domestica] | Palpalis | Expansion | 1 | 1 | 1 | 1 | 5 | 6 | 0.015 |
| VBGT00190000012100 | PREDICTED: integrator complex subunit 4 [Musca domestica] | Palpalis | Expansion | 1 | 1 | 1 | 2 | 3 | 5 | 0.004 |
| VBGT00190000012301 | Phosphoglycerate kinase [Lucilia cuprina] | Palpalis | Expansion | 3 | 3 | 1 | 4 | 5 | 9 | 0 |
| VBGT00190000012418 | PREDICTED: phosphoribosylformylglycinamidine synthase [Stomoxys calcitrans] | Palpalis | Expansion | 1 | 3 | 1 | 2 | 5 | 4 | 0.003 |
| VBGT00190000012451 | PREDICTED: SWI/SNF complex subunit SMARCC2 isoform X2 [Stomoxys calcitrans] | Palpalis | Expansion | 2 | 2 | 2 | 3 | 6 | 7 | 0.017 |
| VBGT00190000013378 | hypothetical protein FF38_01314 [Lucilia cuprina] | Palpalis | Expansion | 1 | 1 | 1 | 1 | 4 | 6 | 0.009 |
| VBGT00190000013689 | PREDICTED: ATP-dependent DNA helicase 2 subunit 1 [Stomoxys calcitrans] | Palpalis | Expansion | 4 | 3 | 2 | 2 | 5 | 7 | 0.011 |
| VBGT00190000013845 | hypothetical protein FF38_01981 [Lucilia cuprina] | Palpalis | Expansion | 1 | 1 | 1 | 1 | 7 | 5 | 0.003 |
| VBGT00190000014141 | PREDICTED: WD repeat and HMG-box DNA-binding protein 1 [Stomoxys calcitrans] | Palpalis | Expansion | 1 | 1 | 1 | 1 | 5 | 7 | 0.003 |
| VBGT00190000014373 | PREDICTED: cilia- and flagella-associated protein 44 [Stomoxys calcitrans] | Palpalis | Expansion | 2 | 1 | 1 | 1 | 14 | 10 | 0 |
| VBGT00190000014659 | PREDICTED: transmembrane protein 131 homolog [Stomoxys calcitrans] | Palpalis | Expansion | 1 | 1 | 1 | 1 | 3 | 6 | 0.003 |
| VBGT00770000031191 | PREDICTED: WD repeat-containing protein 78 [Musca domestica] | Palpalis | Expansion | 4 | 4 | 3 | 4 | 10 | 15 | 0 |
| VBGT00770000031281 | PREDICTED: recombination repair protein 1 isoform X1 [Musca domestica] | Palpalis | Expansion | 1 | 1 | 1 | 1 | 5 | 4 | 0.043 |
| VBGT00770000031399 | Nuclear pore complex protein Nup153 [Lucilia cuprina] | Palpalis | Expansion | 3 | 3 | 1 | 3 | 4 | 6 | 0.003 |
| VBGT00780000038281 | PREDICTED: cell division cycle protein 27 homolog [Stomoxys calcitrans] | Palpalis | Expansion | 2 | 3 | 2 | 2 | 14 | 12 | 0 |
| VBGT00840000047928 | hypothetical protein FF38_09159, partial [Lucilia cuprina] | Palpalis | Expansion | 3 | 4 | 2 | 3 | 8 | 8 | 0.008 |
