## Supplemental Table 5 Count of Glossina orthologs to Immune Activity Associated Drosophila genes for "The *Glossina* Genome Cluster: Comparative Genomic Analysis of the Vectors of African Trypanosomes"

**Supplemental Table 5: Count of Glossina orthologs to Immune Activity Associated *Drosophila* genes.** Table contains Drosophila genes associated with the Immune Process GO term GO:0002376 and counts of orthologus sequences in *Glossina* genomes.

| Drosophila Immune Associated Gene Name | D. melanogaster | G. austeni | G. brevipalpis | G. fuscipes | G. morsitans | G. pallidipes | G. palpalis |
| --- | --- | --- | --- | --- | --- | --- | --- |
| 18 wheeler | 1 | 1 | 1 | 1 | 1 | 1 | 1 |
| Abl tyrosine kinase | 1 | 1 | 1 | 2 | 0 | 0 | 1 |
| achaete | 1 | 1 | 0 | 0 | 0 | 1 | 1 |
| Activated Cdc42 kinase | 1 | 1 | 1 | 1 | 1 | 1 | 1 |
| Activated Cdc42 kinase-like | 1 | 1 | 1 | 1 | 1 | 1 | 1 |
| Activating transcription factor 3 | 1 | 1 | 1 | 1 | 1 | 1 | 1 |
| Activating transcription factor-2 | 1 | 1 | 1 | 1 | 1 | 1 | 1 |
| Activator of SUMO 1 | 1 | 1 | 1 | 1 | 1 | 1 | 1 |
| Adenosine deaminase-related growth factor A | 1 | 1 | 1 | 1 | 1 | 1 | 1 |
| ADP ribosylation factor at 79F | 1 | 1 | 1 | 1 | 1 | 1 | 1 |
| akirin | 1 | 2 | 1 | 1 | 2 | 3 | 1 |
| Akt1 | 1 | 1 | 1 | 1 | 1 | 2 | 1 |
| alpha-Mannosidase class I a | 1 | 0 | 0 | 0 | 0 | 0 | 0 |
| alpha-Mannosidase class I b | 1 | 2 | 2 | 2 | 3 | 0 | 1 |
| alpha-Mannosidase class II a | 1 | 1 | 1 | 1 | 1 | 1 | 1 |
| alpha-Mannosidase class II b | 1 | 0 | 1 | 1 | 1 | 1 | 1 |
| alpha-Tubulin at 84B | 1 | 1 | 1 | 0 | 1 | 1 | 1 |
| Amyotrophic lateral sclerosis 2 | 1 | 1 | 1 | 1 | 1 | 1 | 1 |
| Anaplastic lymphoma kinase | 1 | 1 | 1 | 1 | 0 | 1 | 1 |
| Andropin | 1 | 0 | 0 | 0 | 0 | 0 | 0 |
| Another transcription unit | 1 | 1 | 1 | 1 | 1 | 1 | 1 |
| Antennapedia | 1 | 1 | 1 | 1 | 0 | 1 | 1 |
| anterior open | 1 | 1 | 1 | 1 | 1 | 1 | 0 |
| Argonaute 2 | 1 | 3 | 3 | 3 | 4 | 5 | 3 |
| asrij | 1 | 1 | 1 | 1 | 1 | 1 | 1 |
| Attacin-A | 1 | 5 | 3 | 3 | 4 | 4 | 2 |
| Attacin-B | 1 | 5 | 3 | 3 | 4 | 4 | 2 |
| Attacin-C | 1 | 0 | 0 | 0 | 0 | 0 | 0 |
| atypical protein kinase C | 1 | 1 | 1 | 1 | 0 | 1 | 1 |
| aubergine | 1 | 1 | 4 | 1 | 1 | 1 | 1 |
| Autophagy-related 18a | 1 | 1 | 1 | 2 | 1 | 1 | 1 |
| Autophagy-related 6 | 1 | 1 | 0 | 1 | 1 | 1 | 1 |
| Autophagy-related 7 | 1 | 3 | 2 | 2 | 1 | 1 | 3 |
| B52 | 1 | 2 | 2 | 2 | 2 | 1 | 2 |
| basket | 1 | 1 | 1 | 2 | 1 | 1 | 1 |
| big bang | 1 | 1 | 1 | 1 | 1 | 4 | 1 |
| Big brother | 1 | 2 | 1 | 1 | 1 | 1 | 1 |
| bip1 | 1 | 1 | 1 | 1 | 1 | 1 | 1 |
| brahma | 1 | 1 | 1 | 1 | 1 | 1 | 1 |
| Brother | 1 | 0 | 0 | 0 | 0 | 0 | 0 |
| Btk family kinase at 29A | 1 | 1 | 1 | 1 | 0 | 1 | 1 |
| cactus | 1 | 5 | 4 | 5 | 5 | 4 | 4 |
| Cadherin 99C | 1 | 1 | 1 | 1 | 1 | 1 | 1 |
| Calcineurin A1 | 1 | 1 | 1 | 1 | 1 | 1 | 1 |
| Calmodulin-binding protein related to a Rab3 GDP/GTP exchange protein | 1 | 1 | 1 | 1 | 1 | 1 | 1 |
| caspar | 1 | 1 | 1 | 1 | 1 | 1 | 1 |
| Catalase | 1 | 1 | 1 | 1 | 1 | 1 | 1 |
| caudal | 1 | 1 | 1 | 1 | 0 | 1 | 1 |
| Cdc42 | 1 | 1 | 2 | 1 | 1 | 0 | 1 |
| Cdk5 activator-like protein | 1 | 1 | 1 | 1 | 1 | 1 | 1 |
| Cecropin A1 | 1 | 2 | 1 | 4 | 2 | 1 | 1 |
| Cecropin A2 | 1 | 2 | 1 | 4 | 2 | 1 | 1 |
| Cecropin B | 1 | 2 | 1 | 4 | 2 | 1 | 1 |
| Cecropin C | 1 | 2 | 1 | 4 | 2 | 1 | 1 |
| CG10333 | 1 | 1 | 1 | 1 | 1 | 1 | 1 |
| CG10764 | 1 | 0 | 0 | 0 | 0 | 0 | 0 |
| CG1129 | 1 | 1 | 1 | 1 | 0 | 1 | 1 |
| CG11313 | 1 | 1 | 1 | 1 | 1 | 1 | 1 |
| CG11459 | 1 | 0 | 0 | 0 | 0 | 0 | 0 |
| CG12780 | 1 | 1 | 1 | 0 | 0 | 0 | 0 |
| CG13559 | 1 | 0 | 0 | 1 | 1 | 0 | 1 |
| CG13994 | 1 | 1 | 1 | 1 | 1 | 1 | 2 |
| CG14132 | 1 | 1 | 0 | 1 | 1 | 1 | 0 |
| CG15199 | 1 | 1 | 0 | 1 | 1 | 1 | 0 |
| CG15202 | 1 | 2 | 2 | 2 | 2 | 2 | 2 |
| CG15203 | 1 | 2 | 1 | 1 | 2 | 2 | 1 |
| CG15529 | 1 | 1 | 0 | 1 | 1 | 1 | 1 |
| CG1572 | 1 | 1 | 1 | 1 | 1 | 1 | 1 |
| CG1667 | 1 | 1 | 1 | 1 | 1 | 1 | 1 |
| CG1840 | 1 | 0 | 1 | 0 | 1 | 0 | 0 |
| CG32795 | 1 | 1 | 1 | 1 | 1 | 1 | 1 |
| CG33155 | 1 | 0 | 0 | 0 | 0 | 0 | 0 |
| CG33223 | 1 | 0 | 0 | 0 | 0 | 0 | 0 |
| CG34215 | 1 | 3 | 0 | 4 | 2 | 4 | 4 |
| CG4036 | 1 | 0 | 1 | 1 | 2 | 1 | 1 |
| CG42259 | 1 | 1 | 0 | 1 | 1 | 1 | 1 |
| CG42339 | 1 | 1 | 1 | 1 | 1 | 1 | 2 |
| CG42359 | 1 | 1 | 1 | 1 | 1 | 1 | 1 |
| CG43055 | 1 | 0 | 8 | 1 | 0 | 1 | 1 |
| CG4306 | 1 | 1 | 1 | 1 | 1 | 1 | 1 |
| CG4325 | 1 | 10 | 10 | 10 | 5 | 11 | 13 |
| CG4793 | 1 | 0 | 0 | 0 | 0 | 0 | 0 |
| CG5033 | 1 | 2 | 1 | 1 | 1 | 1 | 0 |
| CG5641 | 1 | 1 | 1 | 1 | 1 | 1 | 1 |
| CG5721 | 1 | 1 | 1 | 1 | 1 | 1 | 1 |
| CG8149 | 1 | 1 | 1 | 1 | 1 | 1 | 1 |
| CG9008 | 1 | 1 | 1 | 1 | 1 | 1 | 1 |
| CG9636 | 1 | 1 | 1 | 1 | 1 | 1 | 0 |
| CG9925 | 1 | 3 | 1 | 1 | 0 | 1 | 1 |
| Charon | 1 | 1 | 1 | 1 | 2 | 0 | 1 |
| cheerio | 1 | 2 | 1 | 1 | 1 | 1 | 2 |
| chico | 1 | 1 | 1 | 1 | 1 | 1 | 1 |
| chromosome bows | 1 | 1 | 1 | 2 | 1 | 2 | 1 |
| Coat Protein (coatomer) beta | 1 | 1 | 1 | 1 | 1 | 1 | 2 |
| connector enhancer of ksr | 1 | 0 | 0 | 0 | 0 | 0 | 0 |
| COP9 signalosome subunit 5 | 1 | 2 | 1 | 2 | 2 | 3 | 2 |
| Core 1 Galactosyltransferase A | 1 | 1 | 1 | 1 | 1 | 1 | 1 |
| couch potato | 1 | 0 | 0 | 0 | 0 | 0 | 0 |
| croquemort | 1 | 1 | 0 | 1 | 1 | 1 | 1 |
| C-terminal Src kinase | 1 | 1 | 1 | 1 | 1 | 1 | 1 |
| CTP synthase | 1 | 1 | 1 | 1 | 1 | 1 | 1 |
| cubitus interruptus | 1 | 1 | 0 | 2 | 0 | 1 | 1 |
| Cuticular protein 49Ac | 1 | 1 | 1 | 1 | 1 | 1 | 1 |
| Cyclin-dependent kinase 5 | 1 | 1 | 1 | 1 | 0 | 1 | 1 |
| Cyclin-dependent kinase 9 | 1 | 1 | 2 | 1 | 0 | 1 | 1 |
| Cylindromatosis | 1 | 1 | 1 | 1 | 1 | 1 | 1 |
| Cytochrome b5 | 1 | 1 | 1 | 1 | 1 | 1 | 1 |
| Darkener of apricot | 1 | 1 | 1 | 1 | 1 | 1 | 1 |
| daughter of sevenless | 1 | 1 | 1 | 1 | 1 | 1 | 1 |
| Death regulator Nedd2-like caspase | 1 | 1 | 1 | 1 | 1 | 1 | 1 |
| Death related ced-3/Nedd2-like caspase | 1 | 1 | 1 | 1 | 1 | 1 | 1 |
| Death-associated APAF1-related killer | 1 | 2 | 1 | 1 | 1 | 1 | 1 |
| Death-associated inhibitor of apoptosis 2 | 1 | 1 | 1 | 1 | 1 | 1 | 1 |
| decapentaplegic | 1 | 1 | 1 | 1 | 1 | 1 | 1 |
| defense repressor 1 | 1 | 1 | 1 | 1 | 1 | 1 | 1 |
| Defensin | 1 | 0 | 0 | 0 | 0 | 0 | 0 |
| Deformed epidermal autoregulatory factor-1 | 1 | 1 | 1 | 1 | 1 | 1 | 1 |
| Dephosphocoenzyme A carrier | 1 | 1 | 1 | 1 | 1 | 0 | 1 |
| diaphanous | 1 | 1 | 2 | 1 | 1 | 1 | 1 |
| Dicer-2 | 1 | 1 | 3 | 2 | 1 | 1 | 1 |
| Diedel | 1 | 0 | 0 | 0 | 0 | 0 | 0 |
| Diptericin A | 1 | 0 | 0 | 0 | 0 | 0 | 0 |
| Diptericin B | 1 | 0 | 0 | 0 | 0 | 0 | 0 |
| Dis3 | 1 | 2 | 1 | 1 | 3 | 2 | 2 |
| DISCO Interacting Protein 1 | 1 | 1 | 2 | 1 | 1 | 1 | 1 |
| Discs large 5 | 1 | 1 | 1 | 1 | 0 | 1 | 2 |
| dissatisfaction | 1 | 1 | 1 | 1 | 1 | 1 | 1 |
| DNA methyltransferase 1 associated protein 1 | 1 | 1 | 1 | 1 | 1 | 1 | 1 |
| DnaJ-like-1 | 1 | 2 | 1 | 2 | 2 | 3 | 2 |
| domeless | 1 | 3 | 3 | 3 | 3 | 3 | 4 |
| domino | 1 | 1 | 1 | 1 | 1 | 1 | 1 |
| dorsal | 1 | 1 | 1 | 1 | 1 | 1 | 1 |
| Dorsal interacting protein 3 | 1 | 1 | 1 | 2 | 1 | 1 | 2 |
| Dorsal switch protein 1 | 1 | 2 | 2 | 2 | 1 | 2 | 3 |
| Dorsal-related immunity factor | 1 | 1 | 1 | 1 | 1 | 1 | 0 |
| double parked | 1 | 2 | 2 | 2 | 4 | 2 | 3 |
| Doublecortin-domain-containing echinoderm-microtubule-associated protein | 1 | 2 | 1 | 1 | 1 | 1 | 1 |
| Downstream of raf1 | 1 | 1 | 1 | 0 | 0 | 1 | 1 |
| downstream of receptor kinase | 1 | 1 | 1 | 0 | 1 | 0 | 1 |
| Drosocin | 1 | 0 | 0 | 0 | 0 | 0 | 0 |
| Drosomycin | 1 | 0 | 0 | 0 | 0 | 0 | 0 |
| Dual oxidase | 1 | 1 | 1 | 1 | 2 | 1 | 1 |
| Dynactin 1, p150 subunit | 1 | 1 | 1 | 1 | 1 | 1 | 2 |
| Dynactin 3, p24 subunit | 1 | 1 | 1 | 2 | 1 | 1 | 1 |
| Dynein heavy chain 64C | 1 | 1 | 1 | 1 | 1 | 1 | 1 |
| E2F transcription factor 1 | 1 | 1 | 1 | 1 | 1 | 1 | 1 |
| eater | 1 | 3 | 1 | 1 | 1 | 1 | 1 |
| Eb1 | 1 | 3 | 1 | 6 | 3 | 3 | 7 |
| Ecdysone receptor | 1 | 1 | 1 | 1 | 1 | 1 | 1 |
| Ecdysone-induced gene 71Ee | 1 | 0 | 0 | 0 | 0 | 0 | 0 |
| Ecdysone-induced protein 75B | 1 | 1 | 1 | 1 | 1 | 1 | 1 |
| ecdysoneless | 1 | 1 | 1 | 1 | 1 | 1 | 1 |
| ECSIT | 1 | 1 | 1 | 3 | 2 | 1 | 3 |
| eiger | 1 | 0 | 0 | 0 | 0 | 0 | 0 |
| elevated during infection | 1 | 0 | 0 | 0 | 0 | 0 | 0 |
| Elongator complex protein 3 | 1 | 1 | 1 | 0 | 2 | 1 | 2 |
| Enhancer of bithorax | 1 | 1 | 1 | 0 | 1 | 1 | 1 |
| Eph receptor tyrosine kinase | 1 | 1 | 1 | 1 | 1 | 1 | 2 |
| Ephexin | 1 | 1 | 1 | 1 | 0 | 1 | 1 |
| Ephrin | 1 | 1 | 1 | 1 | 1 | 1 | 1 |
| Epidermal growth factor receptor | 1 | 1 | 1 | 1 | 1 | 1 | 1 |
| Essential MCU regulator | 1 | 0 | 0 | 0 | 1 | 0 | 1 |
| Etl1 | 1 | 1 | 1 | 1 | 1 | 1 | 1 |
| eyes absent | 1 | 1 | 1 | 1 | 1 | 1 | 1 |
| Fas-associated death domain | 1 | 1 | 1 | 1 | 1 | 1 | 1 |
| fat facets | 1 | 1 | 1 | 1 | 1 | 1 | 1 |
| fates-shifted | 1 | 1 | 1 | 1 | 1 | 1 | 1 |
| FER tyrosine kinase | 1 | 1 | 1 | 1 | 1 | 1 | 1 |
| Ferritin 2 light chain homologue | 1 | 1 | 1 | 1 | 1 | 1 | 1 |
| Flotillin 2 | 1 | 2 | 4 | 1 | 1 | 1 | 1 |
| Focal adhesion kinase | 1 | 2 | 0 | 1 | 0 | 1 | 1 |
| fondue | 1 | 1 | 1 | 1 | 1 | 1 | 1 |
| Forkhead box K | 1 | 0 | 0 | 1 | 1 | 1 | 0 |
| forkhead box, sub-group O | 1 | 1 | 2 | 1 | 1 | 1 | 1 |
| frizzled | 1 | 1 | 0 | 1 | 1 | 1 | 1 |
| frizzled 2 | 1 | 1 | 1 | 1 | 1 | 1 | 1 |
| G protein alpha f subunit | 1 | 1 | 1 | 1 | 1 | 1 | 1 |
| G protein alpha q subunit | 1 | 1 | 1 | 1 | 1 | 1 | 2 |
| G protein-coupled receptor kinase 2 | 1 | 1 | 1 | 1 | 1 | 1 | 1 |
| G9a | 1 | 0 | 1 | 1 | 1 | 1 | 1 |
| Galactose-specific C-type lectin | 1 | 0 | 8 | 1 | 0 | 1 | 1 |
| GATAe | 1 | 1 | 1 | 1 | 1 | 1 | 1 |
| gcm2 | 1 | 1 | 1 | 1 | 1 | 1 | 1 |
| glial cells missing | 1 | 1 | 1 | 1 | 1 | 1 | 1 |
| Glutathione S transferase O2 | 1 | 1 | 1 | 1 | 1 | 1 | 1 |
| Glycogen phosphorylase | 1 | 1 | 1 | 1 | 1 | 1 | 1 |
| Glycyl-tRNA synthetase | 1 | 1 | 1 | 1 | 1 | 1 | 1 |
| Gram-negative bacteria binding protein 1 | 1 | 1 | 1 | 1 | 2 | 1 | 1 |
| Gram-negative bacteria binding protein 2 | 1 | 0 | 0 | 0 | 0 | 0 | 0 |
| Gram-negative bacteria binding protein 3 | 1 | 0 | 0 | 0 | 1 | 1 | 1 |
| Gram-positive Specific Serine protease | 1 | 1 | 1 | 1 | 1 | 1 | 1 |
| grim | 1 | 0 | 0 | 0 | 0 | 0 | 0 |
| Growth-blocking peptide 1 | 1 | 0 | 0 | 0 | 0 | 0 | 0 |
| Gustatory receptor 28b | 1 | 2 | 3 | 1 | 1 | 2 | 1 |
| Hand | 1 | 1 | 1 | 1 | 1 | 1 | 1 |
| Hayan | 1 | 1 | 1 | 1 | 1 | 1 | 1 |
| HBS1 | 1 | 1 | 1 | 1 | 1 | 1 | 1 |
| heartless | 1 | 1 | 1 | 1 | 1 | 1 | 2 |
| HECT and RLD domain containing E3 ubiquitin ligase 4 | 1 | 1 | 1 | 1 | 1 | 1 | 1 |
| hedgehog | 1 | 1 | 1 | 1 | 1 | 1 | 1 |
| heixuedian | 1 | 1 | 1 | 1 | 1 | 1 | 1 |
| Helical Factor | 1 | 0 | 0 | 0 | 0 | 0 | 0 |
| Helicase 89B | 1 | 1 | 1 | 0 | 1 | 1 | 1 |
| Hemese | 1 | 0 | 0 | 0 | 0 | 0 | 0 |
| hemipterous | 1 | 0 | 1 | 1 | 0 | 1 | 0 |
| Hemolectin | 1 | 0 | 1 | 1 | 1 | 1 | 1 |
| Histone acetyltransferase 1 | 1 | 1 | 1 | 1 | 1 | 1 | 1 |
| hopscotch | 1 | 1 | 2 | 1 | 1 | 1 | 1 |
| Hormone-receptor-like in 78 | 1 | 1 | 0 | 0 | 1 | 1 | 1 |
| I-kappaB kinase beta | 1 | 1 | 1 | 1 | 1 | 0 | 1 |
| I-kappaB kinase epsilon | 1 | 1 | 1 | 2 | 1 | 1 | 3 |
| immune deficiency | 1 | 1 | 1 | 1 | 1 | 0 | 1 |
| Immune induced molecule 23 | 1 | 0 | 0 | 0 | 0 | 0 | 0 |
| Immune induced molecule 3 | 1 | 0 | 0 | 0 | 0 | 0 | 0 |
| Immune induced molecule 4 | 1 | 0 | 0 | 0 | 0 | 0 | 0 |
| Immune induced molecule prepropeptide | 1 | 0 | 0 | 0 | 0 | 0 | 0 |
| immune response deficient 10 | 1 | 0 | 0 | 0 | 0 | 0 | 0 |
| immune response deficient 21 | 1 | 0 | 0 | 0 | 0 | 0 | 0 |
| immune response deficient 24 | 1 | 0 | 0 | 0 | 0 | 0 | 0 |
| immune response deficient 25 | 1 | 0 | 0 | 0 | 0 | 0 | 0 |
| immune response deficient 26 | 1 | 0 | 0 | 0 | 0 | 0 | 0 |
| immune response deficient 27 | 1 | 0 | 0 | 0 | 0 | 0 | 0 |
| immune response deficient 6 | 1 | 0 | 0 | 0 | 0 | 0 | 0 |
| immune response deficient 8 | 1 | 0 | 0 | 0 | 0 | 0 | 0 |
| immune response deficient 9 | 1 | 0 | 0 | 0 | 0 | 0 | 0 |
| inflated | 1 | 1 | 1 | 1 | 1 | 1 | 1 |
| Insulin degrading metalloproteinase | 1 | 1 | 1 | 2 | 2 | 2 | 3 |
| Integrin betanu subunit | 1 | 1 | 0 | 1 | 1 | 1 | 1 |
| Jun-related antigen | 1 | 0 | 1 | 1 | 2 | 1 | 1 |
| kayak | 1 | 1 | 1 | 1 | 1 | 1 | 1 |
| kenny | 1 | 1 | 1 | 2 | 1 | 1 | 1 |
| kinase suppressor of ras | 1 | 1 | 1 | 1 | 1 | 1 | 2 |
| kismet | 1 | 1 | 1 | 1 | 1 | 1 | 2 |
| knot | 1 | 1 | 1 | 1 | 1 | 1 | 2 |
| Kruppel | 1 | 1 | 1 | 1 | 1 | 1 | 1 |
| kugelei | 1 | 1 | 1 | 1 | 1 | 1 | 1 |
| kurtz | 1 | 1 | 1 | 1 | 1 | 1 | 1 |
| kuzbanian | 1 | 1 | 1 | 1 | 1 | 1 | 1 |
| Lamin | 1 | 2 | 3 | 2 | 1 | 2 | 2 |
| Laminin A | 1 | 1 | 1 | 1 | 1 | 1 | 1 |
| LanB1 | 1 | 1 | 1 | 1 | 1 | 1 | 0 |
| lesswright | 1 | 1 | 1 | 1 | 1 | 2 | 1 |
| lethal (2) 34Fc | 1 | 0 | 1 | 0 | 1 | 0 | 0 |
| lethal (2) k06524 | 1 | 0 | 0 | 0 | 0 | 0 | 0 |
| lethal (2) k08634 | 1 | 0 | 0 | 0 | 0 | 0 | 0 |
| Leucine-rich repeat | 1 | 1 | 1 | 1 | 1 | 1 | 1 |
| licorne | 1 | 1 | 1 | 1 | 1 | 1 | 1 |
| Listericin | 1 | 0 | 0 | 0 | 0 | 0 | 0 |
| lola like | 1 | 1 | 1 | 1 | 1 | 1 | 1 |
| longitudinals lacking | 1 | 1 | 1 | 1 | 1 | 1 | 1 |
| lozenge | 1 | 1 | 2 | 1 | 0 | 1 | 1 |
| Lysozyme E | 1 | 0 | 0 | 0 | 0 | 0 | 0 |
| male-specific lethal 3 | 1 | 1 | 1 | 1 | 1 | 1 | 1 |
| Mannosyl (alpha-1,3-)-glycoprotein beta-1,2-N-acetylglucosaminyltransferase | 1 | 1 | 1 | 1 | 1 | 1 | 1 |
| maternal expression at 31B | 1 | 1 | 1 | 1 | 1 | 1 | 1 |
| Matrix metalloproteinase 2 | 1 | 1 | 1 | 2 | 1 | 2 | 2 |
| Mediator complex subunit 25 | 1 | 1 | 1 | 1 | 1 | 1 | 1 |
| Mediator complex subunit 6 | 1 | 1 | 1 | 1 | 1 | 1 | 1 |
| Mekk1 | 1 | 1 | 1 | 1 | 1 | 1 | 1 |
| Melanization Protease 1 | 1 | 1 | 1 | 1 | 1 | 1 | 1 |
| members only | 1 | 1 | 2 | 1 | 1 | 1 | 1 |
| Metchnikowin | 1 | 0 | 0 | 0 | 0 | 0 | 0 |
| Methyltransferase 2 | 1 | 1 | 1 | 1 | 1 | 1 | 1 |
| midline fasciclin | 1 | 1 | 1 | 3 | 1 | 1 | 3 |
| Mig-2-like | 1 | 2 | 1 | 1 | 1 | 1 | 1 |
| mir-959 stem loop | 1 | 0 | 0 | 0 | 0 | 0 | 0 |
| mir-960 stem loop | 1 | 0 | 0 | 0 | 0 | 0 | 0 |
| mir-961 stem loop | 1 | 0 | 0 | 0 | 0 | 0 | 0 |
| mir-962 stem loop | 1 | 0 | 0 | 0 | 0 | 0 | 0 |
| mir-963 stem loop | 1 | 0 | 0 | 0 | 0 | 0 | 0 |
| mir-964 stem loop | 1 | 0 | 0 | 0 | 0 | 0 | 0 |
| Mitf | 1 | 0 | 1 | 1 | 1 | 1 | 1 |
| mitochondrial ribosomal protein L53 | 1 | 1 | 1 | 1 | 1 | 1 | 1 |
| Mitogen-activated protein kinase phosphatase 3 | 1 | 1 | 0 | 1 | 1 | 1 | 1 |
| modular serine protease | 1 | 0 | 0 | 0 | 0 | 0 | 0 |
| moira | 1 | 2 | 2 | 7 | 2 | 3 | 6 |
| Mtr4 helicase | 1 | 1 | 6 | 2 | 1 | 2 | 1 |
| multi sex combs | 1 | 1 | 1 | 1 | 0 | 1 | 1 |
| multiple ankyrin repeats single KH domain | 1 | 1 | 1 | 1 | 1 | 1 | 0 |
| mustard | 1 | 1 | 1 | 1 | 1 | 1 | 1 |
| Myd88 | 1 | 1 | 2 | 1 | 1 | 1 | 1 |
| Myelodysplasia/myeloid leukemia factor | 1 | 1 | 1 | 1 | 1 | 1 | 1 |
| Myocyte enhancer factor 2 | 1 | 1 | 1 | 0 | 2 | 1 | 1 |
| myopic | 1 | 1 | 1 | 1 | 1 | 1 | 2 |
| myospheroid | 1 | 1 | 1 | 1 | 2 | 1 | 1 |
| necrotic | 1 | 1 | 1 | 1 | 1 | 1 | 0 |
| nejire | 1 | 1 | 1 | 2 | 1 | 1 | 3 |
| neuralized | 1 | 1 | 1 | 1 | 1 | 1 | 1 |
| Neuroglian | 1 | 1 | 1 | 1 | 1 | 1 | 1 |
| Neuropeptide-like precursor 2 | 1 | 0 | 0 | 0 | 0 | 0 | 0 |
| Niemann-Pick type C-2g | 1 | 1 | 1 | 1 | 1 | 1 | 1 |
| Niemann-Pick type C-2h | 1 | 1 | 1 | 1 | 1 | 1 | 1 |
| no receptor potential A | 1 | 1 | 0 | 1 | 1 | 0 | 1 |
| Noa36 | 1 | 1 | 1 | 1 | 0 | 0 | 1 |
| Nop2/Sun-like domain containing protein 5 | 1 | 1 | 1 | 1 | 1 | 1 | 1 |
| Notch | 1 | 1 | 1 | 1 | 1 | 1 | 2 |
| NTF2-related export protein 1 | 1 | 1 | 1 | 2 | 1 | 1 | 2 |
| nubbin | 1 | 1 | 1 | 1 | 1 | 1 | 1 |
| Nuclear transport factor-2 | 1 | 1 | 1 | 1 | 2 | 2 | 1 |
| Nucleoporin 98-96kD | 1 | 1 | 1 | 2 | 2 | 2 | 1 |
| Nucleosome remodeling factor - 38kD | 1 | 2 | 1 | 1 | 1 | 2 | 1 |
| Oligosaccharide transferase Delta subunit | 1 | 1 | 1 | 2 | 1 | 1 | 2 |
| Oligosaccharide transferase gamma subunit | 1 | 1 | 1 | 1 | 1 | 1 | 1 |
| over compensating males | 1 | 1 | 1 | 1 | 1 | 1 | 1 |
| overgrown hematopoietic organs 51 | 1 | 0 | 0 | 0 | 0 | 0 | 0 |
| overgrown hematopoietic organs 55DE | 1 | 0 | 0 | 0 | 0 | 0 | 0 |
| p38a MAP kinase | 1 | 1 | 1 | 1 | 1 | 1 | 1 |
| p38b MAP kinase | 1 | 1 | 1 | 1 | 1 | 1 | 1 |
| pangolin | 1 | 2 | 1 | 1 | 1 | 1 | 1 |
| pannier | 1 | 1 | 1 | 1 | 1 | 1 | 1 |
| par-1 | 1 | 1 | 1 | 1 | 1 | 1 | 1 |
| PAX transcription activation domain interacting protein | 1 | 1 | 1 | 1 | 1 | 0 | 1 |
| PDGF- and VEGF-receptor related | 1 | 1 | 1 | 1 | 0 | 1 | 1 |
| PDGF- and VEGF-related factor 1 | 1 | 1 | 1 | 1 | 1 | 1 | 1 |
| PDGF- and VEGF-related factor 2 | 1 | 0 | 0 | 0 | 0 | 0 | 0 |
| PDGF- and VEGF-related factor 3 | 1 | 1 | 1 | 1 | 1 | 1 | 2 |
| PDZ domain-containing guanine nucleotide exchange factor | 1 | 1 | 1 | 1 | 1 | 1 | 1 |
| pelle | 1 | 1 | 2 | 1 | 1 | 1 | 1 |
| Pellino | 1 | 1 | 1 | 1 | 1 | 1 | 1 |
| Pendulin | 1 | 2 | 2 | 2 | 2 | 4 | 2 |
| Peptidoglycan recognition protein LA | 1 | 1 | 0 | 1 | 1 | 1 | 1 |
| Peptidoglycan recognition protein LB | 1 | 1 | 1 | 1 | 1 | 1 | 1 |
| Peptidoglycan recognition protein LC | 1 | 1 | 0 | 0 | 1 | 1 | 0 |
| Peptidoglycan recognition protein LD | 1 | 1 | 1 | 1 | 0 | 0 | 1 |
| Peptidoglycan recognition protein LE | 1 | 0 | 0 | 0 | 0 | 0 | 0 |
| Peptidoglycan recognition protein LF | 1 | 1 | 1 | 1 | 0 | 0 | 1 |
| Peptidoglycan recognition protein SA | 1 | 1 | 1 | 1 | 1 | 1 | 2 |
| Peptidoglycan recognition protein SB1 | 1 | 1 | 1 | 1 | 0 | 1 | 1 |
| Peptidoglycan recognition protein SB2 | 1 | 0 | 0 | 0 | 0 | 0 | 0 |
| Peptidoglycan recognition protein SC1a | 1 | 0 | 0 | 0 | 0 | 0 | 0 |
| Peptidoglycan recognition protein SC1b | 1 | 0 | 0 | 0 | 0 | 0 | 0 |
| Peptidoglycan recognition protein SC2 | 1 | 0 | 0 | 0 | 0 | 0 | 0 |
| Peptidoglycan recognition protein SD | 1 | 0 | 0 | 0 | 0 | 0 | 0 |
| Peroxiredoxin 5 | 1 | 1 | 2 | 2 | 3 | 1 | 1 |
| persephone | 1 | 1 | 1 | 1 | 1 | 1 | 1 |
| phagocyte signaling impaired | 1 | 1 | 1 | 1 | 1 | 1 | 1 |
| Phosphatase and tensin homolog | 1 | 1 | 1 | 1 | 1 | 1 | 1 |
| Phosphatidylethanolamine-binding protein 1 | 1 | 0 | 0 | 0 | 0 | 0 | 1 |
| Pi3K92E | 1 | 1 | 1 | 1 | 1 | 1 | 1 |
| Pitslre | 1 | 0 | 1 | 1 | 0 | 0 | 0 |
| Plenty of SH3s | 1 | 1 | 1 | 1 | 1 | 1 | 1 |
| pointed | 1 | 1 | 1 | 1 | 1 | 1 | 0 |
| poly | 1 | 1 | 0 | 0 | 1 | 1 | 1 |
| Poly-(ADP-ribose) polymerase | 1 | 1 | 1 | 2 | 1 | 1 | 2 |
| poor Imd response upon knock-in | 1 | 0 | 1 | 1 | 0 | 0 | 1 |
| Programmed Cell Death 5 | 1 | 1 | 1 | 0 | 0 | 1 | 2 |
| Proliferating cell nuclear antigen | 1 | 4 | 1 | 1 | 1 | 1 | 1 |
| proliferation disrupter | 1 | 1 | 1 | 1 | 1 | 1 | 1 |
| Prophenoloxidase 1 | 1 | 1 | 1 | 1 | 1 | 1 | 1 |
| Prophenoloxidase 2 | 1 | 3 | 3 | 3 | 3 | 2 | 1 |
| Prophenoloxidase 3 | 1 | 3 | 3 | 3 | 3 | 2 | 1 |
| Proteasome alpha6 subunit, Testis-specific | 1 | 1 | 1 | 1 | 1 | 1 | 1 |
| Protein C kinase 53E | 1 | 1 | 1 | 1 | 0 | 1 | 1 |
| Protein phosphatase 1alpha at 96A | 1 | 1 | 1 | 1 | 1 | 2 | 1 |
| puckered | 1 | 1 | 0 | 1 | 1 | 0 | 1 |
| puffyeye | 1 | 1 | 1 | 1 | 1 | 1 | 1 |
| pxb | 1 | 1 | 1 | 1 | 1 | 1 | 1 |
| r2d2 | 1 | 2 | 2 | 3 | 2 | 2 | 2 |
| Rab11 | 1 | 2 | 2 | 2 | 2 | 2 | 2 |
| Rac1 | 1 | 1 | 1 | 0 | 1 | 1 | 1 |
| Rac2 | 1 | 1 | 1 | 1 | 1 | 1 | 1 |
| Raf oncogene | 1 | 1 | 1 | 1 | 1 | 1 | 1 |
| Rap1 GTPase | 1 | 0 | 0 | 0 | 0 | 0 | 0 |
| Ras oncogene at 85D | 1 | 1 | 0 | 1 | 1 | 1 | 1 |
| Ras-like protein A | 1 | 1 | 1 | 1 | 1 | 1 | 1 |
| Relish | 1 | 1 | 1 | 1 | 1 | 1 | 1 |
| Ret oncogene | 1 | 1 | 1 | 2 | 1 | 1 | 2 |
| Rho guanine nucleotide exchange factor 3 | 1 | 1 | 1 | 1 | 1 | 1 | 1 |
| Rho1 | 1 | 1 | 1 | 1 | 1 | 1 | 1 |
| Rho-like | 1 | 1 | 1 | 1 | 1 | 1 | 1 |
| Rho-related BTB domain containing | 1 | 1 | 1 | 1 | 1 | 1 | 1 |
| Ribosomal protein S21 | 1 | 1 | 1 | 1 | 1 | 1 | 1 |
| Ribosomal protein S6 | 1 | 1 | 2 | 1 | 1 | 1 | 1 |
| Ribosomal protein S8 | 1 | 1 | 1 | 2 | 1 | 1 | 2 |
| Ribosomal RNA processing 40 | 1 | 0 | 2 | 0 | 0 | 0 | 0 |
| Ring and YY1 Binding Protein | 1 | 1 | 1 | 1 | 1 | 1 | 1 |
| RluA pseudouridine synthase 2 | 1 | 0 | 0 | 0 | 0 | 0 | 0 |
| Rm62 | 1 | 1 | 1 | 1 | 1 | 1 | 1 |
| rolled | 1 | 1 | 1 | 1 | 1 | 1 | 1 |
| Rrp4 | 1 | 1 | 1 | 1 | 1 | 0 | 1 |
| Rrp6 | 1 | 1 | 1 | 1 | 1 | 1 | 1 |
| Ryanodine receptor | 1 | 1 | 1 | 1 | 1 | 1 | 1 |
| sallimus | 1 | 1 | 1 | 1 | 1 | 1 | 0 |
| saxophone | 1 | 1 | 1 | 1 | 1 | 1 | 1 |
| Scavenger receptor class C, type I | 1 | 1 | 1 | 1 | 0 | 1 | 1 |
| Scavenger receptor class C, type II | 1 | 1 | 1 | 1 | 0 | 1 | 1 |
| Scavenger receptor class C, type IV | 1 | 1 | 1 | 1 | 0 | 1 | 1 |
| scrawny | 1 | 1 | 0 | 1 | 1 | 1 | 1 |
| Secreted Wg-interacting molecule | 1 | 1 | 1 | 1 | 1 | 1 | 1 |
| Secretory 31 | 1 | 1 | 1 | 1 | 1 | 1 | 1 |
| Secretory 5 | 1 | 1 | 1 | 1 | 1 | 1 | 1 |
| Selenoprotein G | 1 | 0 | 1 | 0 | 1 | 0 | 0 |
| senju | 1 | 0 | 1 | 0 | 0 | 0 | 0 |
| senseless-2 | 1 | 1 | 1 | 1 | 1 | 1 | 1 |
| Serine protease 7 | 1 | 1 | 1 | 1 | 1 | 2 | 1 |
| Serine Protease Immune Response Integrator | 1 | 2 | 1 | 2 | 2 | 2 | 2 |
| serpent | 1 | 0 | 0 | 0 | 0 | 0 | 0 |
| Serpin 27A | 1 | 1 | 1 | 1 | 1 | 1 | 1 |
| Serpin 28Dc | 1 | 2 | 1 | 1 | 1 | 1 | 1 |
| Serpin 42Dd | 1 | 0 | 0 | 0 | 0 | 0 | 0 |
| Serpin 77Ba | 1 | 0 | 1 | 0 | 1 | 1 | 1 |
| Serrate | 1 | 1 | 1 | 1 | 1 | 1 | 2 |
| SH2 ankyrin repeat kinase | 1 | 1 | 1 | 2 | 1 | 1 | 4 |
| shaggy | 1 | 1 | 1 | 1 | 0 | 1 | 1 |
| SHC-adaptor protein | 1 | 1 | 1 | 1 | 1 | 1 | 1 |
| shotgun | 1 | 1 | 1 | 1 | 1 | 1 | 1 |
| Signal-transducer and activator of transcription protein at 92E | 1 | 2 | 2 | 3 | 4 | 3 | 2 |
| similar | 1 | 1 | 0 | 1 | 1 | 1 | 1 |
| Sin3A | 1 | 1 | 1 | 1 | 1 | 1 | 1 |
| singed | 1 | 1 | 1 | 1 | 1 | 1 | 1 |
| Ski6 | 1 | 1 | 0 | 1 | 2 | 2 | 1 |
| slimfast | 1 | 2 | 2 | 2 | 2 | 2 | 2 |
| Small ribonucleoprotein particle protein SmD3 | 1 | 1 | 1 | 1 | 1 | 1 | 1 |
| smog | 1 | 1 | 1 | 1 | 1 | 1 | 1 |
| smoothened | 1 | 1 | 1 | 1 | 1 | 3 | 1 |
| smt3 | 1 | 0 | 1 | 0 | 0 | 0 | 1 |
| Son of sevenless | 1 | 1 | 1 | 2 | 1 | 1 | 2 |
| SP2353 | 1 | 2 | 1 | 4 | 1 | 1 | 3 |
| spastin | 1 | 1 | 2 | 1 | 1 | 1 | 1 |
| spatzle | 1 | 2 | 2 | 4 | 1 | 3 | 2 |
| Spatzle-Processing Enzyme | 1 | 1 | 1 | 1 | 0 | 1 | 0 |
| spheroide | 1 | 0 | 0 | 1 | 0 | 0 | 1 |
| sphinx1 | 1 | 0 | 0 | 0 | 0 | 0 | 0 |
| sphinx2 | 1 | 0 | 0 | 0 | 0 | 0 | 0 |
| Spt6 | 1 | 1 | 3 | 1 | 1 | 1 | 1 |
| Src oncogene at 42A | 1 | 2 | 0 | 1 | 1 | 1 | 1 |
| Src oncogene at 64B | 1 | 1 | 1 | 1 | 1 | 1 | 1 |
| stonewall | 1 | 1 | 1 | 0 | 0 | 0 | 1 |
| Superoxide dismutase 2 (Mn) | 1 | 1 | 1 | 1 | 0 | 1 | 1 |
| Suppressor of Hairless | 1 | 1 | 1 | 2 | 1 | 1 | 2 |
| Suppressor of variegation 2-10 | 1 | 1 | 1 | 4 | 3 | 3 | 2 |
| Syntaxin 5 | 1 | 1 | 1 | 1 | 1 | 1 | 1 |
| tailup | 1 | 1 | 1 | 1 | 1 | 1 | 1 |
| TAK1-associated binding protein 2 | 1 | 1 | 0 | 1 | 0 | 1 | 2 |
| Tehao | 1 | 0 | 0 | 0 | 0 | 0 | 0 |
| telomere fusion | 1 | 1 | 1 | 1 | 1 | 0 | 1 |
| Tetraspanin 68C | 1 | 1 | 0 | 1 | 1 | 1 | 1 |
| TGF-beta activated kinase 1 | 1 | 1 | 1 | 1 | 1 | 1 | 1 |
| Thor | 1 | 1 | 1 | 1 | 1 | 1 | 1 |
| tinman | 1 | 1 | 1 | 1 | 1 | 1 | 2 |
| TNF-receptor-associated factor 6 | 1 | 1 | 1 | 1 | 1 | 1 | 1 |
| Toll | 1 | 1 | 1 | 1 | 1 | 1 | 1 |
| Toll-7 | 1 | 1 | 1 | 1 | 1 | 1 | 1 |
| Toll-9 | 1 | 1 | 0 | 1 | 1 | 1 | 1 |
| Tollo | 1 | 1 | 1 | 1 | 1 | 1 | 1 |
| Tousled-like kinase | 1 | 1 | 1 | 1 | 1 | 1 | 1 |
| Transglutaminase | 1 | 1 | 1 | 1 | 1 | 1 | 1 |
| Translocase of inner membrane 10 | 1 | 0 | 1 | 0 | 1 | 1 | 1 |
| Transmembrane 9 superfamily protein member 4 | 1 | 1 | 1 | 1 | 1 | 1 | 1 |
| Transportin | 1 | 1 | 1 | 1 | 1 | 1 | 2 |
| tube | 1 | 2 | 1 | 1 | 1 | 2 | 1 |
| Tyrosyl-tRNA synthetase | 1 | 1 | 1 | 1 | 1 | 1 | 1 |
| Ubiquitin activating enzyme 2 | 1 | 2 | 3 | 3 | 2 | 3 | 2 |
| Ubiquitin conjugating enzyme 7 | 1 | 1 | 1 | 1 | 0 | 1 | 1 |
| ubiquitin like | 1 | 1 | 0 | 1 | 1 | 0 | 1 |
| Ubiquitin specific protease 2 | 1 | 1 | 1 | 1 | 1 | 0 | 1 |
| Ubr3 ubiquitin ligase | 1 | 1 | 1 | 1 | 1 | 1 | 2 |
| UDP-galactose 4'-epimerase | 1 | 1 | 0 | 1 | 1 | 1 | 1 |
| Ulp1 | 1 | 4 | 3 | 4 | 5 | 4 | 3 |
| uncoordinated 45 | 1 | 1 | 1 | 1 | 2 | 1 | 1 |
| unpaired 1 | 1 | 1 | 1 | 1 | 1 | 1 | 1 |
| unpaired 3 | 1 | 1 | 1 | 1 | 1 | 1 | 1 |
| u-shaped | 1 | 1 | 1 | 1 | 1 | 1 | 1 |
| Vacuolar protein sorting 15 | 1 | 2 | 2 | 1 | 1 | 2 | 1 |
| Vacuolar protein sorting 16B | 1 | 1 | 1 | 1 | 1 | 1 | 1 |
| Vacuolar protein sorting 33B | 1 | 1 | 0 | 1 | 1 | 1 | 0 |
| Vago | 1 | 1 | 1 | 1 | 1 | 1 | 1 |
| Vav guanine nucleotide exchange factor | 1 | 1 | 1 | 1 | 1 | 1 | 1 |
| veloren | 1 | 4 | 2 | 1 | 3 | 3 | 2 |
| ventral veins lacking | 1 | 1 | 1 | 1 | 1 | 1 | 1 |
| virus-induced RNA 1 | 1 | 3 | 1 | 2 | 1 | 2 | 2 |
| wingless | 1 | 2 | 1 | 1 | 1 | 1 | 2 |
| wispy | 1 | 1 | 1 | 1 | 1 | 1 | 1 |
| Wnt oncogene analog 4 | 1 | 1 | 1 | 1 | 1 | 1 | 1 |
| Wsck | 1 | 1 | 1 | 1 | 1 | 1 | 1 |
| wunen | 1 | 1 | 1 | 1 | 2 | 1 | 1 |
| yantar | 1 | 1 | 1 | 1 | 0 | 1 | 1 |
| Zinc finger CCHC-type containing 7 | 1 | 1 | 1 | 1 | 1 | 1 | 1 |
| Zinc finger protein RP-8 | 1 | 1 | 1 | 1 | 1 | 1 | 1 |
| Zinc-finger protein | 1 | 1 | 1 | 0 | 1 | 2 | 0 |
| Zizimin-related | 1 | 1 | 1 | 1 | 1 | 1 | 1 |
| Zn finger homeodomain 1 | 1 | 1 | 1 | 1 | 1 | 1 | 1 |
