## Supplemental Table 6 Glossina Milk Protein Genes for "The *Glossina* Genome Cluster: Comparative Genomic Analysis of the Vectors of African Trypanosomes"

| Gene ID | Species | Milk Protein | Scaffold |
| --- | --- | --- | --- |
| GAUT009632 | *G. austeni* | mgp1 | Scaffold14:1466112-1467216:1 |
| GAUT032292 | *G. austeni* | transferrin | Scaffold3:2777421-2780032:-1 |
| GAUT007910 | *G. austeni* | acid sphingomyelinase | Scaffold137:624881-628940:-1 |
| GAUT052949 | *G. austeni* | Mgp2 | Scaffold45:761783:762385:-1 |
| GAUT035464 | *G. austeni* | Mgp3 | Scaffold45:769597:770234:1 |
| GAUT035467 | *G. austeni* | Mgp4 | Scaffold45:758397:758897:1 |
| GAUT035461 | *G. austeni* | Mgp5 | Scaffold45:746356:747009:1 |
| GAUT035463 | *G. austeni* | Mgp6 | Scaffold45:763332:763925:1 |
| GAUT035459 | *G. austeni* | Mgp7 | Scaffold45:736728:737546:1 |
| GAUT035466 | *G. austeni* | Mgp8 | Scaffold45:741110:741706:1 |
| GAUT035460 | *G. austeni* | Mgp9 | Scaffold45:754790:755367:-1 |
| GAUT052948 | *G. austeni* | Mgp10 | Scaffold45:750561:751146:-1 |
| GBRI000512 | *G. brevipalpis* | mgp1 | Scaffold0:865214-866251:1 |
| GBRI004611 | *G. brevipalpis* | transferrin | Scaffold119:130438-132960:-1 |
| GBRI043675 | *G. brevipalpis* | acid sphingomyelinase | Scaffold92:340653-344890:-1 |
| Annotation Pending | *G. brevipalpis* | Mgp3 | Scaffold1:4526048:4525401:-1 |
| GBRI015315 | *G. brevipalpis* | Mgp3 | Scaffold1:4519399:4520451:-1 |
| Annotation Pending | *G. brevipalpis* | Mgp4 | Scaffold1:4533884:4534511:1 |
| GBRI015319 | *G. brevipalpis* | Mgp5 | Scaffold1:4544267:4545482:-1 |
| GBRI015316 | *G. brevipalpis* | Mgp7 | Scaffold1:4552325:4553194:-1 |
| GBRI015317 | *G. brevipalpis* | Mgp8 | Scaffold1:4547741:4549433:-1 |
| GBRI015314 | *G. brevipalpis* | Mgp9 | Scaffold1:4536820:4537630:1 |
| GBRI015312 | *G. brevipalpis* | Mgp10 | Scaffold1:4541300:4541988:1 |
| GFUI006902 | *G. fuscipes* | mgp1 | Scaffold132:123438-124472:1 |
| GFUI040165 | *G. fuscipes* | transferrin | Scaffold57:1030317-1032868:1 |
| GFUI042860 | *G. fuscipes* | acid sphingomyelinase | Scaffold647:119443-123334:1 |
| Annotation Pending partial sequence | *G. fuscipes* | Mgp2 | Scaffold88:162426:162689:1 |
| GFUI050434 | *G. fuscipes* | Mgp3 | Scaffold88:155204:155839:-1 |
| GFUI050428 | *G. fuscipes* | Mgp4 | Scaffold88:166089:166691:1 |
| GFUI050421 | *G. fuscipes* | Mgp5 | Scaffold88:181207:182002:-1 |
| GFUI050422 | *G. fuscipes* | Mgp6 | Scaffold88:159615:161264:-1 |
| GFUI050451 | *G. fuscipes* | Mgp7 | Scaffold88:190717:191338:-1 |
| GFUI050436 | *G. fuscipes* | Mgp8 | Scaffold88:186333:186929:-1 |
| GFUI050420 | *G. fuscipes* | Mgp9 | Scaffold88:170223:170806:1 |
| GFUI050429 | *G. fuscipes* | Mgp10 | Scaffold88:174414:174999:1 |
| GMOY009745 | *G. morsitans* | mgp1 | scf7180000651870:191183-192614:1 |
| GMOY004228 | *G. morsitans* | transferrin | scf7180000648003-44929:47549:1 |
| GMOY002246 | *G. morsitans* | acid sphingomyelinase | scf7180000643139-13861:46922:-1 |
| GMOY001342 | *G. morsitans* | Mgp2 | scf7180000641289:61241:61943:-1 |
| GMOY012125 | *G. morsitans* | Mgp3 | scf7180000641289:68753:72286:1 |
| GMOY012368 | *G. morsitans* | Mgp4 | scf7180000641289:57776:58406:-1 |
| GMOY012370 | *G. morsitans* | Mgp5 | scf7180000641289:42695:43570:1 |
| GMOY001343 | *G. morsitans* | Mgp6 | scf7180000641289:62719:63396:1 |
| GMOY012377 | *G. morsitans* | Mgp7 | scf7180000641289:33723:34473:1 |
| GMOY012016 | *G. morsitans* | Mgp8 | scf7180000641289:38091:38719:1 |
| GMOY012371 | *G. morsitans* | Mgp9 | scf7180000641289:53684:54281:-1 |
| GMOY012369 | *G. morsitans* | Mgp10 | scf7180000641289:49291:50000:-1 |
| GPAI016437 | *G. pallidipes* | mgp1 | Scaffold1:4107747-4108724:-1 |
| GPAI033230 | *G. pallidipes* | transferrin | Scaffold43:693965-696554:-1 |
| GPAI027011 | *G. pallidipes* | acid sphingomyelinase | Scaffold326:50752-54245:-1 |
| GPAI032324 | *G. pallidipes* | Mgp2 | Scaffold41:591935:592530:1 |
| GPAI032303 | *G. pallidipes* | Mgp3 | Scaffold41:584457:585141:-1 |
| GPAI032318 | *G. pallidipes* | Mgp4 | Scaffold41:595496:596201:1 |
| GPAI032322 | *G. pallidipes* | Mgp5 | Scaffold41:609867:610986:-1 |
| GPAI032321 | *G. pallidipes* | Mgp6 | Scaffold41:590355:591123:-1 |
| GPAI032319 | *G. pallidipes* | Mgp7 | Scaffold41:617126:619713:-1 |
| GPAI032320 | *G. pallidipes* | Mgp8 | Scaffold41:614648:615378:-1 |
| GPAI032317 | *G. pallidipes* | Mgp9 | Scaffold41:599399:600206:1 |
| GPAI032316 | *G. pallidipes* | Mgp10 | Scaffold41:603788:604438:1 |
| GPPI015121 | *G. palpalis* | mgp1 | Scaffold68:323777-324973:-1 |
| GPPI043228 | *G. palpalis* | transferrin | Scaffold559:96887-99438:-1 |
| GPPI044202 | *G. palpalis* | acid sphingomyelinase | Scaffold599:29316-34911:-1 |
| GPPI010382 | *G. palpalis* | Mgp2 | Scaffold35:660762:661357:1 |
| GPPI010391 | *G. palpalis* | Mgp3 | Scaffold35:650473:658538:-1 |
| Annotation Pending partial sequence | *G. palpalis* | Mgp4 | Scaffold35:664314:664756:1 |
| GPPI052422 | *G. palpalis* | Mgp5 | Scaffold35:679454:680501:-1 |
| GPPI010392 | *G. palpalis* | Mgp6 | Scaffold35:658759:659920:-1 |
| GPPI052421 | *G. palpalis* | Mgp7 | Scaffold35:688827:689869:-1 |
| GPPI052420 | *G. palpalis* | Mgp8 | Scaffold35:684486:685455:-1 |
| GPPI010383 | *G. palpalis* | Mgp9 | Scaffold35:668551:669134:1 |
| GPPI010380 | *G. palpalis* | Mgp10 | Scaffold35:672681:673401:1 |
