## Supplemental Table 7 Glossina Male Accessory Protein Genes for "The *Glossina* Genome Cluster: Comparative Genomic Analysis of the Vectors of African Trypanosomes"

| *Glossina morsitans* Seminal Protein Genes | *Glossina pallidipes* Orthologs | *Glossina austeni* Orthologs | *Glossina fuscipes* Orthologs | *Glossina palpalis* Orthologs | *Glossina brevipalpis* Orthologs | Gene description |
| --- | --- | --- | --- | --- | --- | --- |
| GMOY000899 | GPAI014669 | GAUT028232 | GFUI042427 | GPPI014847 | GBRI008346 |  |
| GMOY000930 | GPAI013363 |  |  | GPPI005671 | GBRI027522 | serine protease inhibitor (serpin) 9 [Source:VB Community Annotation] |
| GMOY000990 | GPAI018055 | GAUT027968 | GFUI023610 | GPPI046815 | GBRI042139 | serine protease inhibitor (serpin) 5 [Source:VB Community Annotation] |
| GMOY002262 | GPAI003186 | GAUT020005 | GFUI015527 | GPPI041930 | GBRI036070 | serine protease inhibitor (serpin) 7 [Source:VB Community Annotation] |
| GMOY002279 | GPAI019961 |  |  | GPPI013616 | GBRI033168 |  |
| GMOY002399 | GPAI026065 | GAUT036613 | GFUI045750 | GPPI027571 |  |  |
| GMOY002442 | GPAI039816 | GAUT046193 | GFUI043086 | GPPI017645 | GBRI029243 | serine protease inhibitor (serpin) 1 [Source:VB Community Annotation] |
| GMOY002443 | GPAI039813 | GAUT046196 | GFUI043085 | GPPI017647 | GBRI011243 | serine protease inhibitor (serpin) 2 [Source:VB Community Annotation] |
| GMOY002444 | GPAI039809 | GAUT046192 |  | GPPI017642 | GBRI011241 | serine protease inhibitor (serpin) 3 [Source:VB Community Annotation] |
| GMOY003382 | GPAI030837 | GAUT006156 | GFUI013890 | GPPI025543 | GBRI020025 | serine protease inhibitor (serpin) 11 [Source:VB Community Annotation] |
| GMOY003656 | GPAI039671 | GAUT021607 | GFUI034371 | GPPI048779 | GBRI002517 | serine protease inhibitor (serpin) 4 [Source:VB Community Annotation] |
| GMOY003656 | GPAI039671 | GAUT021607 | GFUI034371 | GPPI048779 | GBRI002522 | serine protease inhibitor (serpin) 4 [Source:VB Community Annotation] |
| GMOY003657 | GPAI039670 | GAUT021609 | GFUI034370 |  | GBRI002518 | serine protease inhibitor (serpin) 5 [Source:VB Community Annotation] |
| GMOY004505 | GPAI006745 | GAUT028678 | GFUI033436 | GPPI037293 | GBRI045069 |  |
| GMOY004506 | GPAI006749 | GAUT028675 | GFUI033435 | GPPI021164 |  |  |
| GMOY004724 | GPAI035555 | GAUT022204 | GFUI052930 | GPPI025518 |  |  |
| GMOY004725 |  | GAUT022209 |  |  | GBRI034889 |  |
| GMOY004726 | GPAI035537 | GAUT022203 |  |  | GBRI034889 |  |
| GMOY004727 |  | GAUT022202 |  |  | GBRI034889 |  |
| GMOY004969 | GPAI009113 |  | GFUI048292 | GPPI031227 | GBRI045069 |  |
| GMOY005771 | GPAI012918 | GAUT025963 |  |  | GBRI005448 |  |
| GMOY005874 | GPAI017999 | GAUT029311 | GFUI008562 | GPPI028532 | GBRI010920 |  |
| GMOY005875 | GPAI018000 | GAUT029310 | GFUI008563 | GPPI028531 | GBRI010919 |  |
| GMOY005876 | GPAI018009 | GAUT029308 | GFUI008564 | GPPI028521 | GBRI010929 |  |
| GMOY005876 | GPAI018009 | GAUT029308 | GFUI008564 | GPPI028521 | GBRI010924 |  |
| GMOY006016 |  | GAUT021504 | GFUI051598 | GPPI013618 | GBRI033168 | serine proteinase inhibitor [Source:VB Community Annotation] |
| GMOY006927 | GPAI037620 | GAUT021884 | GFUI021198 | GPPI018038 | GBRI023575 |  |
| GMOY006928 | GPAI037622 | GAUT021882 | GFUI021195 | GPPI018036 | GBRI023572 |  |
| GMOY007314 | GPAI006440 | GAUT028974 | GFUI007906 | GPPI023932 | GBRI036202 | odorant binding protein 17 [Source:VB Community Annotation] |
| GMOY007757 | GPAI008777 | GAUT040992 | GFUI008988 | GPPI008631 | GBRI016436 | odorant binding protein 4 [Source:VB Community Annotation] |
| GMOY007759 | GPAI008749 | GAUT040977 | GFUI008965 | GPPI052283 | GBRI016473 |  |
| GMOY007760 | GPAI008750 | GAUT005680 |  |  |  |  |
| GMOY008627 | GPAI041316 | GAUT048422 | GFUI011319 | GPPI037728 |  |  |
| GMOY008628 | GPAI041313 | GAUT048423 | GFUI011316 | GPPI037724 | GBRI040455 |  |
| GMOY008942 | GPAI029944 | GAUT039250 | GFUI015916 | GPPI027927 | GBRI008095 | serine protease inhibitor (serpin) 6 [Source:VB Community Annotation] |
| GMOY009481 | GPAI041306 | GAUT044880 | GFUI011310 |  |  |  |
| GMOY009777 |  | GAUT012797 |  | GPPI028287 | GBRI032716 |  |
| GMOY009778 | GPAI004192 |  | GFUI008929 | GPPI028285 |  |  |
| GMOY010053 | GPAI020254 | GAUT051462 | GFUI012429 | GPPI016282 | GBRI025356 | serine protease inhibitor (serpin) 12 [Source:VB Community Annotation] |
| GMOY012007 | GPAI011576 | GAUT014044 | GFUI019208 | GPPI033766 | GBRI004383 | serine protease inhibitor (serpin) 10 [Source:VB Community Annotation] |
| GMOY012229 | GPAI008752 | GAUT052951 | GFUI054345 | GPPI052451 | GBRI016471 |  |
| GMOY012237 | GPAI006441 | GAUT028973 | GFUI007905 | GPPI023929 | GBRI036200 |  |
| GMOY012237 | GPAI006441 | GAUT028973 | GFUI007905 | GPPI023929 | GBRI036198 |  |
| GMOY013003 | GPAI041312 | GAUT048424 | GFUI011318 |  | GBRI040454 |  |
| GMOY013035 | GPAI018055 | GAUT027968 | GFUI023610 | GPPI046815 | GBRI042139 | Serine protease inhibitor (serpin) 5 [Source:UniProtKB/TrEMBL;Acc:A0A1B0GGD0] |
| GMOY013054 | GPAI049061 | GAUT028235 | GFUI042426 | GPPI014848 | GBRI045553 |  |
| GMOY013055 | GPAI014675 | GAUT028233 | GFUI042425 | GPPI002594 | GBRI008349 |  |
| GMOY013055 | GPAI014675 | GAUT028233 | GFUI042425 | GPPI052269 | GBRI008349 |  |
| GMOY013075 | GPAI041873 |  | GFUI038743 | GPPI041861 | GBRI005448 |  |
| GMOY013229 | GPAI001679 | GAUT015080 | GFUI038739 | GPPI041864 | GBRI005448 |  |
| GMOY013305 |  | GAUT052823 | GFUI038747 | GPPI041862 | GBRI005448 |  |
| GMOY013306 |  | GAUT052825 |  |  | GBRI005448 |  |
| GMOY013307 | GPAI041874 | GAUT052824 | GFUI038743 | GPPI041861 | GBRI005448 | Putative uncharacterized protein [Source:UniProtKB/TrEMBL;Acc:D3TQH3] |
| GMOY013308 |  |  |  |  | GBRI005448 |  |
