## Supplemental Table 8 Glossina strains and colonies of origin for "The *Glossina* Genome Cluster: Comparative Genomic Analysis of the Vectors of African Trypanosomes"

**Supplemental Table 8:** *Glossina* strains and colonies of origin for sequenced *Glossina* species

| Species | Total Sequence coverage | Strain | Sample IDs | Location |
| --- | --- | --- | --- | --- |
| *Glossina fuscipes* | 52X | IPCL* | MD7, MD7-2, MD8, MD8-1 | Institute of Zoology  Slovak Academy of Sciences |
| *Glossina austeni* | 50X | TTRI |  | Tsetse and Trypanosomosis Research Institute (TTRI) laboratories in Tanga, Tanzania |
| *Glossina pallidipes* | 46X | IPCL* | MD2 1-4 | Institute of Zoology  Slovak Academy of Sciences |
| *Glossina brevipalpis* | 45X | IPCL* |  | IPCL* laboratories in Seibersdorf, Austria |
| *Glossina palpalis* | 58X | IPCL* |  | Institute of Zoology  Slovak Academy of Sciences |

* IPCL = Insect Pest Control Laboratory of the Joint FAO/IAEA Division of Nuclear Techniques in Food and Agriculture, Seibersdorf, Austria
