## Supplemental Table 9 Glossina species contig and scaffold assembly statistics for "The *Glossina* Genome Cluster: Comparative Genomic Analysis of the Vectors of African Trypanosomes"

| Scaffold length | *Glossina morsitans* | *Glossina pallidipes* | *Glossina austeni* | *Glossina fuscipes* | *Glossina palpalis* | *Glossina brevipalpis* |
| --- | --- | --- | --- | --- | --- | --- |
| *> 1 Mb* | 13 | 102 | 78 | 70 | 63 | 81 |
| *250 kb-1 Mb* | 138 | 248 | 316 | 393 | 395 | 202 |
| *100 – 250 kb* | 605 | 184 | 248 | 330 | 326 | 136 |
| *10 -100 kb* | 3,663 | 290 | 379 | 496 | 709 | 257 |
| *5 -10 kb* | 737 | 106 | 94 | 165 | 507 | 85 |
| *2 -5 kb* | 1,933 | 255 | 206 | 252 | 978 | 156 |
| *< 2 kb* | 6,718 | 541 | 884 | 689 | 948 | 734 |
| *Total # of Contigs* | 24,071 | 7,275 | 18,748 | 13,688 | 31,320 | 16,993 |
| *N50^*^ Contig Length(kb)* | 49 | 167 | 46 | 64 | 24 | 62 |
| *Total # of Scaffolds* | 13,807 | 1,726 | 2,205 | 2,395 | 3,926 | 1,651 |
| *N50^*^ Scaffold Length(kb)* | 120 | 1,038 | 812 | 561 | 575 | 1,209 |

^*^N50 metric is that percentage of the assembled genome of the measured length metric or greater
