## Supplemental Table 10 Total measured repetitive elements among Glossina genomes for "The *Glossina* Genome Cluster: Comparative Genomic Analysis of the Vectors of African Trypanosomes"

|  | *Glossina morsitans* | *Glossina pallidipes* | *Glossina austeni* | *Glossina fuscipes* | *Glossina palpalis* | *Glossina brevipalpis* |
| --- | --- | --- | --- | --- | --- | --- |
| DNA | 4.99% | 6.49% | 6.56% | 6.01% | 5.27% | 7.28% |
| LINE | 3.97% | 5.59% | 5.23% | 5.84% | 5.47% | 2.97% |
| LTR | 0.31% | 0.52% | 0.46% | 0.83% | 0.73% | 0.26% |
| RC | 7.46% | 8.93% | 9.07% | 8.00% | 7.89% | 6.19% |
| rRNA | 0.00% | 0.00% | 0.02% | 0.03% | 0.02% | 0.00% |
| Simple repeat | 0.36% | 0.22% | 0.19% | 0.19% | 0.14% | 0.18% |
| SINE | 0.12% | 0.11% | 0.07% | 0.05% | 0.05% | 0.00% |
| Unknown | 10.18% | 5.79% | 5.43% | 5.91% | 5.16% | 3.97% |
| Satellite | 0.00% | 0.00% | 0.00% | 0.00% | 0.00% | 0.02% |
| Total | **27.38%** | **27.65%** | **27.03%** | **26.87%** | **24.72%** | **20.87%** |
