## Supplemental Table 11 Predicted gene sets for new Glossina genomes for "The *Glossina* Genome Cluster: Comparative Genomic Analysis of the Vectors of African Trypanosomes"

Gene set versions are maintained at ([www.vectorbase.org](http://www.vectorbase.org)) for each organism. All highlighted cells relate to the current gene set version indicated in the table. Statistics for older gene set versions are given along with the relevant version number.

| Species | Initial MAKER protein coding genes (version) | Filtered MAKER protein coding genes (version) | Current gene set version | Predicted RNA genes | Manual gene edits | Total protein coding genes | Total protein coding transcripts |
| --- | --- | --- | --- | --- | --- | --- | --- |
| *G. austeni* | 22,814 (GausT1.1) | 19,763 (GausT1.2) | Gaust1.3 | 483 | 125 | 19,739 | 19,752 |
| *G. brevipalpis* | 19,521 (Gbrel1.1) | 14,627 (Gbrel1.2) | Gbrel1.3 | 371 | 180 | 14,637 | 14,643 |
| *G. fuscipes* | 23,264 (Gfusl1.1) | 20,141 (Gfusl1.2) | Gfusl1.3 | 493 | 199 | 20,129 | 20,145 |
| *G. pallidipes* | 21,935 (Gpall1.1) | 19,282 (Gpall1.2) | Gpall1.3 | 436 | 189 | 19,290 | 19,303 |
| *G. palpalis* | 20,726 (Gpapl1.0) | 20,152 (Gpapl1.0) | Gpapl1.1 | 462 | 90 | 20,153 | 20,158 |
