## Supplemental Table 12 BUSCO Genomic Analysis Results for "The *Glossina* Genome Cluster: Comparative Genomic Analysis of the Vectors of African Trypanosomes"

| BUSCO Gene Analysis Results (Percentage) (diptera_odb9 geneset) | | | | | | |
| --- | --- | --- | --- | --- | --- | --- |
| Species | **Complete BUSCOs** | **Complete and single-copy BUSCOS** | **Complete and duplicated BUSCOs** | **Fragmented BUSCOs** | **Missing BUSCOs** | **Total BUSCO groups searched** |
| *G. austeni* | 97.11% | 93.00% | 4.11% | 2.18% | 0.71% | 100.00% |
| *G. morsitans* | 93.53% | 88.00% | 5.54% | 3.22% | 3.25% | 100.00% |
| *G. pallidipes* | 95.53% | 90.78% | 4.75% | 2.72% | 1.75% | 100.00% |
| *G. palpalis* | 95.00% | 87.53% | 7.47% | 3.32% | 1.68% | 100.00% |
| *G. fuscipes* | 96.50% | 91.14% | 5.36% | 2.32% | 1.18% | 100.00% |
| *G. brevipalpis* | 95.14% | 89.03% | 6.11% | 2.97% | 1.89% | 100.00% |

| BUSCO Genomic Analysis Results (Percentage) (diptera_odb9 geneset) | | | | | | |
| --- | --- | --- | --- | --- | --- | --- |
| Species | **Complete BUSCOs** | **Complete and single-copy BUSCOS** | **Complete and duplicated BUSCOs** | **Fragmented BUSCOs** | **Missing BUSCOs** | **Total BUSCO groups searched** |
| *G. austeni* | 98.07% | 97.18% | 0.89% | 1.25% | 0.68% | 100.00% |
| *G. morsitans* | 92.03% | 91.25% | 0.79% | 3.32% | 4.64% | 100.00% |
| *G. pallidipes* | 98.43% | 97.36% | 1.07% | 1.07% | 0.50% | 100.00% |
| *G. palpalis* | 97.07% | 92.85% | 4.22% | 1.86% | 1.07% | 100.00% |
| *G. fuscipes* | 98.32% | 97.21% | 1.11% | 1.18% | 0.50% | 100.00% |
| *G. brevipalpis* | 97.96% | 97.11% | 0.86% | 1.25% | 0.79% | 100.00% |
