## Supplemental Table 13 Primers used to amplify the mitochondrial genome for "The *Glossina* Genome Cluster: Comparative Genomic Analysis of the Vectors of African Trypanosomes"

**Supplemental Table 13:** Details of the primers used to amplify the mitochondrial genome of the seven-tsetse species and for HRM analysis.

| Primers to amplify tsetse species mtDNA genome | |
| --- | --- |
| Primer pair | **Primer sequence (5’🡺3’)** |
| MtDNAtseF1 | TGATAAAWTTTGTGCCAGCAATCG |
| MtDNAtseR1 | TCCAGTACAYCTACTATGTTACGACT |
| MtDNAtseF2 | ATGCACATATCGCCCGTCRCTCTT |
| MtDNAtseR2 | ATAGAAACCAACCTGGCTYACR |
| MtDNAtseF3 | AGCCAGGTTGGTTTCTATCTTT |
| MtDNAtseR3 | TTAACHTGAATTGGAGCYCGACCW |
| MtDNAtseF4 | WGGTCGRGCTCCAATTCADGTT |
| MtDNAtseR4 | TTAGTHCAATGAGTWTGAGGWGGH |
| MtDNAtseF5 | DCCWCCTCAWACTCATTGDACTA |
| MtDNAtseR5 | AATCATTHCCATGWGTDCGAATW |
| MtDNAtseF6 | WATTCGHACWCATGGDAATGAT |
| MtDNAtseR6 | CAGGRGCTTCTACATGAGCTTTAGG |
| MtDNAtseF7 | TAAAGCTCATGTAGAAGCYCCTGT |
| MtDNAtseR7 | TCAAAYTCATARTTAGCCCCTAAHCCN |
| MtDNAtseF8 | NGGDTTAGGGGCTAAYTATGARTTTGA |
| MtDNAtseR8 | CAATCTWCCTTGAATATGAAGCG |
| MtDNAtseF9 | GCAGCTTTTWCTTGRACTTCATACTT |
| MtDNAtseR9 | TCAGGDATTACTGTDACTTGAGCY |
| MtDNAtseF10 | GCTCAAGTHACAGTAATHCCTGADGT |
| MtDNAtseR10 | AGATGACTGAAAGCAAGTAYTGGTC |
| MtDNAtseF11 | GACCARTACTTGCTTTCAGTCATC |
| MtDNAtseR11 | YTAGCNGGWATACCTCGTCGY |
| MtDNAtseF12 | RCGACGAGGTATWCCNGCTAR |
| MtDNAtseR12 | TTCAGCCATTTAATCGCRACARTGA |
| MtDNAtseF13 | YGCGATTAAATGGCTGAAKW |
| MtDNAtseR13 | GCTAAHHAAGCTAMTGGGTTCATACC |
| MtDNAtseF14 | RATGGGGTATGAACCCAKTAGC |
| MtDNAtseR14 | CCTCTRAATAGACTAAAATACCGCCA |
| Primers for HRM analysis of tsetse populations/haplotypes | |
| TsetseHRM-F | TAGCCCCTAACCCCGCTATAAAT |
| TsetseHRM-R | TTCCATTTTCTTCTTGATTGCCTGC |
