## Supplemental Table 14 SRA Numbers for "The *Glossina* Genome Cluster: Comparative Genomic Analysis of the Vectors of African Trypanosomes"

**Supplemental Table 16: SRA Numbers and Associated Information on *Glossina* RNA-seq Datasets**

| **Accession** | **Sample Name** | **Link to Datafile** | **Organism** | **Tax ID** |
| --- | --- | --- | --- | --- |
| SAMN09839318 | Whole Male *G. brevipalpis* RNA-seq | <https://www.ncbi.nlm.nih.gov/sra/9839318> | *Glossina brevipalpis* | 37001 |
| SAMN09839319 | Non-lactating Whole Female *G. brevipalpis* RNA-seq | <https://www.ncbi.nlm.nih.gov/sra/9839319> | *Glossina brevipalpis* | 37001 |
| SAMN09839320 | Lactating Whole Female *G. brevipalpis* RNA-seq | <https://www.ncbi.nlm.nih.gov/sra/9839320> | *Glossina brevipalpis* | 37001 |
| SAMN09839321 | Whole Male *G. fuscipes* RNA-seq | <https://www.ncbi.nlm.nih.gov/sra/9839321> | *Glossina fuscipes* | 7396 |
| SAMN09839322 | Non-lactating Whole Female *G. fuscipes* RNA-seq | <https://www.ncbi.nlm.nih.gov/sra/9839322> | *Glossina fuscipes* | 7396 |
| SAMN09839323 | Lactating Whole Female *G. fuscipes* RNA-seq | <https://www.ncbi.nlm.nih.gov/sra/9839323> | *Glossina fuscipes* | 7396 |
| SAMN09839324 | Whole Male *G. pallidipes* RNA-seq | <https://www.ncbi.nlm.nih.gov/sra/9839324> | *Glossina pallidipes* | 7398 |
| SAMN09839325 | Non-lactating Whole Female *G. pallidipes* RNA-seq | <https://www.ncbi.nlm.nih.gov/sra/9839325> | *Glossina pallidipes* | 7398 |
| SAMN09839326 | Lactating Whole Female *G. pallidipes* RNA-seq | <https://www.ncbi.nlm.nih.gov/sra/9839326> | *Glossina pallidipes* | 7398 |
| SAMN09839327 | Whole Male *G. palpalis gambiensis* RNA-seq | <https://www.ncbi.nlm.nih.gov/sra/9839327> | *Glossina palpalis gambiensis* | 67801 |
| SAMN09839328 | Non-lactating Whole Female *G. palpalis* *gambiensis* RNA-seq | <https://www.ncbi.nlm.nih.gov/sra/9839328> | *Glossina palpalis gambiensis* | 67801 |
| SAMN09839329 | Lactating Whole Female *G. palpalis gambiensis* RNA-seq | <https://www.ncbi.nlm.nih.gov/sra/9839329> | *Glossina palpalis gambiensis* | 67801 |
| SRS2364381 | Male Reproductive tract *G. morsitans* RNA-seq | <https://www.ncbi.nlm.nih.gov/sra/SRS2364381> | *Glossina morsitans morsitans* | 37546 |
| SRS430099 | Non-lactating Whole Female *G. morsitans* RNA-seq | <https://www.ncbi.nlm.nih.gov/sra/SRS430099> | *Glossina morsitans morsitans* | 37546 |
| SRS430097 | Lactating Whole Female *G. morsitans* RNA-seq | <https://www.ncbi.nlm.nih.gov/sra/SRS430097> | *Glossina morsitans morsitans* | 37546 |
| SRS686445 | Whole Male *G. austeni* RNA-seq | <https://www.ncbi.nlm.nih.gov/sra/SRX682955> | *Glossina austeni* | 7395 |
| SRS686473 | Non-lactating Whole Female *G. austeni* RNA-seq | <https://www.ncbi.nlm.nih.gov/sra/SRX682983> | *Glossina austeni* | 7395 |
