## Supplemental Table 16 Neuropeptide genes for "The *Glossina* Genome Cluster: Comparative Genomic Analysis of the Vectors of African Trypanosomes"

**Supplemental Table 16:** Neuropeptide genes identified in various tsetse flies, including *Glossina morsitans*, as deduced by homology searches of the respective genomes *in silico*. The gene identification number is supplied, or listed as “not located” when not found. Where more than one gene is identified in one species and not in another, the latter is indicated as “no” for no other gene identified. Rows colored in brown are lacking representation across all *Glossina* species while yellow rows represent partial representation

| **Neuropeptide^1^** | ***G.morsitans*** | ***G. pallidipes*** | ***G. austeni*** | ***G. palpalis*** | ***G. fuscipes*** | ***G. brevipalpis*** |
| --- | --- | --- | --- | --- | --- | --- |
| ACP | not located | not located | not located | not located | not located | not located |
| Adipokinetic hormone | GMOY003470 | GPAI036121 | GAUT013261 | GPPI030617 | GFUI054167 | GBRI027509 |
|  | GMOY003469 | GPAI049064 | GAUT013267 | GPPI030614 | GFUI054166 | GBRI045557 |
| Allatotropin | not located | not located | not located | not located | not located | not located |
| Allatostatin A | GMOY002621 | GPAI037922 | GAUT042383 | GPPI044384 | GFUI015852 | GBRI014005 |
| Allatostatin B | GMOY012235 | GPAI046483 | GAUT051974 | GPPI036833 | GFUI028593 | GBRI016323 no confirmation now. |
| Allatostatin C | GMOY012106 | GPAI030796 | GAUT018803 | GPPI008165 | GFUI031621 | GBRI013358 |
|  | no | no | GAUT018801 | no | no | no |
| Bursicon-α | GMOY011521 | GPAI045899 | GAUT041352 | GPPI024687 | GFUI031519 | GBRI004157 |
| Bursicon-β | GMOY004325 | GPAI005625 | GAUT008609 | GPPI032353 | GFUI051829 | GBRI001434 |
|  | no | no | no | GPPI047490 | GFUI051824 | GBRI001449 |
| Capa/Pyrokinin | GMOY003803 | GPAI048036 | GAUT035627 | GPPI042459 | GFUI023068 | GBRI037790 |
| CCAP | GMOY009420 | GPAI014903 | GAUT031720 | GPPI035716 | GFUI009406 | GBRI021868 |
| CCHamide-1 | GMOY003581 | GPAI017333 | GAUT048738 | GPPI050261 | GFUI047419 | not located |
|  | no | no | GAUT000929 | no | no | no |
| CCHamide-2 | GMOY010764 | GPAI006272 | GAUT047058 | GPPI027237 | GFUI034473 | GBRI028417 |
| Corazonin | GMOY012060 | GPAI024124 | GAUT003311 | GPPI019152 | GFUI053499 | GBRI045556 |
| DH (calcitonin) | GMOY003573 | GPAI011088 | GAUT009941 | GPPI036049 | GFUI008406 | GBRI018412: not verified |
| DH (CRF) | GMOY012107 | GPAI020401 | GAUT031801 | GPPI047047 | GFUI034251 | GBRI021944 |
| EH | GMOY012086 | not located | not located | GPPI046027 | not located | GBRI026411 |
| ETH | GMOY012358 | GPAI036755 | GAUT019585 | GPPI024407 | GFUI016014 | GBRI013026: not confirmed |
| FMRFamide | GMOY003883 | GPAI028872 | GAUT023561 | GPPI011945 | GFUI022977 | GBRI025526 |
| Glycoprotein α-2 | GMOY006315 | GPAI012868 | GAUT035273 | GPPI035304 | GFUI018308 | GBRI007065 |
| Glycoprotein β-5 | GMOY006313 | GPAI012869 | GAUT035261 | GPPI035289 | GFUI018328 | GBRI007062 |
| Hugin | GMOY003497 | GPAI029434 | GAUT035879 | GPPI016885 | GFUI053332 | GBRI037427 |
| ILP | GMOY003945 | GPAI002313 | GAUT046586 | GPPI022721 | GFUI049575 | GBRI018552 |
|  | GMOY003946 | GPAI001883 | GAUT006766 | GPPI015490 | GFUI041241 | GBRI033759 |
|  | GMOY005180 | no | no | no | no | GBRI004816 |
|  | GMOY002258 | no | no | no | no | no |
|  | GMOY006879 | no | no | no | no | no |
| Inotocin | not located | not located | not located | not located | not located | not located |
| ITP | GMOY012155 | GPAI043749 | GAUT047668 | GPPI029894 | GFUI042528 | GBRI044413 |
| Leucokinin | GMOY012112 | GPAI044412 | GAUT037790 | GPPI015558 | GFUI005571 | GBRI009975: not confirmed |
| Myosuppressin | GMOY011892 | GPAI035646 | GAUT034236 | GPPI027780 | GFUI041280 | GBRI017368 |
| Natalisin | GMOY006483 | GPAI024429 | GAUT036435 | GPPI022174 | GFUI014204 | GBRI016581 |
| Neuroparsin/crimpy | GMOY011055 | GPAI027170 | GAUT026343 | GPPI005114 | GFUI045135 | GBRI013110 |
| NPF | GMOY012182 | not located | GAUT035914 | GPPI016939 | GFUI053299 | not located |
| NPLP-1 | GMOY005894 | GPAI004353 | GAUT049217 | GPPI029397 | GFUI033879 | GBRI033649 |
| Orcokinin | GMOY009230 | GPAI025930 | GAUT027488 | GPPI023490 | GFUI026883 | GBRI002434 |
| PDF | GMOY000603 | GPAI046761 | GAUT031282 | GPPI022815 | GFUI049486 | not located |
| Proctolin | GMOY012104 | GPAI007711 | GAUT023108 | GPPI011465 | GFUI031914 | GBRI000660 |
| PTTH | GMOY011991 | GPAI007078 | GAUT033621 | GPPI021954 | GFUI025672 | GBRI015505 |
| RYamide | not located | not located | GAUT023352 | GPPI019391 | GFUI018333 | GBRI044269 |
| Short NPF | GMOY012142 | GPAI020549 | GAUT011332 | GPPI043755 | GFUI032056 | GBRI022854 |
| SIFamide | GMOY005296 | GPAI045409 | GAUT032331 | GPPI020456 | GFUI039970 | GBRI045710 |
|  | no | GPAI033266 | no | GPPI014368 | GFUI040137 | GBRI004649 |
| Sulfakinin | not located | not located | not located | not located | not located | not located |
| Tachykinin | GMOY008567 | GPAI020386 | GAUT031793 | GPPI048084 | GFUI034255 | GBRI021939 |
| Trissin | GMOY009417 | GPAI014911 | GAUT031727 | GPPI035715 | GFUI009405 | GBRI021869 |

^1^Abbreviated peptide names: ACP, adipokinetic hormone/corazonin-like neuropeptide; CCAP, crustacean cardioactive peptide; DH (calcitonin), calcitonin-like diuretic hormone; DH (CRF), corticotropin releasing factor-like diuretic hormone; EH, eclosion hormone; ETH, ecdysis triggering hormone; ILP, insulin-like peptide; ITP, ion-transporting peptide; NPF, neuropeptide F; NPLP, neuropeptide-like precursor; PDF, pigment-dispersing factor; PTTH, prothoracicotropic hormone.

CNMamide [Source:Projected from *Drosophila melanogaster* (FBgn0035282) FlyBase gene name;Acc:FBgn0035282]; *Musca domestica* = MDOA011180 & *Stomoxys calcitrans* = SCAU014405. Not in *Glossina spp* nor in mosquitoes.
