## Supplemental Table 17 Protein receptor genes for bioactive neuropeptides for "The *Glossina* Genome Cluster: Comparative Genomic Analysis of the Vectors of African Trypanosomes"

**Supplemental Table 17:** Protein receptor genes for bioactive neuropeptides identified in various tsetse flies, including *Glossina morsitans*, as well as the common housefly *Musca domestica*, as deduced by homology searches of the respective genomes *in silico*. The gene identification number is supplied, or listed as “not located” when not found. Where more than one gene is identified in one species and not in another, the latter is indicated as “no” for no other gene identified.

| **Protein Receptor for^1^** | ***G. morsitans*** | ***G. pallidipes*** | ***G. austeni*** | ***G. palpalis*** | ***G. fuscipes*** | ***G. brevipalpis*** | ***M. domestica*** |
| --- | --- | --- | --- | --- | --- | --- | --- |
| ACP | not located | not located | not located |  | not located | not located | not located |
| Adipokinetic hormone | GMOY008368 | GPAI026595 | GAUT019061 | GPPI021443 | GFUI027776 | GBRI000966 | MDOA004853 |
| Allatotropin | not located | not located | not located | not located | not located | not located | not located |
| Allatostatin A | GMOY012001 | GPAI029482 | GAUT035837 | GPPI017451 | GFUI053372 | GBRI037476 | MDOA015456 |
|  | GMOY005179 | GPAI010186 | GAUT000491 | GPPI037221 | GFUI036826 | GBRI033753 | MDOA007526 |
| Allatostatin B | not located | not located | not located | not located | not located | not located | not located |
| Allatostatin C | GMOY002651 | GPAI046718 | GAUT041867 | GPPI012370 | GFUI036377 | GBRI004988 | MDOA007493 |
|  | GMOY012068 | GPAI019597 | no | GPPI032911 | GFUI036404 | GBRI004982 | MDOA005747 |
| Bursicon-α | GMOY011521 | GPAI045899 | not located | GPPI024687 | GFUI031519 | GBRI004157 | MDOA000854 |
|  | GMOY003161 | GPAI014560 | GAUT050652 | GPPI014972 | GFUI006325 | GBRI032052 | MDOA005834 |
| Bursicon-β | GMOY004325 | GPAI005625 | not located | GPPI032353 | GFUI051829 | not located | MDOA014861 |
|  | no | no | no | GPPI047490 | no | no | no |
| Capa /Cap2B/ Pyrokinin/Hugin | GMOY012161 | GPAI046497 | GAUT051954 | GPPI036820 | GFUI028612 | not located | MDOA015286 |
|  | GMOY012119 | GPAI017332 | GAUT018604 | GPPI050255 | GFUI047418 | GBRI031007 | MDOA014609 |
|  | GMOY009301 | GPAI045130 | GAUT048737 | no | GFUI035668 | GBRI032091 | MDOA010837 |
|  | no | no | GAUT051726 | no | no | no | no |
| CCAP | GMOY009062 | GPAI039220 | GAUT010112 | GPPI038279 | GFUI046430 | GBRI009844 | MDOA013475 |
| CCHamide-1 | GMOY006903 | GPAI028941 | GAUT030352 | GPPI040785 | GFUI016889 | GBRI002327 | MDOA013652 |
| CCHamide-2 | GMOY009111 | GPAI044844 | GAUT023072 | GPPI016208 | GFUI015341 | GBRI028932 | MDOA013613 |
| CNMamide | GMOY005917 | GPAI017658 | GAUT003612 | GPPI043177 | GFUI025643 | GBRI031527 | MDOA000808 |
|  | no | no | no | no | GFUI027056 | no | MDOA016759 |
| Corazonin | GMOY006527 | GPAI013494 | GAUT015004 | not located | GFUI040669 | GBRI016365 | MDOA014451 |
|  | no | no | no | no | no | no | MDOA008078 |
| DH 31 (calcitonin) | GMOY010919 | GPAI028216 | GAUT023385 | GPPI048644 | GFUI017241 | GBRI044319 | MDOA000230 |
| DH 44 (CRF) | GMOY012160 | GPAI034419 | GAUT043673 | GPPI028529 | GFUI008579 | GBRI044266 | MDOA009175 |
|  | GMOY012023 | GPAI017992 | GAUT029316 | GPPI028527 | GFUI008577 | GBRI010922 | MDOA006927 |
|  | no | GPAI027700 | GAUT023363 | GPPI043324 | GFUI017282 | no | MDOA006619 |
| ETH | GMOY012065 | GPAI007847 | GAUT007145 | GPPI045513 | GFUI000307 | GBRI036878 | MDOA002914 |
| FMRFamide | GMOY012135 | not located | not located | GPPI020871 | GFUI044848 | not located | MDOA002590 |
| Glycoprotein 2α / 5β | GMOY008940 | GPAI038760 | GAUT035203 | GPPI015990 | GFUI026094 | GBRI013835 | MDOA014289 |
|  | GMOY006337 | no | no | no | no | no | no |
| Hector | GMOY009447 | GPAI031364 | GAUT050139 | GPPI012503 | GFUI051144 | not located | MDOA006082 |
|  | no | no | GAUT050138 | no | no | no | no |
| Leucokinin | GMOY007765 | GPAI032398 | GAUT035408 | GPPI047278 | not located | GBRI042364 | MDOA009703 |
| Lgr3/Orphan GPCR 6 | GMOY012010 | GPAI005664 | GAUT009324 | GPPI021840 | GFUI017878 | GBRI041157 | MDOA002739 |
| Moody | GMOY003467 | GPAI021938 | GAUT014222 | GPPI025997 | GFUI045655 | GBRI045038 | MDOA001450 |
| Myosuppressin | GMOY009670 | GPAI005116 | GAUT047489 | GPPI008887 | GFUI034109 | GBRI039573 | MDOA003714 |
|  | no | no | no | no | no | no | MDOA012230 |
| NPF | GMOY006767 | GPAI023998 | GAUT038865 | GPPI019011 | GFUI050788 | GBRI007095 | MDOA001172 |
| Orphan GPCR 2 | GMOY001972 | not located | GAUT015610 | not located | GFUI047692 | GBRI034114 | MDOA006597 |
| Orphan GPCR 3 | GMOY005600 | GPAI021925 | GAUT014227 | GPPI025988 | GFUI045672 | GBRI045041 | MDOA002331 |
| Orphan GPCR 7 | GMOY012103 | GPAI034951 | GAUT027762 | GPPI009196 | GFUI024162 | GBRI044968 | MDOA005523 |
| Orphan GPCR 8 | GMOY011869 | GPAI020655 | GAUT008659 | GPPI034431 | GFUI004073 | GBRI012070 | MDOA009098 |
| Orphan GPCR 9 | GMOY012305 | GPAI024566 | GAUT033522 | GPPI044476 | GFUI027275 | GBRI012466 | MDOA007874 |
| Orphan GPCR 10 | GMOY012043 | GPAI001438 | GAUT021415 | GPPI029024 | GFUI034321 | GBRI003919 | MDOA005504 |
| PDF | GMOY007422 | GPAI031267 | GAUT050246 | GPPI002487 | GFUI003580 | GBRI010392 | MDOA001908 |
| Proctolin | GMOY011680 | GPAI048197 | GAUT015414 | GPPI001122 | GFUI001648 | GBRI003761 | MDOA010087 |
| RYamide/mammalian NPY-like | GMOY011997 | GPAI021450 | GAUT014810 | GPPI032837 | GFUI043225 | GBRI000226 | MDOA001259 |
|  | GMOY012052 | GPAI021430 | GAUT014822 | GPPI017273 | GFUI043227 | GBRI000235 | MDOA012402 |
| MIP Receptor | GMOY004860 | GPAI042600 | GAUT040731 | GPPI002955 | GFUI026358 | GBRI023627 | MDOA006144 |
|  | no | no | no | no | GFUI026351 | no | no |
| Short NPF | GMOY006636 | not located | GAUT037859 | GPPI019355 | GFUI005605 | GBRI010003 | MDOA011011 |
| SIFamide | GMOY008798 | GPAI007832 | GAUT007169 | GPPI044149 | GFUI047709 | GBRI036855 | MDOA007765 |
| Sulfakinin | not located | not located | not located | not located | not located | GBRI027745 | MDOA011233 |
|  | no | no | no | no | no | GBRI027746 | MDOA008617 |
| Tachykinin (Putative Natalisin Receptor?) | GMOY011823 | GPAI019001 | GAUT046129 | GPPI009923 | GFUI038255 | GBRI012822 | MDOA014354 |
|  | GMOY004521 | no | no | no | no | no | no |
| Trissin | GMOY010450 | GPAI037343 | GAUT022572 | GPPI034168 | GFUI034351 | GBRI035755 | MDOA015069 |

^1^Abbreviated peptide names: ACP, adipokinetic hormone/corazonin-like neuropeptide; CCAP, crustacean cardioactive peptide; DH (calcitonin), calcitonin-like diuretic hormone; DH (CRF), corticotropin releasing factor-like diuretic hormone; EH, eclosion hormone; ETH, ecdysis triggering hormone; ILP, insulin-like peptide; ITP, ion-transporting peptide; NPF, neuropeptide F; NPLP, neuropeptide-like precursor; PDF, pigment-dispersing factor; PTTH, prothoracicotropic hormone. NPY neuropeptide Y-like

CNMamide Receptor *D. melanogaster* annotation symbol CG33696, Flybase ID	FBgn0053696, **CNMamide** (**CNMa**) is a cyclic [neuropeptide](https://en.wikipedia.org/wiki/Neuropeptide) identified by [computational analysis](https://en.wikipedia.org/w/index.php?title=Computational_analysis&action=edit&redlink=1) of [Drosophila melanogaster](https://en.wikipedia.org/wiki/Drosophila_melanogaster) protein sequences and named after its C-terminal ending motif. A gene encoding CNMa was found in most [arthropods](https://en.wikipedia.org/wiki/Arthropod) and comparison among the precursor sequences of several representative species revealed high conservation, particularly in the region of the predicted mature peptide.[^[1]^](https://en.wikipedia.org/wiki/CNMa#cite_note-cnma-1) Two conserved [cysteine](https://en.wikipedia.org/wiki/Cysteine) residues enveloping four amino acids form a [disulfide bond](https://en.wikipedia.org/wiki/Disulfide_bond) and were shown to be important for binding of the peptide to its receptor. Expression of CNMa was confirmed in the larval and adult brain of D. melanogaster but the function of the peptide has not been elucidated yet.

Hector is a G-protein coupled receptor involved in male courtship behavior. [Date last reviewed: 2017-06-15] *Drosophila melanogaster*: CG4395; FBgn0030437.

Lgr3 = Leucine-rich repeat-containing G protein-coupled receptor 3; FBgn0039354

*Drosophila melanogaster*: moody (FBgn0025631), annotation symbol CG4322. For ventral nerve cord development, Isoform A and isoform B are required in glia to regulate the acute sensitivity to cocaine and to continuously maintain the proper blood-brain barrier (BBB) function. A moody-mediated signaling pathway functions in glia to regulate nervous system insulation and drug-related behaviors. Galphai and Galphao, and the regulator of G protein signaling, loco, are required in the surface glia to achieve effective.
