## Supplemental Table 18 Family specific counts of putative cuticle protein genes for "The *Glossina* Genome Cluster: Comparative Genomic Analysis of the Vectors of African Trypanosomes"

**Supplemental Table 18. Family specific counts of putative cuticle protein genes across *Glossina* genomes.**

|  | CPR^a^ | | |  |  |  |  |  |  |  |  |
| --- | --- | --- | --- | --- | --- | --- | --- | --- | --- | --- | --- |
|  | **RR-1** | **RR-2** | **RR-Uncl** | **CPAP1** | **CPAP3** | **CPF** | **CPCFC** | **CPLCA** | **CPLCG** | **TWDL** | **Total** |
| *G. austeni* | 23 | 23 | 14 | 12 | 4 | 1 | 1 | 6 | 5 | 12 | 101 |
| *G. brevipalpis* | 33 | 25 | 16 | 10 | 5 | 1 | 1 | 7 | 5 | 10 | 113 |
| *G. fuscipes* | 37 | 28 | 20 | 13 | 5 | 1 | 1 | 7 | 5 | 13 | 130 |
| *G. pallidipes* | 34 | 27 | 19 | 11 | 6 | 1 | 1 | 9 | 4 | 10 | 122 |
| *G. palpalis* | 36 | 26 | 18 | 10 | 4 | 1 | 2 | 8 | 5 | 8 | 118 |

^a^Sequences that scored above the assigned cutoffs for the RR-1 and RR-2 models were classified as the corresponding type, whereas sequences with scores below the assigned cutoffs but above 0 were characterized as “unclassified” (Uncl). For more information, see (121).
