## Supplemental Table 19 List of all peritrophin and peritrophin-like proteins for "The *Glossina* Genome Cluster: Comparative Genomic Analysis of the Vectors of African Trypanosomes"

**Supplemental Table 19.** List of all peritrophin and peritrophin-like proteins (PLPs), containing one or more chitin binding domains (CBDs) across the six major *Glossina* species, and their orthologues in *Drosophila*.

Suggested gene names are those shown under the heading ‘symbol and synonym’.

*Drosophila* genes highlighted in yellow are those that have been shown to be constitutes of the peritrophic matrix (GO:0016490) either through experimental evidence or by electronic inference.

| FlyBase ID | VectorBase ID | Species | Symbol | Synonym | Description |
| --- | --- | --- | --- | --- | --- |
| FBgn0052036 | GMOY000880 | Morsitans | GmmPLP1 | Peritrophin-Like Protein | Chitin-binding, Peritrophin-A-Domain |
|  | GAUT003677 | Austeni | GauPLP1 | Peritrophin-Like Protein | Chitin-binding, Peritrophin-A-Domain |
|  | GBRI031179 | Brevipalpis | GbrPLP1 | Peritrophin-Like Protein | Chitin-binding, Peritrophin-A-Domain |
|  | GFUI027404 | Fuscipes | GfuPLP1 | Peritrophin-Like Protein | Chitin-binding, Peritrophin-A-Domain |
|  | GPAI017590 | Palldipes | GpaPLP1 | Peritrophin-Like Protein | Chitin-binding, Peritrophin-A-Domain |
|  | GPPI025392 | Palpalis | GppPLP1 | Peritrophin-Like Protein | Chitin-binding, Peritrophin-A-Domain |
| FBgn0052024 | GMOY000968 | Morsitans | GmmPLP2 | Peritrophin-Like Protein | Chitin-binding, Peritrophin-A-Domain |
|  | GAUT010978 | Austeni | GauPLP2 | Peritrophin-Like Protein | Chitin-binding, Peritrophin-A-Domain |
|  | GBRI005049 | Brevipalpis | GbrPLP2 | Peritrophin-Like Protein | Chitin-binding, Peritrophin-A-Domain |
|  | GFUI013195 | Fuscipes | GfuPLP2 | Peritrophin-Like Protein | Chitin-binding, Peritrophin-A-Domain |
|  | GPAI009989 | Palldipes | GpaPLP2 | Peritrophin-Like Protein | Chitin-binding, Peritrophin-A-Domain |
|  | GPPI003158 | Palpalis | GppPLP1 | Peritrophin-Like Protein | Chitin-binding, Peritrophin-A-Domain |
| FBgn0030999 | GMOY001312 | Morsitans | GmmPLP3 | Peritrophin-Like Protein | Chitin-binding, Peritrophin-A-Domain |
|  | GAUT020830 | Austeni | GauPLP3 | Peritrophin-Like Protein | Chitin-binding, Peritrophin-A-Domain |
|  | GBRI007737 | Brevipalpis | GbrPLP3 | Peritrophin-Like Protein | Chitin-binding, Peritrophin-A-Domain |
|  | GFUI048226 | Fuscipes | GfuPLP3 | Peritrophin-Like Protein | Chitin-binding, Peritrophin-A-Domain |
|  | GPAI009178 | Palldipes | GpaPLP3 | Peritrophin-Like Protein | Chitin-binding, Peritrophin-A-Domain |
| FBgn0035430 | GMOY001522 | Morsitans | GmmPLP4 | Peritrophin-Like Protein | Chitin-binding, Peritrophin-A-Domain |
|  | GAUT026255 | Austeni | GauPLP4 | Peritrophin-Like Protein | Chitin-binding, Peritrophin-A-Domain |
|  | GBRI023956 | Brevipalpis | GbrPLP4 | Peritrophin-Like Protein | Chitin-binding, Peritrophin-A-Domain |
|  | GFUI039257 | Fuscipes | GfuPLP4 | Peritrophin-Like Protein | Chitin-binding, Peritrophin-A-Domain |
|  | GPAI027229 | Palldipes | GpaPLP4 | Peritrophin-Like Protein | Chitin-binding, Peritrophin-A-Domain |
|  | GPPI005038 | Palpalis | GppPLP4 | Peritrophin-Like Protein | Chitin-binding, Peritrophin-A-Domain |
|  | GMOY001662 | Morsitans | GmmPLP5a | Peritrophin-Like Protein | Chitin-binding, Peritrophin-A-Domain |
|  | GMOY002339 | Morsitans | GmmPLP5b | Peritrophin-Like Protein | Chitin-binding, Peritrophin-A-Domain |
|  | GAUT014249 | Austeni | GauPLP5 | Peritrophin-Like Protein | Chitin-binding, Peritrophin-A-Domain |
|  | GBRI000180 | Brevipalpis | GbrPLP5 | Peritrophin-Like Protein | Chitin-binding, Peritrophin-A-Domain |
|  | GFUI007094 | Fuscipes | GfuPLP5a | Peritrophin-Like Protein | Chitin-binding, Peritrophin-A-Domain |
|  | GFUI016406 | Fuscipes | GfuPLP5b | Peritrophin-Like Protein | Chitin-binding, Peritrophin-A-Domain |
|  | GPAI001697 | Palldipes | GpaPLP5a | Peritrophin-Like Protein | Chitin-binding, Peritrophin-A-Domain |
|  | GPAI021942 | Palldipes | GpaPLP5b | Peritrophin-Like Protein | Chitin-binding, Peritrophin-A-Domain |
| FBgn0038632 | GMOY002084 | Morsitans | GmmPLP6 | Peritrophin-Like Protein | Chitin-binding, Peritrophin-A-Domain |
|  | GAUT027066 | Austeni | GauPLP6 | Peritrophin-Like Protein | Chitin-binding, Peritrophin-A-Domain |
|  | GBRI004278 | Brevipalpis | GbrPLP6 | Peritrophin-Like Protein | Chitin-binding, Peritrophin-A-Domain |
|  | GFUI036017 | Fuscipes | GfuPLP6 | Peritrophin-Like Protein | Chitin-binding, Peritrophin-A-Domain |
|  | GPAI012018 | Palldipes | GpaPLP6 | Peritrophin-Like Protein | Chitin-binding, Peritrophin-A-Domain |
| FBgn0038629 | GMOY002086 | Morsitans | GmmPLP7 | Peritrophin-Like Protein | Chitin-binding, Peritrophin-A-Domain |
|  | GAUT027081 | Austeni | GauPLP7 | Peritrophin-Like Protein | Chitin-binding, Peritrophin-A-Domain |
|  | GBRI004277 | Brevipalpis | GbrPLP7 | Peritrophin-Like Protein | Chitin-binding, Peritrophin-A-Domain |
|  | GPPI029843 | Palpalis | GppPLP7 | Peritrophin-Like Protein | Chitin-binding, Peritrophin-A-Domain |
| FBgn0037488 | GMOY002141 | Morsitans | GmmPLP8 | Peritrophin-Like Protein | Chitin-binding, Peritrophin-A-Domain |
|  | GAUT044450 | Austeni | GauPLP8 | Peritrophin-Like Protein | Chitin-binding, Peritrophin-A-Domain |
|  | GBRI023660 | Brevipalpis | GbrPLP8 | Peritrophin-Like Protein | Chitin-binding, Peritrophin-A-Domain |
|  | GFUI027472 | Fuscipes | GfuPLP8a | Peritrophin-Like Protein | Chitin-binding, Peritrophin-A-Domain |
|  | GFUI048634 | Fuscipes | GfuPLP8b | Peritrophin-Like Protein | Chitin-binding, Peritrophin-A-Domain |
|  | GPAI005401 | Palldipes | GpaPLP8 | Peritrophin-Like Protein | Chitin-binding, Peritrophin-A-Domain |
|  | GPPI048548 | Palpalis | GppPLP8 | Peritrophin-Like Protein | Chitin-binding, Peritrophin-A-Domain |
| FBgn0037487 | GMOY002142 | Morsitans | GmmPLP9 | Peritrophin-Like Protein | Chitin-binding, Peritrophin-A-Domain |
| FBgn0263748 | GMOY002708 | Morsitans | GmmPLP10  GmmPer66 | Peritrophin-Like Protein/GmmPer66 | Chitin-binding, Peritrophin-A-Domain, mucin domains |
| FBgn0085353 | GAUT001531 | Austeni | GauPLP10a | Peritrophin-Like Protein | Chitin-binding, Peritrophin-A-Domain, mucin domains |
|  | GAUT001532 | Austeni | GauPLP10b | Peritrophin-Like Protein | Chitin-binding, Peritrophin-A-Domain, mucin domains |
|  | GBRI011740 | Brevipalpis | GbrPLP10a | Peritrophin-Like Protein | Chitin-binding, Peritrophin-A-Domain, mucin domains |
|  | GBRI011741 | Brevipalpis | GbrPLP10b | Peritrophin-Like Protein | Chitin-binding, Peritrophin-A-Domain, mucin domains |
|  | GFUI006263 | Fuscipes | GfuPLP10a | Peritrophin-Like Protein | Chitin-binding, Peritrophin-A-Domain, mucin domains |
|  | GFUI006267 | Fuscipes | GfuPLP10b | Peritrophin-Like Protein | Chitin-binding, Peritrophin-A-Domain, mucin domains |
|  | GPAI021775 | Palldipes | GpaPLP10a | Peritrophin-Like Protein | Chitin-binding, Peritrophin-A-Domain, mucin domains |
|  | GPAI021777 | Palldipes | GpaPLP10b | Peritrophin-Like Protein | Chitin-binding, Peritrophin-A-Domain, mucin domains |
|  | GPPI024565 | Palpalis | GppPLP10 | Peritrophin-Like Protein | Chitin-binding, Peritrophin-A-Domain, mucin domains |
| FBgn0260386 | GMOY002812 | Morsitans | GmmPLP11 | Peritrophin-Like Protein | Chitin-binding, Peritrophin-A-Domain |
|  | GAUT028770 | Austeni | GauPLP11 | Peritrophin-Like Protein | Chitin-binding, Peritrophin-A-Domain |
|  | GBRI019411 | Brevipalpis | GbrPLP11 | Peritrophin-Like Protein | Chitin-binding, Peritrophin-A-Domain |
|  | GFUI040609 | Fuscipes | GfuPLP11 | Peritrophin-Like Protein | Chitin-binding, Peritrophin-A-Domain |
|  | GPAI030232 | Palldipes | GpaPLP11 | Peritrophin-Like Protein | Chitin-binding, Peritrophin-A-Domain |
|  | GPPI040873 | Palpalis | GppPLP11 | Peritrophin-Like Protein | Chitin-binding, Peritrophin-A-Domain |
| FBgn0031737 | GMOY003840 | Morsitans | GmmPLP12 | Peritrophin-Like Protein | Chitin-binding, Peritrophin-A-Domain |
|  | GAUT040598 | Austeni | GauPLP12 | Peritrophin-Like Protein | Chitin-binding, Peritrophin-A-Domain |
|  | GBRI039371 | Brevipalpis | GbrPLP12 | Peritrophin-Like Protein | Chitin-binding, Peritrophin-A-Domain |
|  | GFUI049678 | Fuscipes | GfuPLP12 | Peritrophin-Like Protein | Chitin-binding, Peritrophin-A-Domain |
|  | GPAI033642 | Palldipes | GpaPLP12 | Peritrophin-Like Protein | Chitin-binding, Peritrophin-A-Domain |
|  | GPPI003309 | Palpalis | GppPLP12 | Peritrophin-Like Protein | Chitin-binding, Peritrophin-A-Domain |
|  | GMOY004823 | Morsitans | GmmPLP13 | Peritrophin-Like Protein | Chitin-binding, Peritrophin-A-Domain |
|  | GAUT015903 | Austeni | GauPLP13 | Peritrophin-Like Protein | Chitin-binding, Peritrophin-A-Domain |
|  | GFUI012030 | Fuscipes | GfuPLP13a | Peritrophin-Like Protein | Chitin-binding, Peritrophin-A-Domain |
|  | GFUI017945 | Fuscipes | GfuPLP13b | Peritrophin-Like Protein | Chitin-binding, Peritrophin-A-Domain |
|  | GPAI009564 | Palldipes | GpaPLP13 | Peritrophin-Like Protein | Chitin-binding, Peritrophin-A-Domain |
| FBgn0027600 | GMOY004893 | Morsitans | GmmPLP14 | Peritrophin-Like Protein | Chitin-binding, Peritrophin-A-Domain |
|  | GAUT011300 | Austeni | GauPLP14 | Peritrophin-Like Protein | Chitin-binding, Peritrophin-A-Domain |
|  | GBRI022913 | Brevipalpis | GbrPLP14 | Peritrophin-Like Protein | Chitin-binding, Peritrophin-A-Domain |
|  | GFUI015986 | Fuscipes | GfuPLP14 | Peritrophin-Like Protein | Chitin-binding, Peritrophin-A-Domain |
|  | GPAI048004 | Palldipes | GpaPLP14 | Peritrophin-Like Protein | Chitin-binding, Peritrophin-A-Domain |
|  | GPPI007754 | Palpalis | GppPLP14 | Peritrophin-Like Protein | Chitin-binding, Peritrophin-A-Domain |
| FBgn0025390 | GMOY005251 | Morsitans | GmmPLP15 | Peritrophin-Like Protein | Chitin-binding, Peritrophin-A-Domain |
|  | GBRIO26889 | Brevipalpis | GbrPLP15 | Peritrophin-Like Protein | Chitin-binding, Peritrophin-A-Domain |
|  | GFUI045474 | Fuscipes | GfuPLP15a | Peritrophin-Like Protein | Chitin-binding, Peritrophin-A-Domain |
|  | GFUI050311 | Fuscipes | GfuPLP15b | Peritrophin-Like Protein | Chitin-binding, Peritrophin-A-Domain |
|  | GPAI039542 | Palldipes | GpaPLP15 | Peritrophin-Like Protein | Chitin-binding, Peritrophin-A-Domain |
|  | GPPI018985 | Palpalis | GppPLP15 | Peritrophin-Like Protein | Chitin-binding, Peritrophin-A-Domain |
| FBgn0038422 | GMOY005235 | Morsitans | GmmPLP16 | Peritrophin-Like Protein | Chitin-binding, Peritrophin-A-Domain |
|  | GAUT010029 | Austeni | GauPLP16 | Peritrophin-Like Protein | Chitin-binding, Peritrophin-A-Domain |
|  | GBRI008201 | Brevipalpis | GbrPLP16 | Peritrophin-Like Protein | Chitin-binding, Peritrophin-A-Domain |
|  | GFUI023663 | Fuscipes | GfuPLP16 | Peritrophin-Like Protein | Chitin-binding, Peritrophin-A-Domain |
|  | GPAI033432 | Palldipes | GpaPLP16 | Peritrophin-Like Protein | Chitin-binding, Peritrophin-A-Domain |
|  | GPPI023739 | Palpalis | GppPLP16 | Peritrophin-Like Protein | Chitin-binding, Peritrophin-A-Domain |
| FBgn0038492 | GMOY005278 | Morsitans | GmmPLP17 | Peritrophin-Like Protein | Chitin-binding, Peritrophin-A-Domain |
|  | GAUT029524 | Austeni | GauPLP17a | Peritrophin-Like Protein | Chitin-binding, Peritrophin-A-Domain |
|  | GAUT029525 | Austeni | GauPLP17b | Peritrophin-Like Protein | Chitin-binding, Peritrophin-A-Domain |
|  | GBRI026281 | Brevipalpis | GbrPLP17 | Peritrophin-Like Protein | Chitin-binding, Peritrophin-A-Domain |
|  | GFUI015251 | Fuscipes | GfuPLP17 | Peritrophin-Like Protein | Chitin-binding, Peritrophin-A-Domain |
|  | GPAI036992 | Palldipes | GpaPLP17 | Peritrophin-Like Protein | Chitin-binding, Peritrophin-A-Domain |
|  | GPPI048759 | Palpalis | GppPLP17a | Peritrophin-Like Protein | Chitin-binding, Peritrophin-A-Domain |
|  | GPPI042690 | Palpalis | GppPLP17b | Peritrophin-Like Protein | Chitin-binding, Peritrophin-A-Domain |
| FBgn0034030 | GMOY006713 | Morsitans | GmmPLP18 | Peritrophin-Like Protein | Chitin-binding, Peritrophin-A-Domain |
|  | GAUT023971 | Austeni | GauPLP18 | Peritrophin-Like Protein | Chitin-binding, Peritrophin-A-Domain |
|  | GFUI042924 | Fuscipes | GfuPLP18 | Peritrophin-Like Protein | Chitin-binding, Peritrophin-A-Domain |
|  | GPAI039717 | Palldipes | GpaPLP18 | Peritrophin-Like Protein | Chitin-binding, Peritrophin-A-Domain |
|  | GPPI013212 | Palpalis | GppPLP18 | Peritrophin-Like Protein | Chitin-binding, Peritrophin-A-Domain |
| FBgn0036203 | GMOY007191 | Morsitans | GmmPLP19  GmmPer108 | Peritrophin-Like Protein | Chitin-binding, Peritrophin-A-Domain |
|  | GAUT013406 | Austeni | GauPLP19 | Peritrophin-Like Protein | Chitin-binding, Peritrophin-A-Domain |
|  | GBRI016187 | Brevipalpis | GbrPLP19 | Peritrophin-Like Protein | Chitin-binding, Peritrophin-A-Domain |
|  | GFUI029181 | Fuscipes | GfuPLP19 | Peritrophin-Like Protein | Chitin-binding, Peritrophin-A-Domain |
|  | GPAI035996 | Palldipes | GpaPLP19 | Peritrophin-Like Protein | Chitin-binding, Peritrophin-A-Domain |
|  | GPPI010524 | Palpalis | GppPLP19 | Peritrophin-Like Protein | Chitin-binding, Peritrophin-A-Domain |
| FBgn0040601 | GMOY007476 | Morsitans | GmmPLP20 | Peritrophin-Like Protein | Chitin-binding, Partial Peritrophin-C-Domain |
|  | GAUT039321 | Austeni | GauPLP20 | Peritrophin-Like Protein | Chitin-binding, Partial Peritrophin-C-Domain |
|  | GBRI029781 | Brevipalpis | GbrPLP20 | Peritrophin-Like Protein | Chitin-binding, Partial Peritrophin-C-Domain |
|  | GFUI006871 | Fuscipes | GfuPLP20 | Peritrophin-Like Protein | Chitin-binding, Partial Peritrophin-C-Domain |
|  | GPAI013818 | Palldipes | GpaPLP20 | Peritrophin-Like Protein | Chitin-binding, Partial Peritrophin-C-Domain |
| FBgn0036845 | GMOY008030 | Morsitans | GmmPLP21 | Peritrophin-Like Protein | Chitin-binding, Peritrophin-A-Domain |
|  | GAUT042200 | Austeni | GauPLP21 | Peritrophin-Like Protein | Chitin-binding, Peritrophin-A-Domain |
|  | GBRI039692 | Brevipalpis | GbrPLP21 | Peritrophin-Like Protein | Chitin-binding, Peritrophin-A-Domain |
|  | GFUI011847 | Fuscipes | GfuPLP21 | Peritrophin-Like Protein | Chitin-binding, Peritrophin-A-Domain |
|  | GPAI019290 | Palldipes | GpaPLP21 | Peritrophin-Like Protein | Chitin-binding, Peritrophin-A-Domain |
|  | GPPI004759 | Palpalis | GppPLP21 | Peritrophin-Like Protein | Chitin-binding, Peritrophin-A-Domain |
| FBgn0035844 | GMOY008032 | Morsitans | GmmPLP22 | Peritrophin-Like Protein | Chitin-binding, Peritrophin-A-Domain |
|  | GAUT042214 | Austeni | GauPLP22 | Peritrophin-Like Protein | Chitin-binding, Peritrophin-A-Domain |
|  | GBRI039674 | Brevipalpis | GbrPLP22 | Peritrophin-Like Protein | Chitin-binding, Peritrophin-A-Domain |
|  | GFUI011826 | Fuscipes | GfuPLP22 | Peritrophin-Like Protein | Chitin-binding, Peritrophin-A-Domain |
|  | GPAI019295 | Palldipes | GpaPLP22 | Peritrophin-Like Protein | Chitin-binding, Peritrophin-A-Domain |
|  | GPPI004758 | Palpalis | GppPLP22 | Peritrophin-Like Protein | Chitin-binding, Peritrophin-A-Domain |
| FBgn0040959 | GMOY009587 | Morsitans | GmmPLP23/Pro2 | Peritrophin-Like Protein | Chitin-binding, Partial Peritrophin-C-Domain |
|  | GAUT008830 | Austeni | GauPLP23a | Peritrophin-Like Protein | Chitin-binding, Partial Peritrophin-C-Domain |
|  | GAUT024681 | Austeni | GauPLP23b | Peritrophin-Like Protein | Chitin-binding, Partial Peritrophin-C-Domain |
|  | GBRI017781 | Brevipalpis | GbrPLP23 | Peritrophin-Like Protein | Chitin-binding, Partial Peritrophin-C-Domain |
|  | GFUI019092 | Fuscipes | GfuPLP23 | Peritrophin-Like Protein | Chitin-binding, Partial Peritrophin-C-Domain |
|  | GPAI002755 | Palldipes | GpaPLP23 | Peritrophin-Like Protein | Chitin-binding, Partial Peritrophin-C-Domain |
|  | GPPI039564 | Palpalis | GppPLP23a | Peritrophin-Like Protein | Chitin-binding, Partial Peritrophin-C-Domain |
|  | GPPI022981 | Palpalis | GppPLP23b | Peritrophin-Like Protein | Chitin-binding, Partial Peritrophin-C-Domain |
|  | GPPI039559 | Palpalis | GppPLP23c | Peritrophin-Like Protein | Chitin-binding, Partial Peritrophin-C-Domain |
| FBgn0036230 | GMOY009788 | Morsitans | GmmPLP24 | Peritrophin-Like Protein | Chitin-binding, Peritrophin-A-Domain |
| FBgn0262986 | GAUT012373 | Austeni | GauPLP24 | Peritrophin-Like Protein | Chitin-binding, Peritrophin-A-Domain |
|  | GFUI027349 | Brevipalpis | GbrPLP24 | Peritrophin-Like Protein | Chitin-binding, Peritrophin-A-Domain |
|  | GPAI002186 | Palldipes | GpaPLP24 | Peritrophin-Like Protein | Chitin-binding, Peritrophin-A-Domain |
|  | GPPI017724 | Palpalis | GppPLP24 | Peritrophin-Like Protein | Chitin-binding, Peritrophin-A-Domain |
| FBgn0036230 | GMOY009805 | Morsitans | GmmPLP25 | Peritrophin-Like Protein | Chitin-binding, Peritrophin-A-Domain |
| FBgn0262986 | GAUT012335 | Austeni | GauPLP25 | Peritrophin-Like Protein | Chitin-binding, Peritrophin-A-Domain |
|  | GFUI026256 | Fuscipes | GfuPLP25 | Peritrophin-Like Protein | Chitin-binding, Peritrophin-A-Domain |
|  | GPAI002122 | Palldipes | GpaPLP25a | Peritrophin-Like Protein | Chitin-binding, Peritrophin-A-Domain |
|  | GPAI002124 | Palldipes | GpaPLP25b | Peritrophin-Like Protein | Chitin-binding, Peritrophin-A-Domain |
|  | GPPI016003 | Palpalis | GppPLP25 | Peritrophin-Like Protein | Chitin-binding, Peritrophin-A-Domain |
| FBgn0036226 | GMOY009806 | Morsitans | GmmPLP26a | Peritrophin-Like Protein | Chitin-binding, Peritrophin-A-Domain |
|  | GMOY009807 | Morsitans | GmmPLP26b | Peritrophin-Like Protein | Chitin-binding, Peritrophin-A-Domain |
|  | GAUT012312 | Austeni | GauPLP26a | Peritrophin-Like Protein | Chitin-binding, Peritrophin-A-Domain |
|  | GAUT012316 | Austeni | GauPLP26b | Peritrophin-Like Protein | Chitin-binding, Peritrophin-A-Domain |
|  | GBRI029501 | Brevipalpis | GbrPLP26 | Peritrophin-Like Protein | Chitin-binding, Peritrophin-A-Domain |
|  | GFUI026249 | Fuscipes | GfuPLP26a | Peritrophin-Like Protein | Chitin-binding, Peritrophin-A-Domain |
|  | GFUI026252 | Fuscipes | GfuPLP26b | Peritrophin-Like Protein | Chitin-binding, Peritrophin-A-Domain |
|  | GPAI002108 | Palldipes | GpaPLP26 | Peritrophin-Like Protein | Chitin-binding, Peritrophin-A-Domain |
|  | GPPI016000 | Palpalis | GppPLP26 | Peritrophin-Like Protein | Chitin-binding, Peritrophin-A-Domain |
| FBgn0022770 | GMOY009571 | Morsitans | GmmPLP27 | Peritrophin-Like Protein | Chitin-binding, Peritrophin-A-Domain |
|  | GAUT032039 | Austeni | GauPLP27 | Peritrophin-Like Protein | Chitin-binding, Peritrophin-A-Domain |
|  | GFUI025256 | Fuscipes | GfuPLP27 | Peritrophin-Like Protein | Chitin-binding, Peritrophin-A-Domain |
|  | GPAI004770 | Palldipes | GpaPLP27 | Peritrophin-Like Protein | Chitin-binding, Peritrophin-A-Domain |
|  | GPPI004131 | Palpalis | GppPLP27 | Peritrophin-Like Protein | Chitin-binding, Peritrophin-A-Domain |
| FBgn0031097 | GMOY009572 | Morsitans | GmmPLP28 | Peritrophin-Like Protein | Chitin-binding, Peritrophin-A-Domain |
|  | GAUT032040 | Austeni | GauPLP28 | Peritrophin-Like Protein | Chitin-binding, Peritrophin-A-Domain |
|  | GBRI003555 | Brevipalpis | GbrPLP28 | Peritrophin-Like Protein | Chitin-binding, Peritrophin-A-Domain |
|  | GFUI025254 | Fuscipes | GfuPLP28 | Peritrophin-Like Protein | Chitin-binding, Peritrophin-A-Domain |
|  | GPAI004766 | Palldipes | GpaPLP28 | Peritrophin-Like Protein | Chitin-binding, Peritrophin-A-Domain |
|  | GPPI004133 | Palpalis | GppPLP28 | Peritrophin-Like Protein | Chitin-binding, Peritrophin-A-Domain |
| FBgn0026077 | GMOY009826 | Morsitans | GmmPLP29 | Peritrophin-Like Protein | Chitin-binding, Peritrophin-A-Domain |
|  | GAUT041130 | Austeni | GauPLP29 | Peritrophin-Like Protein | Chitin-binding, Peritrophin-A-Domain |
|  | GBRI031719 | Brevipalpis | GbrPLP29 | Peritrophin-Like Protein | Chitin-binding, Peritrophin-A-Domain |
|  | GFUI037199 | Fuscipes | GfuPLP29a | Peritrophin-Like Protein | Chitin-binding, Peritrophin-A-Domain |
|  | GFUI049166 | Fuscipes | GfuPLP29b | Peritrophin-Like Protein | Chitin-binding, Peritrophin-A-Domain |
|  | GPA1031701 | Palldipes | GpaPLP29 | Peritrophin-Like Protein | Chitin-binding, Peritrophin-A-Domain |
|  | GMOY011054 | Morsitans | GmmPLP30 | Peritrophin-Like Protein | Chitin-binding, Peritrophin-A-Domain |
|  | GAUT026340 | Austeni | GauPLP30 | Peritrophin-Like Protein | Chitin-binding, Peritrophin-A-Domain |
|  | GBRI013108 | Brevipalpis | GbrPLP30 | Peritrophin-Like Protein | Chitin-binding, Peritrophin-A-Domain |
|  | GFUI045125 | Fuscipes | GfuPLP30 | Peritrophin-Like Protein | Chitin-binding, Peritrophin-A-Domain |
|  | GPAI027171 | Palldipes | GpaPLP30 | Peritrophin-Like Protein | Chitin-binding, Peritrophin-A-Domain |
|  | GPPI005116 | Palpalis | GppPLP30 | Peritrophin-Like Protein | Chitin-binding, Peritrophin-A-Domain |
| FBgn0039452 | GMOY011777 | Morsitans | GmmPLP31 | Peritrophin-Like Protein | Chitin-binding, Partial Peritrophin-C-Domain |
|  | GAUT036980 | Austeni | GauPLP31a | Peritrophin-Like Protein | Chitin-binding, Partial Peritrophin-C-Domain |
|  | GAUT039327 | Austeni | GauPLP31b | Peritrophin-Like Protein | Chitin-binding, Partial Peritrophin-C-Domain |
|  | GBRI013425 | Brevipalpis | GbrPLP31a | Peritrophin-Like Protein | Chitin-binding, Partial Peritrophin-C-Domain |
|  | GBRI034385 | Brevipalpis | GbrPLP31b | Peritrophin-Like Protein | Chitin-binding, Partial Peritrophin-C-Domain |
|  | GFUI006839 | Fuscipes | GfuPLP31a | Peritrophin-Like Protein | Chitin-binding, Partial Peritrophin-C-Domain |
|  | GFUI006847 | Fuscipes | GfuPLP31b | Peritrophin-Like Protein | Chitin-binding, Partial Peritrophin-C-Domain |
|  | GPAI013834 | Palldipes | GpaPLP31a | Peritrophin-Like Protein | Chitin-binding, Partial Peritrophin-C-Domain |
|  | GPAI013840 | Palldipes | GpaPLP31b | Peritrophin-Like Protein | Chitin-binding, Partial Peritrophin-C-Domain |
|  | GPPI050938 | Palpalis | GppPLP31a | Peritrophin-Like Protein | Chitin-binding, Partial Peritrophin-C-Domain |
|  | GPPI050939 | Palpalis | GppPLP31b | Peritrophin-Like Protein | Chitin-binding, Partial Peritrophin-C-Domain |
| FBgn0038643 | GMOY011809 | Morsitans | GmmPLP32/Pro1 | Peritrophin-Like Protein | Chitin-binding, Partial Peritrophin-C-Domain |
|  | GBRI007305 | Brevipalpis | GbrPLP32 | Peritrophin-Like Protein | Chitin-binding, Partial Peritrophin-C-Domain |
|  | GPPI040997 | Palpalis | GpaPLP32 | Peritrophin-Like Protein | Chitin-binding, Partial Peritrophin-C-Domain |
| FBgn0085311 | GMOY011810 | Morsitans | GmmPLP33  GmmPer12 | Peritrophin-Like Protein | Chitin-binding, Partial Peritrophin-C-Domain |
|  | GAUT038910 | Austeni | GauPLP33 | Peritrophin-Like Protein | Chitin-binding, Partial Peritrophin-C-Domain |
|  | GBRI007300 | Brevipalpis | GbrPLP33a | Peritrophin-Like Protein | Chitin-binding, Partial Peritrophin-C-Domain |
|  | GBRI007301 | Brevipalpis | GbrPLP33b | Peritrophin-Like Protein | Chitin-binding, Partial Peritrophin-C-Domain |
|  | GFUI000259 | Fuscipes | GfuPLP33 | Peritrophin-Like Protein | Chitin-binding, Partial Peritrophin-C-Domain |
|  | GPPI040977 | Palpalis | GppPLP33 | Peritrophin-Like Protein | Chitin-binding, Partial Peritrophin-C-Domain |
|  | GAUT012314 | Austeni | GauPLP34 | Peritrophin-Like Protein | Chitin-binding, Peritrophin-A-Domain, mucin domains |
|  | GBRI029506 | Brevipalpis | GbrPLP34 | Peritrophin-Like Protein | Chitin-binding, Peritrophin-A-Domain, mucin domains |
|  | GFUI026251 | Fuscipes | GfuPLP34 | Peritrophin-Like Protein | Chitin-binding, Peritrophin-A-Domain, mucin domains |
|  | GPAI002117 | Palldipes | GpaPLP34 | Peritrophin-Like Protein | Chitin-binding, Peritrophin-A-Domain, mucin domains |
|  | GPPI016008 | Palpalis | GppPLP34 | Peritrophin-Like Protein | Chitin-binding, Peritrophin-A-Domain, mucin domains |
|  | GAUT044451 | Austeni | GauPLP35 | Peritrophin-Like Protein | Chitin-binding, Peritrophin-A-Domain |
|  | GBRI023665 | Brevipalpis | GbrPLP35 | Peritrophin-Like Protein | Chitin-binding, Peritrophin-A-Domain |
|  | GFUI048633 | Fuscipes | GfuPLP35 | Peritrophin-Like Protein | Chitin-binding, Peritrophin-A-Domain |
|  | GPPI048544 | Palpalis | GppPLP35 | Peritrophin-Like Protein | Chitin-binding, Peritrophin-A-Domain |
|  | GAUT036465 | Austeni | GauPLP36 | Peritrophin-Like Protein | Chitin-binding, Partial Peritrophin-C-Domain |
|  | GBRI019611 | Brevipalpis | GbrPLP36 | Peritrophin-Like Protein | Chitin-binding, Partial Peritrophin-C-Domain |
|  | GPAI033924 | Palldipes | GppPLP36 | Peritrophin-Like Protein | Chitin-binding, Partial Peritrophin-C-Domain |
| FBgn0040958 | GAUT051817 | Austeni | GauPLP37 | Peritrophin-Like Protein | Chitin-binding, Partial Peritrophin-C-Domain |
|  | GBRI017597 | Brevipalpis | GbrPLP37 | Peritrophin-Like Protein | Chitin-binding, Partial Peritrophin-C-Domain |
|  | GFUI036185 | Fuscipes | GfuPLP37 | Peritrophin-Like Protein | Chitin-binding, Partial Peritrophin-C-Domain |
| FBgn0259192 | GBRI023004 | Brevipalpis | GbrPLP38 | Peritrophin-Like Protein | Chitin-binding, Peritrophin-A-Domain |
|  | GFUI007992 | Fuscipes | GfuPLP38 | Peritrophin-Like Protein | Chitin-binding, Peritrophin-A-Domain |
|  | GPAI006518 | Palldipes | GpaPLP38 | Peritrophin-Like Protein | Chitin-binding, Peritrophin-A-Domain |
|  | GPPI046699 | Palpalis | GppPLP38 | Peritrophin-Like Protein | Chitin-binding, Peritrophin-A-Domain |
|  | GBRI024198 | Brevipalpis | GbrPLP39 | Peritrophin-Like Protein | Chitin-binding, Peritrophin-A-Domain |
|  | GFUI022163 | Fuscipes | GfuPLP39 | Peritrophin-Like Protein | Chitin-binding, Peritrophin-A-Domain |
|  | GPAI033201 | Palldipes | GpaPLP39 | Peritrophin-Like Protein | Chitin-binding, Peritrophin-A-Domain |
|  | GPPI000357 | Palpalis | GppPLP39 | Peritrophin-Like Protein | Chitin-binding, Peritrophin-A-Domain |
|  | GBRI027291 | Brevipalpis | GbrPLP40 | Peritrophin-Like Protein | Chitin-binding, Peritrophin-A-Domain |
|  | GPAI016225 | Palldipes | GpaPLP40 | Peritrophin-Like Protein | Chitin-binding, Peritrophin-A-Domain |
|  | GFUI037693 | Fuscipes | GfuPLP41 | Peritrophin-Like Protein | Chitin-binding, Peritrophin-A-Domain |
|  | GPPI030394 | Palpalis | GppPLP41 | Peritrophin-Like Protein | Chitin-binding, Peritrophin-A-Domain |
|  | GFUI022165 | Fuscipes | GfuPLP42 | Peritrophin-Like Protein | Chitin-binding, Peritrophin-A-Domain |
