## Supplemental Data 7 Transcription Factor Data for "The *Glossina* Genome Cluster: Comparative Genomic Analysis of the Vectors of African Trypanosomes": README.txt

This directory contains the following files, for all six Glossina species:

1. TF_motif_inferences_SPECIES.txt

This file (tab delimited) contains the predicted TFs, and all of their predicted DNA binding motifs.  The columns in this file indicate (left to right):
1 Gene ID
2 Protein ID
3 DNA binding domains contained within the TF (N to C terminus order).  PFAM IDs are provided.
4 Species from which the DNA binding motif was mapped.
5 Gene from which the DNA binding motif was mapped.
6 Number of mismatches in the amino acid alignment between the two DBDs.
7 Percent identity between the two DBDs (i.e., the fraction of identical amino acids in the DBD alignment).
8 Study from which the DNA binding motif was obtained.
9 ID of the DNA binding motif from the indicated study.
10 Type of experiment used to derive the motif. B1H = Bacteria one-hypbrid; Compiled = compiled from various sources; PBM = protein binding microarray.
11 IUPAC sequence of the DNA binding motif (forward direction).
12 IUPAC sequence of the DNA binding motif (reverse direction).
13-15 Source of the motif


2. TF_no_motif_SPECIES.txt
This file contains all predicted TFs that do not have a predicted DNA binding motif.


3. Logos_SPECIES (directory of .png files)
This directory contains "sequence logos" (forward and reverse) for the best motif for each TF.


4. PWMs_SPECIES (directory of .txt files)
This directory contains position weight matrices (PWMs) in the for of frequencies for the best motif for each TF. (This is the same format used by the CisBP database).


An updated list of motifs can be obtained from the CisBP website (http://cisbp.ccbr.utoronto.ca/).

Questions can be directed to Matt Weirauch
