## Supplementary figures and images for "The *Glossina* Genome Cluster: Comparative Genomic Analysis of the Vectors of African Trypanosomes"

### GAUT012024-PA_Homeobox_fwd.png

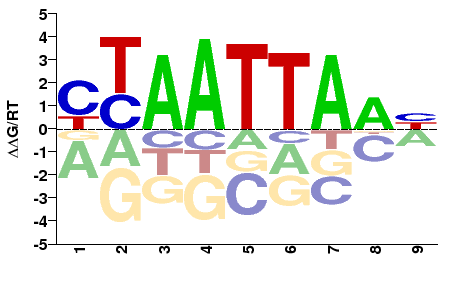

### GAUT012024-PA_Homeobox_rev.png

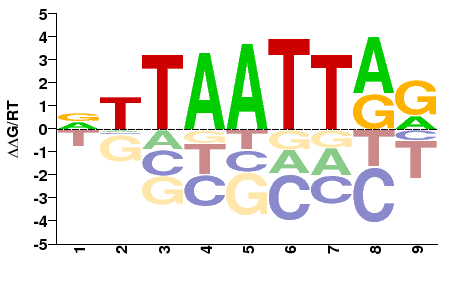

### GAUT012144-PA_T-box_fwd.png

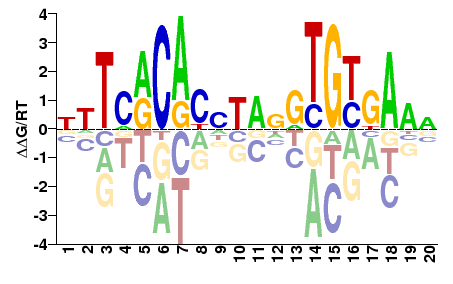

### GAUT012144-PA_T-box_rev.png

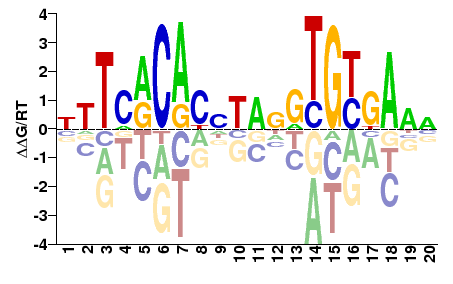

### GAUT012315-PA_zf-C2H2_fwd.png

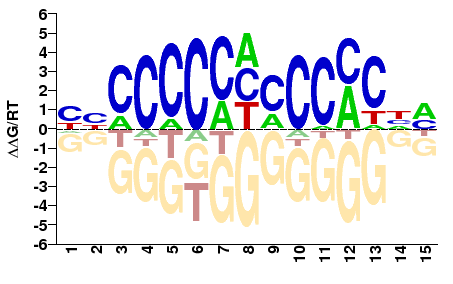

### GAUT012315-PA_zf-C2H2_rev.png

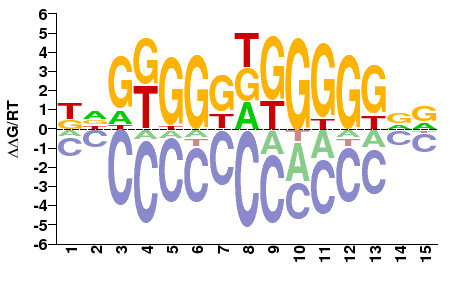

### GAUT012582-PA_HLH_fwd.png

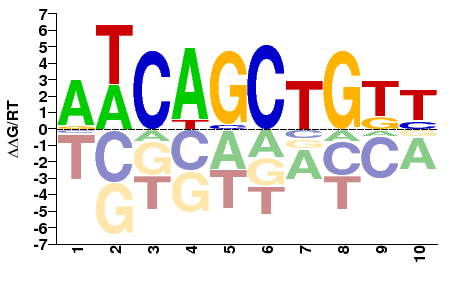

### GAUT012582-PA_HLH_rev.png

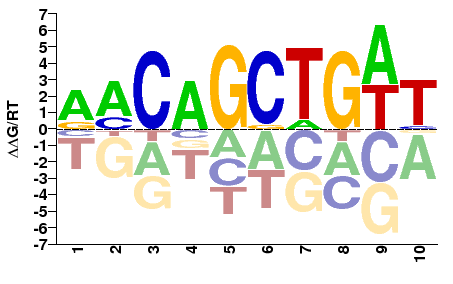

### GAUT013196-PA_zf-C2H2_fwd.png

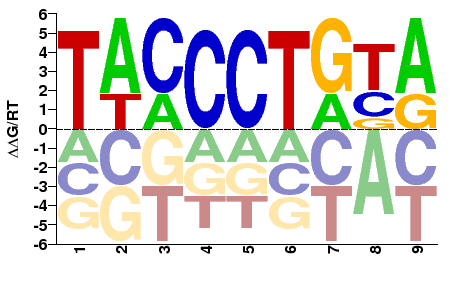

### GAUT013196-PA_zf-C2H2_rev.png

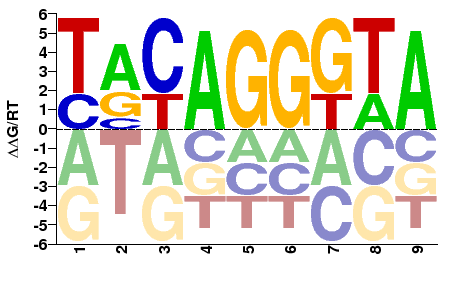

### GAUT013277-PA_Homeobox_fwd.png

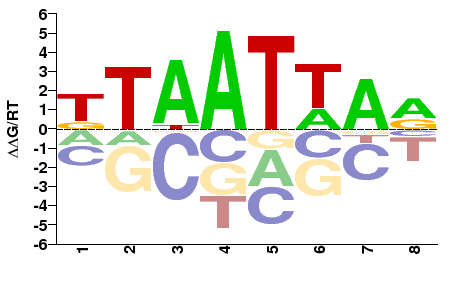

### GAUT013277-PA_Homeobox_rev.png

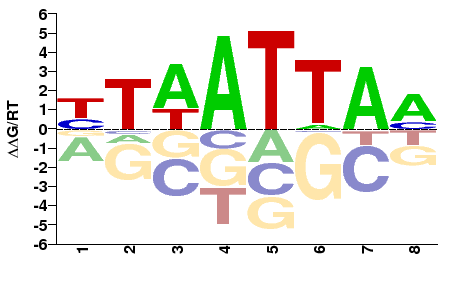

### GAUT013292-PA_Homeobox_fwd.png

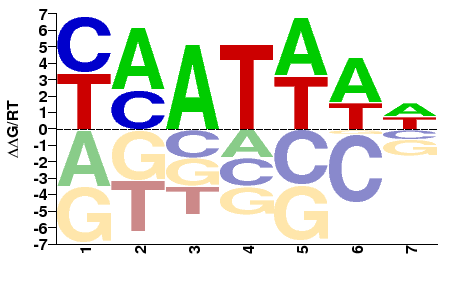

### GAUT013292-PA_Homeobox_rev.png

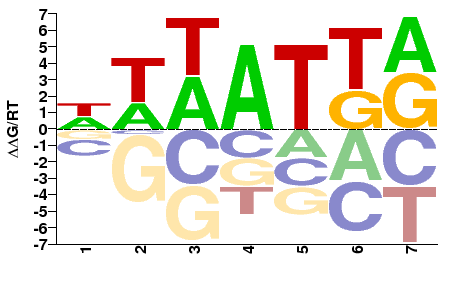

### GAUT013430-PA_MADF_DNA_bdg_fwd.png

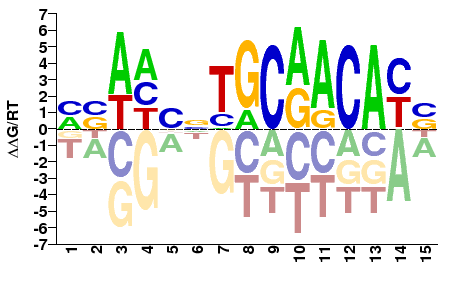

### GAUT013430-PA_MADF_DNA_bdg_rev.png

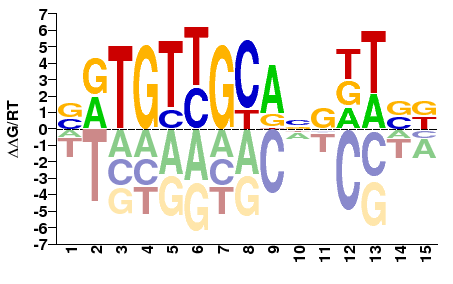

### GAUT013481-PA_Homeobox_fwd.png

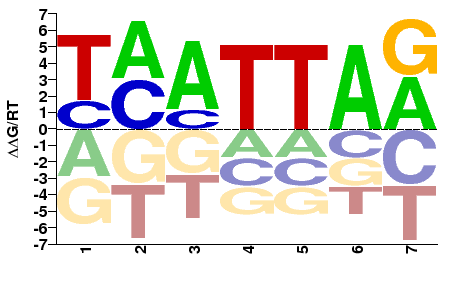

### GAUT013481-PA_Homeobox_rev.png

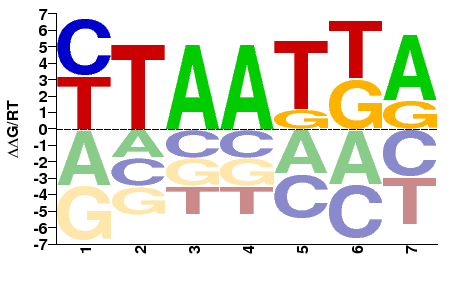

### GAUT013608-PA_zf-C2H2_fwd.png

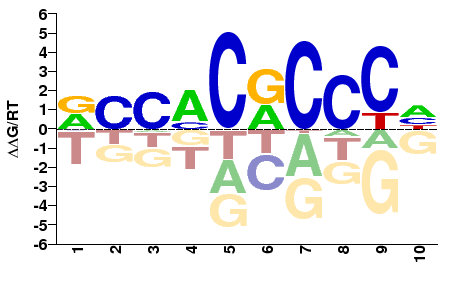

### GAUT013608-PA_zf-C2H2_rev.png

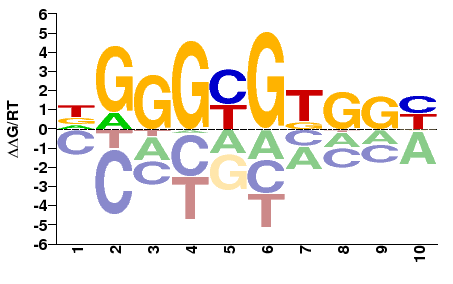

### GAUT013656-PA_Homeobox_fwd.png

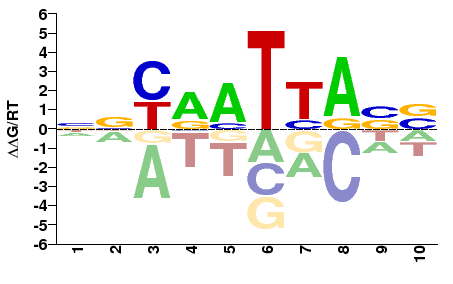

### GAUT013656-PA_Homeobox_rev.png

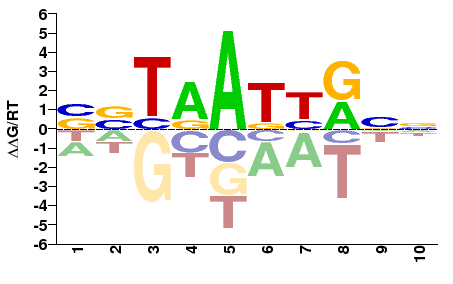

### GAUT014094-PA_MADF_DNA_bdg_fwd.png

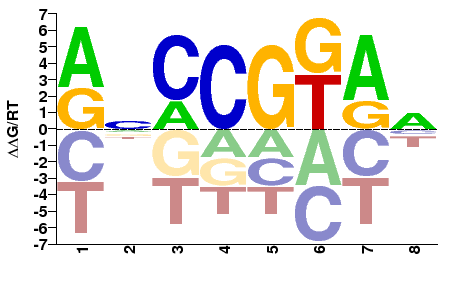

### GAUT014094-PA_MADF_DNA_bdg_rev.png

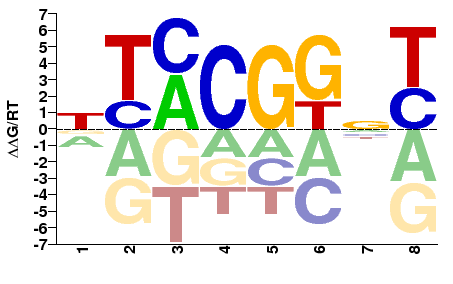

### GAUT014130-PA_zf-C2H2_fwd.png

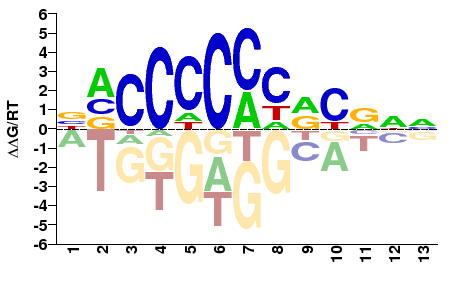

### GAUT014130-PA_zf-C2H2_rev.png

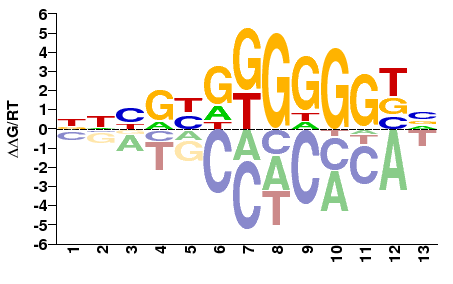

### GAUT015377-PA_bZIP_1_fwd.png

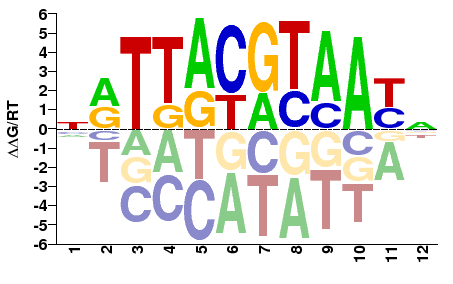

### GAUT015377-PA_bZIP_1_rev.png

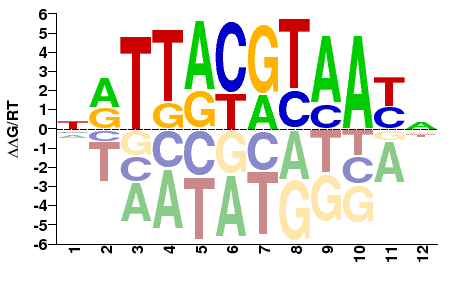

### GAUT015520-PA_Homeobox_fwd.png

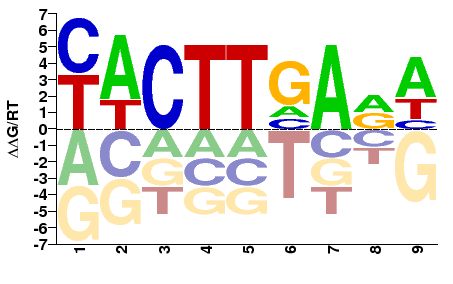

### GAUT015520-PA_Homeobox_rev.png

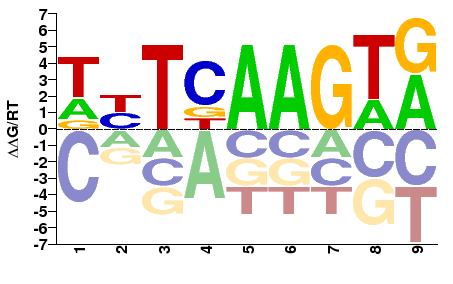

### GAUT016036-PA_HLH_fwd.png

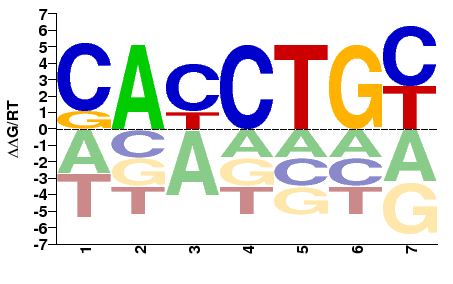

### GAUT016036-PA_HLH_rev.png

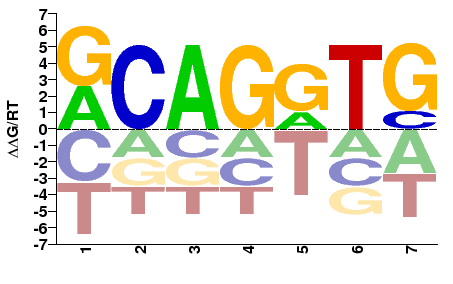

### GAUT016057-PA_HLH_fwd.png

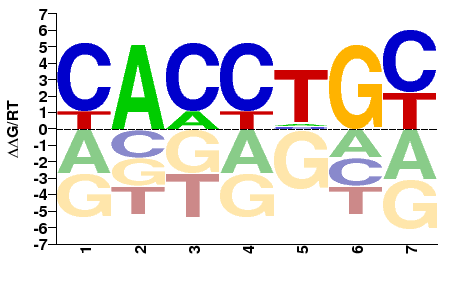

### GAUT016057-PA_HLH_rev.png

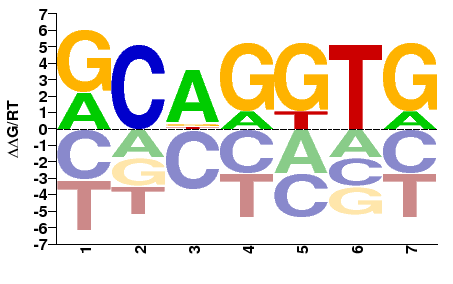

### GAUT016064-PA_HLH_fwd.png

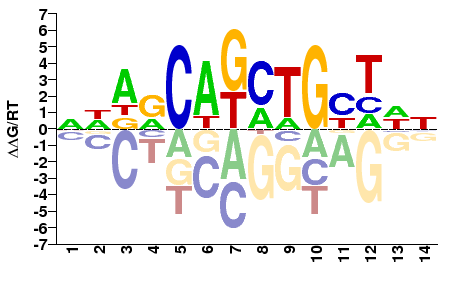

### GAUT016064-PA_HLH_rev.png

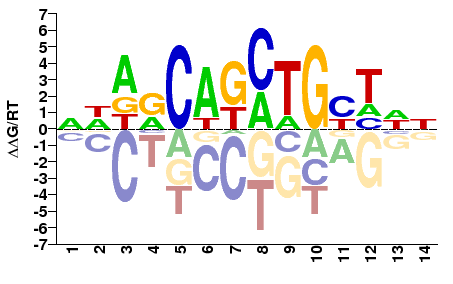

### GAUT016075-PA_HLH_fwd.png

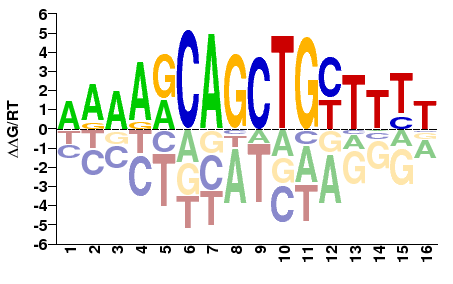

### GAUT016075-PA_HLH_rev.png

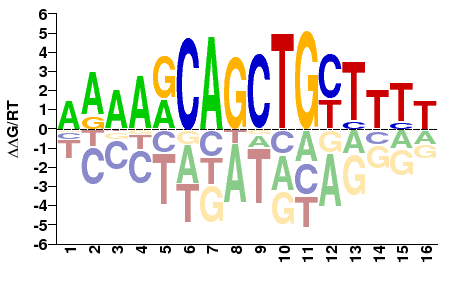

### GAUT016631-PA_MADF_DNA_bdg_fwd.png

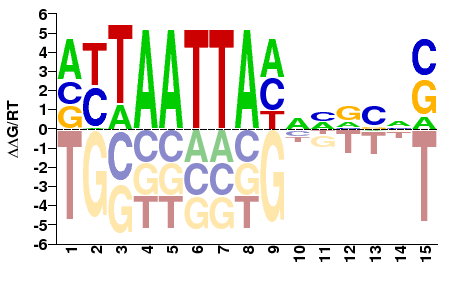

### GAUT016631-PA_MADF_DNA_bdg_rev.png

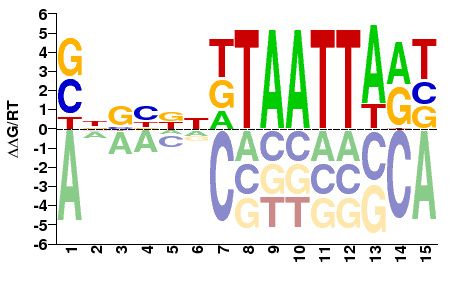

### GAUT017030-PA_zf-C4_fwd.png

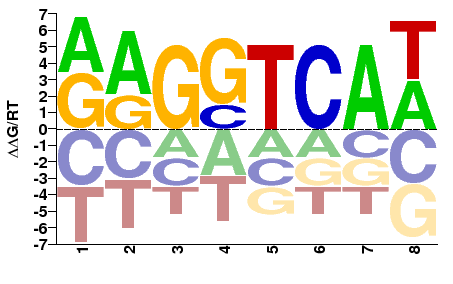

### GAUT017030-PA_zf-C4_rev.png

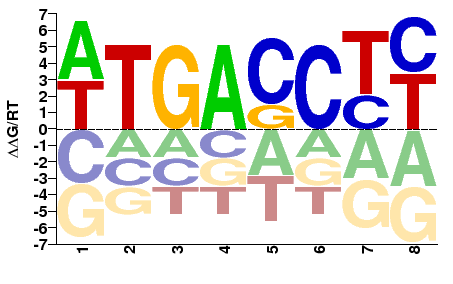

### GAUT017322-PA_HLH_fwd.png

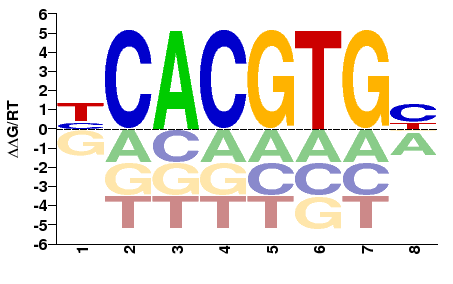

### GAUT017322-PA_HLH_rev.png

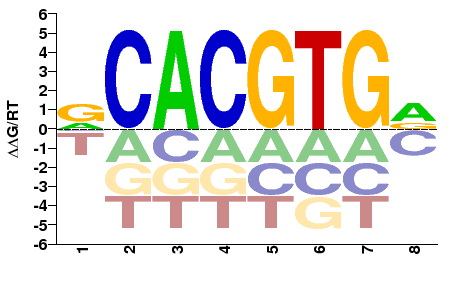

### GAUT017359-PA_MADF_DNA_bdg_fwd.png

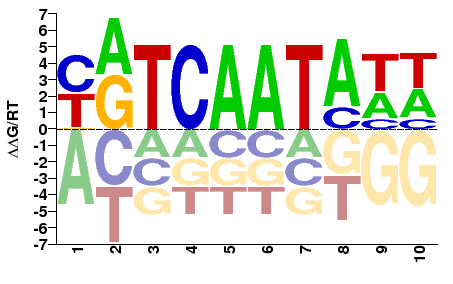

### GAUT017359-PA_MADF_DNA_bdg_rev.png

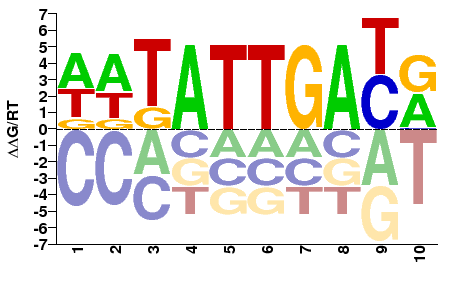

### GAUT017373-PA_Homeobox_fwd.png

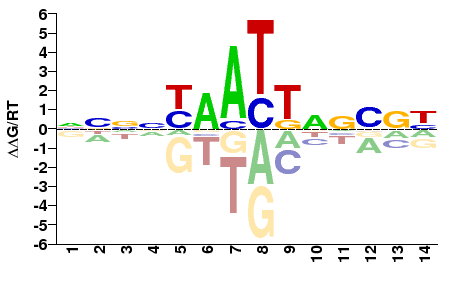

### GAUT017373-PA_Homeobox_rev.png

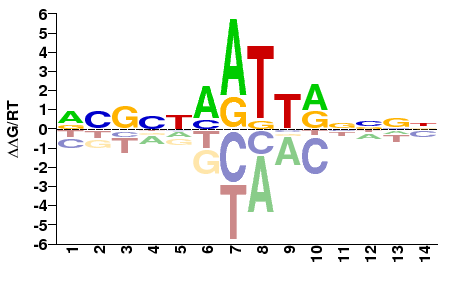

### GAUT017893-PA_Homeobox_fwd.png

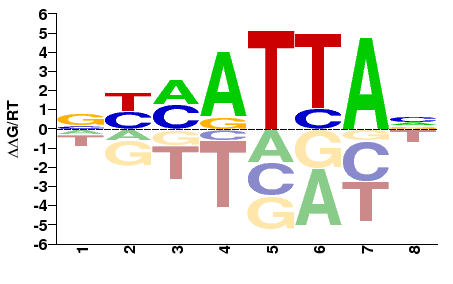

### GAUT017893-PA_Homeobox_rev.png

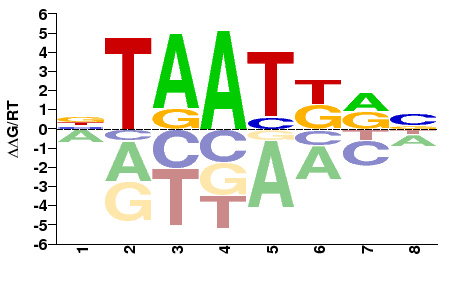

### GAUT017906-PA_Homeobox_fwd.png

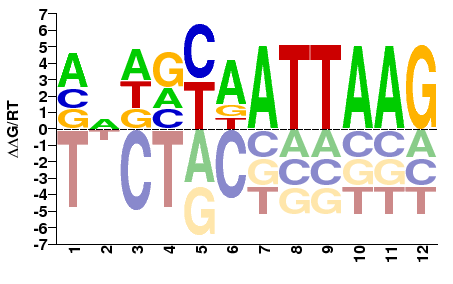

### GAUT017906-PA_Homeobox_rev.png

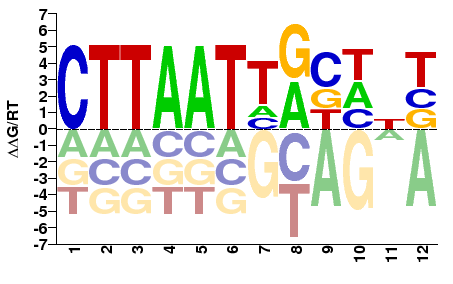

### GAUT018270-PA_HLH_fwd.png

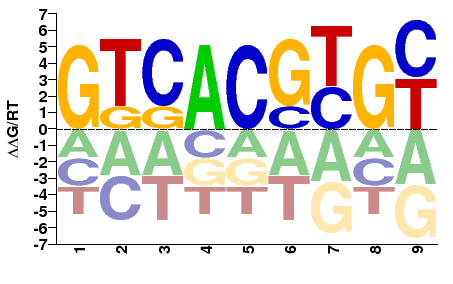

### GAUT018270-PA_HLH_rev.png

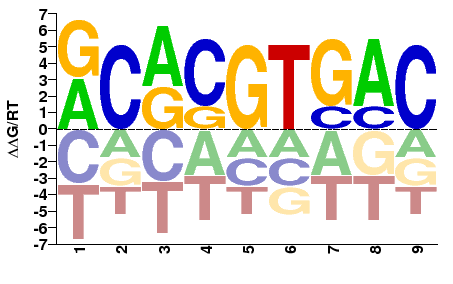

### GAUT018814-PA_RHD_DNA_bind_fwd.png

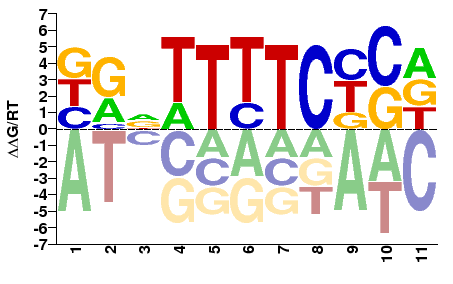

### GAUT018814-PA_RHD_DNA_bind_rev.png

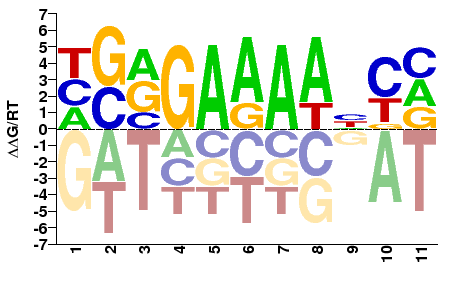

### GAUT018860-PA_zf-C2H2_fwd.png

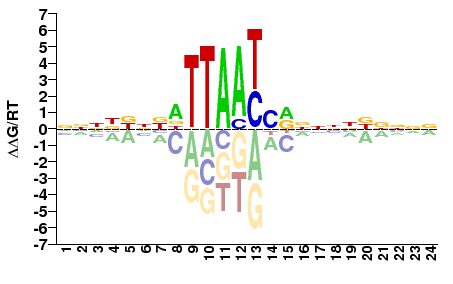

### GAUT018860-PA_zf-C2H2_rev.png

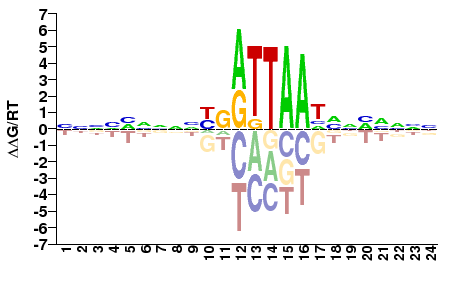

### GAUT018927-PA_Homeobox_fwd.png

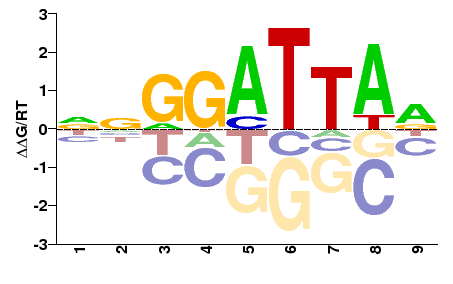

### GAUT018927-PA_Homeobox_rev.png

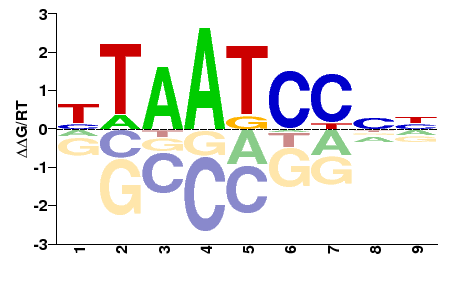

### GAUT018954-PA_zf-C2H2_fwd.png

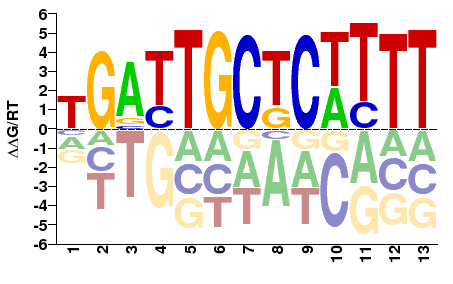

### GAUT018954-PA_zf-C2H2_rev.png

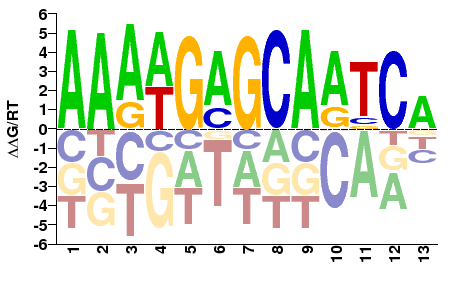

### GAUT019201-PA_zf-C2H2_fwd.png

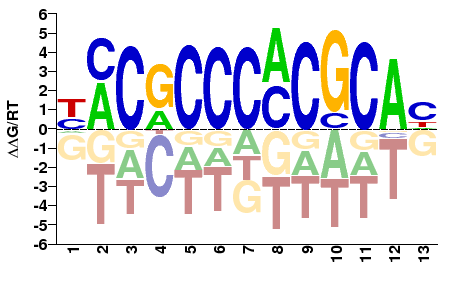

### GAUT019201-PA_zf-C2H2_rev.png

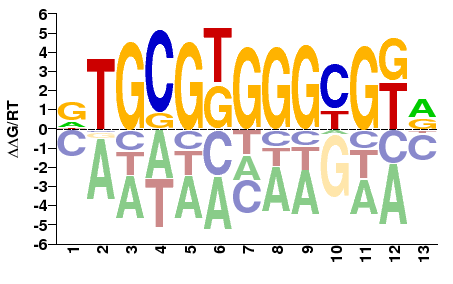

### GAUT019261-PA_Homeobox_fwd.png

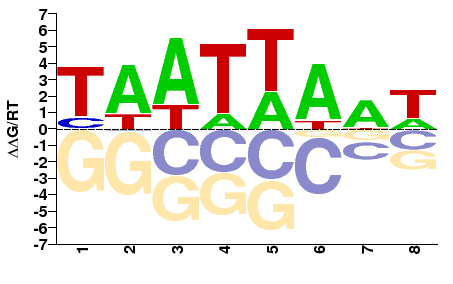

### GAUT019261-PA_Homeobox_rev.png

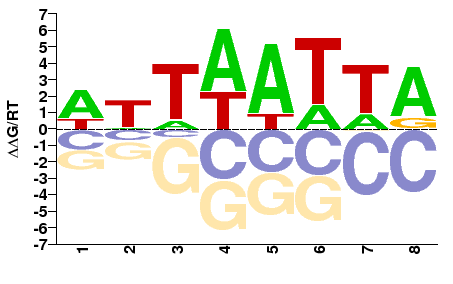

### GAUT019304-PA_zf-C4_fwd.png

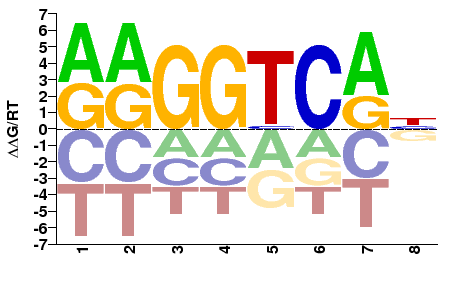

### GAUT019304-PA_zf-C4_rev.png

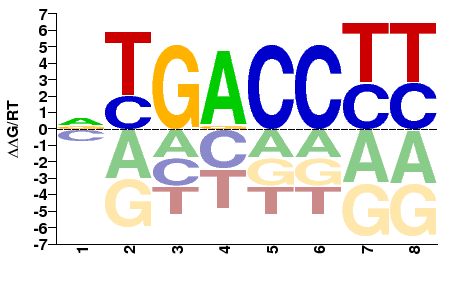

### GAUT019327-PA_HLH_fwd.png

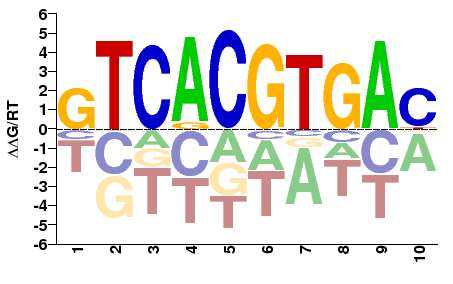

### GAUT019327-PA_HLH_rev.png

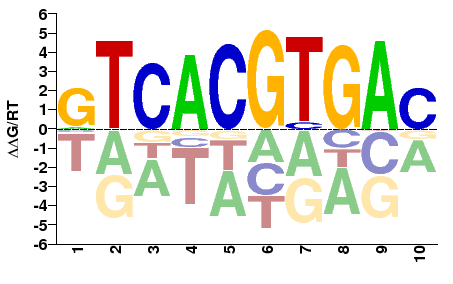

### GAUT019915-PA_Homeobox_fwd.png

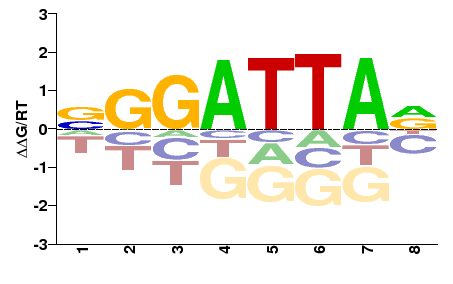

### GAUT019915-PA_Homeobox_rev.png

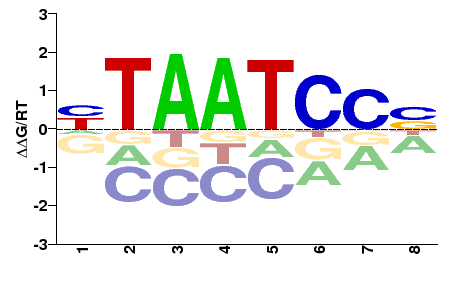

### GAUT020302-PA_MADF_DNA_bdg_fwd.png

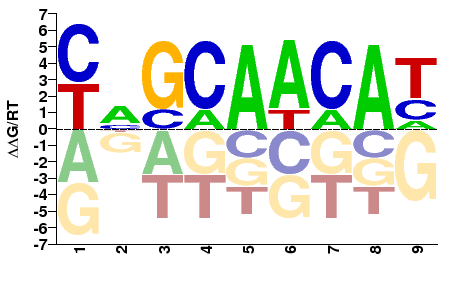

### GAUT020302-PA_MADF_DNA_bdg_rev.png

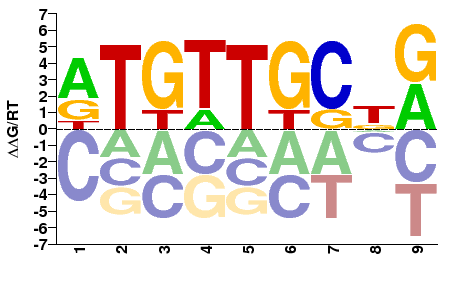

### GAUT020317-PA_HLH_fwd.png

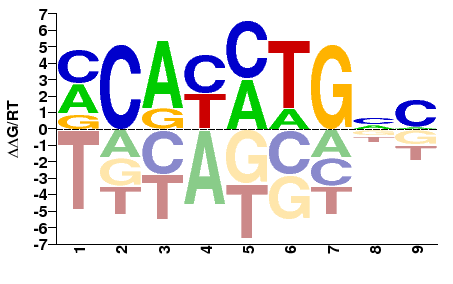

### GAUT020317-PA_HLH_rev.png

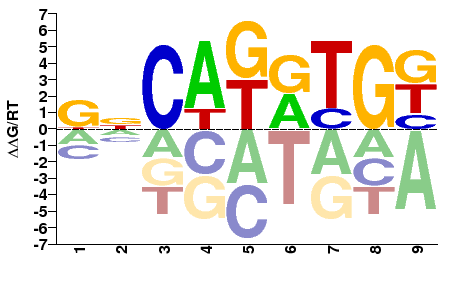

### GAUT020434-PA_HLH_fwd.png

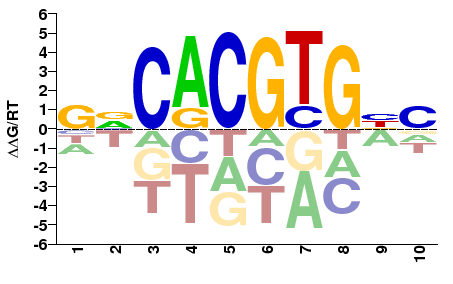

### GAUT020434-PA_HLH_rev.png

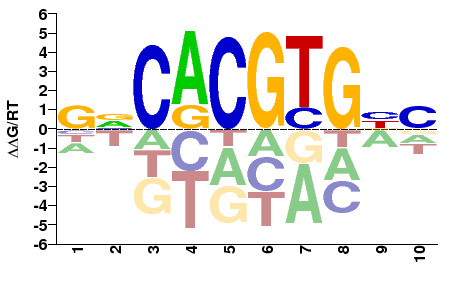

### GAUT021275-PA_Homeobox_fwd.png

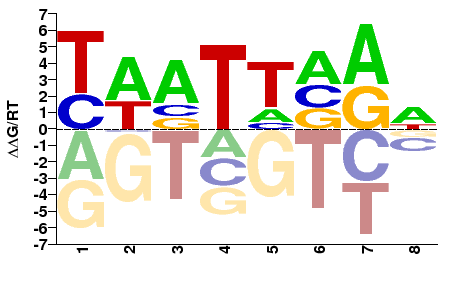

### GAUT021275-PA_Homeobox_rev.png

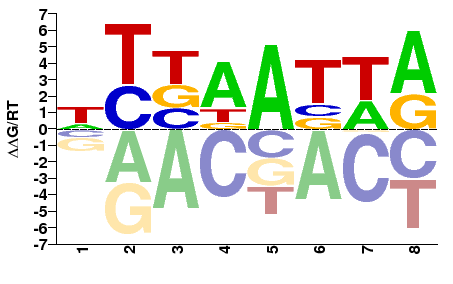

### GAUT021433-PA_zf-C2H2_fwd.png

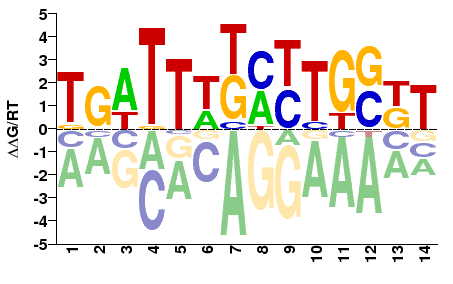

### GAUT021433-PA_zf-C2H2_rev.png

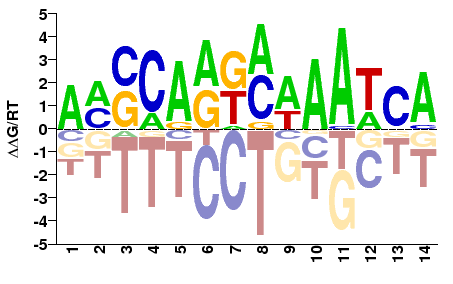

### GAUT021502-PA_zf-C2H2_fwd.png

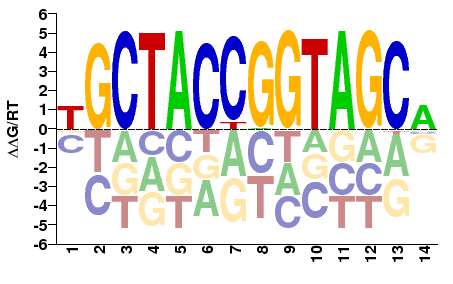

### GAUT021502-PA_zf-C2H2_rev.png

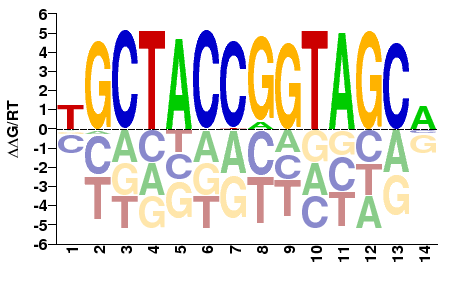

### GAUT021503-PA_zf-C2H2_fwd.png

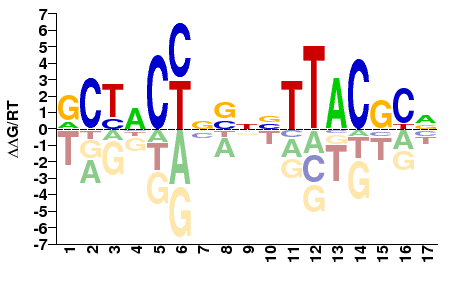

### GAUT021503-PA_zf-C2H2_rev.png

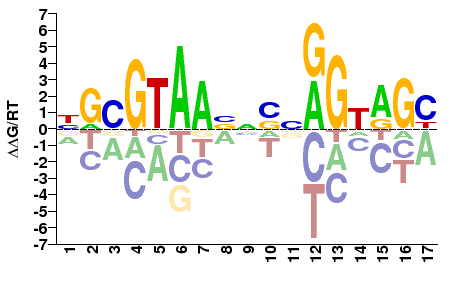

### GAUT022685-PA_Homeobox_fwd.png

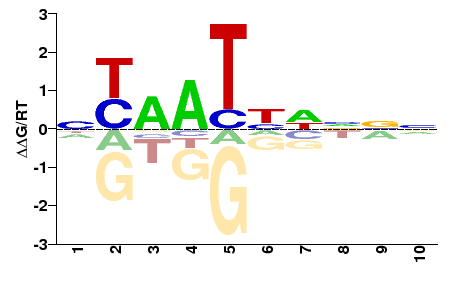

### GAUT022685-PA_Homeobox_rev.png

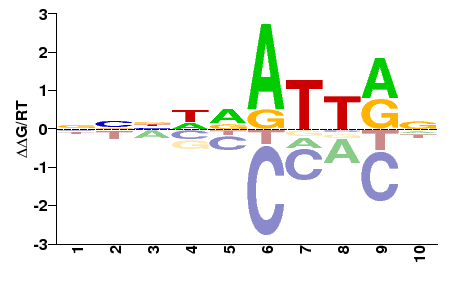

### GAUT023713-PA_HMG_box_fwd.png

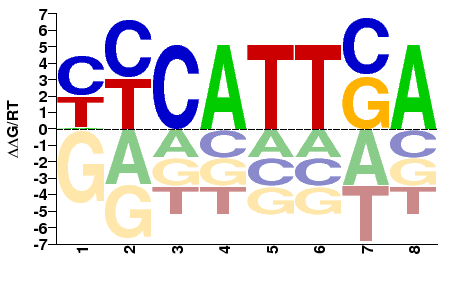

### GAUT023713-PA_HMG_box_rev.png

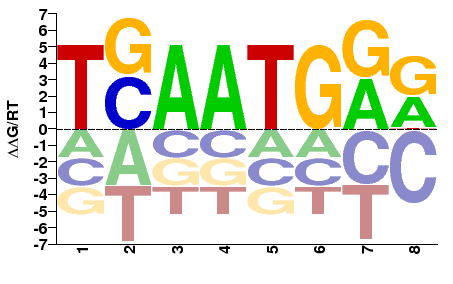

### GAUT023983-PA_HLH_fwd.png

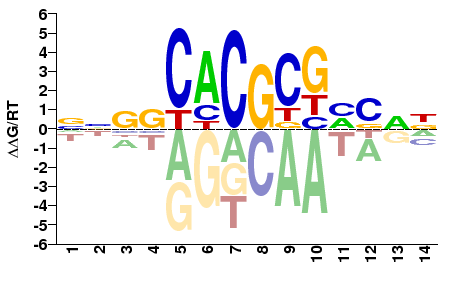

### GAUT023983-PA_HLH_rev.png

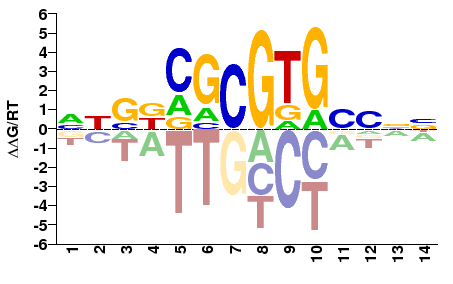

### GAUT024062-PA_Homeobox_fwd.png

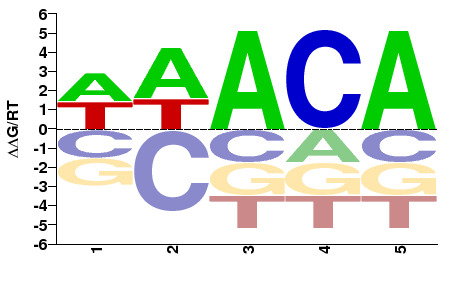

### Supplemental Figure 2

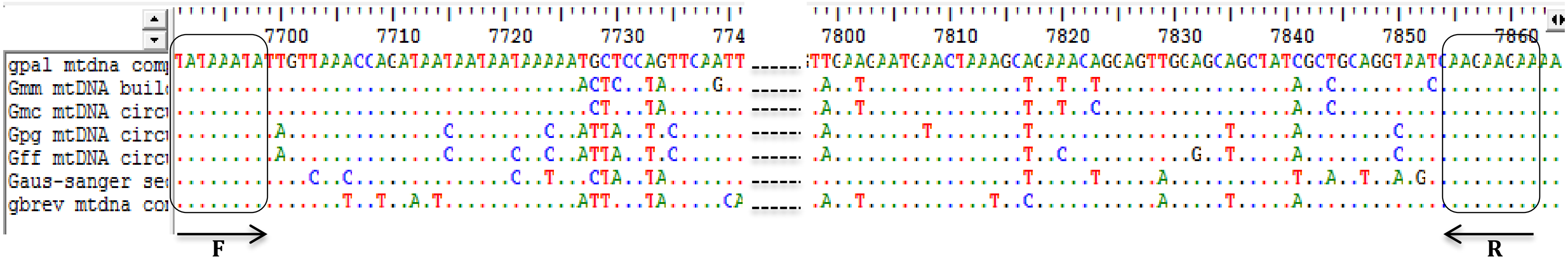

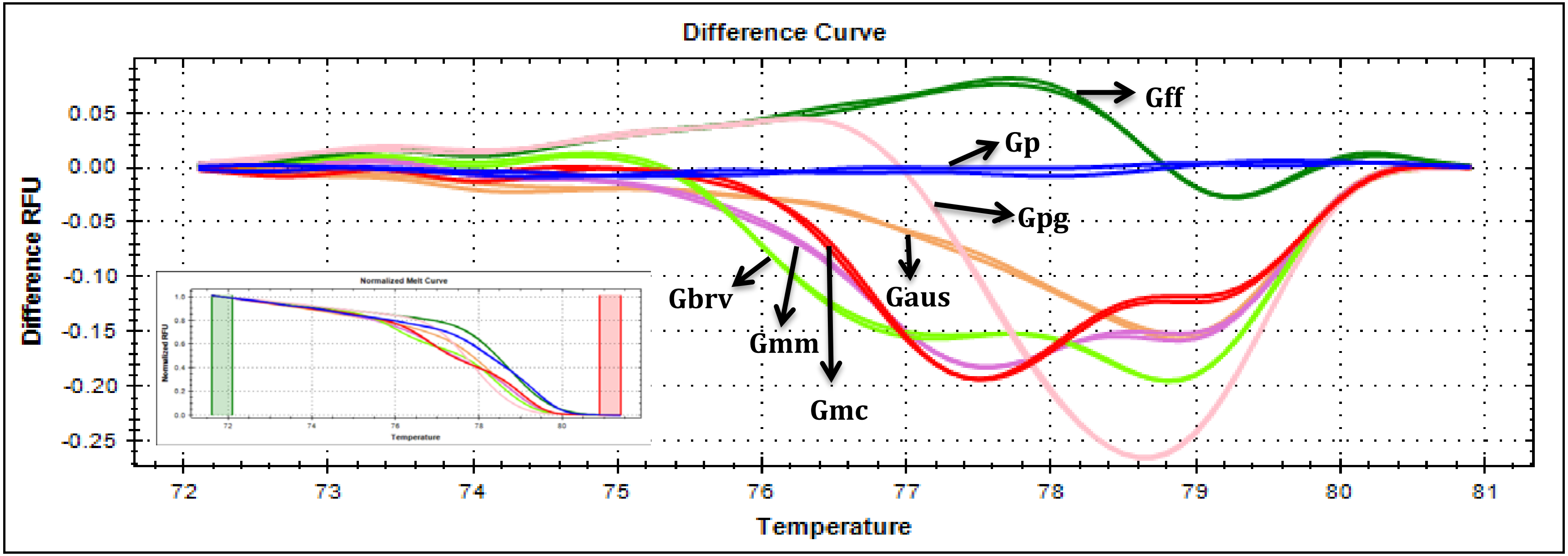

### Supplemental Figure 3

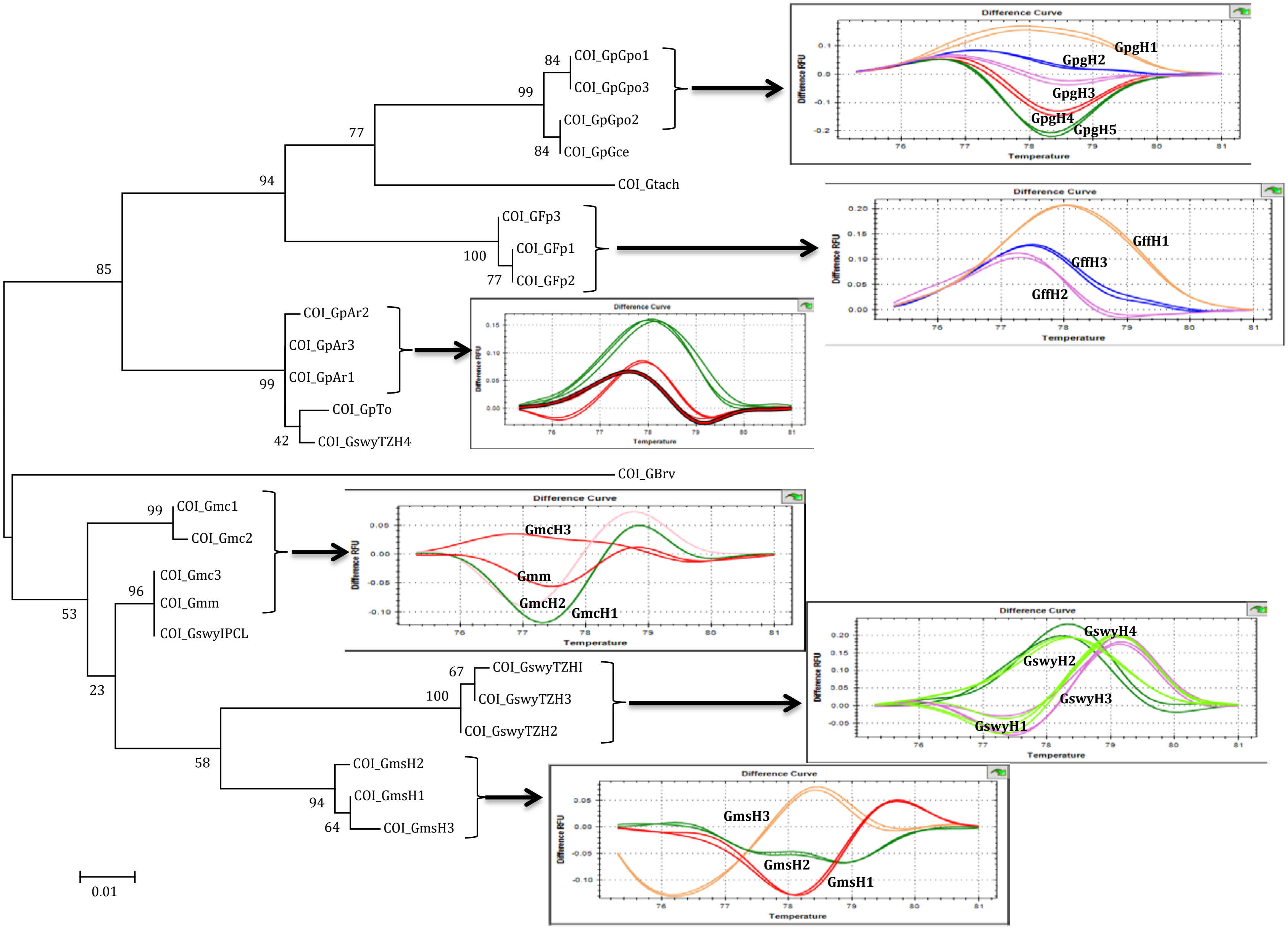

### Supplemental Figure 4 - X-Linkage of Male and Female Biased Genes

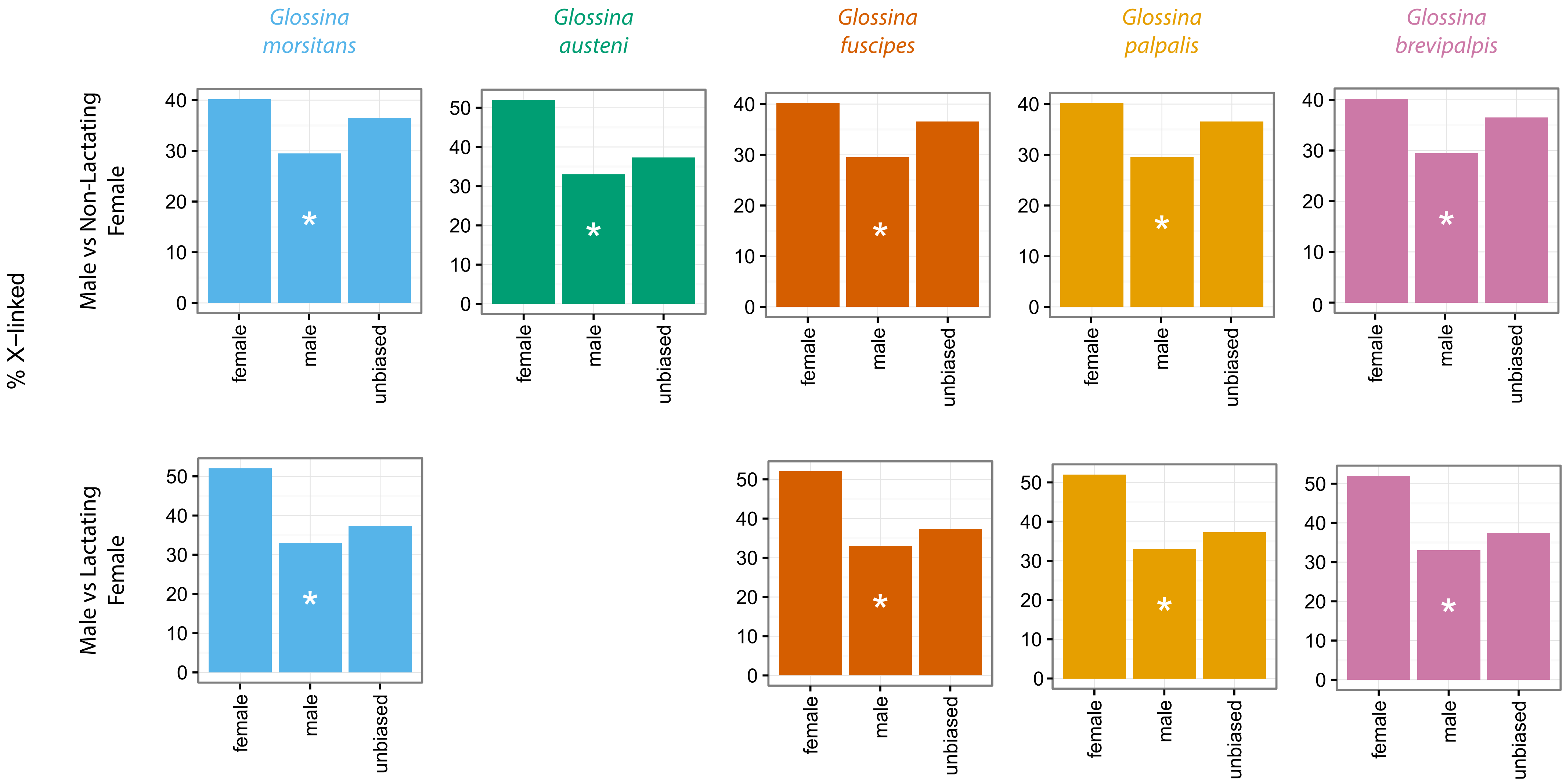

### Supplemental Figure 5 - Autosome-X Expression Levels

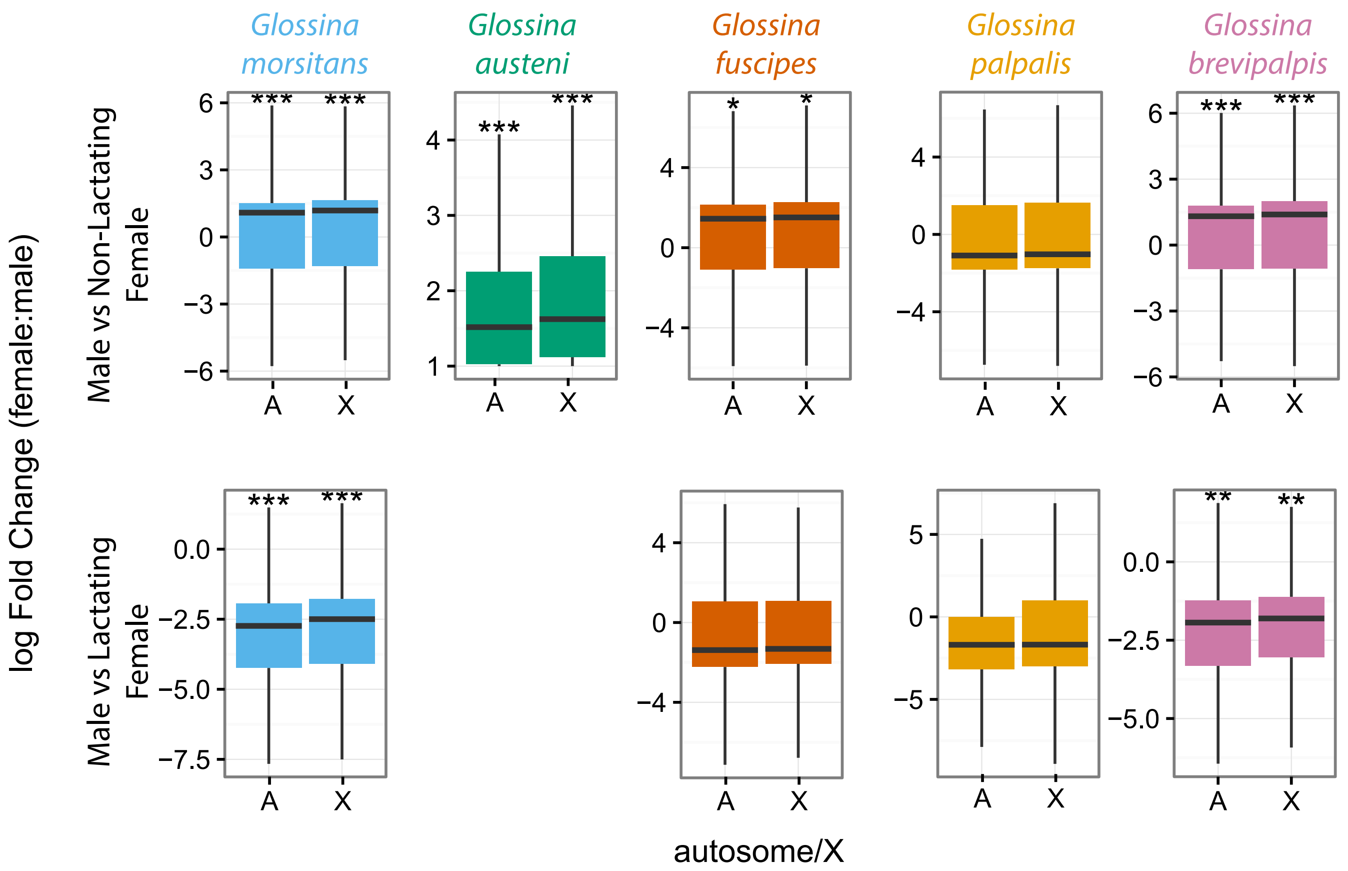

### Supplemental Figure 6 - dN-dS Autosome vs X

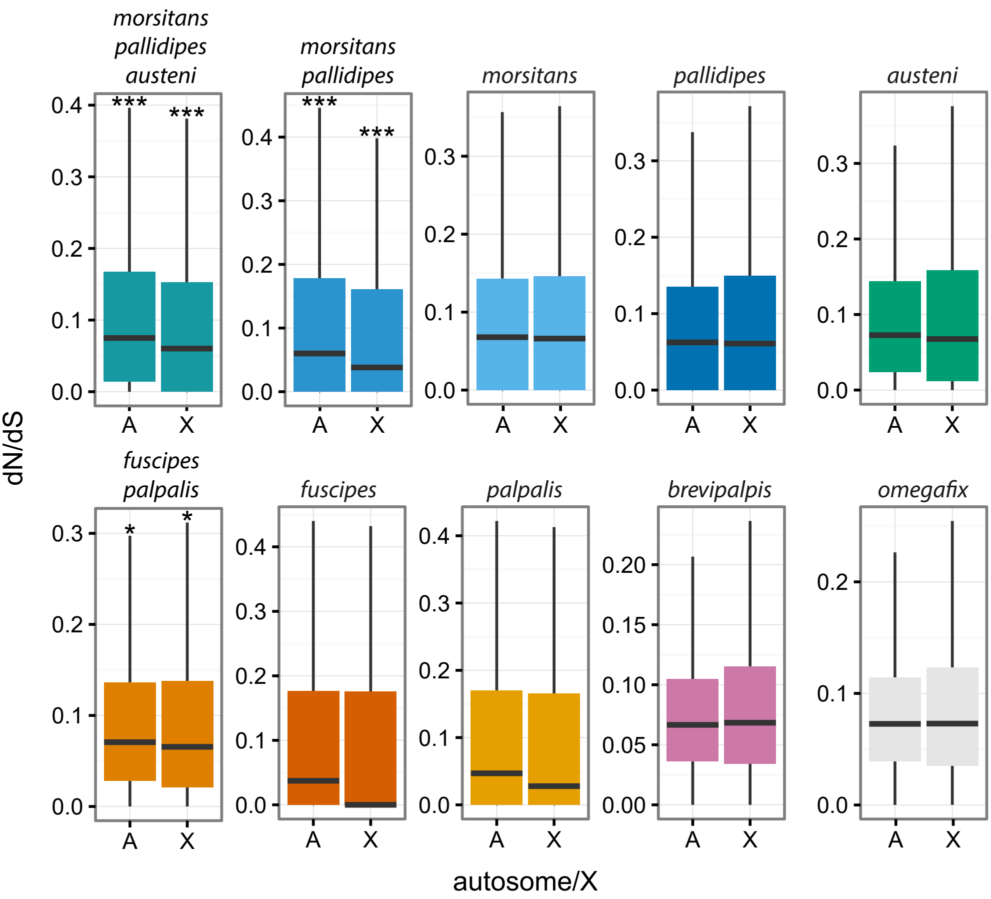

### Supplemental Figure 7_dnds_sexbias_XA_scaff

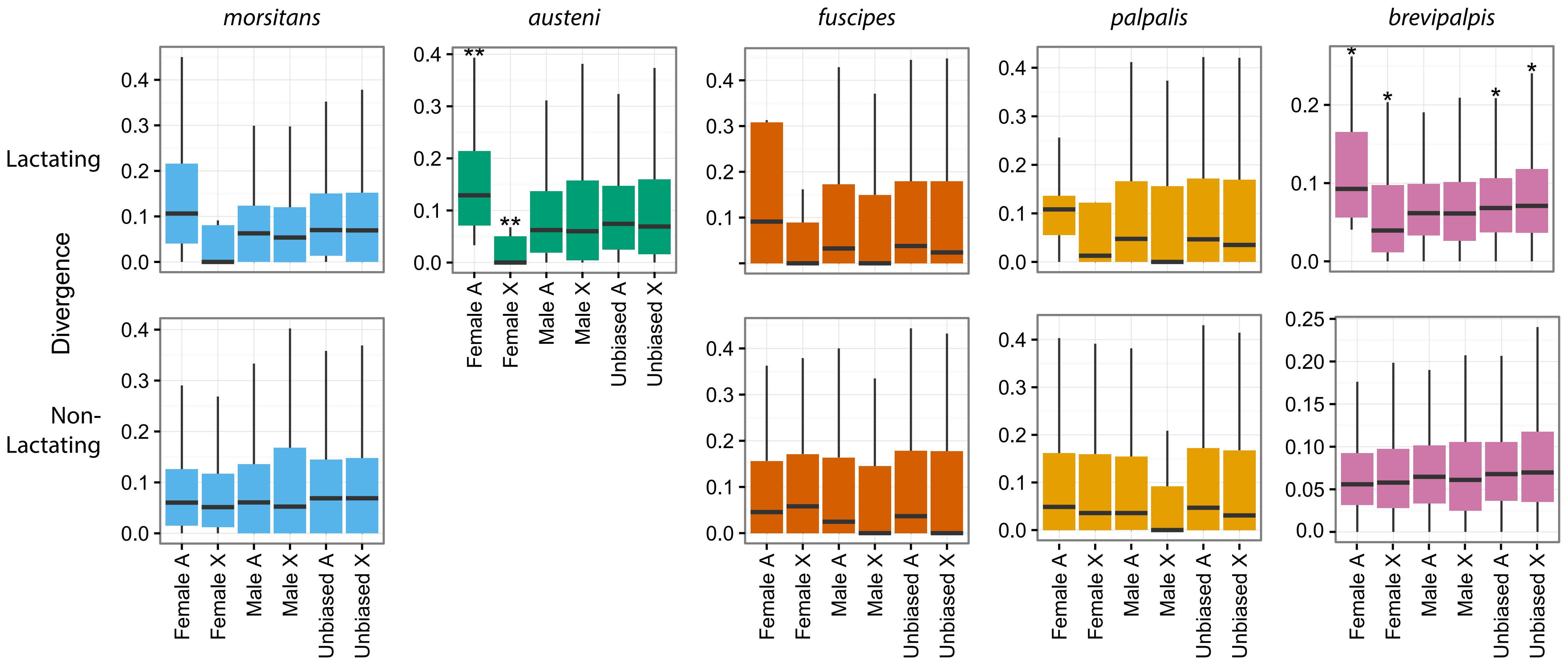

### Supplemental Figure 8 - Male Biased Expression per Muller Element

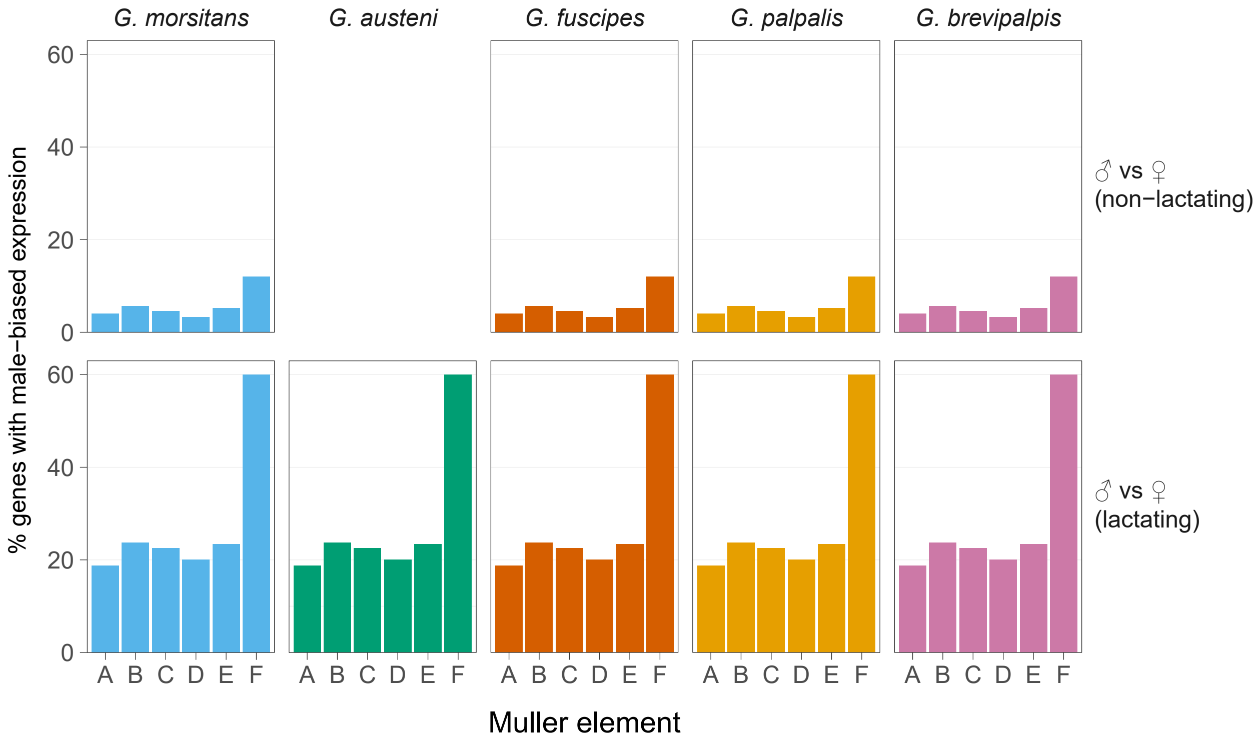
