## Supplementary figures and images for "The *Glossina* Genome Cluster: Comparative Genomic Analysis of the Vectors of African Trypanosomes"

### GAUT000053-PA_Homeobox_fwd.png

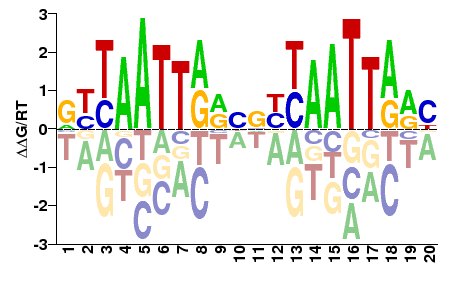

### GAUT000053-PA_Homeobox_rev.png

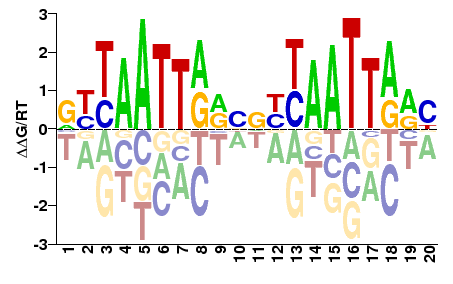

### GAUT000511-PA_zf-C2H2_fwd.png

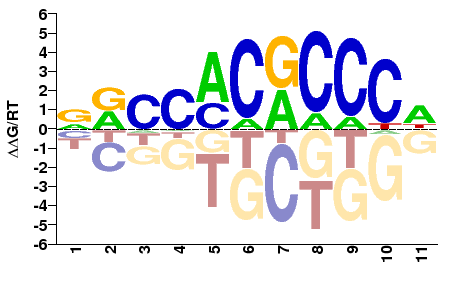

### GAUT000511-PA_zf-C2H2_rev.png

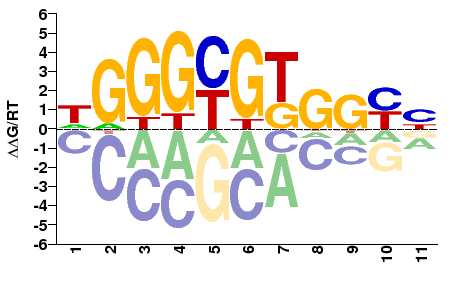

### GAUT000631-PA_CUT,Homeobox_fwd.png

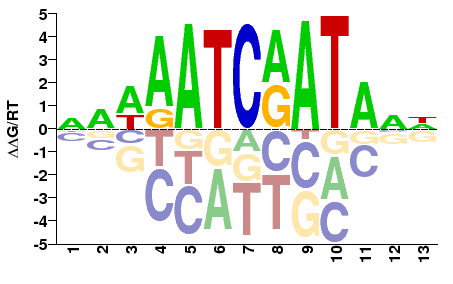

### GAUT000631-PA_CUT,Homeobox_rev.png

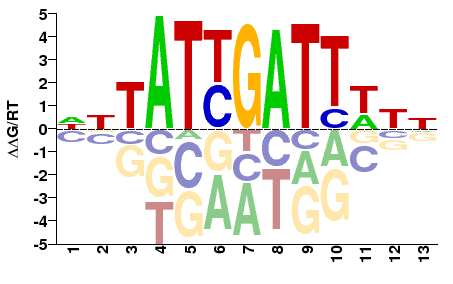

### GAUT001639-PA_Homeobox_fwd.png

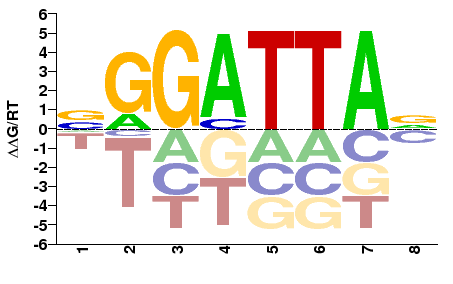

### GAUT001639-PA_Homeobox_rev.png

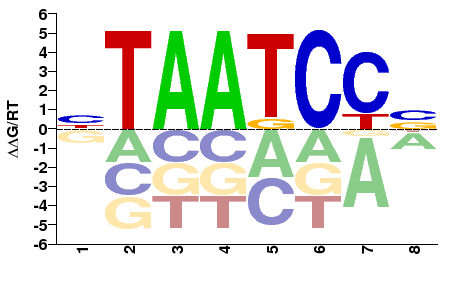

### GAUT001653-PA_Myb_DNA-binding_fwd.png

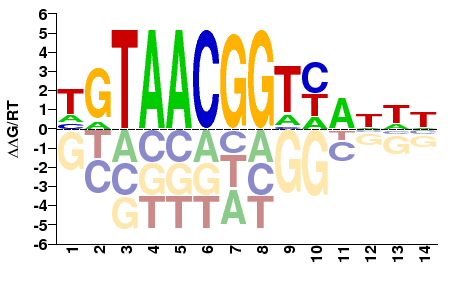

### GAUT001653-PA_Myb_DNA-binding_rev.png

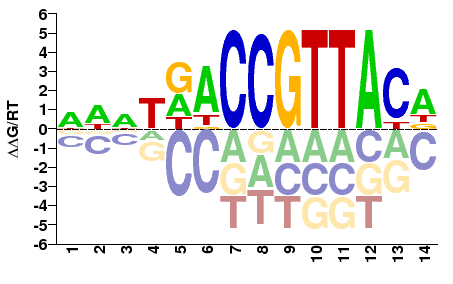

### GAUT001674-PA_HLH_fwd.png

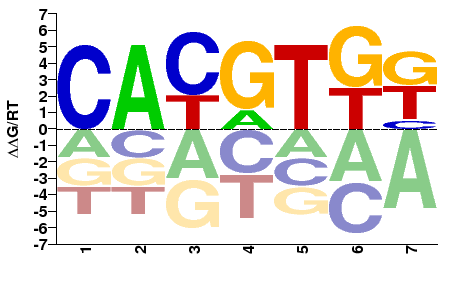

### GAUT001674-PA_HLH_rev.png

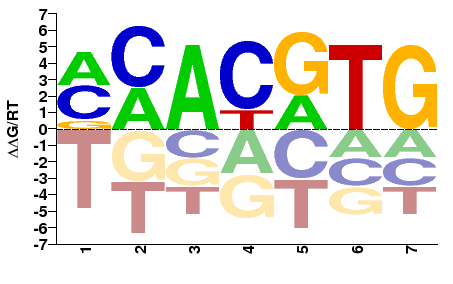

### GAUT001844-PA_zf-C2H2_fwd.png

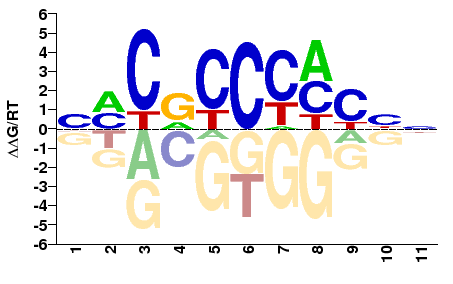

### GAUT001844-PA_zf-C2H2_rev.png

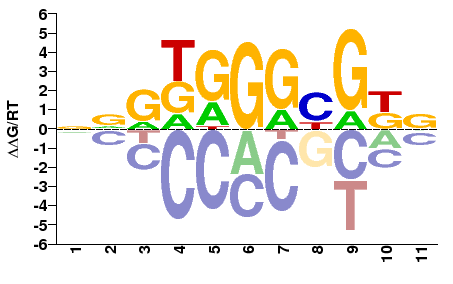

### GAUT001887-PA_zf-C2H2_fwd.png

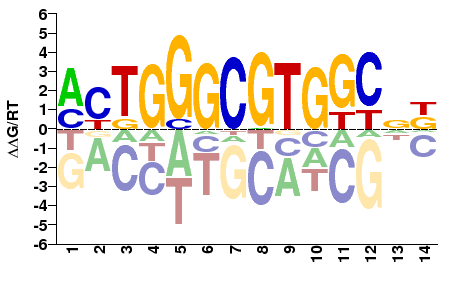

### GAUT001887-PA_zf-C2H2_rev.png

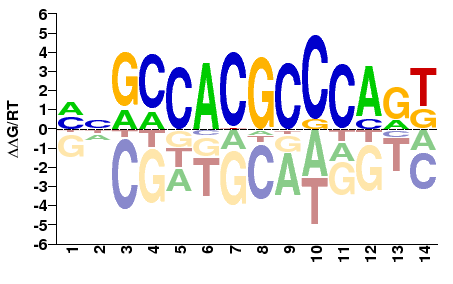

### GAUT002383-PA_Homeobox_fwd.png

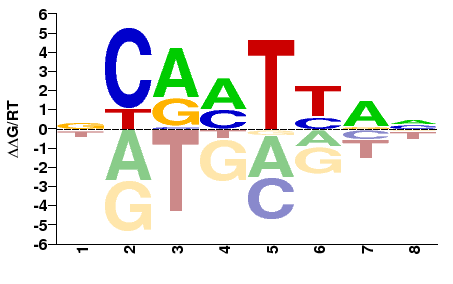

### GAUT002383-PA_Homeobox_rev.png

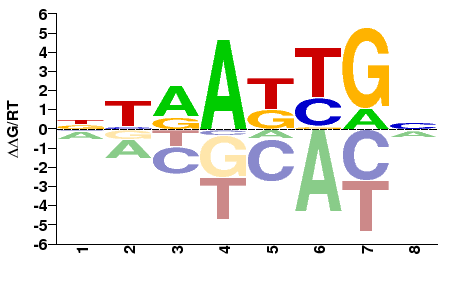

### GAUT002395-PA_Homeobox_fwd.png

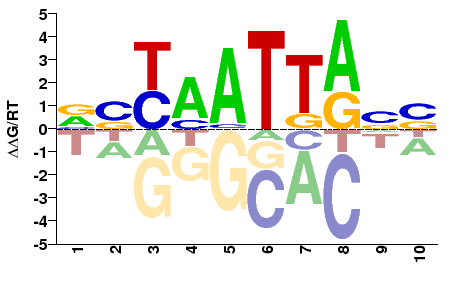

### GAUT002395-PA_Homeobox_rev.png

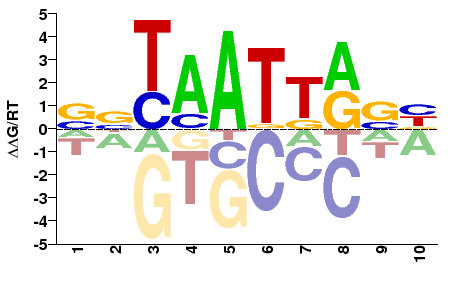

### GAUT002640-PA_bZIP_1_fwd.png

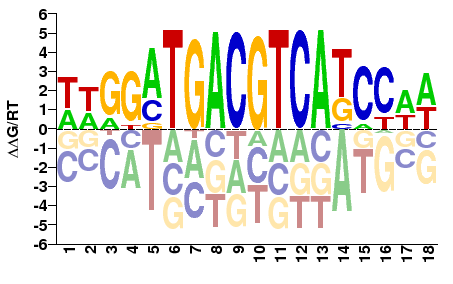

### GAUT002640-PA_bZIP_1_rev.png

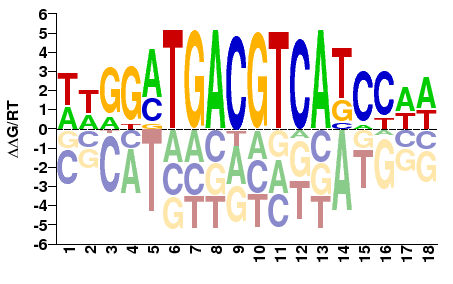

### GAUT002826-PA_GATA_fwd.png

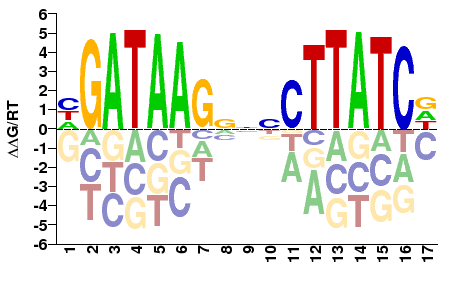

### GAUT002826-PA_GATA_rev.png

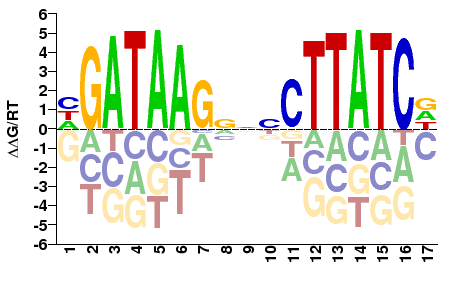

### GAUT002835-PA_GATA_fwd.png

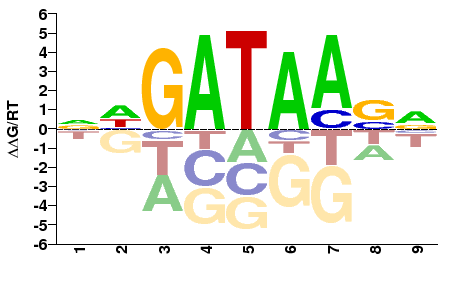

### GAUT002835-PA_GATA_rev.png

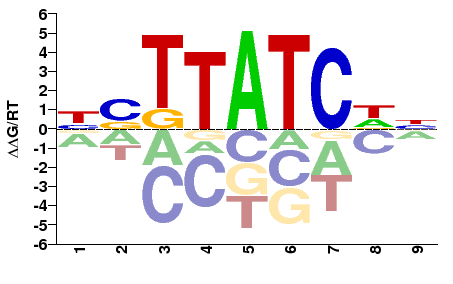

### GAUT003221-PA_PAX_fwd.png

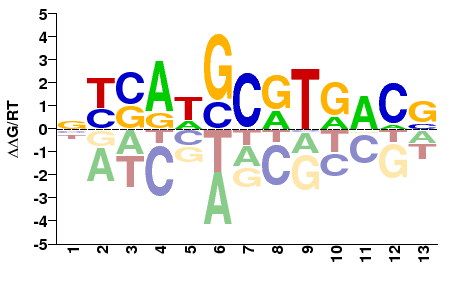

### GAUT003221-PA_PAX_rev.png

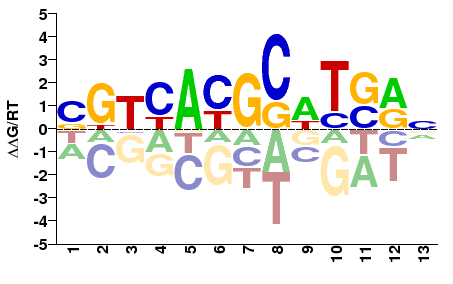

### GAUT003304-PA_Homeobox_fwd.png

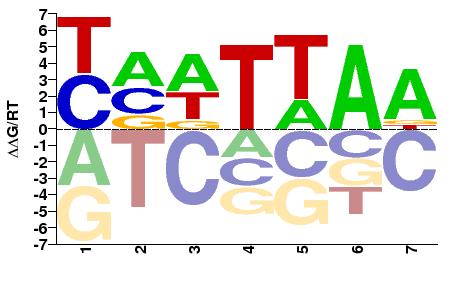

### GAUT003304-PA_Homeobox_rev.png

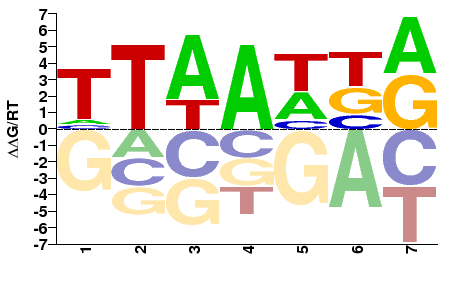

### GAUT003321-PA_Homeobox_fwd.png

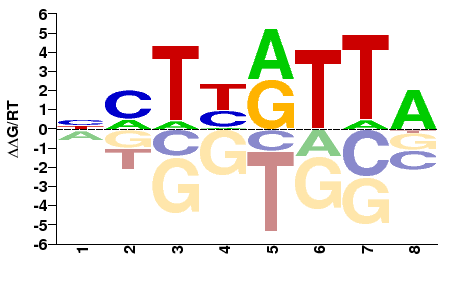

### GAUT003321-PA_Homeobox_rev.png

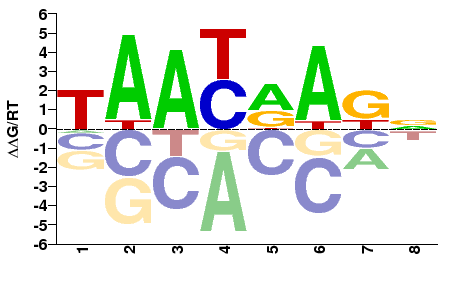

### GAUT003322-PA_Homeobox_fwd.png

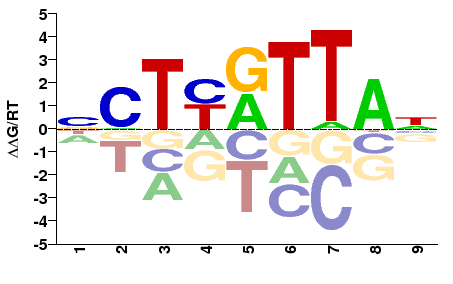

### GAUT003322-PA_Homeobox_rev.png

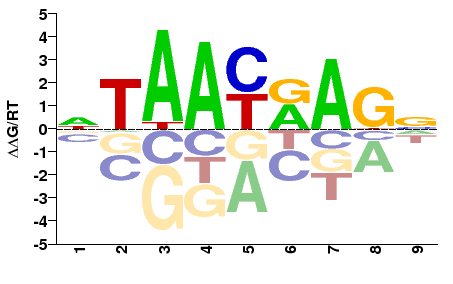

### GAUT004117-PA_zf-C4_fwd.png

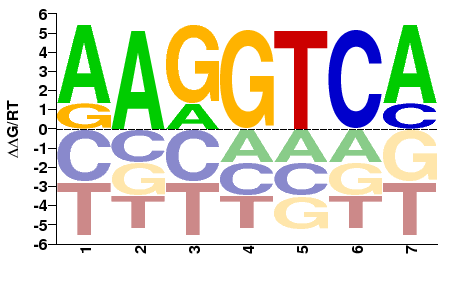

### GAUT004117-PA_zf-C4_rev.png

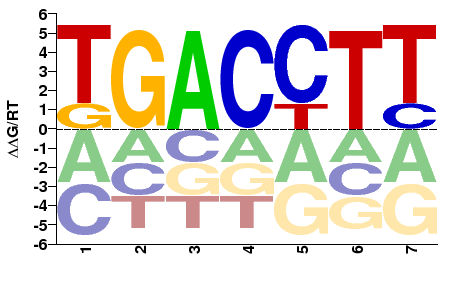

### GAUT004707-PA_LAG1-DNAbind_fwd.png

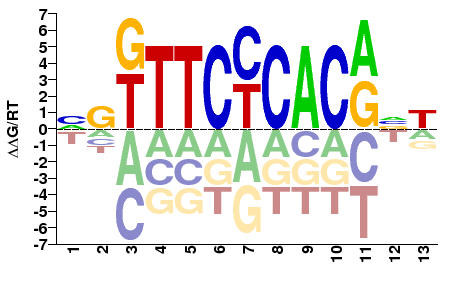

### GAUT004707-PA_LAG1-DNAbind_rev.png

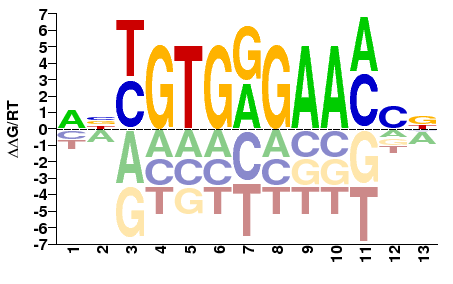

### GAUT004881-PA_zf-C4_fwd.png

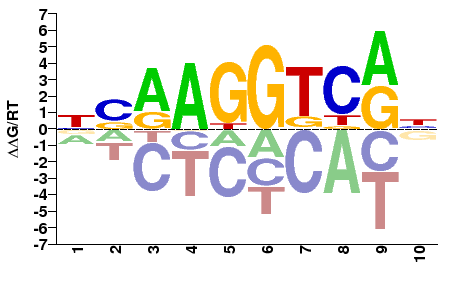

### GAUT004881-PA_zf-C4_rev.png

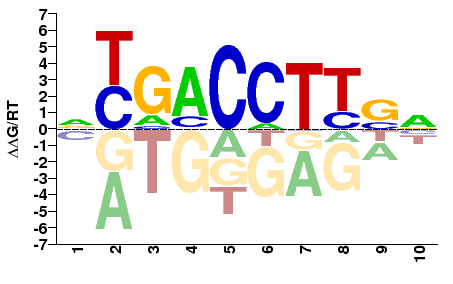

### GAUT006620-PA_Forkhead_fwd.png

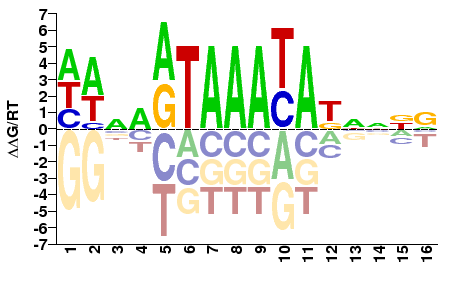

### GAUT006620-PA_Forkhead_rev.png

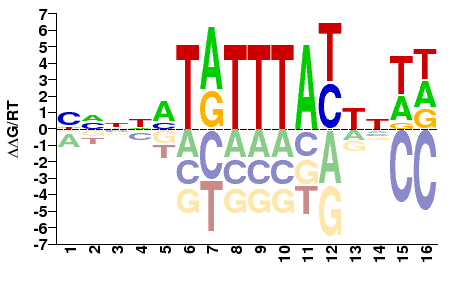

### GAUT006952-PA_Myb_DNA-binding_fwd.png

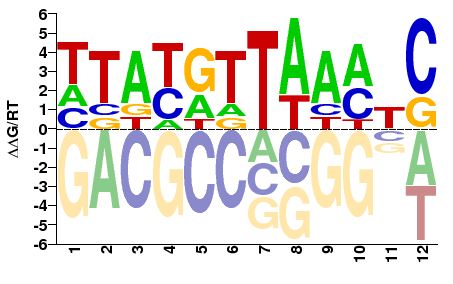

### GAUT006952-PA_Myb_DNA-binding_rev.png

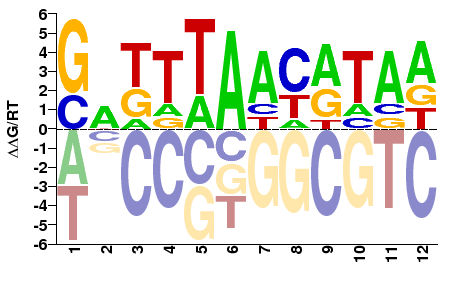

### GAUT007108-PA_zf-C2H2_fwd.png

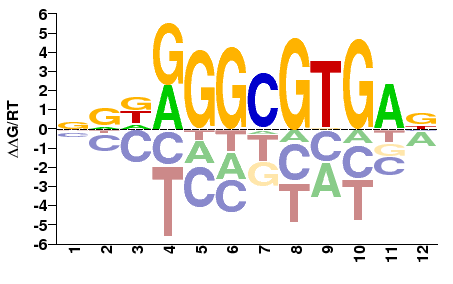

### GAUT007108-PA_zf-C2H2_rev.png

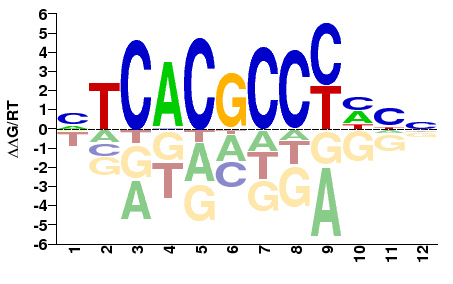

### GAUT007221-PA_MADF_DNA_bdg_fwd.png

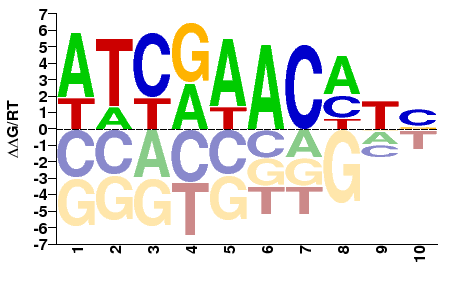

### GAUT007221-PA_MADF_DNA_bdg_rev.png

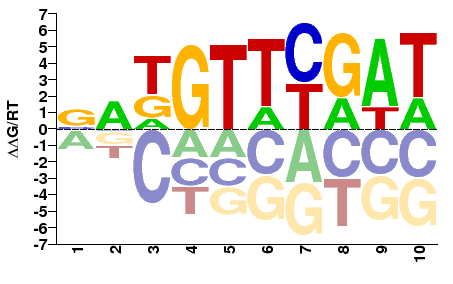

### GAUT007361-PA_Homeobox_fwd.png

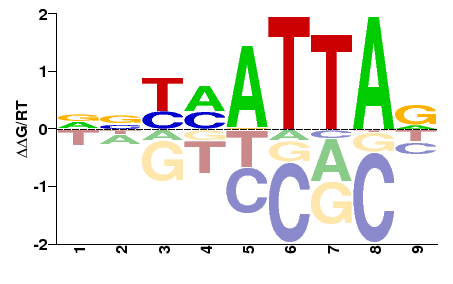

### GAUT007361-PA_Homeobox_rev.png

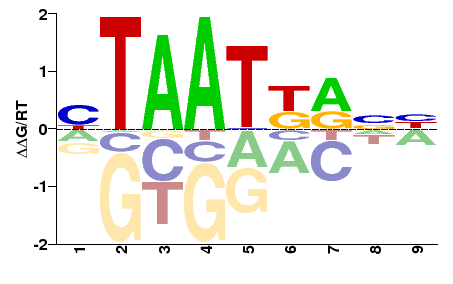

### GAUT008149-PA_HTH_psq_fwd.png

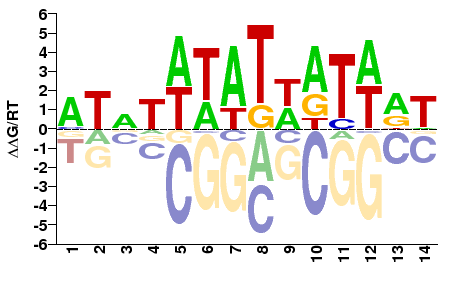

### GAUT008149-PA_HTH_psq_rev.png

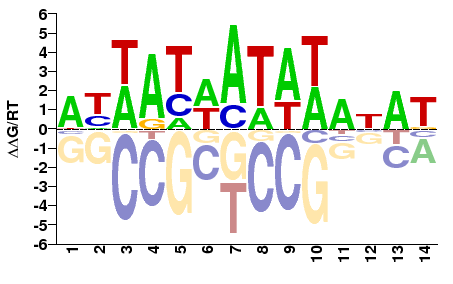

### GAUT008304-PA_zf-C2H2_fwd.png

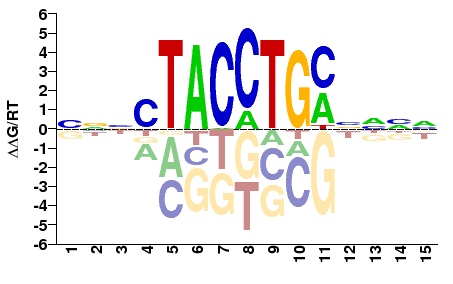

### GAUT008304-PA_zf-C2H2_rev.png

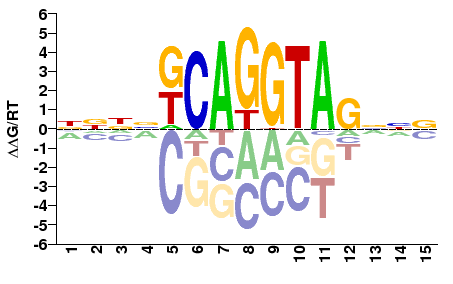

### GAUT008381-PA_HLH_fwd.png

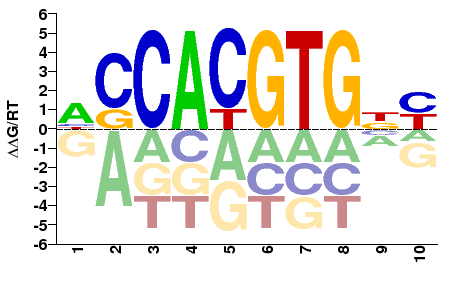

### GAUT008381-PA_HLH_rev.png

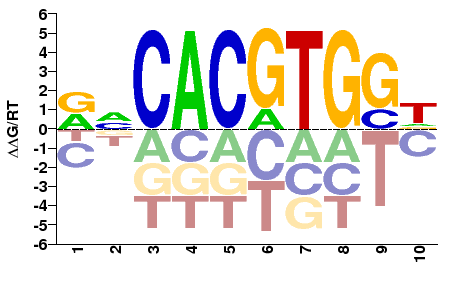

### GAUT008741-PA_HLH_fwd.png

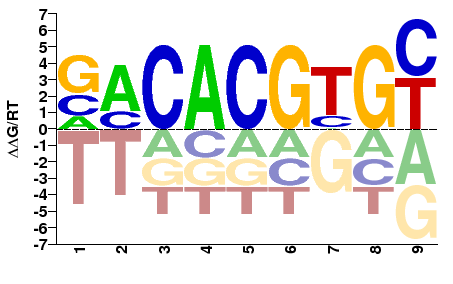

### GAUT008741-PA_HLH_rev.png

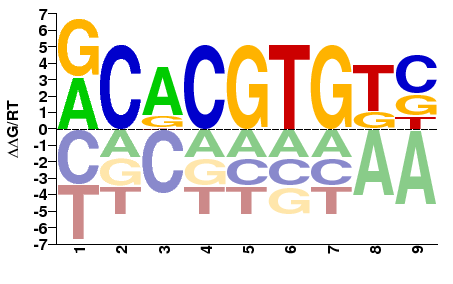

### GAUT008843-PA_Ets_fwd.png

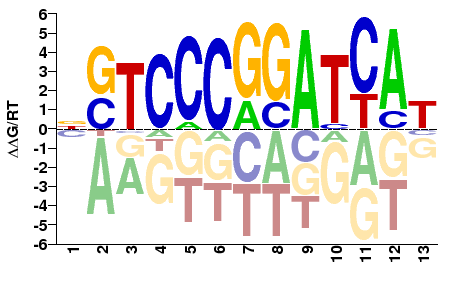

### GAUT008843-PA_Ets_rev.png

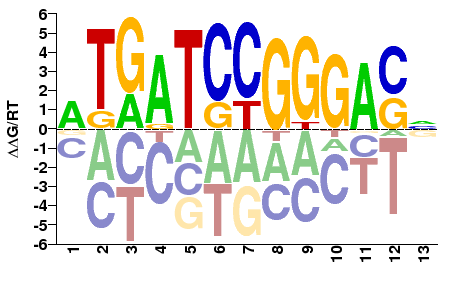

### GAUT009195-PA_Homeobox_fwd.png

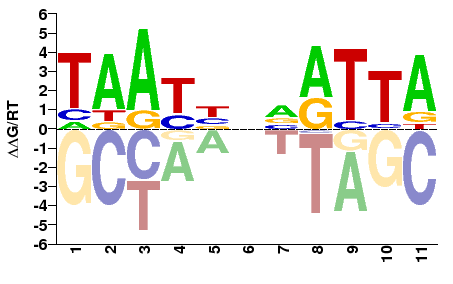

### GAUT009195-PA_Homeobox_rev.png

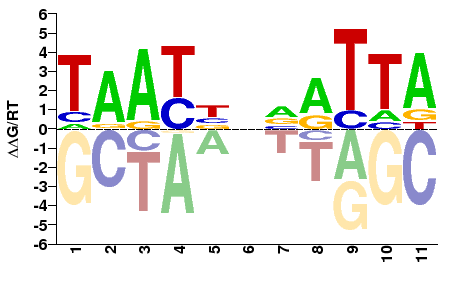

### GAUT009346-PA_HLH_fwd.png

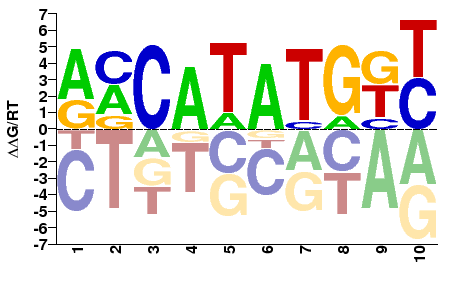

### GAUT009346-PA_HLH_rev.png

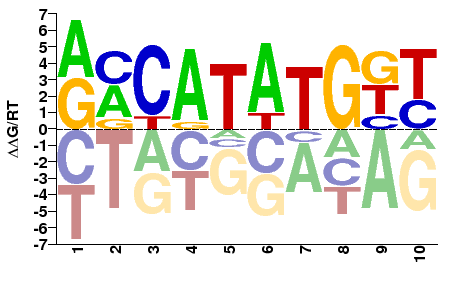

### GAUT009458-PA_zf-C2H2_fwd.png

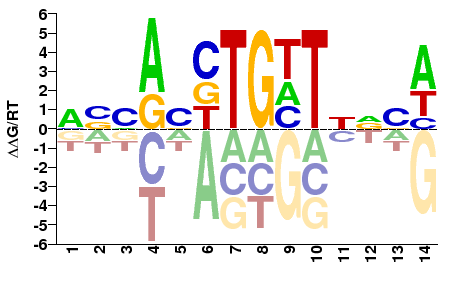

### GAUT009458-PA_zf-C2H2_rev.png

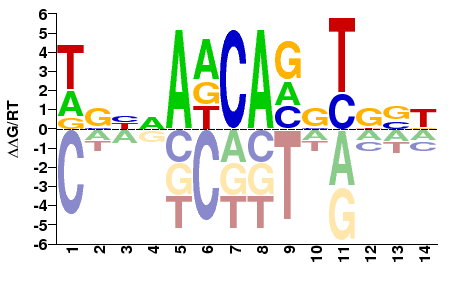

### GAUT009466-PA_zf-C2H2_fwd.png

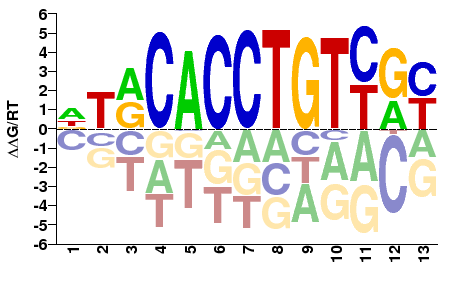

### GAUT009466-PA_zf-C2H2_rev.png

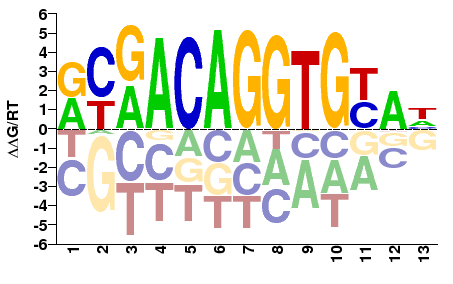

### GAUT009489-PA_zf-C2H2_fwd.png

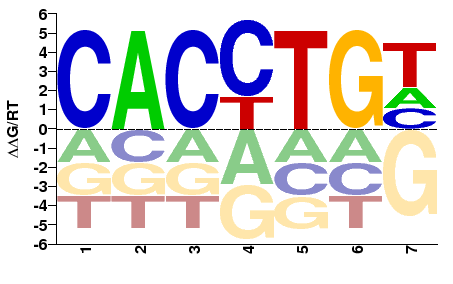

### GAUT009489-PA_zf-C2H2_rev.png

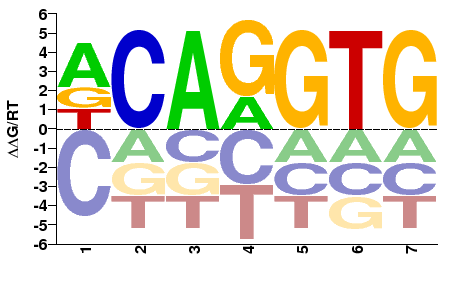

### GAUT009726-PA_HLH_fwd.png

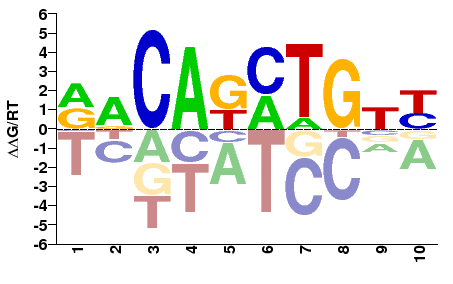

### GAUT009726-PA_HLH_rev.png

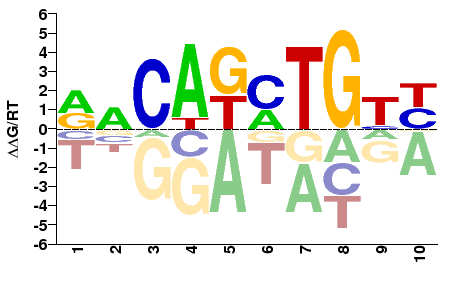

### GAUT010135-PA_HLH_fwd.png

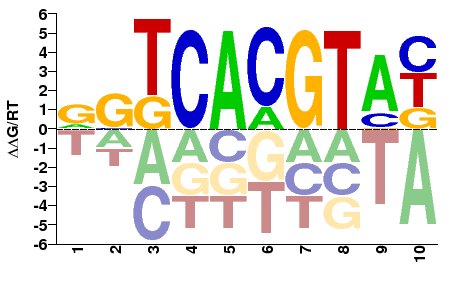

### GAUT010135-PA_HLH_rev.png

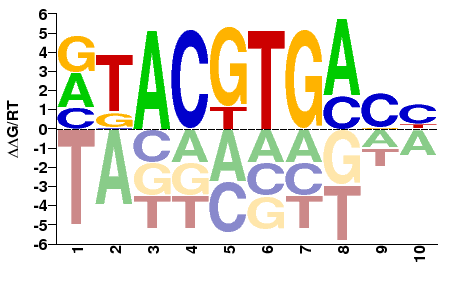

### GAUT010297-PA_zf-C2H2_fwd.png

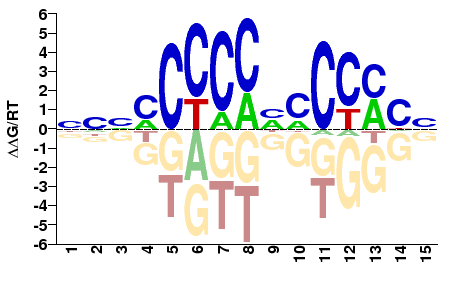

### GAUT010297-PA_zf-C2H2_rev.png

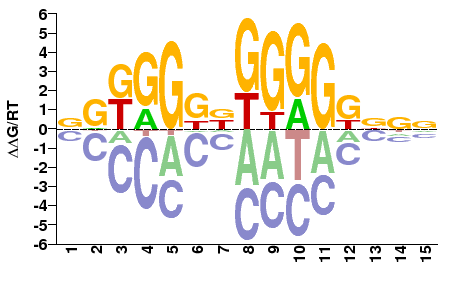

### GAUT010429-PA_Homeobox_fwd.png

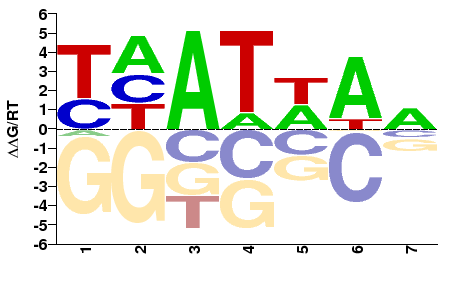

### GAUT010429-PA_Homeobox_rev.png

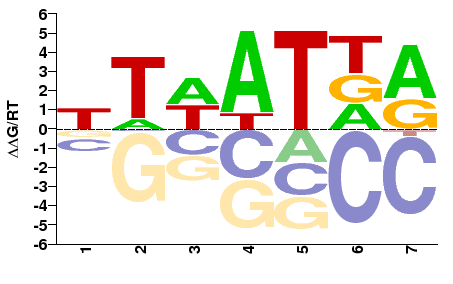

### GAUT010489-PA_Forkhead_fwd.png

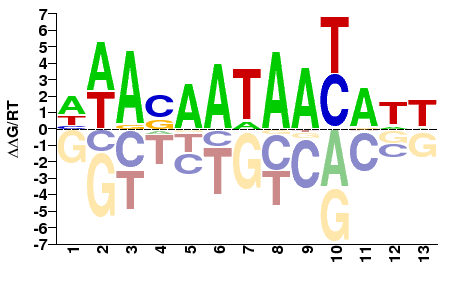

### GAUT010489-PA_Forkhead_rev.png

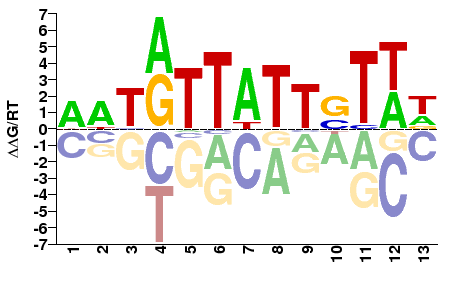

### GAUT010640-PA_AT_hook_fwd.png

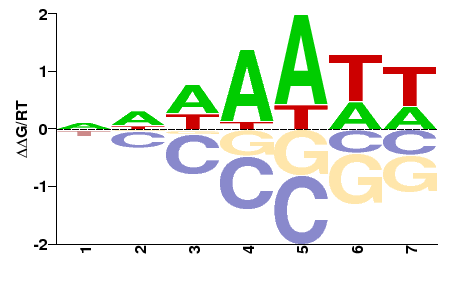

### GAUT010640-PA_AT_hook_rev.png

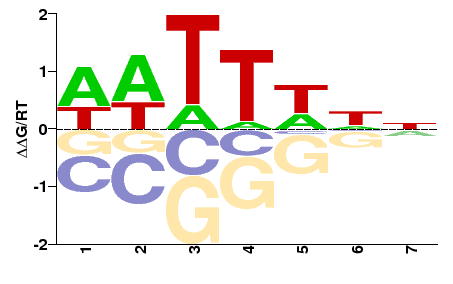

### GAUT011069-PA_zf-C4_fwd.png

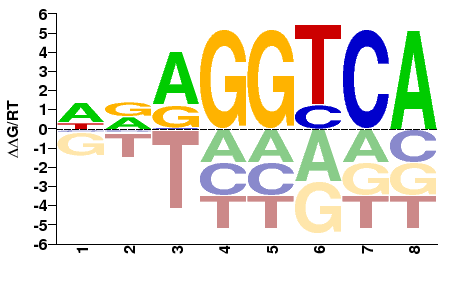

### GAUT011069-PA_zf-C4_rev.png

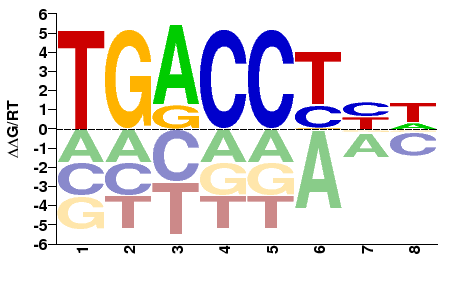

### GAUT011215-PA_CUT,Homeobox_fwd.png

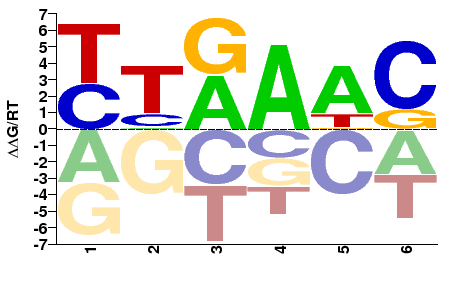

### GAUT011215-PA_CUT,Homeobox_rev.png

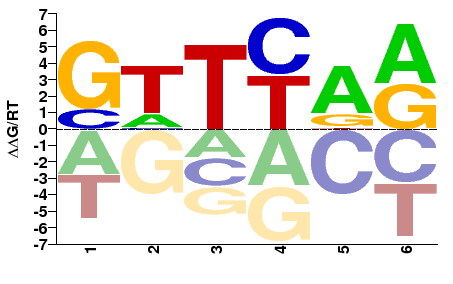

### GAUT011248-PA_Homeobox_fwd.png

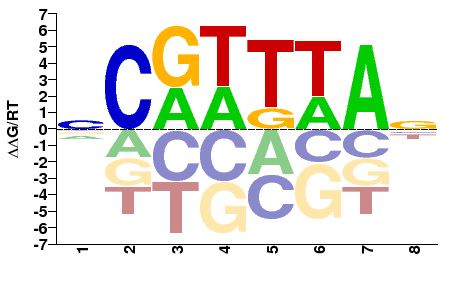

### GAUT011248-PA_Homeobox_rev.png

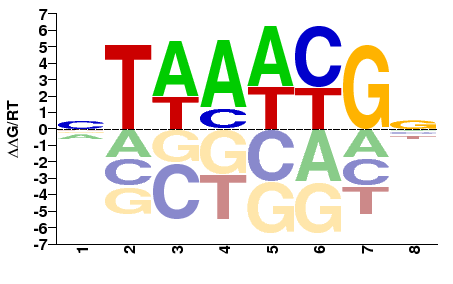

### GAUT011414-PA_PAX_fwd.png

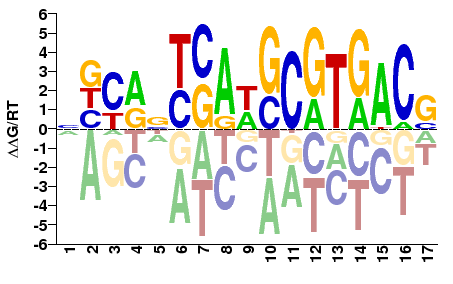

### GAUT011414-PA_PAX_rev.png

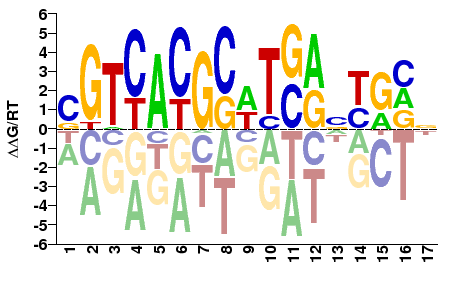

### GAUT011573-PA_Homeobox_fwd.png

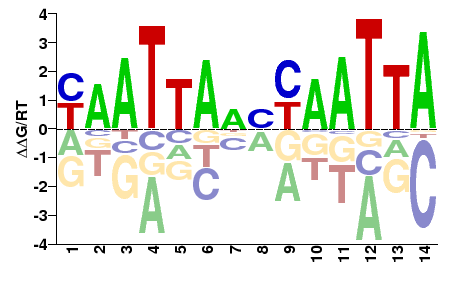

### GAUT011573-PA_Homeobox_rev.png

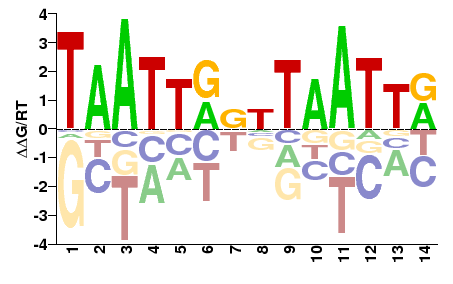

### GAUT011656-PA_zf-C2H2_fwd.png

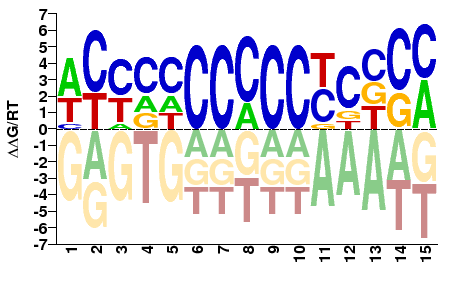

### GAUT011656-PA_zf-C2H2_rev.png

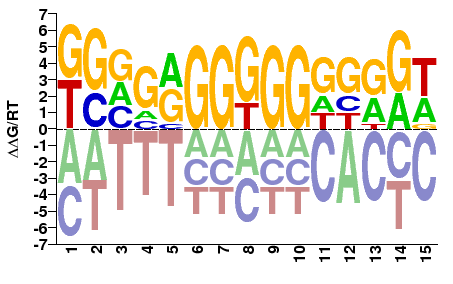

### GAUT011834-PA_bZIP_1_fwd.png

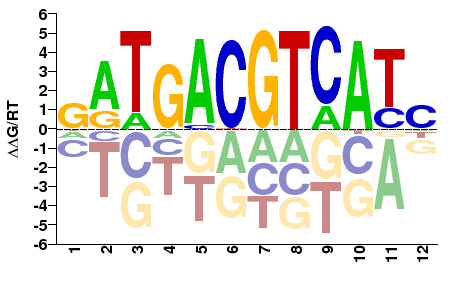

### GAUT011834-PA_bZIP_1_rev.png

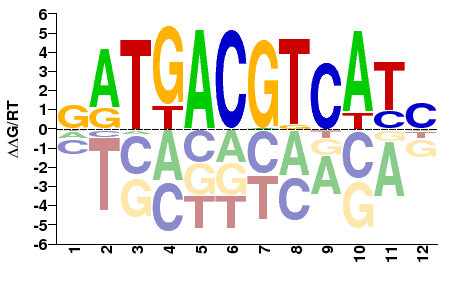

### GAUT011960-PA_Homeobox,Pou_fwd.png

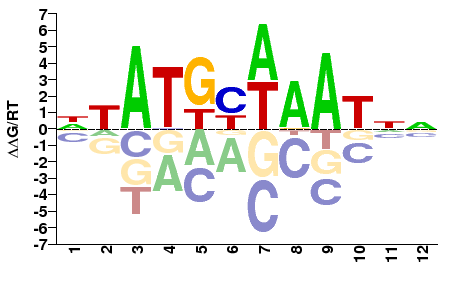

### GAUT011960-PA_Homeobox,Pou_rev.png

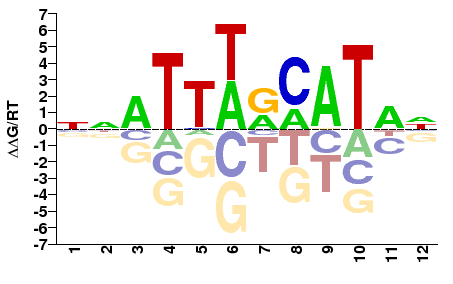

### GAUT012010-PA_Homeobox_fwd.png

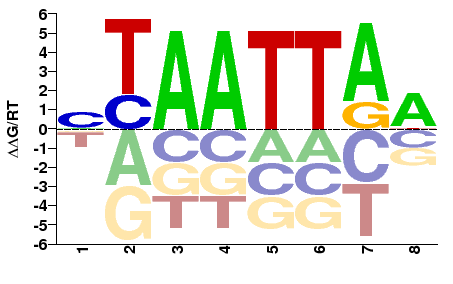

### GAUT012010-PA_Homeobox_rev.png

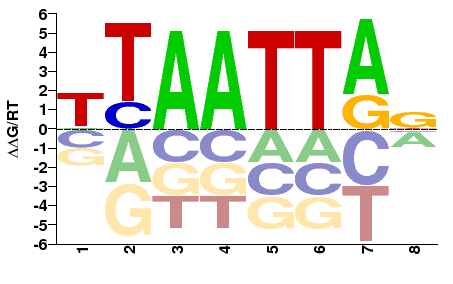
