## Supplemental Figure 1 for "The *Glossina* Genome Cluster: Comparative Genomic Analysis of the Vectors of African Trypanosomes"

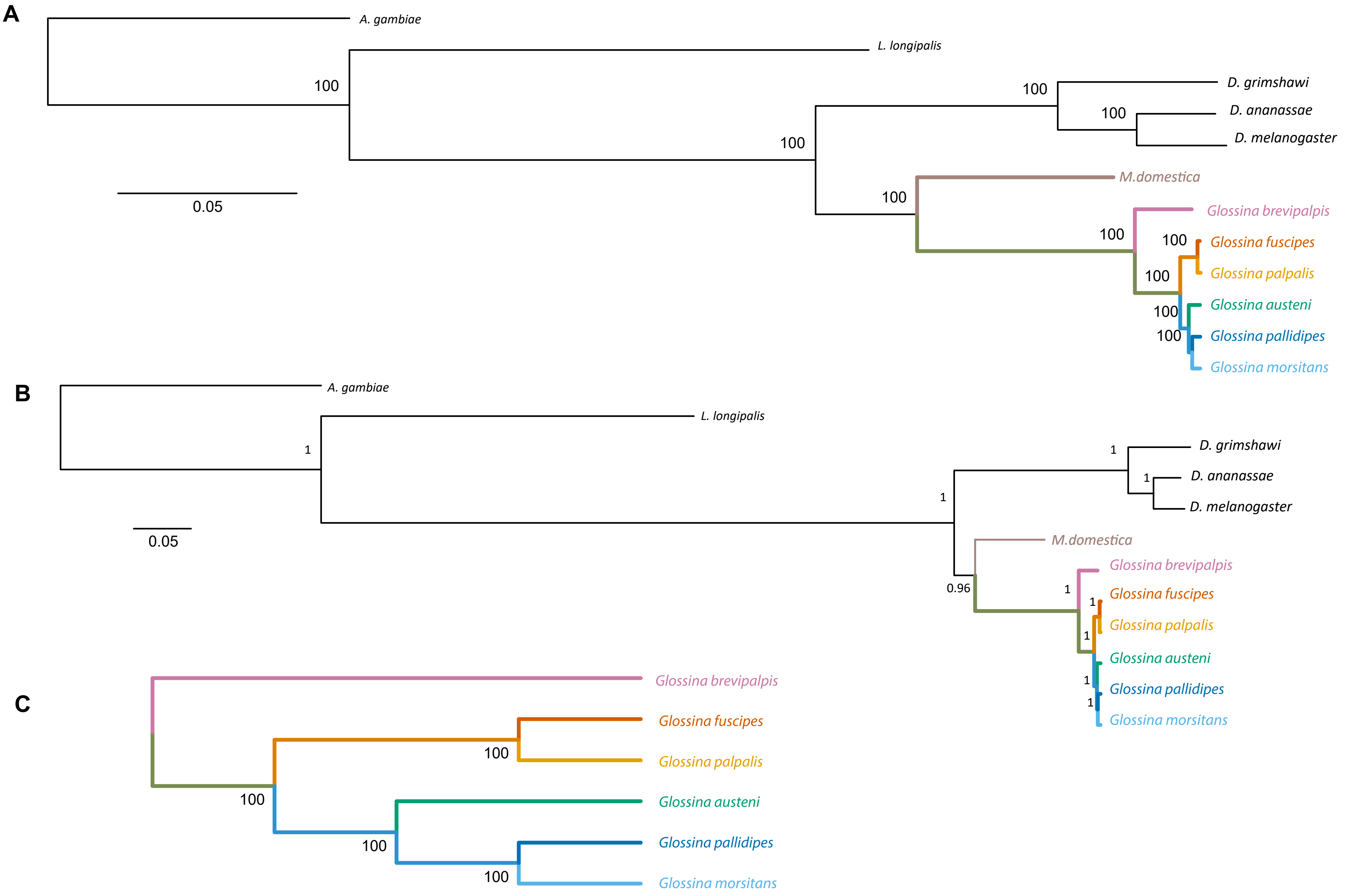


**Suppl. Figure 1.** A: Maximum likelihood tree inferred by Raxml on a concatenated alignment of 117,783 amino acids from 286 genes and employing the LG+G+F model of replacement. Value at nodes are bootstrap supports from the consensus of 100 pseudoreplicates in Raxml. B: Bayesian phylogentic tree inferred with Phylobayes 3 using CAT+G replacment model, supports at nodes are posterior probabilities; C: Astral analysis based on 286 genes with bootstrap support.
