## Supplementary figures and images for "The *Glossina* Genome Cluster: Comparative Genomic Analysis of the Vectors of African Trypanosomes"

### Supplemental Figure 9 - dN-dS per Muller Element

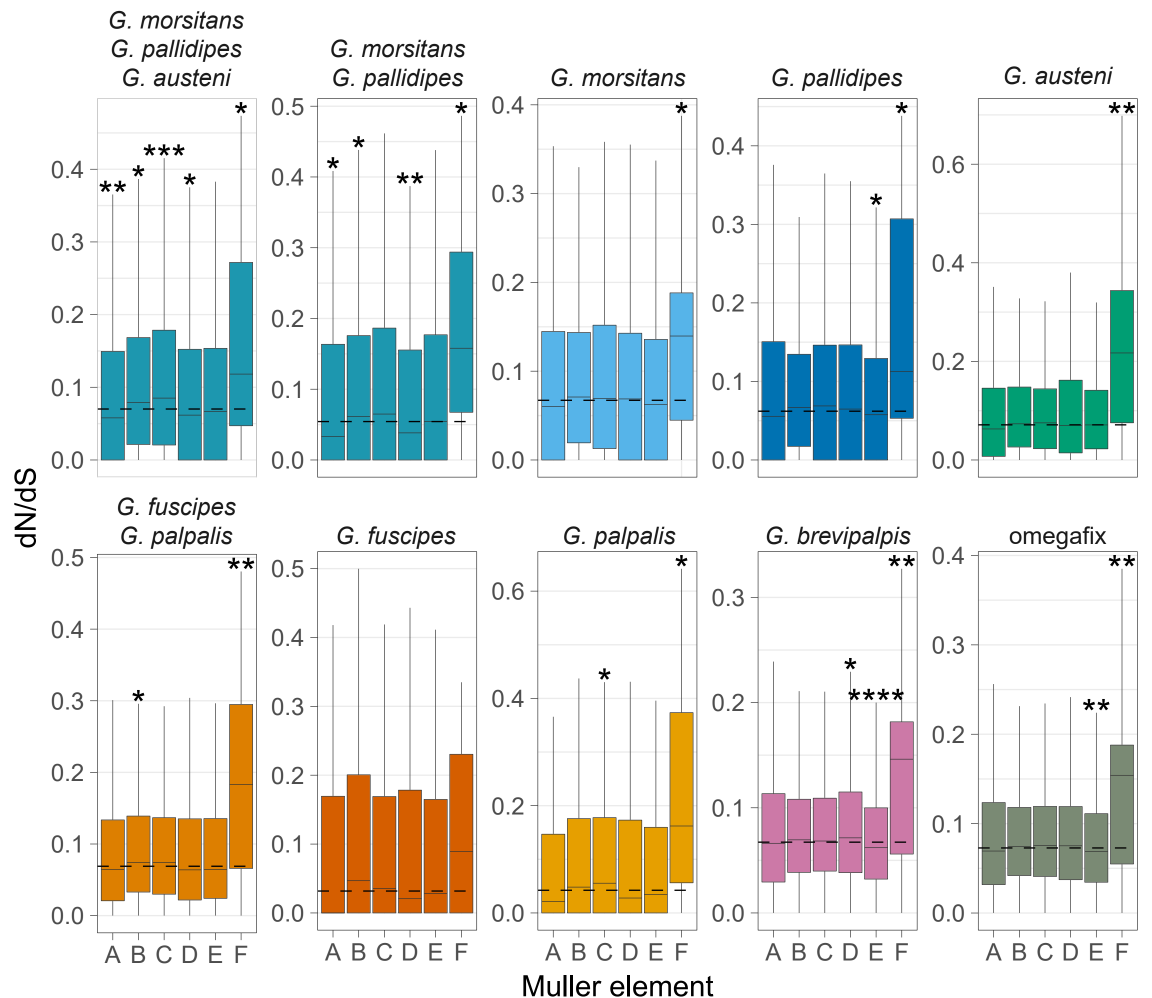

### Supplemental Figure 10 - Genomic Composition of Glossina Transposable Elements

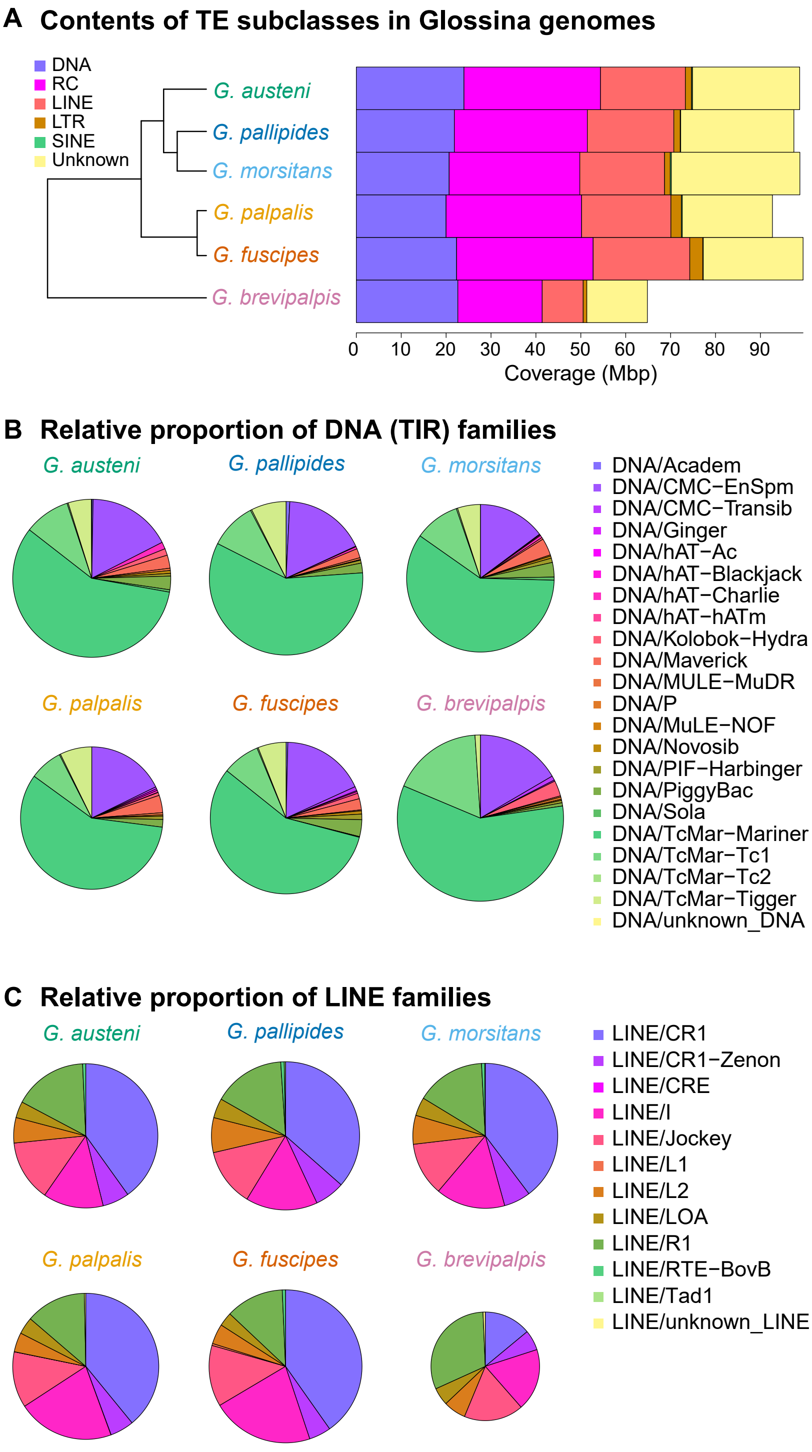

### Supplemental Figure 11 - Sub-Genus specific Gene Families PCA Plot Descriptions

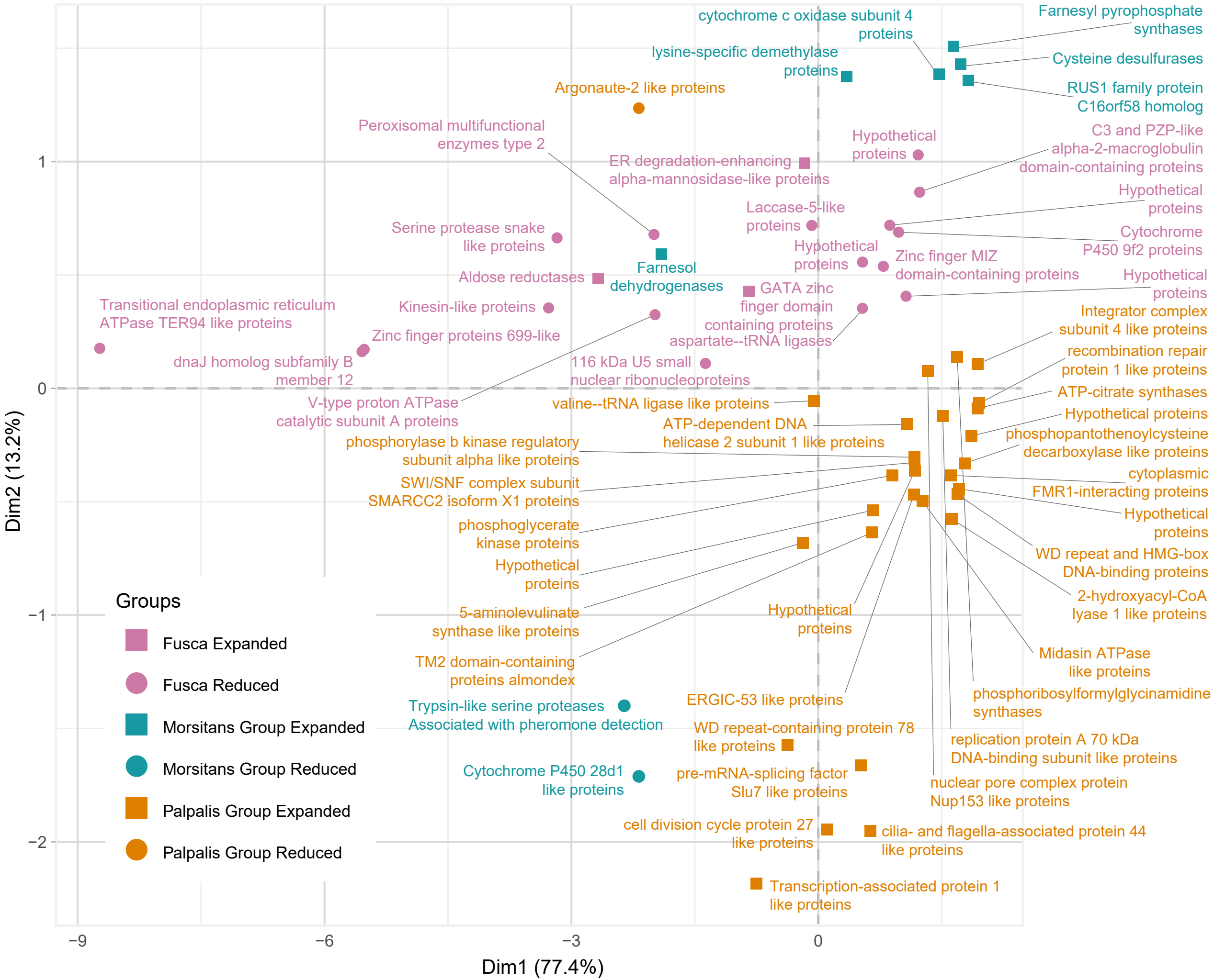

### Supplemental Figure 12 - Sub-Genus specific Gene Families PCA Plot Indentifiers

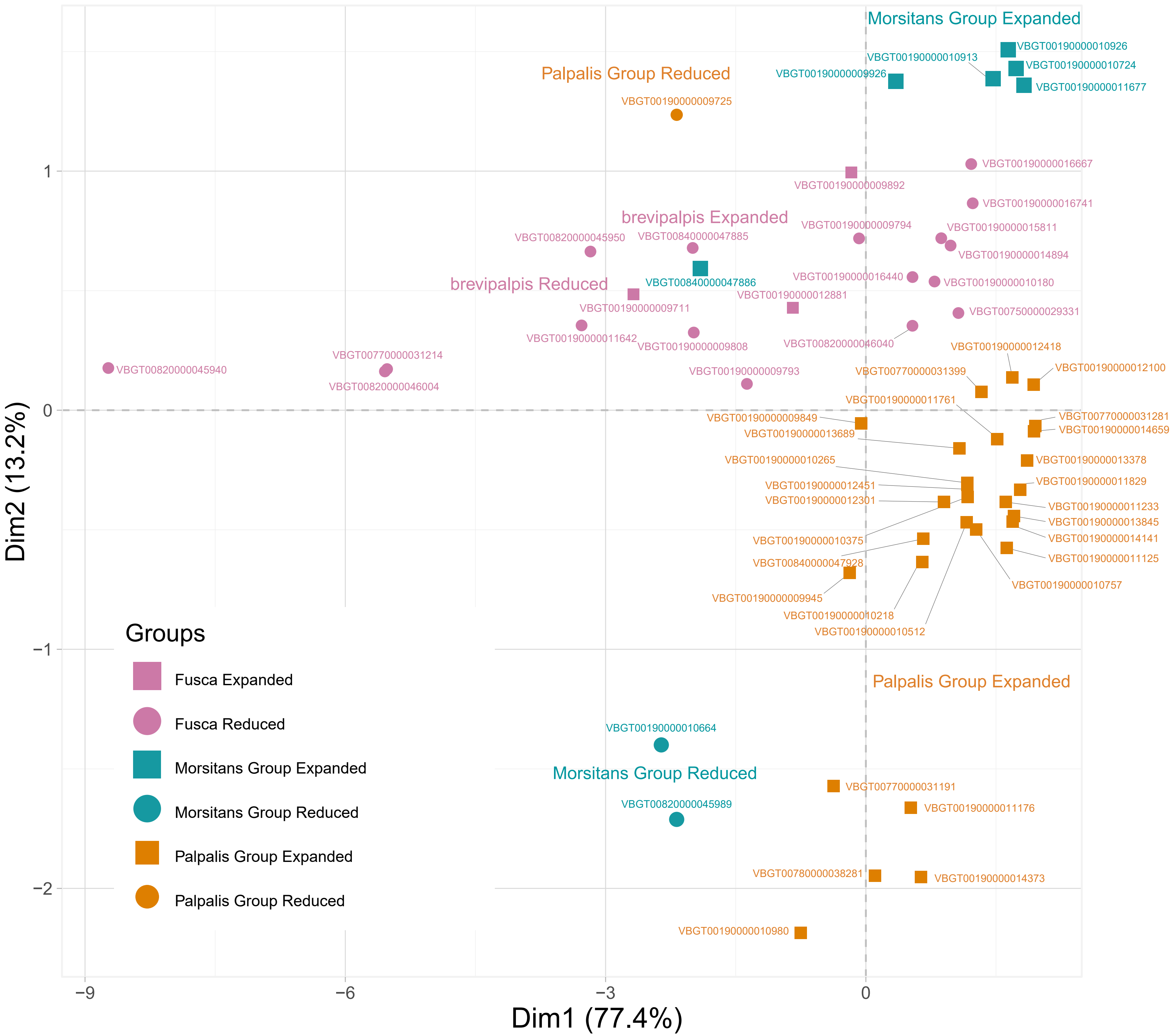

### Supplemental Figure 13 - Transcription Factor Constitution in Glossina

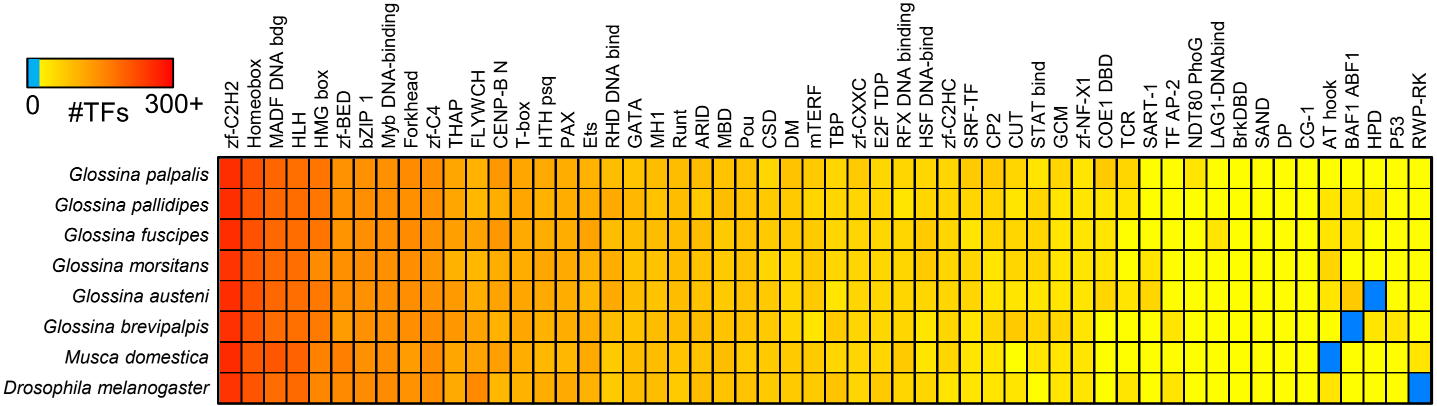
